# The gut metagenome harbors metabolic and antibiotic resistance signatures of moderate-to-severe asthma

**DOI:** 10.1101/2023.01.03.522677

**Authors:** Naomi G. Wilson, Ariel Hernandez-Leyva, Drew J. Schwartz, Leonard B. Bacharier, Andrew L. Kau

## Abstract

Asthma is a common allergic airway disease that develops in association with the human microbiome early in life. Both the composition and function of the infant gut microbiota have been linked to asthma risk, but functional alterations in the gut microbiota of older patients with established asthma remain an important knowledge gap. Here, we performed whole metagenomic shotgun sequencing of 95 stool samples from 59 healthy and 36 subjects with moderate-to-severe asthma to characterize the metagenomes of gut microbiota in children and adults 6 years and older. Mapping of functional orthologs revealed that asthma contributes to 2.9% of the variation in metagenomic content even when accounting for other important clinical demographics. Differential abundance analysis showed an enrichment of long-chain fatty acid (LCFA) metabolism pathways which have been previously implicated in airway smooth muscle and immune responses in asthma. We also observed increased richness of antibiotic resistance genes (ARGs) in people with asthma. One differentially abundant ARG was a macrolide resistance marker, *ermF*, which significantly co-occurred with the *Bacteroides fragilis* toxin, suggesting a possible relationship between enterotoxigenic *B. fragilis*, antibiotic resistance, and asthma. Lastly, we found multiple virulence factor (VF) and ARG pairs that co-occurred in both cohorts suggesting that virulence and antibiotic resistance traits are co-selected and maintained in the fecal microbiota of people with asthma. Overall, our results show functional alterations via LCFA biosynthetic genes and increases in antibiotic resistance genes in the gut microbiota of subjects with moderate-to-severe asthma and could have implications for asthma management and treatment.

## Background

Asthma is a common respiratory disease characterized by symptoms of airway obstruction including wheeze, cough, and shortness of breath. In most cases, asthma onsets in early childhood with the development of sensitization to environmental allergens. Ongoing environmental exposures lead to airway inflammation and ultimately result in asthma symptoms manifesting within the first few years of life. Recent findings support the notion that asthma develops in association with the human gut microbiome composition early in life[1, 2]. This finding is supported by 16S rRNA sequencing surveys demonstrating that alterations in the gut microbiota precede asthma development within the first few months of life[1, 3].

Early childhood gut microbial communities have been proposed to contribute to asthma by several mechanisms. Epoxide hydrolases encoded by enterococci and other gut bacteria produce the lipokine 12,13-diHOME that predisposes towards atopic sensitization and asthma[3, 4]. Similarly, short-chain fatty acids (SCFAs), produced by the metabolism of dietary fibers by diverse members of the gut microbiota, are thought to protect from asthma through their effect on the host G-protein coupled receptor GPR41, shaping immune cell differentiation in the lungs, and ameliorating allergic airway inflammation[1, 5–8].

In addition to microbially-encoded metabolic features, carriage of antibiotic resistance genes (ARGs) within the gut microbiota, termed the resistome, has been associated with asthma risk. In infants, microbial signatures associated with the development of asthma are also associated with increased richness of ARGs in the gut microbiome[9]. These differences in ARG carriage were found to be driven primarily by *E. coli*, which is a common colonizer in the first days of life[9]. These findings are important in understanding the origins of asthma since antibiotic exposure correlates both to the number of ARGs within the gut microbiome[10] and the later development of asthma and other allergic diseases[11–13]. This association between antibiotic exposure and asthma is supported by animal models that found antibiotic treatment worsens allergic airway inflammation (AAI)[14–16].

While there is an abundance of data supporting the idea that asthma susceptibility is associated with features of the gut microbiota in early childhood, the potential effect of gut microbial functions on asthma later in life remains an important knowledge gap. Since asthma often begins in infancy when the gut microbiota composition is highly unstable, disease-causing microbial functions may not persist into older children and adults. Nevertheless, the gut microbiota in older individuals could underlie the variable manifestations of asthma[17] and may hold valuable prognostic and therapeutic significance.

Asthma-associated differences in later childhood and adult gut microbial communities have already been noted in several reports. Studies in preschool-aged children have noted distinct taxonomic composition of gut microbial communities in asthmatic subjects compared to healthy controls[2]. These differences are reported to include reductions in *Akkermansia muciniphila*[18], *Faecalibacterium prausnitzii*[19] as well as *Roseburia* species[20]. Functional characterization of microbial communities by whole metagenomic sequencing from an older population of asthmatic women[19] has shown that pathways related to lipid and amino acid metabolism, as well as carbohydrate utilization were enriched in asthmatics. In contrast, microbial pathways involved in the production of SCFAs, like butyrate, were enriched in the healthy cohort of the same study[19]. These findings are supported by a complementary study designed to test the effect of probiotic supplementation on asthma that found an association of improved asthma symptoms with SCFA biosynthesis as well as tryptophan metabolism pathways in the adult gut microbiota[21].

Here, we describe an analysis of whole metagenomic sequencing data from a cohort of 36 subjects with physician-diagnosed, moderate-severe asthma along with a matched cohort of 59 healthy controls. This study tests the hypothesis that the gut metagenome harbors signatures of asthma later in life. Our results identify global differences in metagenomic functions between healthy and asthmatic subjects and reveal an enrichment in long-chain fatty acid biosynthetic pathways. We also find an increased richness of ARGs in asthmatics and co-occurrence of ARGs with known bacterial virulence factors, suggesting a potential relationship between antibiotic exposure and pathogen colonization in asthmatics.

## Methods

### MARS Study Population

The Microbiome and Asthma Research Study (MARS) consisted of 104 subjects from the St Louis, MO USA area that are either healthy or had physician-diagnosed moderate-to-severe asthma. This study included an adult cohort (ages 18-40 years) and pediatric cohort (ages 6-10 years). As described in previous manuscripts[22, 23], 9 patients were disqualified or did not donate stool samples. The remaining 95 patients donated stool samples either at home or at the recruitment visit and were evaluated with a clinical questionnaire to gather relevant metadata. Stool samples were kept at −20°C and delivered within 24 hours to the study site, Kau Lab at Washington University School of Medicine, where they were stored at −80°C for no more than three years until processing for DNA isolation. All recruitment, follow up, and sample acquisition occurred between November 2015 and December 2017.

### Fecal DNA Isolation

Frozen human stool samples were pulverized in liquid nitrogen using a pestle and mortar. We then homogenized the stool in a mixture of phenol, chloroform, and isoamyl alcohol with a bead beater using sterilized zirconium and steel beads as previously described[24] to extract crude DNA. We purified the fecal DNA with a 96-well QIAGEN PCR Clean up kit and quantitated by measuring the absorbance at 260/280 nm. Sample DNA concentrations were normalized to 0.5 ng/mL. Neither depletion of human DNA sequence nor enrichment of microbial or viral DNA was performed. No experimental quantification like a spike-in were used.

### Whole Metagenomic Sequencing of Fecal Communities

To generate fecal metagenomic sequencing data, we adapter-ligated libraries by tagmentation using an adaptation of the Nextera Library Prep kit (Illumina, cat. No. FC-121-1030/1031)[25]. Individual libraries were then purified with AMPure XP SPRI beads, quantitated using Quant-iT (Invitrogen, cat. Q33130), and then combined in an equimolar ratio. We confirmed that each library was adequately represented in the combined library by preliminary sequencing on a MiSeq instrument at the Washington University in St. Louis Center for Genome Sciences to assess the evenness of the library. Once the quality of the library was assured, we sequenced the combined library on a NovaSeq 6000 S4 with 2×150 bp chemistry to achieve an average of 3.4 Giga-base-pairs (Gb) per sample. NovaSeq services and data demultiplexing were performed by the Genome Technology Access Center at the McDonnell Genome Institute (St Louis, MO). All samples were tagmented simultaneously and sequenced on the same run to avoid batch effects.

### Processing of sequencing data

Metagenomic raw demultiplexed reads were then processed to (1) remove spurious human sequences (human reference database was hg37dec_v0.1.1), (2) remove low quality sequences, and (3) trim remaining adapter content using Kneaddata v. 0.10.0 (huttenhower.sph.harvard.edu/kneaddata) bypassing the tandem repeat finder step (“- -bypass-trf”). FastQC (fastqc v0.11.7) and MultiQC (multiqc v1.2) with default settings were used to create quality reports and visualize processing steps. See Figure S1A and Table S1 for number of reads dropped per processing step. After trimming and filtering, no samples had adaptor content, overrepresented sequences, or an average sequence quality score below Phred 24. Estimated metagenome coverage was calculated with Nonpareil[26, 27] (version 3.4.1) via the online querying tool at http://enve-omics.ce.gatech.edu/nonpareil/submit.

### Read-based metagenome profiling

To obtain functional information about the metagenomic contents of fecal samples, we processed samples using HUMAnN[28] v3.0.0 on filtered reads with default parameters. The marker gene database used by HUMAnN to identify taxonomic identities was ChocoPhlAn v201901b and the protein database used by HUMAnN to identify functions was the UniRef90 full database v201901b. Alpha diversity analysis of Uniref90 genes and two-sample tests of KEGG orthologs were performed on respective genes that were present (>0 copies per million) in at least 16 out of 95 samples, which was the lowest prevalence cutoff that would allow for Bonferroni corrected Wilcoxon p-values below 0.0001. HUMAnN was used to determine the abundance of metagenomic pathways by mapping UniRef90 genes to the MetaCyc database. We performed differential abundance analysis using the Wilcoxon 2-sample tests on pathways that had a minimum of 10% prevalence.

To identify antibiotic resistance genes present in the fecal metagenomes of MARS stools, we used ShortBRED-identify[29] (v0.9.4) with the Comprehensive Antibiotic Resistance Database[30] (downloaded 2021-07-05 16:10:04.04555) and Virulence Factor Database[31] (downloaded Fri Jul 16 10:06:01 2021). ShortBRED-Quantify was run on the filtered reads with default parameters. ARGs or VFs that had an abundance greater than zero in less than 7 out of 95 samples were excluded from downstream analyses. This prevalence cutoff was determined using the binomial distribution to maintain a 95% confidence that enrichment was not due to random chance (using stats::binom in R). In the analyses that compared virulence factor profiles to antibiotic resistance gene profiles, any gene with the same name was excluded from the list of antibiotic resistance and considered a virulence factor only, to prevent spurious results due to co-correlations. Only one gene matched this criterion: *ugd* (UDP-glucose 6-dehydrogenase).

Microbial composition was determined with MetaPhlAn 3.0[28] which is included in the HUMAnN pipeline described[28]. MaasLin[32] (Maaslin2_1.5.1) was used in R to find taxa of any taxonomic level that correlated with asthma by setting asthma as a fixed effect and setting age group and race as random effects.

For PERMANOVA analyses, BMI class refers to two stratifications: Non-obese (underweight, healthy, or overweight) and obese determined for adults by BMI cutoffs and for pediatric patients by BMI-for-age percentile as defined by the Centers for Disease Control and Prevention (see cdc.gov/healthyweight/assessing/bmi/childrens_bmi/about_childrens_bmi.html). Race was reported by the subject and split into the two categories of Caucasian and non-Caucasian.

### Metagenome Assemblies

Filtered reads were assembled into contigs using spades[33] (v3.14.0) with the “meta” flag and k-mers lengths as follows: -k 21,33,55,77. The resulting scaffolds achieved an average N50 of 3525 <u>+/-</u> 178 bp, an average L50 of 7192 +/- 372 and an average total length of 136.8 +/- 4.5 Mbp as measured by QUAST (v 4.5) [34, 35] (see Table S1). Determination of *ermF* location was performed by aligning the 801-bp coding sequence of *ermF* from CARD[30] to all scaffolds. Scaffolds containing BLAST hits with 98% identity or higher to the full-length CARD *ermF* sequence were further annotated by Prokka (v1.14.5) to find open reading frames and annotate them. Manual BLAST was used to annotate “hypothetical protein” open reading frames for the contexts of *ermF* hits.

### Statistics

R version 3.6.3 was used for all analyses downstream of HUMAnN and ShortBRED, and for data visualization. Wilcoxon tests with false discovery rate multiple testing correction or Type II ANOVAs were used to determine statistically significant differences with the car::Anova package in R. PERMANOVAs were performed in R using the vegan::adonis package with default settings and 100,000 iterations. The following symbols were used to designate significance: * p < 0.05, ** p < 0.01, *** p < 0.001 and the following for q values (FDR-adjusted p-values): * q < 0.2, ** q < 0.05.

## Results

### Whole metagenomic shotgun sequencing of fecal samples from adults and children with asthma and healthy controls

We performed whole metagenomic sequencing on fecal samples from asthmatic subjects and healthy controls taking part in the Microbiome & Asthma Research Study (MARS), which we have previously described[22, 23]. MARS participants were recruited from the St. Louis, Missouri area and included pediatric (6-10 years) and adult (18-40 years) age groups. All asthma cohort patients had a physician diagnosis of moderate-to-severe asthma, and history of allergic sensitization as evidenced by positive skin testing or serum specific-IgE to one or more common aeroallergens. In total, we analyzed 95 patient stool samples including 17 adults and 19 school-aged participants with asthma, and 40 adults and 19 school-aged participants without asthma.

NovaSeq S4 sequencing of our libraries yielded 1.69 billion paired-end reads translating to a total of approximately 500 Gigabases (Gb). After filtering for read quality, dropping host contaminants, and trimming adaptor content, we achieved 1.23 billion paired-end reads and an average 3.4 Gb per stool sample with a range of 0.4-9.9 Gb/sample (Figure S1A). Neither host contamination nor sequencing depth differed between asthma and healthy cohorts (t-test p=0.2 and 0.7, Table S1). All samples achieved an estimated average metagenomic coverage of at 89% (range of 61-98%) with the annotation-free redundancy-based metagenome coverage estimator, Nonpareil[26] (Figure S1B). Further, estimated metagenome coverage was not different between the asthma and healthy cohorts, although we noted coverage was slightly reduced in the pediatric cohort (Figure SB, Table S1). We employed the read-based annotation pipeline, HUMAnN[28] to determine the abundance of genes and functional pathways in the stool metagenomes. We found that the most abundant functional pathways (Figure S1C) across all MARS participants are involved in essential processes of gut microbes such as starch degradation and glycolysis, demonstrating that our sequencing captured core functions of the gut metagenome, as expected. Taken together, we concluded that our sequencing is of sufficient depth and quality to be used for further analyses.

### Gut taxonomic composition differs between people with and without asthma

We first leveraged the clade marker annotation tool, MetaPhlAn[28], to analyze the taxonomic composition of the study participants. We found dominate genera typical in gut microbiota communities including *Bacteroides* (phylum Bacteroidota) and *Faecalibacterium* (phylum Bacillota) (Figure S1D). Simpson alpha diversity was slightly higher in the asthma cohort even when taking read depth and age group into account (Figure S1E). Bray-Curtis dissimilarity (Figure 1F) was shifted between the asthma and healthy cohorts (p<0.0004, R^2^=0.029) even when accounting for other covariates including age (p<0.001, R^2^=0.032), race (p=0.0006, R^2^=0.026), recent antibiotic usage (p=0.9, R^2^=0.006), read depth (p=0.2, R^2^=0.013), obesity (p=0.7, R^2^=0.008), sex (p=0.4, R^2^=0.011), and tobacco exposure (p=0.2, R^2^=0.012) by sequential PERMANOVA (Figure 1G). There was also no significant interaction between asthma status and age group (p=0.8, R^2^=0.007), or between asthma status and recent antibiotic usage (p=0.6, R^2^=0.009) (Figure S1G). To determine differentially abundant taxa, we tested the fixed effect of asthma along with the random effects of age group and race in a general linear model[32] and found *Eubacterium rectale* and *Prevotella copri* were enriched in the healthy cohort (Figure S1H, Table S2). All of these findings are consistent with 16S rRNA sequencing performed in a previous study[23] which lent us further confidence that our sequencing data was suitable for functional profiling.

**Figure 1:**
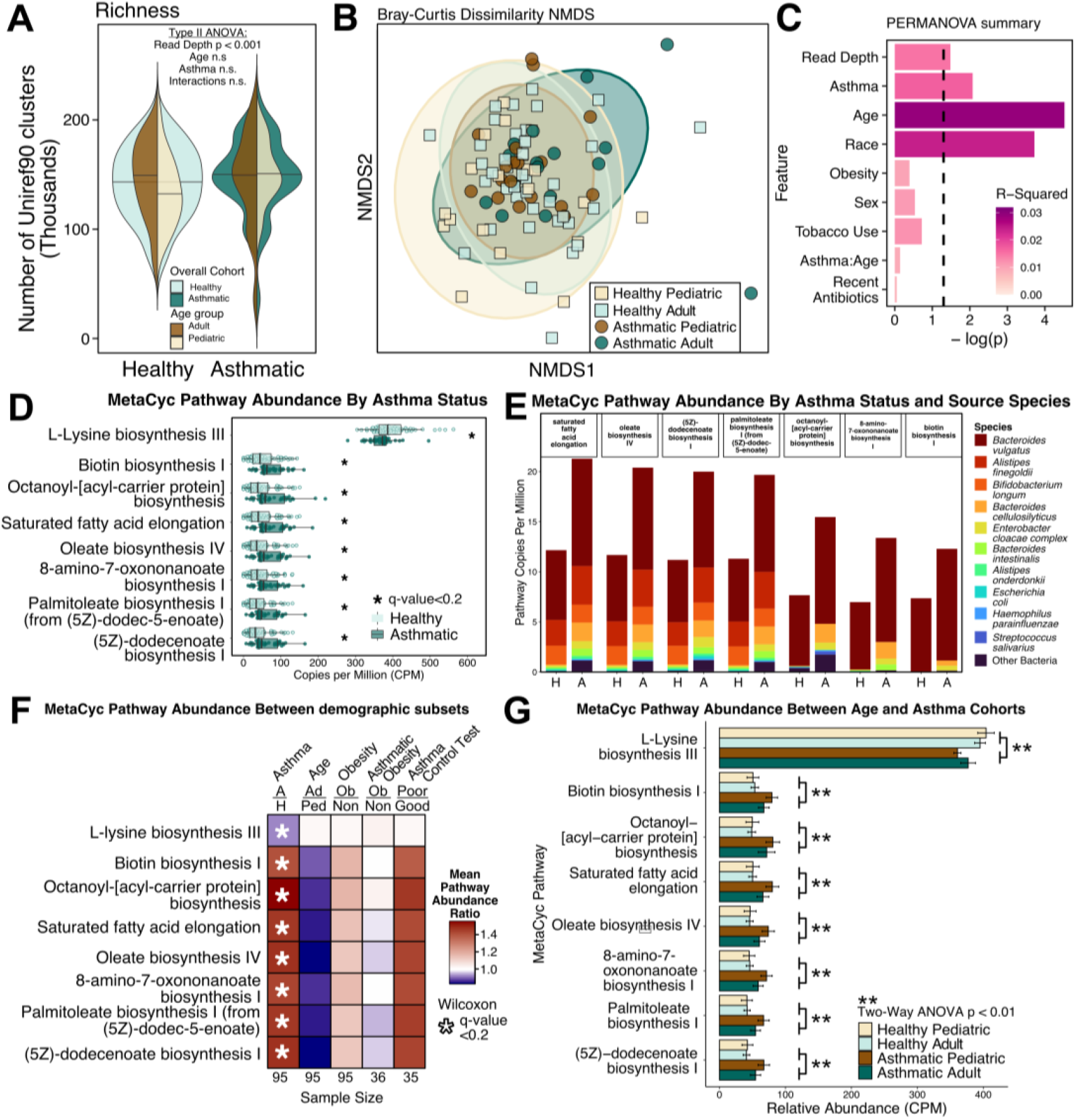
Gut metagenomes from individuals with asthma show increased genes encoding fatty acid metabolism. **A)** Stacked violin plots of Uniref90 cluster richness (unique Uniref90 cluster with CPM>0) grouped by either healthy and asthma cohort (blue green colors in background) or age (brown colors in foreground. **B)** Non-metric multidimensional scaling plot of Bray-Curtis Dissimilarity distance between Uniref90 (copies per million) profiles. Axis 1 and 2 of five total are shown of an NMDS with stress value 0.09. **C)** Sequential PERMANOVA of Bray-Curtis dissimilarities between Uniref90 profiles. Input order of terms to the test is identical to the order of the barplot from top to bottom. **D)** Relative abundance of MetaCyc pathways that were differentially abundant given a Wilcoxon q value below 0.2 (p-value after FDR correction). **E)** Stacked bar plot of differentially abundant fatty acid metabolism pathways mapped to respective taxa by MetaPhlAn3.0/HUMAnN3.0, averaged within asthma or healthy cohorts. **F)** Heatmap of MetaCyc pathway abundance ratios between groups in important clinical demographics: Asthma vs. Healthy, Adult vs. Pediatric, Obese vs Non-Obese, and Well-Controlled Asthmatics vs. Poorly-Controlled Asthmatics. Asterisk denotes a significant differential abundance (*q<0.2) according to Wilcoxon tests controlled for multiple comparison testing within each demographic category. **G)** Differentially abundant MetaCyc pathways plotted as four cohorts: asthma by age with respective Two-Way ANOVAs. Only statistically significant p values shown.

### Fatty acid metabolism pathways are enriched in the gut metagenomes of people with asthma

Given that our samples had adequate coverage to capture expected taxonomic shifts, we started interrogating the differences in metagenomic functions of the gut microbiota attributable to asthma status. The alpha diversity of genes (UniRef90 clusters) was neither different between the asthma and healthy cohorts nor between the pediatric and adult cohorts, suggesting that our gene profiling reached a similar total number of genes in both cohorts (Figure 1A). Using PERMANOVA, we noted that, even while accounting for significant covariates of age (p<0.001, R^2^=0.029), race (p<0.001, R^2^=0.024), and read depth (p=0.03, R^2^=0.015), asthma status also significantly impacted gut microbiome functional composition (p=0.008, R^2^=0.017; Figure 1B, C). We note that age group’s interaction term with asthma did not significantly contribute to the variance in beta diversity, suggesting that the influence of asthma and age on beta diversity is non-overlapping. These findings support the idea that the gut metagenomic content of people with asthma is different than that of healthy individuals, even when accounting for other clinical sources of interpersonal gut microbiome variation.

We next considered which metagenomic functions and metabolic pathways may be involved in the differences between asthma and healthy cohorts. We first examined a list of specific metagenomic functions previously implicated in asthma, including genes related to histamine production, 12-13 diHOME biosynthesis, and tryptophan metabolism, but we were unable to identify a difference between cohorts (Figure S2A). To identify pathways that differed between asthma and healthy subjects, we performed a Wilcoxon Rank Sum test with a false discovery rate q<0.2 on the relative abundance of all pathways annotated by the MetaCyc database that were above 10% prevalence within the population. Using these criteria, we found seven pathways that were enriched in asthma and one that was enriched in the healthy cohort out of 312 total pathways (Figure 1D). To determine if these findings were robust to other analysis methods, we performed additional differential abundance approaches on the 312 MetaCyc pathways, including a Wilcoxon test on centered log-transformed counts and ALDEX2, both of which demonstrated that these pathways differed between healthy and asthmatic cohorts (See Table S3). All differentially abundant pathways enriched in patients with asthma were involved in fatty acid synthesis, and included the production of oleate, palmitoleate, (5Z)-dodecenoate, 8-amino-7-oxononanoate, biotin, and octanoyl acyl-carrier protein, as well as saturated fatty acid elongation. In the healthy cohort, only a single L-lysine biosynthesis pathway was enriched.

Using taxonomically tiered functional mapping, we determined which taxa were driving the observed differences in asthma-associated pathways. For the L-lysine biosynthesis III pathway which was more abundant in healthy subject, we found that it primarily originated from *Blautia obeum*, Figure S2B). In the case of the asthma-enriched pathways, we found that *Bacteroides vulgatus* and *Alistipes finegoldii* account for the largest fraction of complete fatty acid biosynthesis pathways (Figure 1E, Figures S3C). However, the differential abundance of these asthma-associated pathways was probably not due solely to an enrichment of *B. vulgatus* or *A. finegoldii* in asthma stool since neither species was differentially abundant (maaslin2 q-value=0.58 and 0.25, respectively; See Table S2). Further, the majority of mapped pathways were not attributable to any single species and these unmapped pathway counts made up more of the overall pathway richness than *B. vulgatus* (Wilcoxon q values < 0.05 for all seven pathways; see “Community” stratification in Figure S2C). Taken together, these findings indicate that the differences may be either driven by community-level effort (i.e. distinct steps of the pathway are encoded across more than one species), or that current databases are insufficiently granular to identify the key taxa responsible for these differences.

We reviewed the enzymatic steps of each of the eight pathways represented in Figure 1D and found that, of the 78 total reactions in these pathways, only 11 reactions were shared between 2 pathways (Figure S3). The 8-amino-7-oxononanoate biosynthesis I pathway consists of the first 11 reactions of the larger biotin biosynthesis pathway and the latter only has four additional reaction steps past synthesizing 8-amino-7-oxonanoate to produce biotin. Additionally, the (5Z)-dodecenoate pathway can feed directly into the palmitoleate biosynthesis pathway, and that the octanoyl acyl carrier protein pathway shares an upstream substrate (acetoacetyl-acyl carrier protein) with the saturated fatty acid elongation pathway (Figure S3). Together, our findings indicate that long chain fatty acid biosynthesis is differentially abundant in the asthma gut metagenome via related but largely non-redundant pathways.

Given the association between obesity with fatty acid metabolism[36] as well as asthma[37–39], we next wanted to determine whether obesity (which we define here as a BMI greater than 30 in adults or a BMI-for-age percentile of greater than 95% in children) confounds the association of microbial fatty acid metabolism with asthma. We compared the abundance of the differentially abundant fatty acid pathways between all non-obese and obese patients and found no significant difference (Figure 1F). Within the asthma cohort, there was similarly no statistically significant difference between the asthmatic obese and asthmatic non-obese patients, suggesting that obesity is not a confounder for the difference we observed in fatty acid metabolism. To determine whether fatty acid metabolism is related to the intensity of asthma symptoms and their effect on everyday life activities, we utilized a validated survey of asthma control (The Asthma Control Test; ACT)[40]. None of the fatty acid pathways were differentially abundant between well-controlled and poorly-controlled asthmatics (Figure 1F). We tested if age group affects the differentially abundant metabolic pathways and found that these pathways were not differentially abundant between age groups alone (Figure 1F). We also tested the impact of asthma and age as independent variables to differentially abundant metabolic pathways using a Two-way ANOVA. We found that, even while taking age into account, these pathways are differentially abundant between asthma and healthy cohorts, but are not different by age or an interaction between asthma and age (Figure 1G, 2-Way ANOVA). Given that the effect of asthma status on differentially abundant metagenomic functions was distinct from that of age, we primarily focused our subsequent analyses on the asthma and healthy cohorts overall, combining age groups.

### Richness of antibiotic resistance genes is increased in the gut metagenomes of people with asthma

Since people with asthma tend to be prescribed antibiotics frequently[41] and oral antibiotic exposure is a risk factor for the acquisition of ARGs in the gut[10], we wanted to determine if the members of our asthma cohort were more likely to have received antibiotics. To test this, we counted how many subjects had taken a course of antibiotics within one year of their participation in the study. As part of the study design, participants could not take antibiotics in the month prior to fecal donation. We found that a greater proportion of the asthma cohort received antibiotics in the past year compared to that of healthy participants (42% of asthma cohort versus 15% of the healthy cohort, Fisher’s test, p=0.011, Figure 2A). This finding represents evidence of increased antibiotic exposure amongst subjects with asthma in our study.

**Figure 2:**
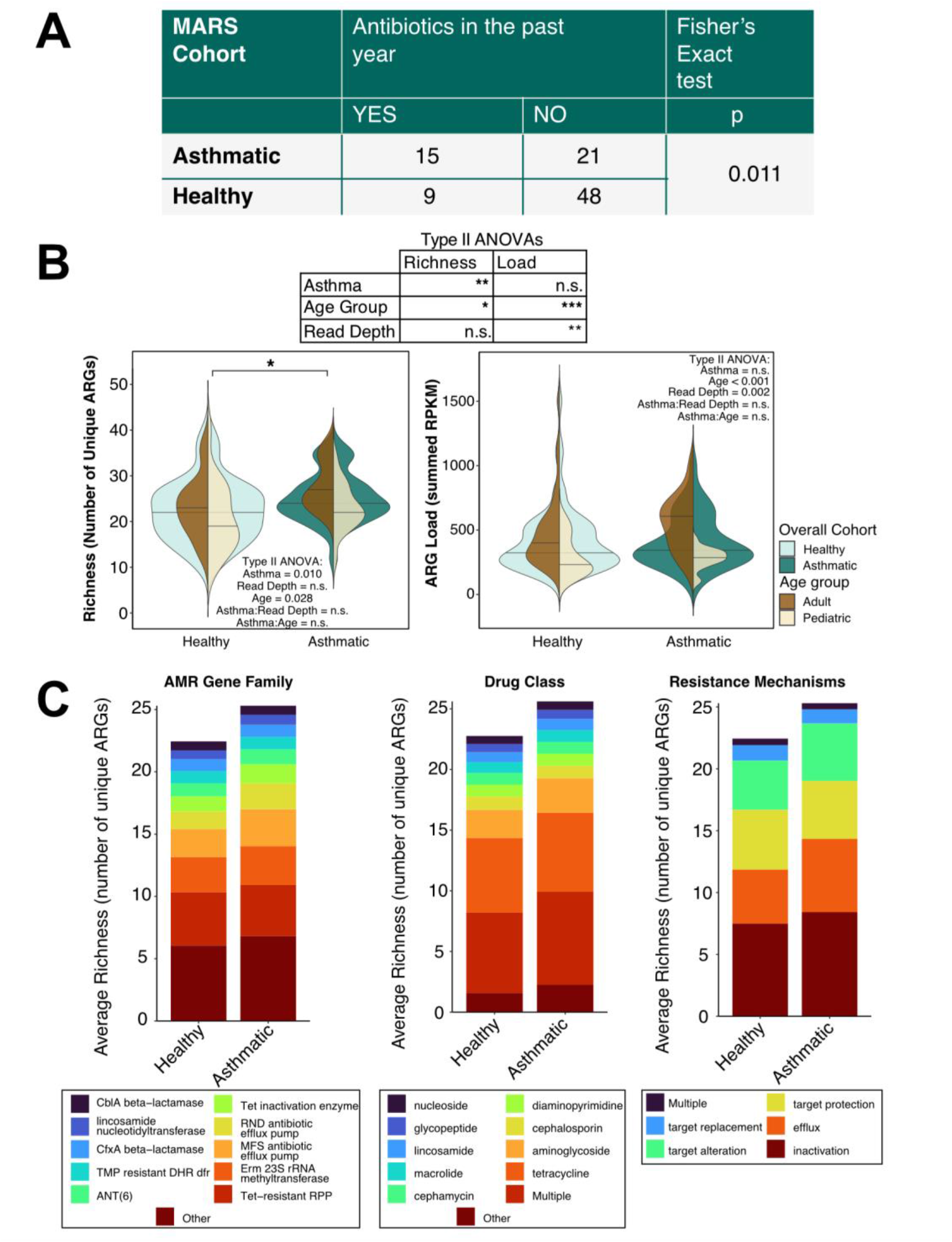
Gut metagenomes from individuals with asthma harbor an increased richness of antibiotic resistance genes. **A)** Table describing short-term antibiotic usage in the MARS cohorts. **B)** Overlapping violin plots of ARG richness and load by grouped by either healthy and asthma cohort (blue green colors in background) or age (brown colors in foreground. **C)** Stacked bar plots of average ARG richness painted by antimicrobial family (AMR), drug class to which the ARG confers resistance, and ARG resistance mechanism.

We next sought to characterize the gut antibiotic resistome in the asthma and healthy cohorts. To test if the increased antibiotic exposure in the asthma cohort was reflected in the gut resistome, we utilized the ShortBRED pipeline[29] to detect reads mapped to the Comprehensive Antibiotic Resistance Database (CARD)[30]. We first asked whether there were more ARGs in our asthma cohort by summarizing our dataset into richness (Total number of unique ARGs detected per sample) and load (Total sum of ARG RPKM per sample). We found that ARG richness was higher in people with asthma even when accounting for differences due to age (p=0.03) and sequencing depth (p=0.09 while ARG load was not different between asthma and healthy cohorts (p=0.4) when accounting for age (p<0.001) and read depth (0.002) (Figure 2B). We note that *E coli* was not differentially abundant between asthma and healthy cohorts (p=0.52, Table S2), so the richness increase we observe in the asthma cohort is not due solely to an increase in *E. coli* relative abundance. These results suggest that there are a higher number of unique ARGs, or a higher diversity, in asthma compared to healthy controls.

From our 95 stool samples, we detected 71 unique ARGs, comprising 32 antimicrobial resistance families, 29 drug classes, and 7 mechanisms of resistance, with 26 ARGs (37% of the total) conferring multi-drug resistance (Figure 2C). Similar to previous studies of gut resistomes, we found that tetracycline resistance markers were the most commonly detected ARGs and inactivation is the most common mechanism of resistance followed by efflux pumps[9] (Figure 2C). Using the abundance data of each detected ARG, we determined that asthma (p=0.005, R^2^=0.028) and age (p<0.001, R^2^=0.053) were the strongest factors contributing to the variance in ARG beta diversity even when accounting for important technical and demographic covariates (Figure 3A and 3B). We next wanted to ascertain to what degree the resistome profile was determined by microbial composition. We used a Procrustes analysis[42] to compare compositional data generated from MetaPhlAn[28] to the antibiotic resistome profile derived from ShortBRED and found that the microbiome composition correlated to the resistome profile (Figure 3C, PROTEST corr = 0.627, p-value < 0.0001), indicating that ARG profiles are directly related to bacterial species composition.

**Figure 3:**
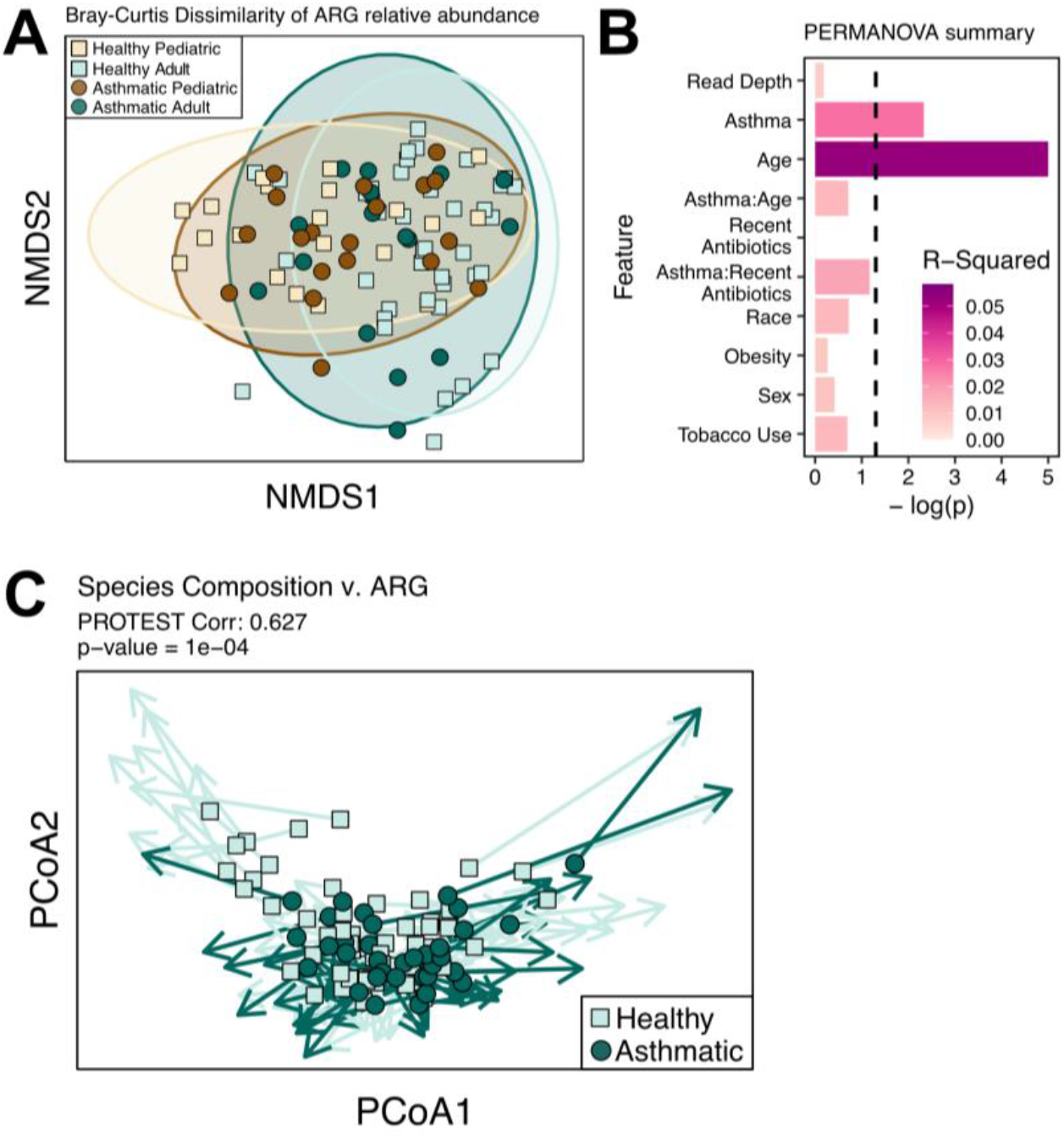
The gut antibiotic resistome is altered in asthma patients. **A)** Non-metric Multidimensional Scaling (NMDS) plot of antibiotic resistome with units in Bray-Curtis dissimilarity of total-sum scaled RPKM, labeled by asthma and age cohorts. Showing two axes out of five with stress value=0.1. **B)** Effect of demographic categories on antibiotic resistome data in A (sequential PERMANOVA). **C)** Procrustes and PROTEST analysis between MetaPhlAn species-level Bray-Curtis dissimilarity distances and CARD ShortBRED Bray-Curtis dissimilarity distances. Arrows connect the two data points belonging to identical samples.

### Macrolide resistance markers are differentially abundant in asthma

To determine ARGs that are differentially abundant between asthmatic and healthy gut metagenomes, we applied negative binomial tests to the abundance of all ARGs detected in at least 7 samples. This prevalence cutoff was chose because it is the minimum number of samples needed to detect a difference using a negative binomial distribution. We found that genes encoding resistance to macrolides (*ermF, ermB* and *ermA*), vancomycin (*vanRO*), tetracycline (*tet(45)*), as well as multi-drug efflux pumps (*smeB, mdtO*, and *oqxA*) were enriched in the asthma cohort (Figure 4A, Table S4). Prominent amongst these was the 23S rRNA methyltransferase *ermF*, which is typically encoded by *Bacteroides* species and confers resistance to macrolides.

**Figure 4:**
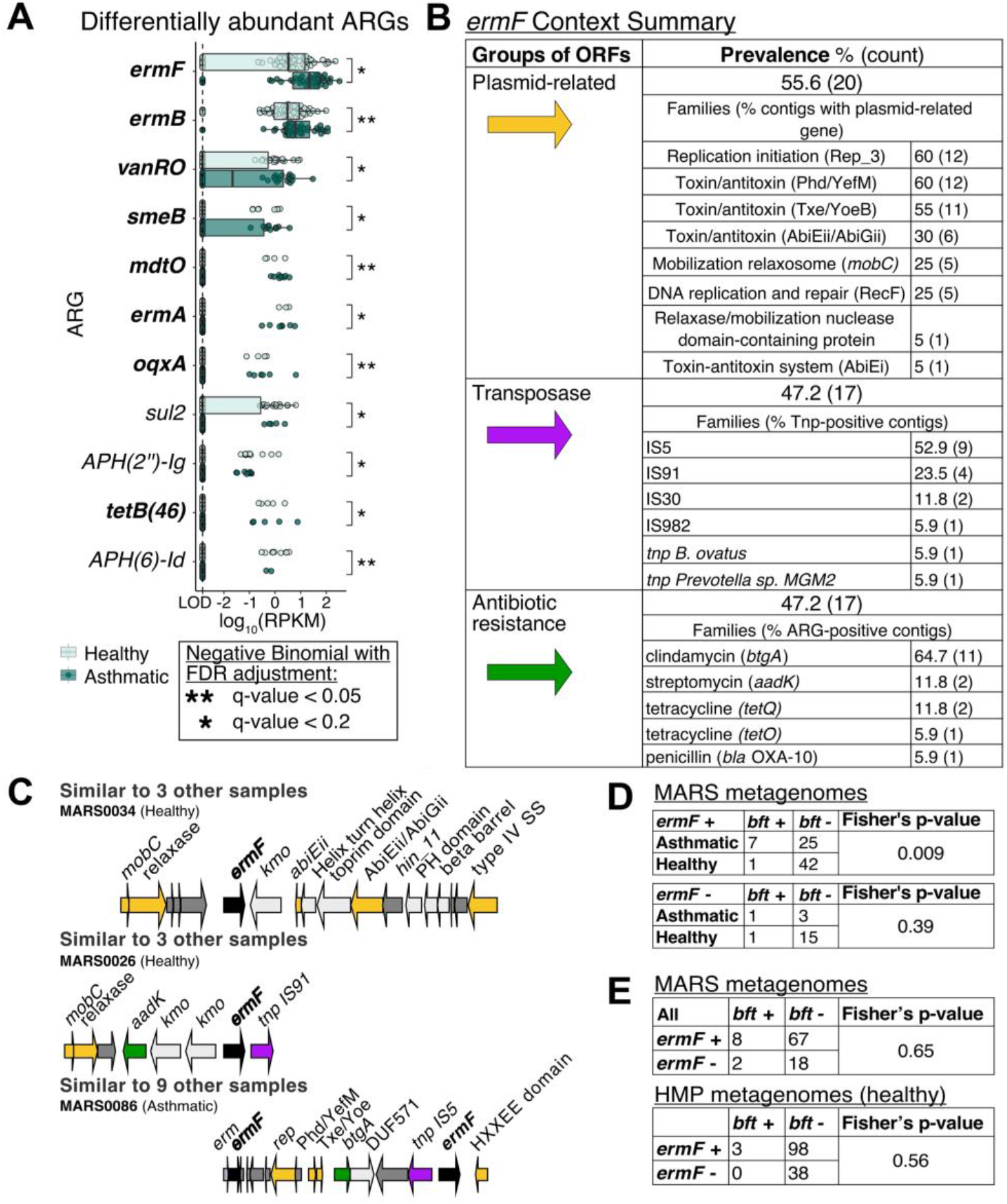
Resistance gene *ermF* is differentially abundant in diverse genomic contexts of gut resistomes belonging to individuals with asthma. **A)** Boxplots of antibiotic resistance gene (ARG) abundance by cohort on log-scale. Showing only ARGs present in at least 7 out of 95 samples and have q-values less than 0.2. A pseudocount of 0.0015 RPKM (designated as the limit of detection “LOD”) was used for the negative binomial tests. Bolded genes are enriched in the asthma cohort while non-bolded are enriched in the healthy cohort. **B)** Summary of *ermF* contexts on contigs from metagenomic assemblies that had at least one detectable open reading frame flanking the *ermF* within 10 kilobases. **C)** Three representative *ermF* context maps generated in GeneSpy. **D)** Count tables of *ermF*+ (top) and *ermF-* (bottom) MARS fecal samples split by *bft* presence and asthma status. Both tables showing two-sided p-value. **E)** Count tables of metagenomes (top from MARS and bottom from Human Microbiome Project) split by the presence of *B. fragilis* toxin (*bft*) and *ermF*. Top: one-sided p-value shown; bottom: two-sided p-value shown.

Next, we explored the genomic context of *ermF* by assembling metagenomic sequencing reads into contigs with metaSPAdes[33] and annotating open reading frames with Prokka[43] and BLAST. We detected full-length *ermF* with 98% or higher identity in 53 out of 95 samples. Out of 53 contigs, the vast majority originated from members of the Bacteroidota, 75.4% originated from the *Bacteroides* genus and 60.3% of them were likely from *B. fragilis* based on the top BLAST homology. Of the contigs that encoded *ermF*, 68% occurred on scaffolds with at least one other open reading frame within ten kilobases (Figure 4B). We found that many *ermF genes* are co-located with genes associated with mobile genetic elements such as transposases, mobilization genes, and toxin/antitoxin systems, as well as with other ARGs like *btgA* which encodes clindamycin resistance (Figure 4B,C). This indicates that *ermF* occurs in multiple different genomic contexts within our cohort and suggests that its presence is not strictly due to propagation of a single *B. fragilis* strain.

### People with asthma have a distinct set of co-existing pairs of antibiotic resistance genes and virulence factors in the gut metagenome

In our prior work on this same cohort of patients, we found that, compared to healthy subjects, a greater portion of asthma subjects were colonized with *B. fragilis* strains harboring the virulence factor *B. fragilis* toxin (*bft*), which we showed has the potential to shape inflammation in the lung[23]. Given that our resistome analysis pointed to an enrichment of a *B. fragilis* ARG, we wanted to test whether the *ermF* gene is co-selected with *bft*. When only taking metagenomes encoding *ermF* into account, we observed an enrichment of *bft* prevalence in the asthma cohort (Figure 4D, p=0.009). In contrast, among metagenomes with no detectable *ermF*, there is no enrichment of *bft* in the asthma cohort (Figure 4D, p=0.39). When reviewing the entire MARS population, we found no statistically significant co-occurrence of *ermF* with *bft* (Figure 4E; one-tailed Fishers test p>0.05) and this was consistent with healthy gut metagenomes from the Human Microbiome Project (Figure 4E, Fisher’s test p=0.56). However, in our MARS samples, we did not find any instances where *bft* and *ermF* occurred on the same scaffold, so it remains unclear whether these two genes are encoded within the same *B. fragilis* strain or within two separate strains. Nevertheless, these results suggest that the environment supporting the gut microbiota of asthmatic individuals presents opportunities or niches for *ermF* and *bft* to co-occur.

To explore the possibility that virulence traits and ARGs are linked in the gut microbiota, we characterized virulence factor (VF) content of all samples using the Virulence Factor Database[31] and compared these data to the antibiotic resistome profiles. We did not find the same overall shift in the virulence factor beta diversity between asthma and healthy that we observed with the resistomes (Figure S4A-C), but we did find differentially abundant VFs belonging to capsule and peritrichous flagella VF families (Table S5, q values<0.2). Further, we found that microbiota composition is highly correlated with virulence factor profile (Figure S4D, Protest correlation coefficient=0.61, p<0.0001). Given that microbiota composition strongly affects both VF and ARG content, we used a partial correlation between VF and ARG richness to test our hypothesis while removing the effect of total metagenomic content. We found a positive partial correlation between VF and ARG richness in both the asthma and healthy cohorts (Figure 5A). Similarly, virulence factor and resistome beta diversity profiles were also positively correlated (Figure 5B, Protest correlation coefficient=0.574, p=1e-4). Together, our results suggest that these two microbial features, virulence and antibiotic resistance, are closely linked within the gut metagenome.

**Figure 5:**
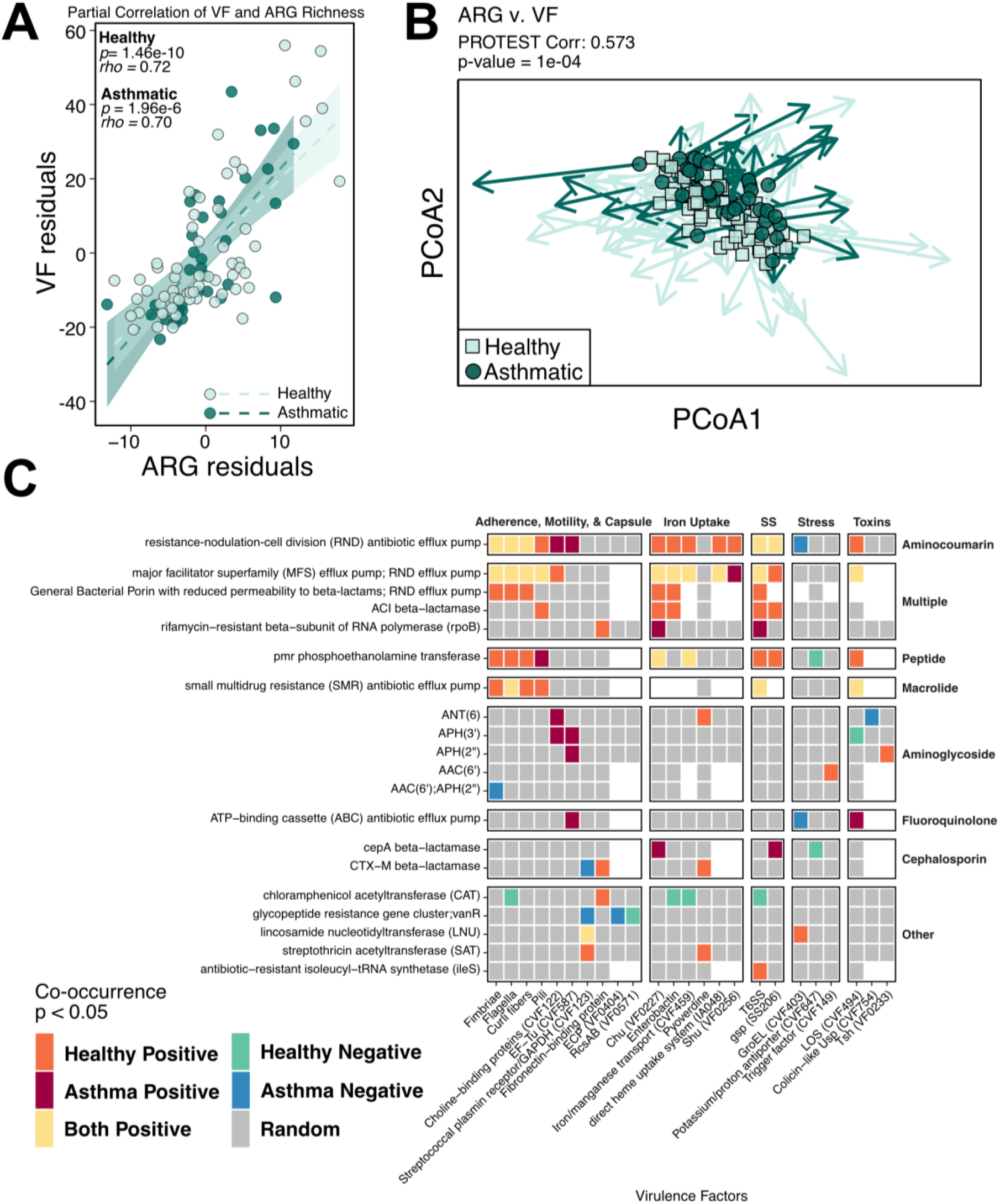
Asthma patients have unique sets of virulence factor and antibiotic resistance gene associations. **A)** Partial correlations split by asthma status between virulence factor richness and ARG richness after accounting for species richness. **B)** Procrustes and PROTEST analysis between Bray-Curtis dissimilarity distances of virulence factors and CARD resistomes. Arrows connect the two data points belonging to identical samples. **C)** Heatmap of statistically significant (cooccur R package p<0.05) co-occurrence relationships between all VFs and ARGs. Colors indicate direction of co-occurrence and in which cohort(s) the respective effect was detected. Grey squares mark pairs with no statistically significant co-occurrence. White squares were pairs filtered out due to a lack of observed co-occurrence.

We next performed a co-occurrence analysis to uncover other linked virulence and antibiotic resistance traits that could be important in gut ecology. We found numerous co-occurring VF-ARG pairs in MARS gut metagenomes (Figure 5C, p<0.05). Several of these positively co-occurring pairs were shared between the two cohorts (yellow), suggesting that these relationships are not dependent on asthma status. In contrast, many pairs specifically co-occur in one cohort and may indicate microbial interactions important in asthma but not healthy gut metagenomes (Figure 5C). In summary, we found that VF and ARG presence is linked in the gut metagenome and that people with asthma have a distinct set of co-occurring functions compared to healthy people.

While our co-occurrence analysis between VFs and ARGs demonstrated multiple examples of virulence and antibiotic resistance traits found in the same gut metagenome, this analysis does not indicate if these genes are present in a single organism. To obtain a more granular view of VF-ARG co-occurrence, we limited our analysis to look for VF-ARG pairs that could be encoded by the same species. This analysis showed that the asthma cohort had a greater number of ARGs (p=0.007 and 0.01) and VFs (p=0.005 and 0.09) annotated as coming from *Klebsiella pneumoniae* and *Escherichia coli*, respectively (Figure S5A). Individual co-occurrences attributable to each of these species are summarized in Figure S5B and show that *cepA*, encoding a beta-lactamase, and *chuU*, a VF involved in iron acquisition, are both putatively encoded by *E. coli* and co-occur in asthmatics, suggesting that the metagenome-wide co-occurrence of CepA and Chu families observed in Figure 5C may be due to enrichment within one or more *E. coli* strains harboring these VF/ARG pairs. Together, our co-occurrence analyses show that there appear to be multiple co-occurring VFs and ARGs, similar to *B. fragilis*-encoded *bft* and *ermF*, in the gut metagenome and within putative individual species that could be important for asthma. The cohort-specific co-occurring VF-ARG pairs found here could serve as candidates for future studies of asthma gut microbiome ecology.

## Discussion

In this study, we present an exploratory analysis of fecal whole metagenomic sequencing contrasting subjects with moderate-to-severe asthma to a group of healthy controls to identify disease-associated microbial genes with the strongest likelihood of affecting disease. Our sequencing and subsequent analyses revealed that the functional content of individuals with asthma differed significantly from that of healthy controls. We found an enrichment of functions associated with saturated and mono-unsaturated fatty acids, including oleate, palmitoleate, 5(Z)-dodecenoate, biotin, 8-amino-oxononanoate, saturated fatty acid elongation, and octanoyl acyl carrier protein pathways. Currently, the functional significance of gut bacterial synthesis of these long-chain fatty acids (LCFA) to asthma has not been well defined. Excess LCFAs, usually studied in the context of dietary fat intake, have been associated with metabolic diseases including diabetes, obesity, and atherosclerosis risk[38] but is also linked to asthma risk in adults[37–39, 44]. Increasing recognition that obesity predisposes to asthma has motivated investigation of the impact of fatty acids on airway biology and has shown that LCFA signaling through free fatty acid receptor 1 (FFAR1, also called GPR40) induces airway smooth muscle cell contraction and proliferation, both of which are important components of asthma pathophysiology[38, 45]. Notably, a study that sequenced airway microbes in children with cystic fibrosis implicated a similar list of LCFA production pathways during exacerbations, suggesting that microbially produced LCFAs may influence airway physiology[46]. To our knowledge, the potential for gut microbes to contribute to the amount of free fatty acids available to the lung has not yet been defined, however, LCFAs are readily absorbed into the circulation[47] and could plausibly reach the airways. Further, previous studies have shown the effect of SCFA (e.g. acetate, butyrate, propionate) produced by gut microbes to directly alter lung inflammation via GPR41 (FFAR3)[7, 8]. While our study did not find a direct enrichment of SCFA production pathways in the healthy cohort as has been previously reported[19], we did observe that lysine biosynthesis was enriched. Since lysine may serve as a precursor to the SCFA butyrate[48], SCFAs may still be more abundant in our healthy cohort but may be subject to transcriptional regulation that would not be detected by metagenomic DNA sequencing. Together, our metabolic pathway analyses of the gut metagenome demonstrate a positive association between LCFAs produced by gut microbes and asthma, in contrast to the negatively associated SCFAs.

In addition to metabolic alterations, analysis of the gut resistome demonstrated that subjects with asthma had a distinct ARG composition. In a recently published prospective gut metagenomic study of infants, asthma-associated taxonomic signatures were associated with a higher number of ARGs[9]. These differences in the resistome were largely driven by a single species of bacteria, *E. coli*, and reveals that acquisition of ARGs in subjects with asthma may begin in early childhood and could affect asthma development. In our study of older subjects with established asthma, we similarly found a higher richness of ARGs that is associated with asthma in both school-aged children and adults, supporting the idea that increased ARG carriage may persist in people with asthma throughout life. Based on our resistome annotation, however, ARGs in our cohort were likely from a diverse assemblage of bacteria in contrast to what was observed in infants. This is likely due to differences in gut dynamics between age groups. The infant microbiome is heavily shaped by limited available niches in the developing gut, which favor transient, facultative anaerobes like *E. coli* [9], whereas the gut resistome in older subjects reflects selective pressures experienced over a lifetime. One important consequence of increased richness of ARGs in people with asthma is that it may promote persistence of some bacterial strains[49, 50] and contribute to the taxonomic differences in the gut microbiota between asthma and healthy people[2, 23].

While asthma was among the important factors accounting for a significant amount of the variance in ARG beta diversity, we found that recent antibiotic exposure (within the past year) was not. Notably, no participant in our cohort received a course of antibiotics in the month prior to fecal sampling since this could have confounded our analyses on asthma-associated microbial community changes. Previous studies have shown that the gut microbiota recovers in approximately a month after perturbation from antibiotics in healthy adults[51]. We interpret these findings to mean that recent exposure (within 1 - 12 months) to antibiotics does not drastically change the resistome, whereas repeated exposures over time may be more important for driving the population-wide shifts we observed in our cohort[50].

Of the ARGs found to be enriched within asthmatic resistomes, the ARG *ermF*, encoding resistance to macrolide antibiotics, was especially prominent amongst the asthmatic cohort. While we did not collect data on the antibiotic drug classes, number of courses and their duration, or the reason for prescription of antibiotics, our subjects received, it is likely that our asthma population has been exposed to macrolides. Macrolide antibiotics, including clarithromycin and azithromycin, are commonly prescribed for upper and lower airway infections which disproportionately affect people with asthma[52]. This class of antibiotics, particularly azithromycin, have been a focus of special concern for driving antibiotic resistance due to their frequent usage and pharmacological properties[53–55]. Nevertheless, azithromycin has been noted to have beneficial effects in asthma, and some[56], but not all[57], studies suggest that azithromycin may prevent exacerbations in asthmatics. Given the interest in azithromycin as a treatment modality in asthma, there will be an urgent need for additional studies to determine the robustness of the association between asthma and macrolide ARG accumulation in the gut to inform parameters for antibiotic selection and prescription in people with asthma.

Additional exploration of the gut metagenomes revealed potential co-selection in people with asthma for *B. fragilis* genes *ermF* and *bft* (*B. fragilis* toxin), the latter of which is more prevalent in fecal samples from the asthma compared to healthy cohort[23]. Untargeted analysis of gut resistomes revealed multiple examples of virulence factor and ARG co-occurrence as well as positive correlations between ARG and VF richness in people with and without asthma. Our findings are consistent with previous reports that found correlations between VFs and ARG richness and VF-ARG cooccurrence relationships in both gut metagenomes[58] and human-associated bacterial genomes[59]. Our findings also add to these studies by demonstrating that, while the correlation between VF and ARG richness does not appear to be any stronger in the asthma cohort after taking gene richness into account, the two MARS cohorts do not have identical sets of statistically significant co-occurring VF-ARG pairs. These data suggest that people with asthma may be experiencing different selection pressures from that of healthy people, leading to accumulation of a distinct set of virulence and antibiotic determinants. Given that antibiotics induce gut inflammation through the disruption of the gut microbiota[60], and strains encoding virulence factors such as *bft* are known to thrive in an inflammatory environment[61], one plausible model for the apparent accumulation of distinct VF-ARG pairs is that antibiotic treatment not only selects for ARGs[10, 50], but simultaneously selects for VFs. Together with evidence that virulence determinants, such as *bft*, are associated with airway inflammation[23], our model implies that heightened antibiotic treatment may contribute to the manifestations of asthma via co-selection for VFs and ARGs. Considering that prenatal and early life antibiotic exposure is linked to asthma risk[12, 60], this model could be used to test whether the initial events driving VF and ARG co-occurrence start with the first vertical transmission events in very early life.

Our study has several limitations that constrain the scope of our claims. First, MARS is an exploratory, cross-sectional study with only a moderate number of subjects recruited from a single site, which is less ideal for identifying disease-associated microbiome differences[62]. As a result, our study had limited statistical power to detect less prevalent or abundant functions. Second, our study focused on school-aged and older subjects with moderate-to-severe asthma, and thus our findings may not be applicable to other younger populations or those with less severe disease. These population differences may explain why we were unable to identify statistically significant differences in microbial metabolic pathways identified from other studies including bile acid metabolism[1], epoxide hydrolases[4], histamine metabolism[63, 64], or tryptophan metabolism[65, 66] (Figure S2A). Third, the factors driving the shift in gut bacterial metabolism to LCFA biosynthesis and whether gut microbiome enrichment of this pathway is sufficient to change the hosts’ LCFA profile is not known. Collecting blood to interrogate host metabolism as well as dietary information at the time of fecal sample collection would have helped to disentangle the effects of diet on host and gut microbiota metabolism. Fourth, a record of the frequency and class of antibiotics administered to our participants would have allowed us to confirm whether macrolide administration associates with the enrichment of *ermF* in our asthma cohort and whether a higher diversity of antibiotic usage correlates with ARG richness. It is likely that antibiotic exposures accumulated throughout life contribute to the resistome, and a complete catalog of exposures is critical to determine patterns of antibiotic prescription most likely to account for the ARG associations to asthma found in this study. Fifth, as with all metagenomic sequencing studies, we are limited by annotation bias in existing databases. This is a concern for our virulence factor and antibiotic resistance profiling especially, where we rely on the database to predict source species for ARGs and VFs. We also recognize that the databases we used for these two analyses are biased towards well-studied human pathogens rather than commensals or opportunistic pathogens. However, we note that other investigators have reported similar co-occurrence of ARGs and VFs[58, 59], and co-selection of these features is biologically plausible.

Despite these constraints on the scope of our study, we provide evidence that there is an increased production of LCFA and an increased richness of ARGs encoded by the gut microbiota in people with asthma. These findings could have applications in the care of patients with asthma. If LCFA pathways are shown to play a causal role in airway inflammation in future studies, microbiota-directed therapeutics in the form of dietary interventions or probiotics, could be developed to modify gut microbial metabolism to protect against asthma. Additionally, our resistome findings add to the growing concern over antibiotic resistance in patients with asthma by suggesting that antibiotic administration may also contribute to gut carriage of virulence factors that can alter airway inflammation. Ultimately, our study shows that the gut microbiota of school-aged and older subjects with moderate-to-severe asthma harbor important functional alterations that could serve as a foundation for future studies investigating how gut microbial functions affect pulmonary diseases.

## Conclusions

Asthma is an airway disease that affects the everyday lives of millions of people and accounts for approximately 1.5 million emergency room visits yearly in the US[67] Both antibiotic usage and gut microbiota dysbiosis have been linked to the development of asthma, however, little is known about the specific gut microbial functions associated with asthma, particularly in older populations. In this study, we characterized the gut microbiota of school-aged children and adults with moderate-to-severe asthma and uncovered asthma-associated microbial functions that may contribute to disease features. We found that people with asthma have an increase in gut microbial genes associated with long-chain fatty acid metabolism as well as an accumulation of antibiotic resistance genes, both of which may have practical consequences for monitoring and treatment of asthma.

## List of Abbreviations

(AAI): Allergic airway inflammation
(ARG): antibiotic resistance gene
(LFCA): long-chain fatty acid
(SCFA): short-chain fatty acid
(VF): virulence factor

## Declarations

### Ethics approval and consent to participate

This study was approved by the Washington University Institutional Review Board (IRB# 201412035). Written informed consent documents were obtained from all MARS subjects or their legal guardians.

### Consent for publication

Not applicable.

### Availability of data and material

The metagenomic sequencing dataset generated during the current study are available at European Nucleotide Archive (https://www.ebi.ac.uk/ena/browser/home) under project accession number PRJEB56741. Demographic data needed to reproduce results can be found in this manuscript (Table S1). A full record of all statistical analyses is included as a PDF document generated by knitr in R[68] in Additional File 1. A STORMS (Strengthening The Organizing and Reporting of Microbiome Studies) checklist[69] is available at doi: 10.5281/zenodo.7492635.

### Author Contributions

N.G.W. and A.L.K. conceptualized the work. L.B.B. and A.L.K. planned the clinical study. N.G.W., A.H-L., D.J.S., and A.L.K. analyzed the data and drafted the manuscript. All authors interpreted the data, and read and approved the manuscript.

## Acknowledgements

We would like to thank our clinical study coordinators Tarisa Mantia, Caitlin O’Shaughnessy, and Shannon Rook; the physicians of Washington University Pediatric and Adolescent Ambulatory Research Consortium, especially Dr. Jane Garbutt; the Volunteer for Health registry; and the MARS participants and their families. We would also like to acknowledge the Center for Genome Sciences Sequencing Core and Genome Technology Access Center at the McDonnell Genome Institute for sequencing services. The authors also thank Dr. Leyao Wang and members of the Kau lab for critical reading of the manuscript.

## Supplemental Figure and Data Table Captions

### Figures

**Figure S1: MARS whole metagenomic shotgun sequencing captures essential functions and taxonomic shifts of the asthma gut microbiota. A)** Summary of select sequencing statistics from NovaSeq shotgun metagenomic sequencing and subsequent filtering steps. **B)** Boxplot of redundancy-based estimated metagenome coverage (%) as calculated by running the forward reads through the Nonpareil tool. Split into asthma and age group and Two-way Type II ANOVA results shown. **C)** Bar plot of MetaCyc pathway copies per million (CPM) in all MARS samples annotated by HUMANnN pipeline, with horizontal length representing mean and bars the standard error. For all panels: N= 20 healthy children, 39 healthy adults, 19 asthmatic children, 17 asthmatic adults. **D)** Relative abundance stacked barplots of top abundant bacterial genera split by age group and asthma cohort. **E)** Simpson alpha diversity boxplots split by asthma and age group cohorts (2-Way Type II ANOVA). **F)** NMDS of Bray-Curtis Dissimilarity of species-level relative abundance grouped by age and asthma. **G)** Sequential PERMANOVA to test effect of demographics on beta diversity. Terms were input into the test as ordered from top to bottom of barplot. Dotted vertical line represents a p value of 0.05. Color scale is mapped to the R^2^ value. **H)** Arcsine transformed relative abundance boxplots of differentially abundant species as determined by Maaslin2 with age group and race modeled as random effects. For all panels: N= 20 healthy children, 39 healthy adults, 19 asthmatic children, 17 asthmatic adults.

**Figure S2: KEGG orthologs, KEGG pathways, and differentially abundant MetaCyc fatty acid pathways. A)** Relative abundance of KEGG orthologs previously implicated in asthma. Copies per million (CPM) are counts normalized by gene size and read depth, then total-sum-scaled to one million. **B-C)** Stacked bar plots of differentially abundant pathways mapped to respective taxa including “Community” bin which accounts for the remaining reads that mapped to the pathway but not to any single species by MetaPhlAn3.0/HUMAnN3.0, averaged within asthma or healthy cohorts. **B)** L-lysine biosynthesis III pathway. Only top 13 taxa shown in addition to Community category. **C)** Seven fatty acid metabolism pathways differentially abundant in the asthma cohort. Only top 9 taxa shown in addition to Community category. Stars represent a q value < 0.05 of Wilcoxon tests between the Community pathway richness and *B. vulgatus*-encoded pathway richness. For all panels: N= 59 healthy, 36 asthmatic individuals.

**Figure S3: Pathway collage for differentially abundant MetaCyc pathways**. “PWY-6519: 8-amino-7-oxononanoate biosynthesis I” is completely overlapping with “BIOTIN-BIOSYNTHESIS-PWY: biotin biosynthesis I” and its steps are highlighted in blue text. Pathway collage made on MetaCyc browser tool.

**Figure S4: Gut virulence factor ecology shifts with age group but not asthma cohort. A)** Total-sum scaled RPKM Bray-Curtis Dissimilarity Non-metric Multidimensional Scaling (NMDS) plot labeled by asthma and age cohorts. Showing two axes out of 5 with stress value=0.09. **B)** Effect of demographic categories on virulence factor profile in A (by sequential PERMANOVA, input terms ordered from top to bottom of barplot). **C)** Stacked violin plots of virulence factor alpha diversity grouped by eithter healthy and asthma cohort (blue green colors in background) or age (brown colors in foreground). Two-Way ANOVA results shown in table above plot. **D)** Procrustes plot and PROTEST analysis between virulence factor profile Bray-Curtis dissimilarity distances and Metaphlan species relative abundance Bray-Curtis dissimilarity distances. Arrows connect the two data points belonging to identical samples. For all panels: N= 20 healthy children, 39 healthy adults, 19 asthmatic children, 17 asthmatic adults.

**Figure S5: Asthma-associated ARG richness and ARG-VF co-occurrence relationships are observed within *K. pneumoniae* and *E. coli*. A)** Richness bar plots between antibiotic resistance genes (ARGs) and virulence factors (VFs) grouped by asthma status. **B)** Heatmap of the co-occurrence of each VF/ARG pair colored by the direction in which (positively or negatively co-occurring) and the cohort for which (asthma vs. healthy) the pair had a p-value less than 0.05 via R cooccur function. Blank squares were pairs filtered out due to a lack of observed co-occurrence. SS: secretion systems. (N=36 Healthy, 59 Asthmatic)

### Tables

Table S1: Fecal shotgun metagenomics filtering and assembly summary statistics

Table S2: Maaslin 2 Analysis of Metaphlan Community Composition

Table S3: Comparison of MetaCyc Pathway Differential Abundance Analyses

Table S4: Negative Binomial tests of antibiotic resistance gene abundance (RPKM) between healthy and asthma cohorts

Table S5: Negative Binomial tests of virulence factor abundance (RPKM) between healthy and asthma cohorts

## Additional Files

**Additional File 1:** Statistical Analyses for ‘The gut metagenome harbors metabolic and antibiotic resistance signatures of moderate-to-severe asthma’ knitr document

