## Supplementary figures and images for "The gut metagenome harbors metabolic and antibiotic resistance signatures of moderate-to-severe asthma"

### Supplementary Figures

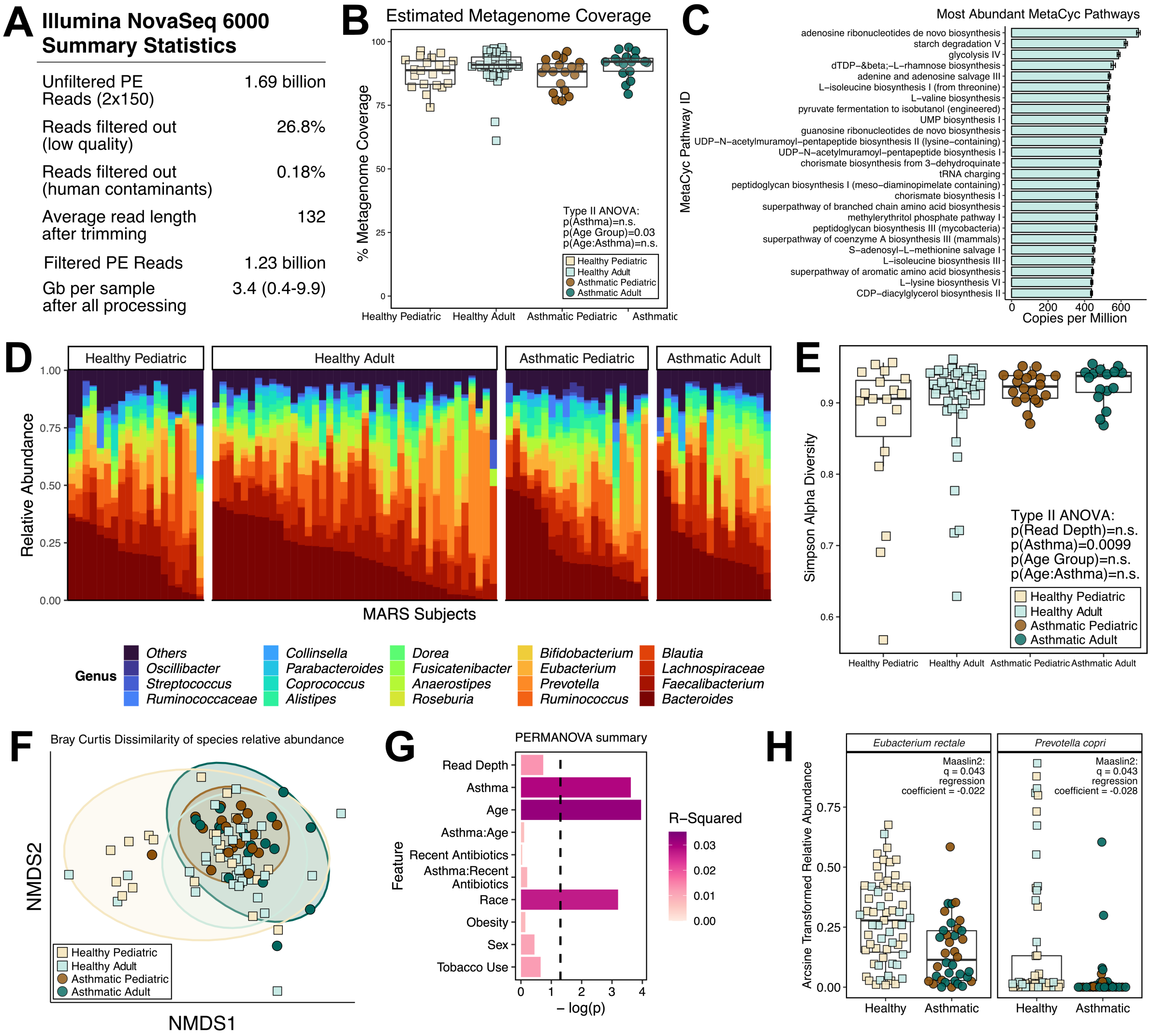

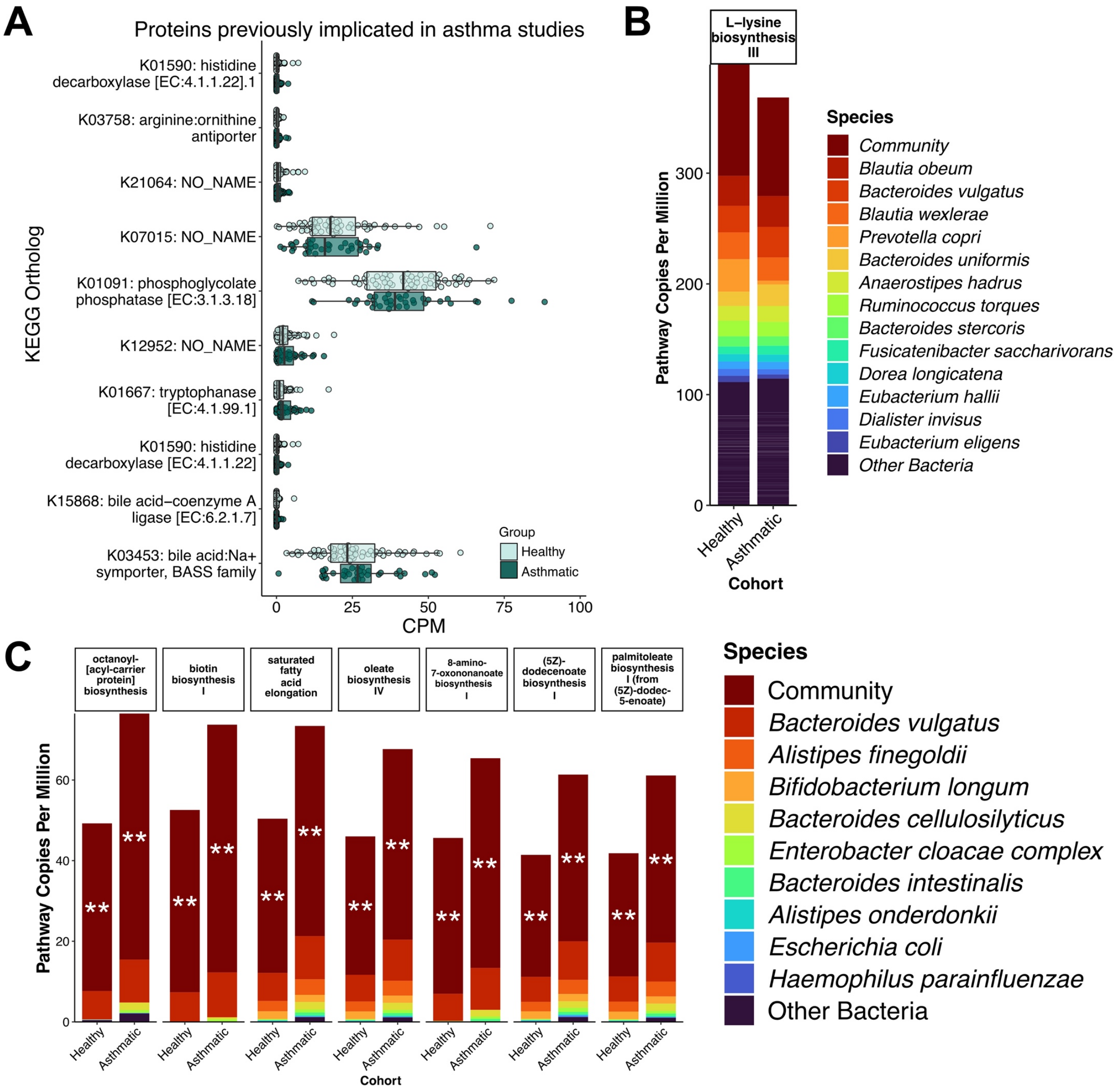

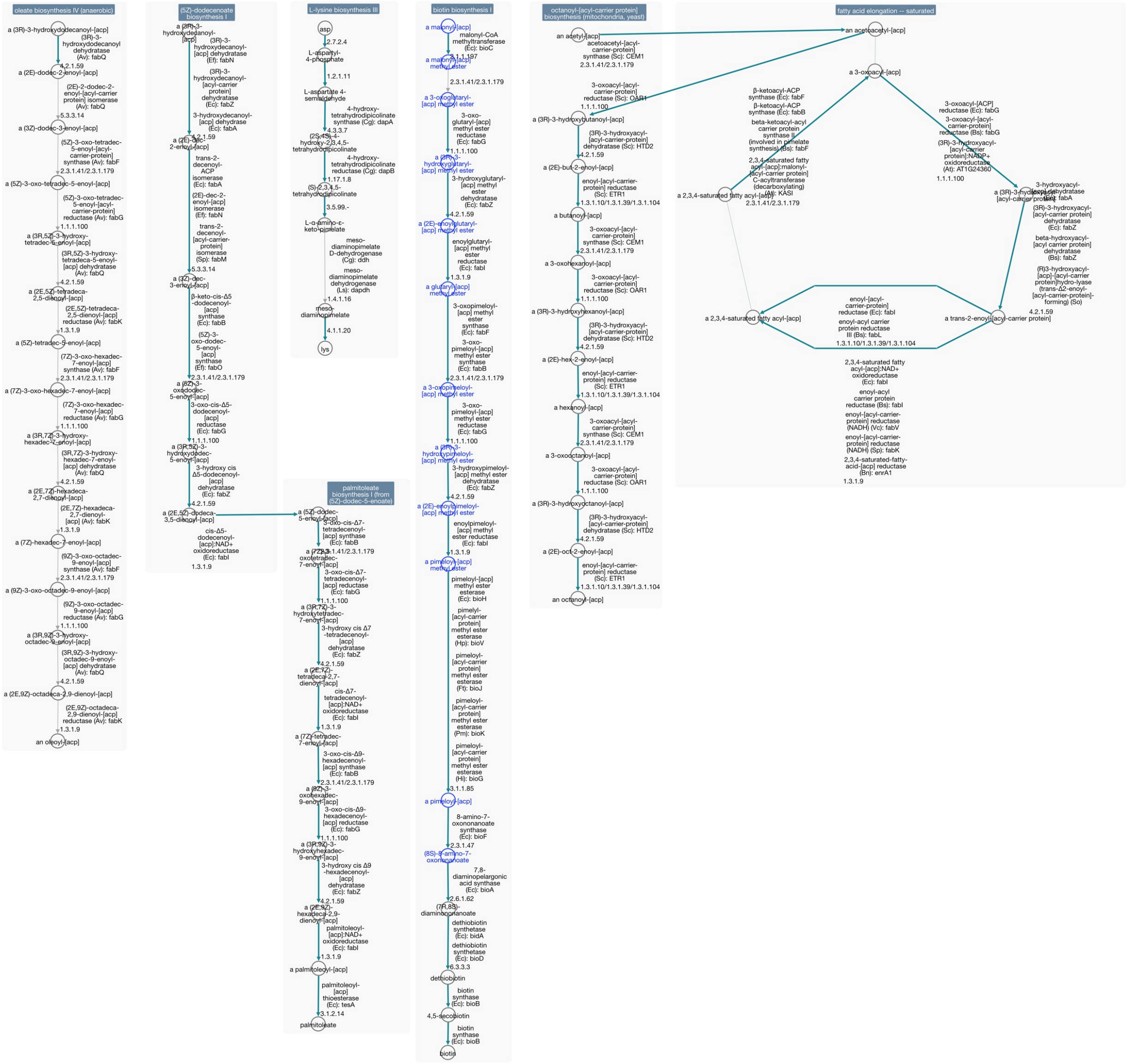

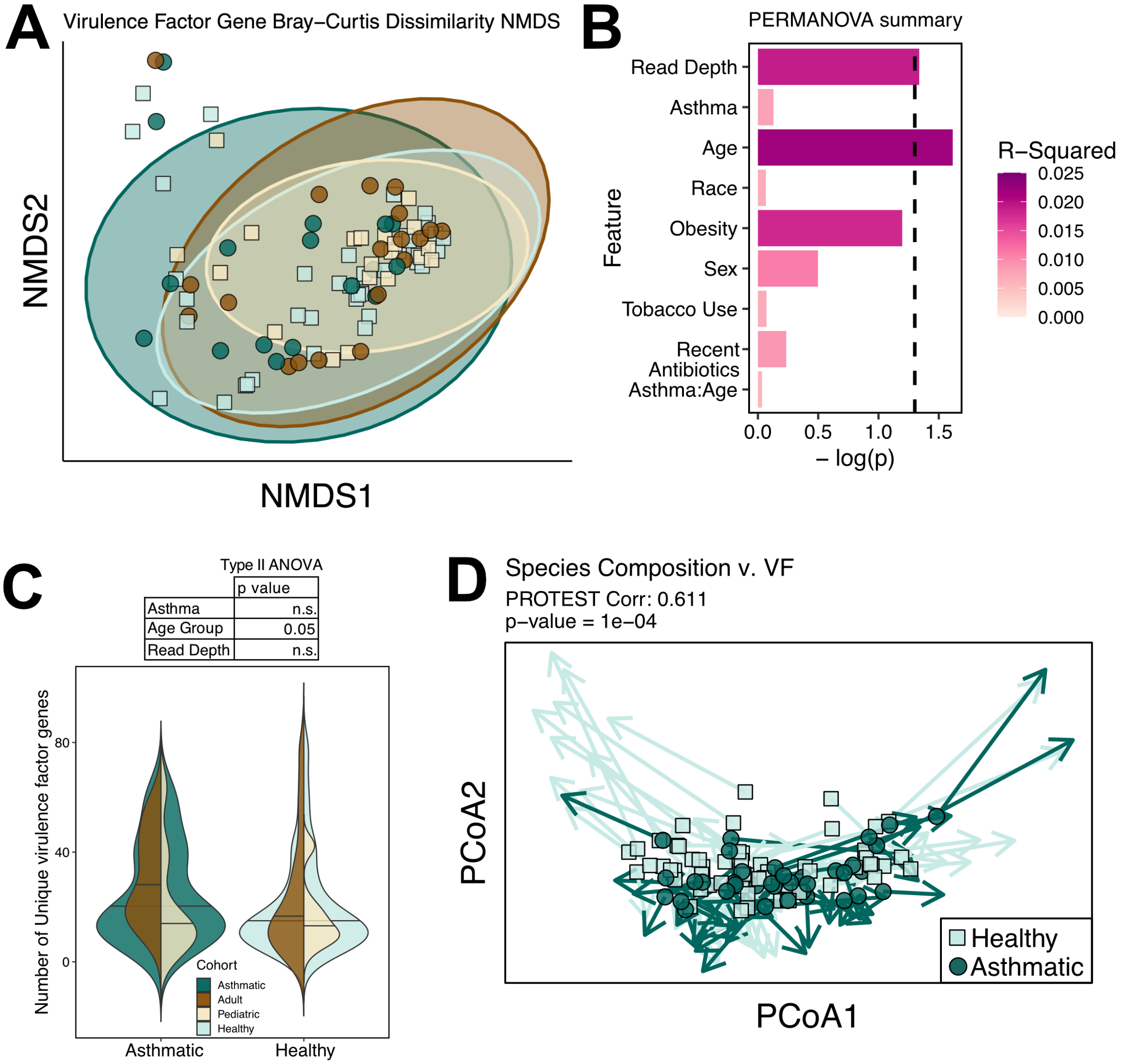

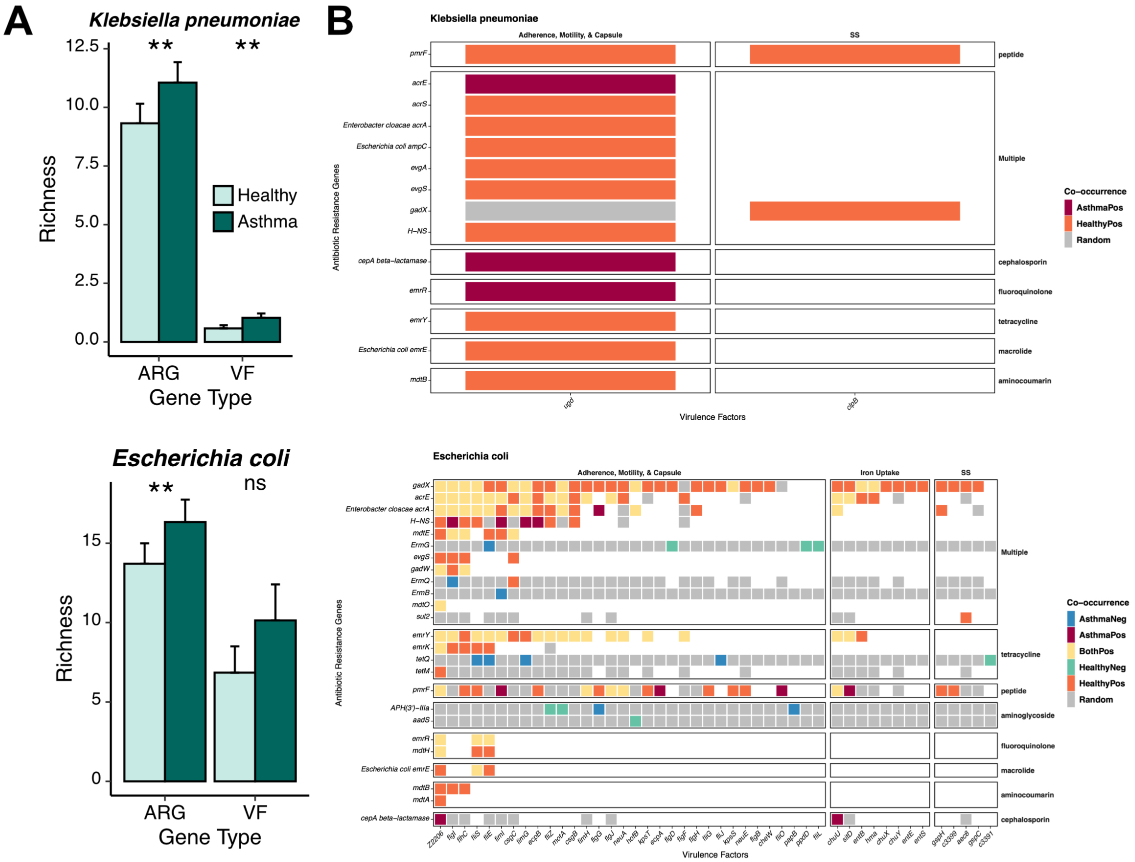
