## Additional File 1 for "The gut metagenome harbors metabolic and antibiotic resistance signatures of moderate-to-severe asthma"

Naomi G Wilson

2023-01-13

#### Functions

```
rm(list=ls())
source("./humann3_210613_functions_ngw.R")
c("hi", "hello", "world", "bye")[c("hi", "hello", "world", "bye") %notin% c("hi", "bye")]
↪  # check- notin is a function in humann3_210613_functions_ngw.R
```

```
## [1] "hello" "world"
```

```
filter_pathway_data <-
  function(indata, mymetadata, mindetect=7) {
  # drop absent genes, dropped unmapped counts
  mapped_dat <- indata[2:dim(indata)[1],] # dropped unmapped counts
  if(all(mymetadata$SampleName %in% names(mapped_dat))){
    # drop disqualifieds and exacerbations (N=95)
    mapped_dat = mapped_dat[,colnames(mapped_dat) %in% mymetadata$SampleName]
    # drop genes of zero abundance
    EmptyGenes = row.names(mapped_dat)[rowSums(mapped_dat) == 0]
    myplotdat = mapped_dat[row.names(mapped_dat) %notin% EmptyGenes,]
    # make sure there is enough data in each column (>=7 instances)
    LowGenes = row.names(myplotdat)[rowSums(ifelse(myplotdat>0, 1, 0)) < mindetect]
    myplotdat = myplotdat[row.names(myplotdat) %notin% LowGenes,]
    # also remove first two pathways since they are a catch-all bucket and skew the
    ↪  plots:
    myplotdat = myplotdat[row.names(myplotdat) %notin% "UNGROUPED",]
    myplotdat = myplotdat[row.names(myplotdat) %notin% "UNINTEGRATED",]
    return(myplotdat)
  } else {
    print("Error: sample names do not match.")
  }
  }


mean_by_group_pwys <- function(my_data = readRDS(file =
↪  "~/Library/CloudStorage/Box-Box/Kau
↪  Lab/Results/MARS/FecalMetagenomics/humann3_210613/metacyc_pwys/MARS_metacyc_pwys_CPM_filtered.rds"),
                                group_by_me="ASTHMA", # column name in humann3_mf_filtered
                                 ↪  (AgeGroup)
                               sample_mf,
                               basedir="~/Library/CloudStorage/Box-Box/Kau
                               ↪  Lab/Results/MARS/FecalMetagenomics/humann3_210613/metacyc_pwys/")
                               ↪  {
```

```
  # get mean relative abundance by specified group
  # my.plotdat <- readRDS(file=paste0(basedir, my_data_table_name, ".plotdat.rds"))
  my.plotdat <- my_data
  my.df.tmp <- as.data.frame(t(my.plotdat))
  if (is.null(sample_mf[row.names(my.df.tmp), group_by_me])) {
    print("Mapping file column not found (group_by_me argument)")
  } else {
    my.df.tmp$Group = sample_mf[row.names(my.df.tmp), group_by_me]
  }
  # tibble is supposed to be fast:
  my.tbl.tmp <- tibble::as_tibble(my.df.tmp)
  my.tbl.tmp.means <- my.tbl.tmp %>%
    group_by(Group) %>%
    summarise_all("mean")
  # make it a data frame again
  my.df.tmp.means <- as.data.frame(my.tbl.tmp.means)
  row.names(my.df.tmp.means) <- my.df.tmp.means$Group
  my.df.tmp.means$Group = NULL
  return(my.df.tmp.means)
}

renameMePathways <- function(id_number, mapping_file){
  # renames an input pathway ID (ex. COG1961) with its respective mapping file and
  ↪  outputs the ID with the human-readable name separated with underscore
  row.names(mapping_file) <- mapping_file$names_out
  name <- mapping_file[as.character(id_number), "fullname_out"]
  # print(name)
  if (is.na(name)) {
    outname <- paste0(as.character(id_number), "_", "Unknown")
  } else {
    outname <- paste0(as.character(id_number), "_", substr(name, 1,50))
  }
  return(outname)
}
```

### upload metadata from Supplemental Table 1

```
library(readxl)
nonpareildat <-
↪  data.frame(readxl::read_excel("/Users/naomiwilson/Library/CloudStorage/Box-Box/Kau
↪  Lab/MARS Paper 2/Microbiome Submission/Table S1.xlsx")) # change to reflect location
↪  of Table S1
```

```
## New names:
## * `` -> `...2`
## * `` -> `...3`
## * `` -> `...4`
## * `` -> `...5`
## * `` -> `...6`
## * `` -> `...7`
## * `` -> `...8`
## * `` -> `...9`
```

```
## * `` -> `...10`
## * `` -> `...11`
## * `` -> `...12`
## * `` -> `...13`
## * `` -> `...14`
## * `` -> `...15`
## * `` -> `...16`
## * `` -> `...17`
## * `` -> `...18`
## * `` -> `...19`
## * `` -> `...20`
## * `` -> `...21`
## * `` -> `...22`
## * `` -> `...23`
## * `` -> `...24`
## * `` -> `...25`
## * `` -> `...26`
## * `` -> `...27`
```

```
nonpareildat <- nonpareildat[4:99,c(1:10, 20,21)] # asthma, age, obesity, race, sex,
↪ recent abx, filtered read depth, nonpareil % metagenome coverage
names(nonpareildat) <-  c("SampleName", "MARSID", "ASTHMA", "AgeGroup", "sex",
↪ "weightClassBinary", "raceBinary", "ACTScoreClass", "subjectTobaccoUse",
↪ "AbxWithinYr", "readDepth", "coverage")
# nonpareildat[1,] # manually check that these ~match first row for sanity
nonpareildat <- nonpareildat[2:96,] # drop old column names
nonpareildat$coverage <- as.numeric(nonpareildat$coverage)
nonpareildat$readDepth <- as.numeric(nonpareildat$readDepth)*2 # account for both PE
↪ sister reads
nonpareildat$filenamePeriod <- paste0(gsub(x = nonpareildat$SampleName, pattern="-",
↪ replacement="."), "_interleaved_Abundance")
nonpareildat$ACTScoreClass <- ifelse(nonpareildat$ACTScoreClass=="NA" |
↪ nonpareildat$ACTScoreClass=="-", NA, nonpareildat$ACTScoreClass)

saveRDS(nonpareildat, "./humann3_mf_filtered.rds")
```

### Figure 1A:

Upload and filter CPM These are the Total Sum Scaled Reads Per Kilobase - so normalized by sequencing depth and gene size, takes minutes

```
# Upload CPM (these are the TSS normalized RPKs - so normalized by sequencing depth and
↪ gene size) takes minutes
system.time(MARSgenefamiliesCPM <- read.csv("~/Library/CloudStorage/Box-Box/Kau
↪ Lab/Results/MARS/FecalMetagenomics/humann3_210613/MARSgenefamiliesCPM_SUMMEDONLY.tsv",
↪ sep="\t"))
```

```
##    user  system elapsed
## 112.115   2.598 115.010
```

```
dim(MARSgenefamiliesCPM)
```

```
## [1] 1660518     105
```

```
row.names(MARSgenefamiliesCPM) <- MARSgenefamiliesCPM[,1]
MARSgenefamiliesCPM$X..Gene.Family = NULL
names(MARSgenefamiliesCPM) <- gsub(names(MARSgenefamiliesCPM), pattern = ".RPKs",
↪  replacement = "")

system.time(koCPM <- read.csv("~/Library/CloudStorage/Box-Box/Kau
↪  Lab/Results/MARS/FecalMetagenomics/humann3_210613/regrouped_data/genefamCPM_KO_regroup_renamed_SUMM
↪  sep="\t"))
```

```
##    user  system elapsed
##   0.292   0.010   0.305
```

```
dim(koCPM)
```

```
## [1] 4769  105
```

```
row.names(koCPM) <- koCPM[,1]
koCPM$X..Gene.Family = NULL
names(koCPM) <- samplelist <- gsub(names(koCPM), pattern = ".RPKs", replacement = "")

humann3_mf_filtered <- readRDS("./humann3_mf_filtered.rds")
# Filter data CPM - drop absent genes, drop unmapped counts (uniref90), NO min sample
↪  detection level
my_min_detect <- 0
uniref90CPM.plotdatU <-  filter_my_data(MARSgenefamiliesCPM,
↪  humann3_mf_filtered$filenamePeriod, min_detect = my_min_detect) # this is for
↪  ordination plots
uniref90CPM.plotdat <- filter_my_data(MARSgenefamiliesCPM,
↪  humann3_mf_filtered$filenamePeriod, min_detect = 16) # this is for alpha diversity
koCPM.plotdatF <- filter_my_data(koCPM, humann3_mf_filtered$filenamePeriod, min_detect =
↪  16)

# save:
saveRDS(uniref90CPM.plotdatU, file=paste0("./uniref90CPM.md", my_min_detect,
↪  ".plotdat.rds"))  # 1633505      95
saveRDS(uniref90CPM.plotdat, file="./uniref90CPM.plotdat.rds") # 348759     95
saveRDS(koCPM.plotdatF, file=paste0("./koCPM.plotdat.rds")) # 2581   95
```

Get Bray-Curtis NMDS - create matrices (this takes several minutes on macbook)

```
rm(list=ls())
source("./humann3_210613_functions_ngw.R")
library(vegan)
```

```
## Loading required package: permute

## Loading required package: lattice

## This is vegan 2.5-7
```

```
library(parallel)
my_norm_type = "CPM"
basedir = "./"
humann3_mf_filtered <- readRDS(paste0(basedir, "humann3_mf_filtered.rds"))
uniref90.plotdat <- readRDS(file=paste0(basedir, "uniref90", my_norm_type,
↪  ".md0.plotdat.rds"))
```

```
# ko.plotdat <- readRDS(file=paste0(basedir, "ko", my_norm_type, ".md0.plotdat.rds"))
MARSgenefamilies.plotdat <- uniref90.plotdat
data_table_name = paste0("uniref90", my_norm_type)
mytitle = paste0(data_table_name, " NMDS")
MARSgenefamilies.plotdat.msds <- metaMDS(t(MARSgenefamilies.plotdat), distance = "bray",
↪  k=5, try=100) # stress 0.09054304
```

```
## Square root transformation
## Wisconsin double standardization
## Run 0 stress 0.09054323
## Run 1 stress 0.09054309
## ... New best solution
## ... Procrustes: rmse 0.0002707565  max resid 0.001200488
## ... Similar to previous best
## Run 2 stress 0.09054716
## ... Procrustes: rmse 0.00107017  max resid 0.004674641
## ... Similar to previous best
## Run 3 stress 0.09054356
## ... Procrustes: rmse 0.0004166262  max resid 0.002514493
## ... Similar to previous best
## Run 4 stress 0.09054347
## ... Procrustes: rmse 0.0004610244  max resid 0.002159301
## ... Similar to previous best
## Run 5 stress 0.09054501
## ... Procrustes: rmse 0.0006138312  max resid 0.002487049
## ... Similar to previous best
## Run 6 stress 0.09054421
## ... Procrustes: rmse 0.0006370248  max resid 0.0028693
## ... Similar to previous best
## Run 7 stress 0.09054348
## ... Procrustes: rmse 0.000444947  max resid 0.001954596
## ... Similar to previous best
## Run 8 stress 0.09056997
## ... Procrustes: rmse 0.002552137  max resid 0.01125545
## Run 9 stress 0.09054333
## ... Procrustes: rmse 0.0003100846  max resid 0.001331
## ... Similar to previous best
## Run 10 stress 0.09054342
## ... Procrustes: rmse 0.0004160945  max resid 0.002242687
## ... Similar to previous best
## Run 11 stress 0.09054355
## ... Procrustes: rmse 0.0005035564  max resid 0.002618363
## ... Similar to previous best
## Run 12 stress 0.09054568
## ... Procrustes: rmse 0.0006354613  max resid 0.003502519
## ... Similar to previous best
## Run 13 stress 0.0905434
## ... Procrustes: rmse 0.0004048833  max resid 0.001922746
## ... Similar to previous best
## Run 14 stress 0.0905466
## ... Procrustes: rmse 0.0009042064  max resid 0.003606264
## ... Similar to previous best
## Run 15 stress 0.09054341
```

```
## ... Procrustes: rmse 0.0003626914  max resid 0.001802955
## ... Similar to previous best
## Run 16 stress 0.09054311
## ... Procrustes: rmse 0.0003926808  max resid 0.00244835
## ... Similar to previous best
## Run 17 stress 0.09054339
## ... Procrustes: rmse 0.0004461252  max resid 0.002907603
## ... Similar to previous best
## Run 18 stress 0.09054319
## ... Procrustes: rmse 0.0003333048  max resid 0.002075319
## ... Similar to previous best
## Run 19 stress 0.09054305
## ... New best solution
## ... Procrustes: rmse 0.0002878917  max resid 0.001619853
## ... Similar to previous best
## Run 20 stress 0.0905445
## ... Procrustes: rmse 0.0006898169  max resid 0.004531119
## ... Similar to previous best
## Run 21 stress 0.09054334
## ... Procrustes: rmse 0.0002854261  max resid 0.001679377
## ... Similar to previous best
## Run 22 stress 0.0905571
## ... Procrustes: rmse 0.001569886  max resid 0.008949549
## ... Similar to previous best
## Run 23 stress 0.09057263
## ... Procrustes: rmse 0.002488899  max resid 0.01490885
## Run 24 stress 0.09054413
## ... Procrustes: rmse 0.0006009875  max resid 0.002740686
## ... Similar to previous best
## Run 25 stress 0.09054365
## ... Procrustes: rmse 0.0004234874  max resid 0.003014502
## ... Similar to previous best
## Run 26 stress 0.09055264
## ... Procrustes: rmse 0.001109701  max resid 0.004548554
## ... Similar to previous best
## Run 27 stress 0.0905439
## ... Procrustes: rmse 0.0003072173  max resid 0.001388834
## ... Similar to previous best
## Run 28 stress 0.09054374
## ... Procrustes: rmse 0.0004175526  max resid 0.002753145
## ... Similar to previous best
## Run 29 stress 0.09054391
## ... Procrustes: rmse 0.0003761491  max resid 0.002458125
## ... Similar to previous best
## Run 30 stress 0.09197773
## Run 31 stress 0.09054364
## ... Procrustes: rmse 0.000493536  max resid 0.002184186
## ... Similar to previous best
## Run 32 stress 0.09054415
## ... Procrustes: rmse 0.0006030159  max resid 0.002785551
## ... Similar to previous best
## Run 33 stress 0.09054336
## ... Procrustes: rmse 0.0003033596  max resid 0.001721624
## ... Similar to previous best
```

```
## Run 34 stress 0.09054391
## ... Procrustes: rmse 0.0004425779  max resid 0.003006805
## ... Similar to previous best
## Run 35 stress 0.09054491
## ... Procrustes: rmse 0.0006055425  max resid 0.003354325
## ... Similar to previous best
## Run 36 stress 0.09054404
## ... Procrustes: rmse 0.0005887122  max resid 0.003768173
## ... Similar to previous best
## Run 37 stress 0.09054315
## ... Procrustes: rmse 0.0001236281  max resid 0.0008795886
## ... Similar to previous best
## Run 38 stress 0.09054393
## ... Procrustes: rmse 0.0005589986  max resid 0.003370057
## ... Similar to previous best
## Run 39 stress 0.09055162
## ... Procrustes: rmse 0.001322724  max resid 0.007631554
## ... Similar to previous best
## Run 40 stress 0.09054442
## ... Procrustes: rmse 0.0003859695  max resid 0.001521933
## ... Similar to previous best
## Run 41 stress 0.09054404
## ... Procrustes: rmse 0.0004329603  max resid 0.003108062
## ... Similar to previous best
## Run 42 stress 0.09054416
## ... Procrustes: rmse 0.000506906  max resid 0.00244511
## ... Similar to previous best
## Run 43 stress 0.09059321
## ... Procrustes: rmse 0.002654721  max resid 0.0205635
## Run 44 stress 0.09054431
## ... Procrustes: rmse 0.0003683709  max resid 0.001582735
## ... Similar to previous best
## Run 45 stress 0.09054414
## ... Procrustes: rmse 0.0004488877  max resid 0.003481846
## ... Similar to previous best
## Run 46 stress 0.09054409
## ... Procrustes: rmse 0.0004625503  max resid 0.00326757
## ... Similar to previous best
## Run 47 stress 0.09054549
## ... Procrustes: rmse 0.0005126802  max resid 0.003261779
## ... Similar to previous best
## Run 48 stress 0.09055479
## ... Procrustes: rmse 0.001546119  max resid 0.007053544
## ... Similar to previous best
## Run 49 stress 0.09057403
## ... Procrustes: rmse 0.002249959  max resid 0.009815433
## ... Similar to previous best
## Run 50 stress 0.0905436
## ... Procrustes: rmse 0.0003475573  max resid 0.002512204
## ... Similar to previous best
## Run 51 stress 0.09054358
## ... Procrustes: rmse 0.0004844256  max resid 0.003380333
## ... Similar to previous best
## Run 52 stress 0.09054318
```

```
## ... Procrustes: rmse 0.0003273211  max resid 0.00219276
## ... Similar to previous best
## Run 53 stress 0.09054347
## ... Procrustes: rmse 0.0001971245  max resid 0.0007997918
## ... Similar to previous best
## Run 54 stress 0.0943834
## Run 55 stress 0.09054404
## ... Procrustes: rmse 0.0005616457  max resid 0.004093378
## ... Similar to previous best
## Run 56 stress 0.09054308
## ... Procrustes: rmse 0.0002111858  max resid 0.001085766
## ... Similar to previous best
## Run 57 stress 0.09054396
## ... Procrustes: rmse 0.00034991  max resid 0.001464794
## ... Similar to previous best
## Run 58 stress 0.09054486
## ... Procrustes: rmse 0.0005654215  max resid 0.003501417
## ... Similar to previous best
## Run 59 stress 0.09054384
## ... Procrustes: rmse 0.0005140804  max resid 0.003280059
## ... Similar to previous best
## Run 60 stress 0.09054396
## ... Procrustes: rmse 0.0002343271  max resid 0.001403422
## ... Similar to previous best
## Run 61 stress 0.09054344
## ... Procrustes: rmse 0.0004443673  max resid 0.001979679
## ... Similar to previous best
## Run 62 stress 0.09054436
## ... Procrustes: rmse 0.000420069  max resid 0.001861278
## ... Similar to previous best
## Run 63 stress 0.09054369
## ... Procrustes: rmse 0.0004029163  max resid 0.002651016
## ... Similar to previous best
## Run 64 stress 0.09054385
## ... Procrustes: rmse 0.0003821255  max resid 0.001956409
## ... Similar to previous best
## Run 65 stress 0.09054411
## ... Procrustes: rmse 0.0004439165  max resid 0.002413479
## ... Similar to previous best
## Run 66 stress 0.09091681
## ... Procrustes: rmse 0.00432462  max resid 0.01796912
## Run 67 stress 0.09054417
## ... Procrustes: rmse 0.0005841531  max resid 0.003801773
## ... Similar to previous best
## Run 68 stress 0.09054386
## ... Procrustes: rmse 0.0003866456  max resid 0.001859779
## ... Similar to previous best
## Run 69 stress 0.09054425
## ... Procrustes: rmse 0.0005288185  max resid 0.003259484
## ... Similar to previous best
## Run 70 stress 0.09054331
## ... Procrustes: rmse 0.0004071596  max resid 0.001945681
## ... Similar to previous best
## Run 71 stress 0.09054406
```

```
## ... Procrustes: rmse 0.0005991234  max resid 0.004107377
## ... Similar to previous best
## Run 72 stress 0.09267917
## Run 73 stress 0.09054317
## ... Procrustes: rmse 0.0003706823  max resid 0.002075508
## ... Similar to previous best
## Run 74 stress 0.09054373
## ... Procrustes: rmse 0.0003067456  max resid 0.001747832
## ... Similar to previous best
## Run 75 stress 0.09054386
## ... Procrustes: rmse 0.0002672777  max resid 0.00153704
## ... Similar to previous best
## Run 76 stress 0.09055098
## ... Procrustes: rmse 0.001172031  max resid 0.007745772
## ... Similar to previous best
## Run 77 stress 0.09054361
## ... Procrustes: rmse 0.0002513514  max resid 0.001464982
## ... Similar to previous best
## Run 78 stress 0.09054348
## ... Procrustes: rmse 0.0003141648  max resid 0.002109032
## ... Similar to previous best
## Run 79 stress 0.09054428
## ... Procrustes: rmse 0.0005557775  max resid 0.003800624
## ... Similar to previous best
## Run 80 stress 0.09054329
## ... Procrustes: rmse 0.0002925033  max resid 0.001304578
## ... Similar to previous best
## Run 81 stress 0.09054417
## ... Procrustes: rmse 0.0004861994  max resid 0.003557013
## ... Similar to previous best
## Run 82 stress 0.09093087
## ... Procrustes: rmse 0.008355134  max resid 0.03091923
## Run 83 stress 0.09054367
## ... Procrustes: rmse 0.0002676707  max resid 0.001509128
## ... Similar to previous best
## Run 84 stress 0.09054339
## ... Procrustes: rmse 0.0002708193  max resid 0.001180398
## ... Similar to previous best
## Run 85 stress 0.09054447
## ... Procrustes: rmse 0.0005414993  max resid 0.00323834
## ... Similar to previous best
## Run 86 stress 0.09054456
## ... Procrustes: rmse 0.0006397422  max resid 0.004118343
## ... Similar to previous best
## Run 87 stress 0.09054398
## ... Procrustes: rmse 0.0004791164  max resid 0.002771346
## ... Similar to previous best
## Run 88 stress 0.09054695
## ... Procrustes: rmse 0.0008614077  max resid 0.004264517
## ... Similar to previous best
## Run 89 stress 0.090544
## ... Procrustes: rmse 0.0005919167  max resid 0.002352417
## ... Similar to previous best
## Run 90 stress 0.09054388
```

```
## ... Procrustes: rmse 0.0002722268  max resid 0.001175299
## ... Similar to previous best
## Run 91 stress 0.09054393
## ... Procrustes: rmse 0.0003224258  max resid 0.001396842
## ... Similar to previous best
## Run 92 stress 0.09054355
## ... Procrustes: rmse 0.0004882498  max resid 0.002689576
## ... Similar to previous best
## Run 93 stress 0.09054495
## ... Procrustes: rmse 0.0007525662  max resid 0.003195707
## ... Similar to previous best
## Run 94 stress 0.09054319
## ... Procrustes: rmse 0.0001566802  max resid 0.001162561
## ... Similar to previous best
## Run 95 stress 0.09054332
## ... Procrustes: rmse 0.0002065387  max resid 0.0007840679
## ... Similar to previous best
## Run 96 stress 0.09054396
## ... Procrustes: rmse 0.0003586862  max resid 0.002101296
## ... Similar to previous best
## Run 97 stress 0.09054388
## ... Procrustes: rmse 0.0004617846  max resid 0.00289609
## ... Similar to previous best
## Run 98 stress 0.09054451
## ... Procrustes: rmse 0.0004688597  max resid 0.003262515
## ... Similar to previous best
## Run 99 stress 0.09054471
## ... Procrustes: rmse 0.0005826958  max resid 0.0041989
## ... Similar to previous best
## Run 100 stress 0.09054414
## ... Procrustes: rmse 0.0003218905  max resid 0.001997528
## ... Similar to previous best
## *** Solution reached
```

```
saveRDS(MARSgenefamilies.plotdat.msds, paste0(basedir, data_table_name, ".msds.rds"))
```

uniref90 cpm nmds

```
library(ggplot2)
library(vegan)
library(ggfortify)
library(RColorBrewer)

basedir = "./"
data_table_name = "uniref90CPM"
data.msds <- readRDS(paste0(basedir, data_table_name, ".msds.rds"))
uniref90CPM.plotdat <- readRDS(file="./uniref90CPM.md0.plotdat.rds")
humann3_mf <- readRDS(file = paste0(basedir, "humann3_mf_filtered.rds"))
humann3_mf <- humann3_mf[rowSums(is.na(humann3_mf)) < dim(humann3_mf)[2],]
dim(humann3_mf)[1] == 95
```

```
## [1] TRUE
```

```
row.names(humann3_mf) <- humann3_mf$filenamePeriod
```

```
mymetadata <- humann3_mf
humann3_mf$AgeGroup <- factor(humann3_mf$AgeGroup)
humann3_mf$ASTHMA <- factor(humann3_mf$ASTHMA)
my_data_table_name <- data_table_name
mytitle = "Bray-Curtis NMDS"
asthma_color_hex= brewer.pal(7, "BrBG")[7] #"#DE4968FF",
healthy_color_hex= brewer.pal(7, "BrBG")[5] #"#51127CFF",
confidenceEllipse = 0.95

brayCurtisMDS <- data.frame(data.msds$points)
brayCurtisMDS$AgeGroup <-
↪  factor(mymetadata[as.character(row.names(brayCurtisMDS)),"AgeGroup"])
brayCurtisMDS$Asthma <-
↪  factor(mymetadata[as.character(row.names(brayCurtisMDS)),"ASTHMA"])
# 4 grouper ####
brayCurtisMDS$Cohort <- factor(paste0(brayCurtisMDS$Asthma, brayCurtisMDS$AgeGroup))
my_color_scheme = c(brewer.pal(7, "BrBG")[c(7,1,5,3)])
# library(viridis)
betaDivPlot4 <- ggplot(data = brayCurtisMDS,
                        aes(x = MDS1, y = MDS2,
                            fill = Cohort,
                            color = Cohort,
                            shape = Asthma)) +
  stat_ellipse(aes(x=MDS1, y=MDS2, color=Cohort, group=Cohort), show.legend = TRUE, type
  ↪  = "t", level = confidenceEllipse, geom = "polygon", alpha = 0.4, size = 1)+
  # geom_point(size = 6, shape = 21, color ="black") +
  geom_point(size = 4.5, color = "black", alpha=0.8) +
    # geom_point(show.legend = FALSE, aes(brayCurtisMDS["MARS0043.F.16s", "MDS1"],
    ↪  brayCurtisMDS["MARS0043.F.16s", "MDS2"]),
    #            colour = asthma_color_hex, pch=5, size =8) +
    # geom_point(show.legend = FALSE, aes(brayCurtisMDS["MARS0022.F.16s", "MDS1"],
    ↪  brayCurtisMDS["MARS0022.F.16s", "MDS2"]),
    #            colour = healthy_color_hex, pch=5, size =8) +
    # scale_fill_viridis(breaks = levels(brayCurtisMDS$Cohort), discrete = TRUE, alpha =
    ↪  1,
    #                    labels = gsub(pattern = " ", replacement = "\n",
    #                                  x = levels(brayCurtisMDS$Cohort))) +
    # scale_fill_viridis(breaks = levels(brayCurtisMDS$Cohort), discrete = TRUE, alpha =
    ↪  1,
    #                    labels = gsub(pattern = " ", replacement = "\n",
    #                                  x = levels(brayCurtisMDS$Cohort))) +
    # scale_color_viridis(discrete = TRUE, alpha = 0.4) +
    scale_color_manual(values = my_color_scheme) +
    scale_fill_manual(values = my_color_scheme) +
    scale_shape_manual(values=c(21, 22)) +
    theme_classic() +
    xlab(label = "NMDS1") + ylab("NMDS2") + ggtitle(mytitle) +

    # geom_point(aes(shape=AgeGroup, color="black", size=6)) +
    # scale_shape_manual(values=c(21, 22)) +

    theme(axis.title.x = element_text(size = 21, vjust = -0.9),
          axis.title.y = element_text(size = 21, vjust = 1),
```

```
        axis.text.x = element_blank(),
        axis.ticks = element_blank(),
        axis.text.y = element_blank(),
        legend.text = element_text(size = 16),
        legend.title = element_text(size = 16, face = "bold"),
        legend.position = "bottom", legend.box = "horizontal")#,
# plot.margin = margin(b = .9, l = 0.9, t = 1, r = 1, unit = "cm"))
betaDivPlot4
```

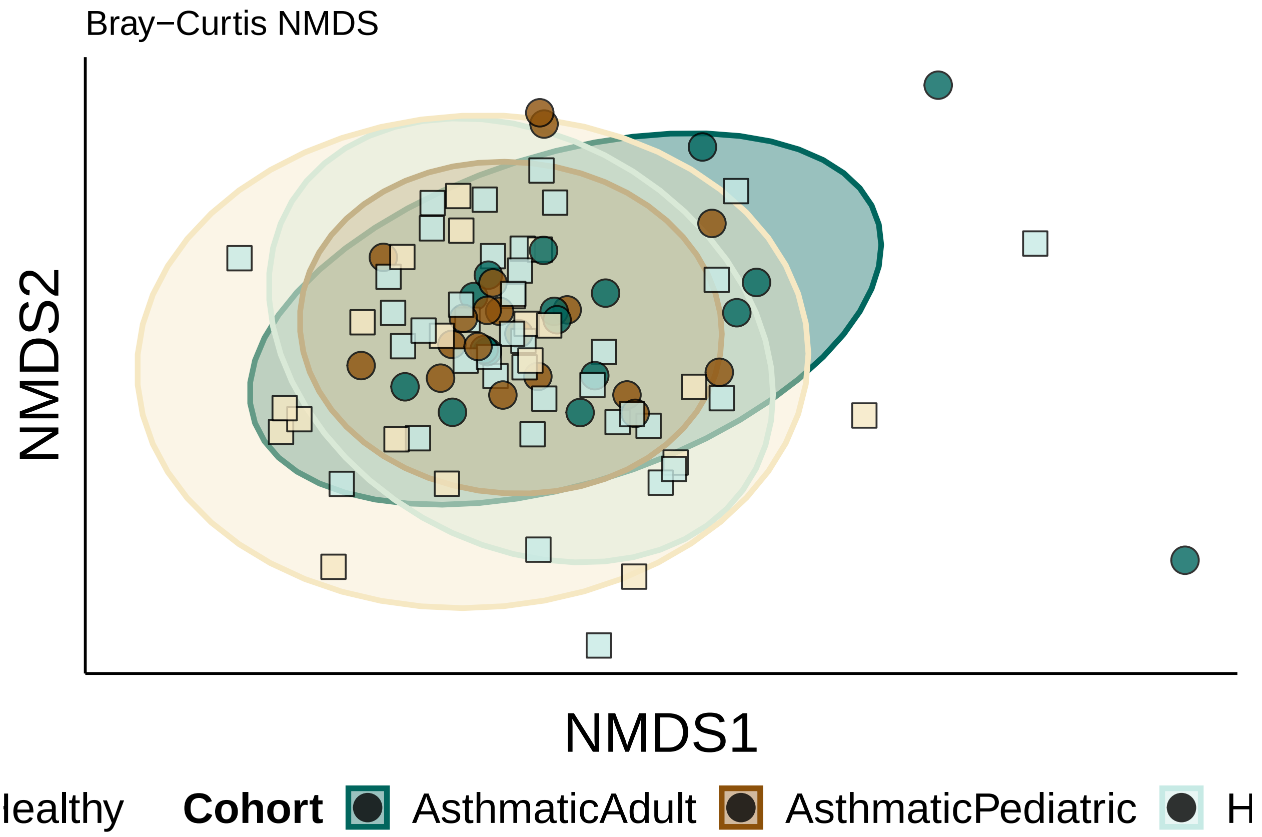


```
# ggsave(plot=betaDivPlot4, filename = paste0(basedir, my_data_table_name,
↪   "_4Group_NMDS_1and2.jpeg"), device = "jpg", width = 5, height = 5, units = "in")
# ggsave(plot=betaDivPlot4, filename = paste0(basedir, my_data_table_name,
↪   "_4Group_NMDS_1and2.pdf"), device = "pdf", width = 5, height = 5, units = "in",
↪   useDingbats = FALSE)
```

#### Figure 1B:

Uniref90 CPM PERMANOVA Bray-Curtis distances barplot

```
source("./humann3_210613_functions_ngw.R")
library(ggplot2)
library(vegan)
library(RColorBrewer)

binaryTrue = FALSE
my_norm_type = "CPM"
```

```
uniref90.plotdat <- readRDS(file=paste0("./uniref90", my_norm_type, ".md0.plotdat.rds"))
# unire90.braydist = readRDS(file="./unire90CPM.braydist.rds")
unire90.braydist <- vegdist(t(uniref90.plotdat), method = "bray", binary=binaryTrue)
pca.dist = unire90.braydist
distance_metric = "BrayCurtis"
gene_analysis_name = "unire90CPM"
basedir="./"

humann3_mf_filtered_perm <- readRDS(file = paste0(basedir, "humann3_mf_filtered.rds"))
row.names(humann3_mf_filtered_perm) <- humann3_mf_filtered_perm$filenamePeriod
# humann3_mf_filtered <- humann3_mf_filtered_perm[as.character(colnames(ko.plotdat)),]
humann3_mf_filtered <-
↪  humann3_mf_filtered_perm[as.character(colnames(uniref90.plotdat)),]
adonisAllIndpRaceBinaryPATHWAYS <- adonis(pca.dist ~
↪  readDepth+ASTHMA+AgeGroup+raceBinary+weightClassBinary+sex+subjectTobaccoUse+ASTHMA*AgeGroup+AbxWitl
                        data = humann3_mf_filtered,
                        permutations = 100000) # sequential permanova
# make table
adonisAllIndpRaceBinaryPATHWAYSTable <-
↪  data.frame(adonisAllIndpRaceBinaryPATHWAYS$aov.tab[c("R2","Pr(>F)")])
names(adonisAllIndpRaceBinaryPATHWAYSTable) <- c("R2", "pval")
adonisAllIndpRaceBinaryPATHWAYSTable$features <-
↪  row.names(adonisAllIndpRaceBinaryPATHWAYSTable)
adonisAllIndpRaceBinaryPATHWAYSTable
↪  <-adonisAllIndpRaceBinaryPATHWAYSTable[!adonisAllIndpRaceBinaryPATHWAYSTable$features
↪  %in% c("Residuals","Total"),]
adonisAllIndpRaceBinaryPATHWAYSTable$pval <-
↪  round(adonisAllIndpRaceBinaryPATHWAYSTable$pval, 5)
adonisAllIndpRaceBinaryPATHWAYSTable$R2 <- round(adonisAllIndpRaceBinaryPATHWAYSTable$R2,
↪  3)
# attr(adonisAllIndpRaceBinaryPATHWAYSTable$R2, "label") <- "R-Squared"
row.names(adonisAllIndpRaceBinaryPATHWAYSTable) <- NULL
adonisAllIndpRaceBinaryPATHWAYSTable <- adonisAllIndpRaceBinaryPATHWAYSTable[,c(3,1,2)]
write.table(adonisAllIndpRaceBinaryPATHWAYSTable, file = paste0(basedir,
↪  gene_analysis_name, "_", distance_metric,
↪  "_adonisAllIndpRaceBinaryweightClassBinary10000.tsv"), sep="\t", row.names = FALSE,
↪  col.names = TRUE)
adonisAllIndpRaceBinaryPATHWAYSTable$neglogpval <-
↪  -log(as.numeric(adonisAllIndpRaceBinaryPATHWAYSTable$pval), base = 10)
adonisAllIndpRaceBinaryPATHWAYSTable$features <-
↪  factor(adonisAllIndpRaceBinaryPATHWAYSTable$features, levels=rev(c("readDepth",
↪  "ASTHMA", "AgeGroup", "raceBinary", "weightClassBinary",  "sex",
↪  "subjectTobaccoUse", "ASTHMA:AgeGroup", "AbxWithinYr")), labels = rev(c("Read Depth",
↪  "Asthma", "Age", "Race", "Obesity", "Sex", "Tobacco Use", "Asthma:Age",
↪  "Recent\nAntibiotics"))) # plot them in order of the permanova input
AdonisBarplotForPaperBinaryRacePATHWAYS <- makeAdonisBarplot(adonis_dat =
↪  adonisAllIndpRaceBinaryPATHWAYSTable, y_colname = "neglogpval", features_colname =
↪  "features", fill_colname = "R2",
↪  max_fill_value=max(adonisAllIndpRaceBinaryPATHWAYSTable$R2)*1.1)
print(AdonisBarplotForPaperBinaryRacePATHWAYS)
```

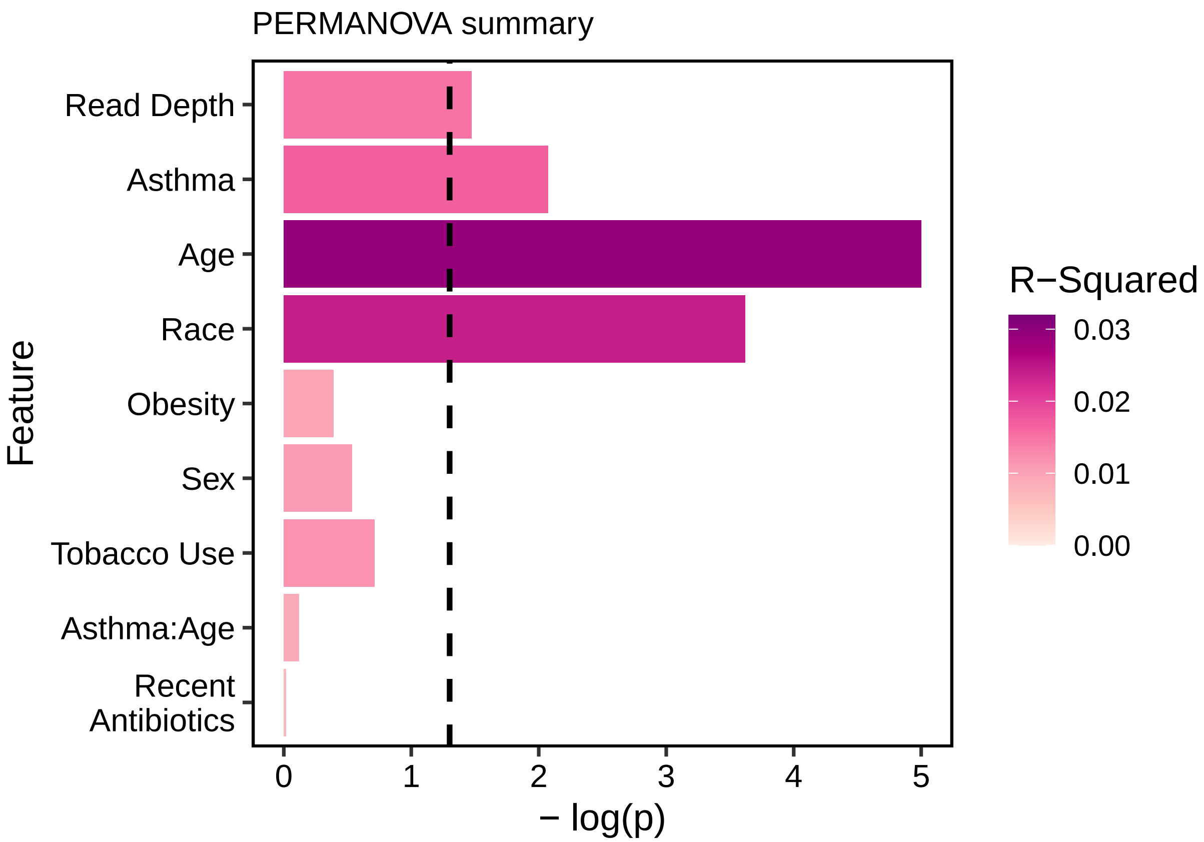


```
# ggsave(AdonisBarplotForPaperBinaryRacePATHWAYS,
#        filename = paste0(basedir, gene_analysis_name, "_", distance_metric,
↪  "_adonis100000.pdf"),
#        height = 4, width = 5,
#        units = "in",
#        device = "pdf", useDingbats=FALSE)
#
# ggsave(AdonisBarplotForPaperBinaryRacePATHWAYS,
#        filename = paste0(basedir, gene_analysis_name, "_", distance_metric,
↪  "_adonis100000.jpg"),
#        height = 4, width = 5,
#        units = "in",
#        device = "jpg")
```

#### Figure 1C:

Alpha Diversity/Richness per cohort on filtered data

```
rm(list = ls())
library(ggpubr)
library(ggplot2)
library(ggsignif)
library(ggbeeswarm)
library(RColorBrewer)
source("./humann3_210613_functions_ngw.R")
```

```r
basedir="./"
getwd()
```

```
## [1] "/Users/naomiwilson/Library/CloudStorage/Box-Box/Kau Lab/MARS Paper 2/Microbiome Submission/Shar
```

```r
my.plotdat <- readRDS(file=paste0(basedir, "uniref90CPM", ".plotdat.rds")) # filtered
↪  data
# check filtering:
sum(rowSums(my.plotdat>0)<16) == 0
```

```
## [1] TRUE
```

```r
library(readxl)
humann3_mf_filtered <- readRDS("humann3_mf_filtered.rds")
row.names(humann3_mf_filtered) <- humann3_mf_filtered$filenamePeriod
humann3_mf = humann3_mf_filtered
my.plotdat[my.plotdat>0]=1 # make binary
if(sum(rowSums(my.plotdat) < 10)){
  print("pls drop with sparse data before doing alpha div")
}
my.plotdat <- colSums(my.plotdat)
plotmypwys <- data.frame(t(rbind(my.plotdat,
↪  as.character(humann3_mf[as.character(names(my.plotdat)), "ASTHMA"]))))
names(plotmypwys) <- c("Richness", "Group")
plotmypwys$Richness <- as.numeric(as.character(plotmypwys$Richness))
plotmypwys$Richness <- plotmypwys$Richness/1000
plotmypwys$Group <- factor(plotmypwys$Group, levels=c("Healthy", "Asthmatic"))
plotmypwys$AgeGroup <- as.character(humann3_mf[as.character(row.names(plotmypwys)),
↪  "AgeGroup"])
plotmypwys$readDepth <- as.numeric(humann3_mf[as.character(row.names(plotmypwys)),
↪  "readDepth"])
plotmypwys$coverage <- as.numeric(humann3_mf[as.character(row.names(plotmypwys)),
↪  "coverage"])

library(car)
```

```
## Loading required package: carData
```

```r
Anova(lm(Richness ~Group*readDepth+Group*AgeGroup, data = plotmypwys),type=2)
```

```
## Anova Table (Type II tests)
##
## Response: Richness
##                   Sum Sq Df F value    Pr(>F)
## Group               1174  1  1.4679    0.2289
## readDepth          35875  1 44.8636 1.844e-09 ***
## AgeGroup             318  1  0.3976    0.5300
## Group:readDepth        1  1  0.0008    0.9772
## Group:AgeGroup      1643  1  2.0552    0.1552
## Residuals          71169 89
## ---
## Signif. codes:  0 '***' 0.001 '**' 0.01 '*' 0.05 '.' 0.1 ' ' 1
```

```r
# Anova(lm(Richness ~Group*coverage+Group*AgeGroup, data = plotmypwys),type=2)
```

```r
# VIOLIN PLOTS
source("./split_violins_ggplot.R")
Uniref90richnessViolin <- ggplot(data = plotmypwys, aes(x=Group, y=Richness, fill =
↪  interaction(Group,AgeGroup))) +
  geom_violin(aes(x=Group, y=Richness, fill=Group), draw_quantiles = c(0.5), inherit.aes
  ↪  = FALSE, scale="width", trim=FALSE, alpha=0.8, show.legend = T) + # draw_quantiles
  ↪  = c(0.5), c(0.25, 0.5, 0.75)
  geom_split_violin(draw_quantiles = c(0.5), scale="width", width=0.5, trim=FALSE,
  ↪  alpha=0.8, show.legend = T) +  # add this to draw quantiles: draw_quantiles =
  ↪  c(0.25, 0.5, 0.75)
  theme_classic() +
  scale_fill_manual(name="Cohort",
                    values = c("#01665E", brewer.pal(7, "BrBG")[1], brewer.pal(7,
                    ↪  "BrBG")[3],
                               "#C7EAE5", brewer.pal(7, "BrBG")[1], brewer.pal(7,
                               ↪  "BrBG")[3]),
                    labels = c('Asthmatic','Adult', 'Pediatric',
                               'Healthy', 'Adult', 'Pediatric'))+ # color order:
                               ↪  A,AA,AP,H,HA,HP
  theme(panel.grid.major = element_blank(),
        panel.grid.minor = element_blank(),
        axis.line.x = element_line(size = 0, colour = "black"),
        axis.line.y = element_line(size = 0, colour = "black"),
        axis.line = element_line(size=1, colour = "black"),
        panel.border = element_rect(fill = NA, colour = "black", size = 1),
        panel.background = element_blank(),
        text=element_text(size = 16),
        axis.text.x=element_text(colour="black", size = 12),
        axis.text.y=element_text(colour="black", size = 12),
        axis.title.x=element_blank())+
        # legend.position = "none")+
  ylab("Number of Unique Uniref90 clusters")

Uniref90richnessViolin
```

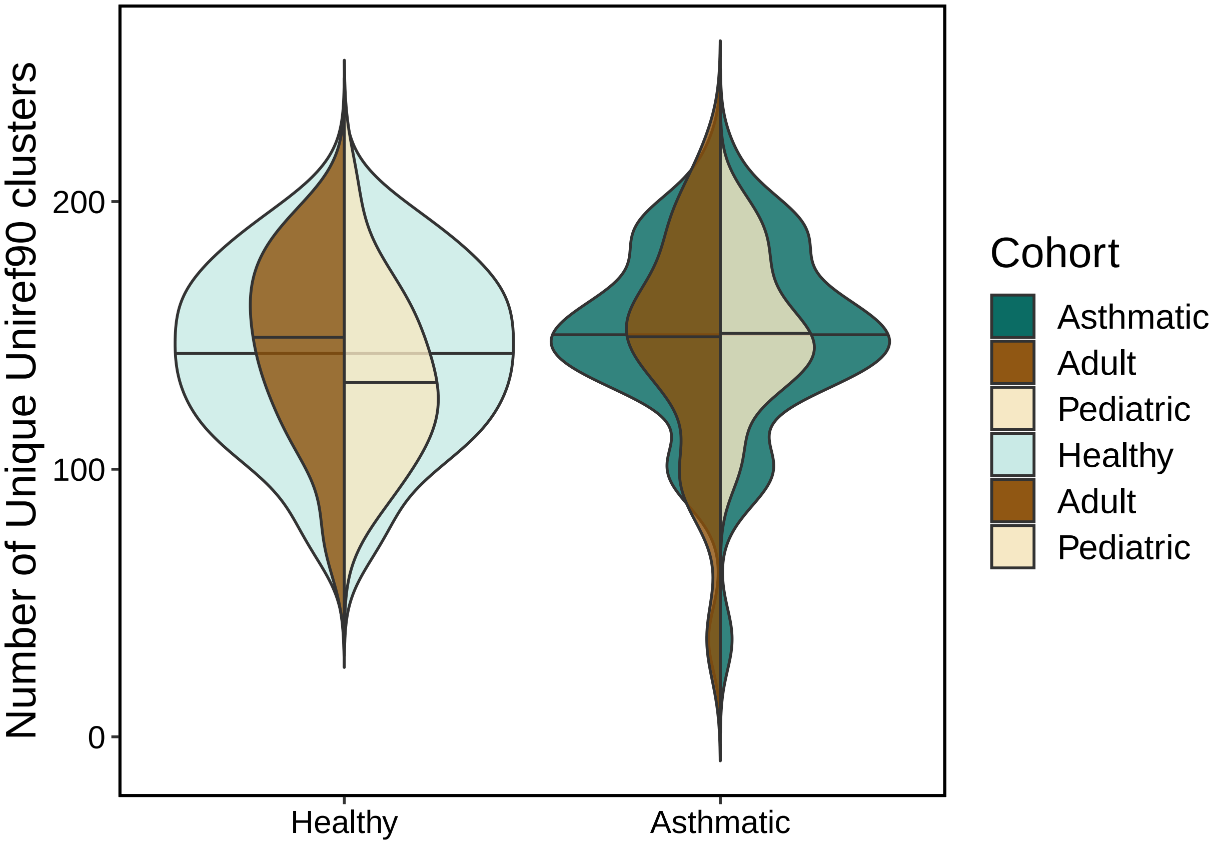


```
# ggsave(plot = Uniref90richnessViolin, filename =
↪  "./Uniref90_Richness_doubleviolin.pdf", width = 6, height = 6, units = "in",
↪  useDingbats = FALSE)
# ggsave(filename = "./Uniref90_Richness_doubleviolin.jpeg", plot =
↪  Uniref90richnessViolin, units = "in", width =6, height = 6, device = "jpg")
```

#### Figure S1B:

nonpareil coverage

```
rm(list=ls())
source("./humann3_210613_functions_ngw.R")
library(RColorBrewer)

datframe<- readRDS("humann3_mf_filtered.rds")

# 4 groups
datframe$cohort <- factor(paste0(datframe$ASTHMA, " ", datframe$AgeGroup),
↪  levels=c("Healthy Pediatric", "Healthy Adult", "Asthmatic Pediatric", "Asthmatic
↪  Adult"))
ageasthmacoverage <- ggplot(datframe, aes(x=cohort, y=coverage)) +
  ggbeeswarm::geom_quasirandom(aes(shape=cohort, fill=cohort), size=3.5, alpha=0.9)+
    geom_boxplot(outlier.shape=NA, alpha=0.2) +
    scale_fill_manual(values = c(brewer.pal(7, "BrBG")[c(3,5,1,7)]))+
    scale_shape_manual(values=c(22, 22, 21, 21)) +
    scale_color_manual(values=c("black", "black","black", "black")) +
```

```
    scale_x_discrete(name = "Microbiota") +
    scale_y_continuous(name="% Metagenome Coverage", limits = c(0,100)) +
    # ylim(0, 100) +
    theme(panel.grid.major = element_blank(),
          panel.grid.minor = element_blank(),
          axis.line.x = element_line(size = 0, colour = "black"),
          axis.line.y = element_line(size = 0, colour = "black"),
          axis.line = element_line(size=1, colour = "black"),
          panel.border = element_rect(fill = NA, colour = "black", size = 1),
          panel.background = element_blank(),
          text=element_text(size = 8),
          axis.text.x=element_text(colour="black", size = 8),
          axis.text.y=element_text(colour="black", size = 8),
          axis.title.x=element_blank(),
          legend.title=element_blank(),
          legend.key = element_rect(color=NA, fill=NA)
    )
ageasthmacoverage
```

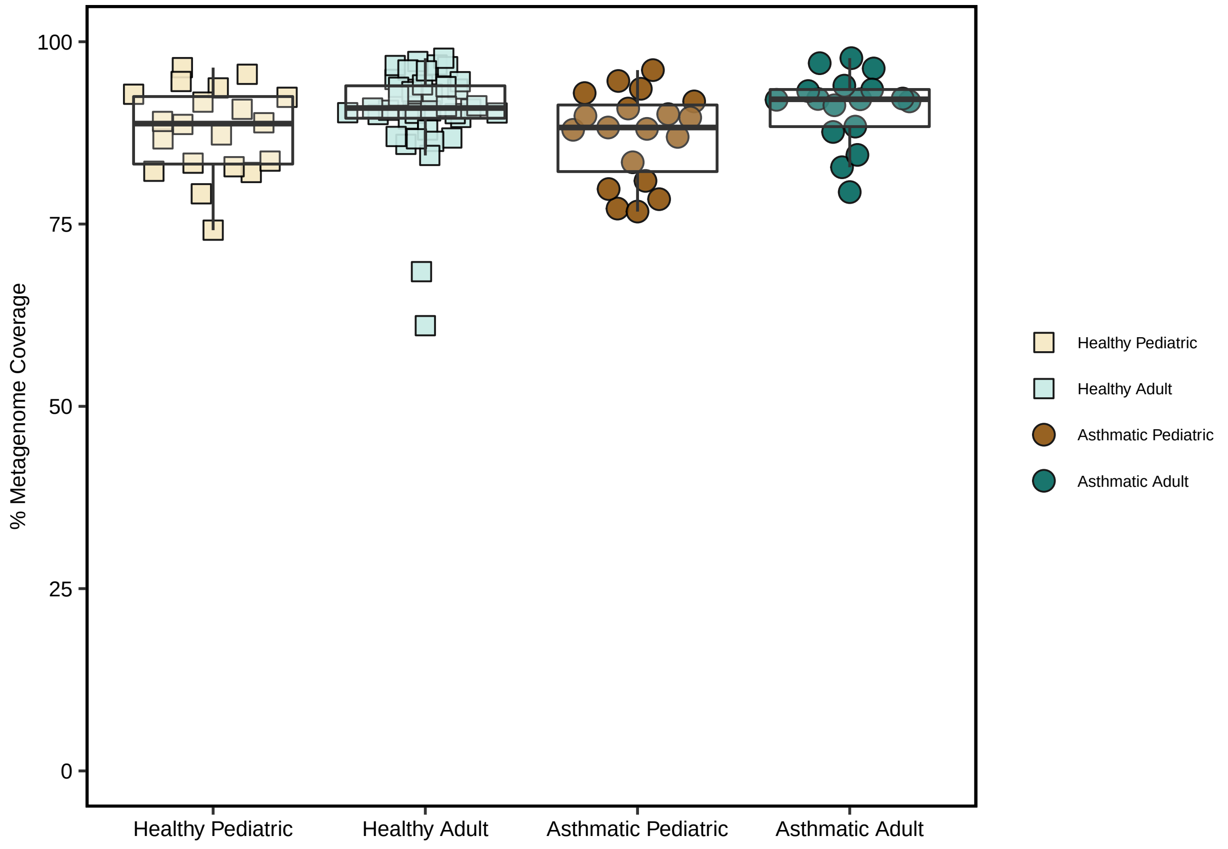


```
# ggsave(plot = ageasthmacoverage, filename =
↪   "nonpareilMetagenomeCoverage_boxplot_AgeAsthmaSplit.pdf",device = "pdf", width = 5.5,
↪   height = 4, units = "in", useDingbats=FALSE)

library(car)
Anova(lm(data = datframe, coverage~ASTHMA*AgeGroup, type=2))
```

```
## Warning: In lm.fit(x, y, offset = offset, singular.ok = singular.ok, ...) :
```

```
##  extra argument 'type' will be disregarded

## Anova Table (Type II tests)
##
## Response: coverage
##                  Sum Sq Df F value  Pr(>F)
## ASTHMA              0.2  1  0.0053 0.94196
## AgeGroup          199.4  1  5.0572 0.02694 *
## ASTHMA:AgeGroup     8.2  1  0.2084 0.64907
## Residuals        3587.4 91
## ---
## Signif. codes:  0 '***' 0.001 '**' 0.01 '*' 0.05 '.' 0.1 ' ' 1
```

```
# Anova(lm(data = datframe, coverage~ASTHMA*AgeGroup+readDepth), type = 2)
# Anova(lm(data = datframe, readDepth~ASTHMA*AgeGroup), type = 2)
```

### Figure S1C:

METACYC CPM pathway data filtering filter to present in at least 10% of samples - >=10

```
rm(list=ls())
source("./humann3_210613_functions_ngw.R")

mypwys <- read.csv("~/Library/CloudStorage/Box-Box/Kau
↪  Lab/Results/MARS/FecalMetagenomics/humann3_210613/MARSpathabundanceCPM_SUMMEDONLY.tsv",
↪  sep = "\t", skip = 0, stringsAsFactors = F, comment.char = "", check.names = F, quote
↪  = "")
dim(mypwys) # 384
```

```
## [1] 384 105
```

```
row.names(mypwys) <- mypwys[,1]
mypwys[,1] <- NULL
length(names(mypwys)) == 104
```

```
## [1] TRUE
```

```
names(mypwys) <- gsub(names(mypwys), pattern = "_interleaved_Abundance", replacement =
↪  "")

humann3_mf_filtered <- readRDS("./humann3_mf_filtered.rds")
row.names(humann3_mf_filtered) <- humann3_mf_filtered$SampleName

# filter
mypwysF <- filter_pathway_data(indata = mypwys, mymetadata = humann3_mf_filtered,
↪  mindetect=10)
length(names(mypwysF)) == 95
```

```
## [1] TRUE
```

```
dim(mypwysF) #78 (328 metacyc)
```

```
## [1] 312  95
```

```
dim(mypwys) #108
```

```
## [1] 384 104
```

```r
saveRDS(mypwys, file = "MARS_metacyc_pwys_CPM.rds")
saveRDS(mypwysF, file = "MARS_metacyc_pwys_CPM_filtered.rds")
```

### Figure S1C:

Plot top pathways overall

```r
library(stringr)
mypwys <- readRDS(file = "MARS_metacyc_pwys_CPM_filtered.rds")
humann3_mf_filtered <- readRDS("humann3_mf_filtered.rds")
row.names(humann3_mf_filtered) <- humann3_mf_filtered$filenamePeriod
humann3_mf = humann3_mf_filtered
means.df <- as.data.frame(apply(mypwys, MARGIN = 1, FUN = mean)) # get means
pwys_to_plot.df <- mypwys[order(means.df, decreasing = TRUE), ] # grab top pwys
pwys_to_plot.df <- pwys_to_plot.df[1:25,]
my_title = "Top Pathways"
myFactor <- as.character(row.names(pwys_to_plot.df))
myFactor <- comprehenr::to_vec( for(i in myFactor) ifelse(grepl(i, pattern=":"),
↪  stringr::str_split(i, pattern = ": ")[[1]][2], i))
myLevels <- rev(myFactor)
pwys_to_plot.df[,"PWY"] <- myFactor
pwys_to_plot.df$PWY <- factor(as.character(pwys_to_plot.df$PWY), levels =
↪  as.character(myLevels))
pwys_to_plot.df.melted <- reshape2::melt(pwys_to_plot.df, id.vars = "PWY")
# calculate means and error
pwys_to_plot.df.melted.forplotErrorBars <- my_data_summary(pwys_to_plot.df.melted,
↪  varname = "value", groupnames = c("PWY"))
```

```
## Loading required package: plyr

##
## Attaching package: 'plyr'

## The following object is masked from 'package:ggpubr':
##
##     mutate
```

```r
pwys_to_plot.df.melted.forplotErrorBars$se <-
  as.numeric(pwys_to_plot.df.melted.forplotErrorBars$sd/sqrt(dim(humann3_mf)[1]))
  pwys_to_plot.df.melted.forplotErrorBars$PWY <-
↪  factor(pwys_to_plot.df.melted.forplotErrorBars$PWY, levels =
↪  unique(pwys_to_plot.df.melted.forplotErrorBars$PWY))
  TopBarPlot <- ggplot(pwys_to_plot.df.melted.forplotErrorBars,
                                     aes(y=value, x=PWY), width=0.75) +
      geom_bar(stat="identity", color="black", fill="#C7EAE5",
               position=position_dodge()) +
      geom_errorbar(aes(ymin=value-se, ymax=value+se), width=.5,
                    position=position_dodge(.9))
  TopBarPlotFinal <- TopBarPlot +
    theme_classic() +
    theme(legend.text = element_text(size = 18, color = "black"),
          axis.text=element_text(size=16, color = "black"),
          axis.title = element_text(size = 18, color = "black"))+
```

```
    labs(title=my_title, #"Wilcoxon<0.05\n(before FDR correction)"
         x="MetaCyc Pathway ID", y="Copies per Million (CPM)")+
    scale_y_continuous(breaks = c(0, 100, 200, 300, 400, 500, 600)) +
    coord_flip()+
    # scale_fill_manual(values = c(brewer.pal(7, "BrBG")[c(7,5)]))+
    scale_color_manual(values = c(brewer.pal(7, "BrBG")[7]), guide="none")
TopBarPlotFinal
```

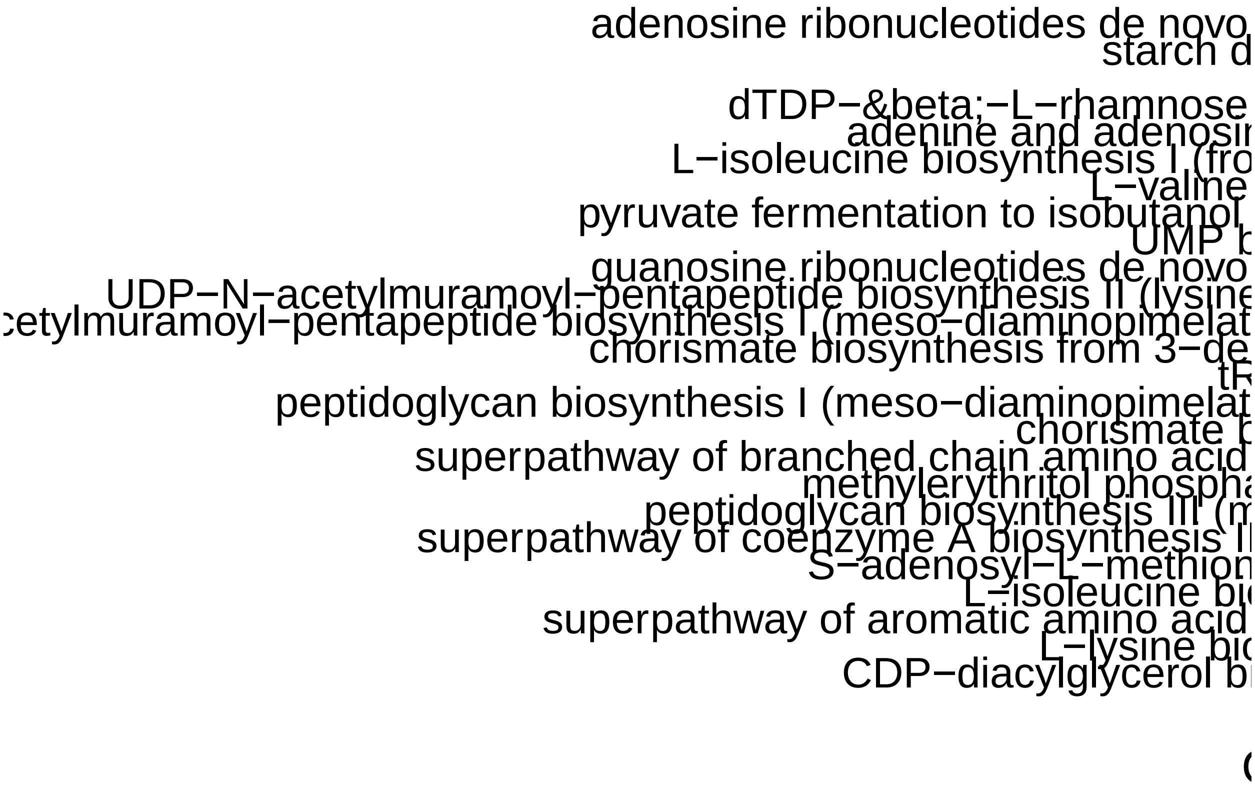


#### Figure 1D:

Get mean by asthma and age values for plotting

```
rm(list = ls())
source("./humann3_210613_functions_ngw.R")

library(tidyverse)
```

```
## -- Attaching packages --------------------------------------- tidyverse 1.3.1 --

## v tibble  3.1.7     v purrr   0.3.4
## v tidyr   1.2.0     v dplyr   1.0.9
## v readr   2.1.2     v forcats 0.5.1

## -- Conflicts ------------------------------------------ tidyverse_conflicts() --
## x dplyr::arrange()   masks plyr::arrange()
## x purrr::compact()   masks plyr::compact()
## x dplyr::count()     masks plyr::count()
## x dplyr::failwith()  masks plyr::failwith()
## x dplyr::filter()    masks stats::filter()
```

```
## x dplyr::id()        masks plyr::id()
## x dplyr::lag()       masks stats::lag()
## x dplyr::mutate()    masks plyr::mutate(), ggpubr::mutate()
## x dplyr::recode()    masks car::recode()
## x dplyr::rename()    masks plyr::rename()
## x purrr::some()      masks car::some()
## x dplyr::summarise() masks plyr::summarise()
## x dplyr::summarize() masks plyr::summarize()
```

```
# library(dplyr)
library(reshape2)
```

```
##
## Attaching package: 'reshape2'

## The following object is masked from 'package:tidyr':
##
##     smiths
```

```
mypwys <- readRDS(file = "MARS_metacyc_pwys_CPM_filtered.rds")
humann3_mf_filtered <- readRDS("./humann3_mf_filtered.rds")
row.names(humann3_mf_filtered) <- humann3_mf_filtered$SampleName
humann3_mf = humann3_mf_filtered

saveRDS(mean_by_group_pwys(my_data= mypwys, group_by_me = "ASTHMA", sample_mf =
↪  humann3_mf, basedir="./"),
        paste0("METACYC_PWYS", ".my.df.means.", "ASTHMA", ".rds"))
saveRDS(mean_by_group_pwys(my_data= mypwys, group_by_me = "AgeGroup", sample_mf =
↪  humann3_mf, basedir="./"),
        paste0("METACYC_PWYS", ".my.df.means.", "AgeGroup", ".rds"))

# asthma only age
mypwysAsthma <- mypwys[, as.character(humann3_mf[humann3_mf$ASTHMA=="Asthmatic",
↪  "SampleName"])]
asthmaAge <- mean_by_group_pwys(my_data= mypwysAsthma, group_by_me = "AgeGroup",
↪  sample_mf = humann3_mf, basedir="./")
row.names(asthmaAge) <- c("Asthmatic Adult", "Asthmatic Pediatric")
# healthy only age
mypwysHealthy <- mypwys[, as.character(humann3_mf[humann3_mf$ASTHMA=="Healthy",
↪  "SampleName"])]
healthyAge <- mean_by_group_pwys(my_data= mypwysHealthy, group_by_me = "AgeGroup",
↪  sample_mf = humann3_mf, basedir="./")
row.names(healthyAge) <- c("Healthy Adult", "Healthy Pediatric")
ASTHMAAGEMEANS = rbind(healthyAge, asthmaAge)
saveRDS(ASTHMAAGEMEANS, paste0("METACYC_PWYS", ".my.df.means.", "ASTHMAAge", ".rds"))

# we also just want mean values overall for ranking and overview barplot purposes:
mypwys.means <- as.data.frame(apply(mypwys, MARGIN = 1, FUN = mean))
# mypwys.means <- mypwys.means[order(mypwys.means, decreasing = TRUE),]
saveRDS(mypwys.means, paste0("METACYC_PWYS", ".my.df.means.", "Individual", ".rds"))
```

Wilcox tests and boxplot

```
rm(list = ls())
source("./humann3_210613_functions_ngw.R")
```

```r
library(ggpubr)
library(ggplot2)
library(ggsignif)
library(ggbeeswarm)
library(RColorBrewer)

mypwys <- readRDS(file = "MARS_metacyc_pwys_CPM_filtered.rds")
humann3_mf_filtered <- readRDS("./humann3_mf_filtered.rds")
row.names(humann3_mf_filtered) <- humann3_mf_filtered$SampleName
humann3_mf = humann3_mf_filtered
plotmypwys <- mypwys[order(rowSums(mypwys), decreasing=T),] #reorder by abundance
plotmypwys[,"PWY"] <- row.names(plotmypwys)
plotmypwys$PWY <- factor(plotmypwys$PWY)
plotmypwys.melted <- reshape2::melt(plotmypwys, id.vars = "PWY")
plotmypwys.melted$Asthma <- humann3_mf[as.character(plotmypwys.melted$variable),"ASTHMA"]

PADJUSTTYPE = "fdr"

# wilcoxon on all pathway normalized counts
allcounts.wilcox <- plotmypwys.melted %>%
  group_by(PWY) %>%
  dplyr::summarise(pval = suppressWarnings(wilcox.test(value~Asthma))$p.value,
            type=suppressWarnings(wilcox.test(value~Asthma))$alternative,
            mean=mean(value),
            incidence=sum(value>0))
allcounts.wilcox$padj <- p.adjust(allcounts.wilcox$pval, method = PADJUSTTYPE, n =
↪  length(allcounts.wilcox$pval))
allcounts.wilcox <- allcounts.wilcox[order(allcounts.wilcox$pval),]
saveRDS(allcounts.wilcox, "metacycpwy_Asthma_wilcox_test_fdr.rds")

# PLOT ####
mean_by_group_pwys_asthma <- readRDS(file = paste0("METACYC_PWYS", ".my.df.means.",
↪  "ASTHMA", ".rds"))
allcounts.wilcox <- data.frame(allcounts.wilcox)
allcounts.wilcox <- allcounts.wilcox[order(allcounts.wilcox$mean, decreasing = F),]
pwysSigWilcox <- mypwys[as.character(allcounts.wilcox[allcounts.wilcox$padj < 0.2,
↪  "PWY"]),]
# pwysSigWilcox
p2 <- pwy_barplot(means.df=mean_by_group_pwys_asthma, pwys_to_plot.df=pwysSigWilcox,
↪  my_title="Wilcoxon FDR-corrected p<0.2", my_format = "boxplot")
```

```
## Loading required package: comprehenr
```

```r
# ggsave(plot = p2,
#        filename = "metacycpwy_wilcox_padj<0.2_boxplot.jpg", device = "jpg",
#        width = 14, height = 6, units = "in")
# ggsave(plot = p2,
#        filename = "metacycpwy_wilcox_padj<0.2_boxplot.pdf", device = "pdf",
#        width = 14, height = 6, units = "in", useDingbats=FALSE)
p2
```

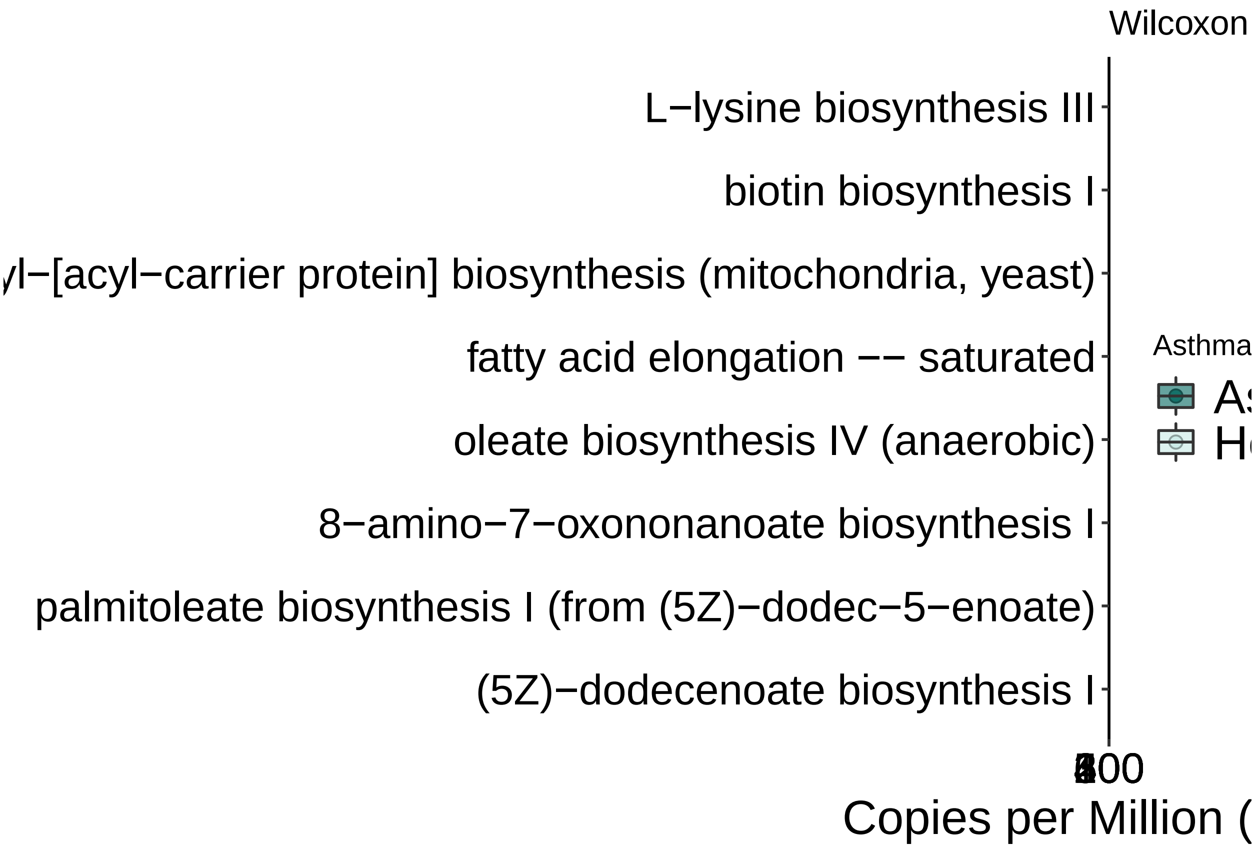


#### Figure 1F:

Wilcoxon heatmap Age, Obesity, ACT wilcoxon tests

```
rm(list = ls())
source("./humann3_210613_functions_ngw.R")
setwd("./")

library(ggpubr)
library(ggplot2)
library(ggsignif)
library(ggbeeswarm)
library(RColorBrewer)

mypwys <- readRDS(file = "MARS_metacyc_pwys_CPM_filtered.rds")
humann3_mf_filtered <- readRDS("./humann3_mf_filtered.rds")
row.names(humann3_mf_filtered) <- humann3_mf_filtered$SampleName
humann3_mf = humann3_mf_filtered
plotmypwys <- mypwys[order(rowSums(mypwys), decreasing=T),] #reorder by abundance
plotmypwys[,"PWY"] <- row.names(plotmypwys)
plotmypwys$PWY <- factor(plotmypwys$PWY)
plotmypwys.melted <- reshape2::melt(plotmypwys, id.vars = "PWY")
plotmypwys.melted$Asthma <- humann3_mf[as.character(plotmypwys.melted$variable),"ASTHMA"]
plotmypwys.melted$AgeGroup <-
↪  factor(humann3_mf[as.character(plotmypwys.melted$variable),"AgeGroup"])
plotmypwys.melted$weightClass <-
↪  factor(humann3_mf[as.character(plotmypwys.melted$variable),"weightClassBinary"])
```

```r
plotmypwys.melted$ACTScoreClass <-
↪  factor(humann3_mf[as.character(plotmypwys.melted$variable),"ACTScoreClass"])
# plotmypwys.melted$ACTScoreClass <-
↪  factor(ifelse(as.numeric(as.character(plotmypwys.melted$ACTScore)) > 19,
↪  "Controlled", "Poorly Controlled"))
PADJUSTTYPE = "fdr"

#AGE
allcounts.wilcox.age <- plotmypwys.melted %>%
  group_by(PWY) %>%
  dplyr::summarise(pval = suppressWarnings(wilcox.test(value~AgeGroup))$p.value,
            type=suppressWarnings(wilcox.test(value~AgeGroup))$alternative,
            mean=mean(value),
            incidence=sum(value>0))
allcounts.wilcox.age$padj <- p.adjust(allcounts.wilcox.age$pval,
                                      method = PADJUSTTYPE,
                                      n = length(allcounts.wilcox.age$pval))
allcounts.wilcox.age <- allcounts.wilcox.age[order(allcounts.wilcox.age$pval),]
saveRDS(allcounts.wilcox.age, "metacycpwy_AgeGroup_wilcox_test_fdr.rds")
# allcounts.wilcox <- readRDS(file = "metacycpwy_AgeGroup_wilcox_test_fdr.rds")

#Obesity
allcounts.wilcox.obesity <- plotmypwys.melted %>%
  group_by(PWY) %>%
  dplyr::summarise(pval = suppressWarnings(wilcox.test(value~weightClass))$p.value,
            type=suppressWarnings(wilcox.test(value~weightClass))$alternative,
            mean=mean(value),
            incidence=sum(value>0))
allcounts.wilcox.obesity$padj <- p.adjust(allcounts.wilcox.obesity$pval,
                                      method = PADJUSTTYPE,
                                      n = length(allcounts.wilcox.obesity$pval))
allcounts.wilcox.obesity <-
↪  allcounts.wilcox.obesity[order(allcounts.wilcox.obesity$pval),]
saveRDS(allcounts.wilcox.obesity, "metacycpwy_weightClass_wilcox_test_fdr.rds")
# allcounts.wilcox <- readRDS(file = "metacycpwy_weightClass_wilcox_test_fdr.rds")

#ACT score
allcounts.wilcox.act <- plotmypwys.melted %>%
  group_by(PWY) %>%
  dplyr::summarise(pval = suppressWarnings(wilcox.test(value~ACTScoreClass))$p.value,
            type=suppressWarnings(wilcox.test(value~ACTScoreClass))$alternative,
            mean=mean(value, na.rm=T),
            incidence=sum(value>0))
allcounts.wilcox.act$padj <- p.adjust(allcounts.wilcox.act$pval,
                                      method = PADJUSTTYPE,
                                      n = length(allcounts.wilcox.act$pval))
allcounts.wilcox.act <- allcounts.wilcox.act[order(allcounts.wilcox.act$pval),]
dim(allcounts.wilcox.act[allcounts.wilcox.act$padj<0.2,])
```

```
## [1] 2 6
```

```r
dim(allcounts.wilcox.act[allcounts.wilcox.act$pval<0.05,])
```

```
## [1] 24  6
```

```
saveRDS(allcounts.wilcox.act, "metacycpwy_ACTClass_wilcox_test_fdr.rds")
# allcounts.wilcox <- readRDS(file = "metacycpwy_AgeGroup_wilcox_test_fdr.rds")


# Obesity for only asthmatics
plotmypwys.melted.asthmaonly <- plotmypwys.melted
plotmypwys.melted.asthmaonly$Asthma <-
↪  humann3_mf[as.character(plotmypwys.melted.asthmaonly$variable),"ASTHMA"]
plotmypwys.melted.asthmaonly <-
↪  plotmypwys.melted.asthmaonly[plotmypwys.melted.asthmaonly$Asthma == "Asthmatic",]
allcounts.wilcox.AsOb <- plotmypwys.melted.asthmaonly %>%
  group_by(PWY) %>%
  dplyr::summarise(pval = suppressWarnings(wilcox.test(value~weightClass))$p.value,
            type=suppressWarnings(wilcox.test(value~weightClass))$alternative,
            mean=mean(value, na.rm=T),
            incidence=sum(value>0))
allcounts.wilcox.AsOb$padj <- p.adjust(allcounts.wilcox.AsOb$pval,
                                      method = PADJUSTTYPE,
                                      n = length(allcounts.wilcox.AsOb$pval))
allcounts.wilcox.AsOb <- allcounts.wilcox.AsOb[order(allcounts.wilcox.AsOb$pval),]
dim(allcounts.wilcox.AsOb[allcounts.wilcox.AsOb$padj<0.2,])
```

```
## [1] 0 6
```

```
dim(allcounts.wilcox.AsOb[allcounts.wilcox.AsOb$pval<0.05,])
```

```
## [1] 9 6
```

```
saveRDS(allcounts.wilcox.AsOb,
↪  "metacycpwy_weightClass_asthmaticsOnly_wilcox_test_fdr.rds")
```

Make Heatmap

```
rm(list = ls())
source("./humann3_210613_functions_ngw.R")

library(ggpubr)
library(ggplot2)
library(ggsignif)
library(RColorBrewer)
library(ggbeeswarm)
library(tidyverse)
library(gplots)
```

```
##
## Attaching package: 'gplots'

## The following object is masked from 'package:stats':
##
##     lowess
```

```
mypwys <- readRDS(file = "MARS_metacyc_pwys_CPM_filtered.rds")
humann3_mf_filtered <- readRDS("./humann3_mf_filtered.rds")
row.names(humann3_mf_filtered) <- humann3_mf_filtered$SampleName
humann3_mf = humann3_mf_filtered
stat.wilcoxon <- data.frame(readRDS(file = "metacycpwy_Asthma_wilcox_test_fdr.rds"))
```

```r
sig.pathways <- as.character(stat.wilcoxon[stat.wilcoxon$padj<.2,"PWY"])
mean_by_group_pwys_asthma <- readRDS(file = paste0("METACYC_PWYS", ".my.df.means.",
↪  "ASTHMA", ".rds"))
foldchange_asthma <-
↪  mean_by_group_pwys_asthma["Asthmatic",]/mean_by_group_pwys_asthma["Healthy",]
mean_by_group_pwys_age <- readRDS(file = paste0("METACYC_PWYS", ".my.df.means.",
↪  "AgeGroup", ".rds"))
foldchange_age <- mean_by_group_pwys_age["Adult",]/mean_by_group_pwys_age["Pediatric",]

# all (healthy and asthma) obese v non obese
humann3_mf$Obesity <- humann3_mf$weightClassBinary
saveRDS(mean_by_group_pwys(my_data= mypwys, group_by_me = "Obesity", sample_mf =
↪  humann3_mf, basedir="./"), paste0("METACYC_PWYS", ".my.df.means.", "weightClass",
↪  ".rds"))
mean_by_group_pwys_obesity <- readRDS(file = paste0("METACYC_PWYS", ".my.df.means.",
↪  "weightClass", ".rds"))
foldchange_obesity_all <-
↪  mean_by_group_pwys_obesity["Obese",]/mean_by_group_pwys_obesity["Non-Obese",]

# only asthma obese vs non-obese
saveRDS(mean_by_group_pwys(my_data=mypwys[,row.names(humann3_mf_filtered[humann3_mf_filtered$ASTHMA=="As
↪  group_by_me = "Obesity", sample_mf = humann3_mf, basedir="./"),
↪  paste0("METACYC_PWYS", ".my.df.means.", "weightClassASTHMATICONLY", ".rds")) # 312 36
mean_by_group_pwys_obesity_asthmaticsonly <- readRDS(file = paste0("METACYC_PWYS",
↪  ".my.df.means.", "weightClassASTHMATICONLY", ".rds"))
foldchange_obesity_asthmatics <-
↪  mean_by_group_pwys_obesity_asthmaticsonly["Obese",]/mean_by_group_pwys_obesity_asthmaticsonly["Non-

# ACT groups
humann3_mf$ACTScoreClass <- factor(humann3_mf$ACTScoreClass)
modified_mypwys  <- mypwys[,names(mypwys) %notin%
↪  gsub(row.names(humann3_mf[is.na(humann3_mf$ACTScoreClass),]), pattern = "[.]",
↪  replacement = "-")]
dim(modified_mypwys)[2] == 36
```

```
## [1] FALSE
```

```r
saveRDS(mean_by_group_pwys(my_data= modified_mypwys, group_by_me = "ACTScoreClass",
↪  sample_mf = humann3_mf, basedir="./"),
        paste0("METACYC_PWYS", ".my.df.36samples.means.", "ACTScoreClass", ".rds"))
mean_by_group_pwys_act <- readRDS(file = paste0("METACYC_PWYS",
↪  ".my.df.36samples.means.", "ACTScoreClass", ".rds"))
foldchange_act <- mean_by_group_pwys_act["Controlled",]/mean_by_group_pwys_act["Poorly
↪  Controlled",]
# foldchange_act

foldchange.df <- rbind(foldchange_asthma,
                       foldchange_age,
                       foldchange_obesity_all,
                       foldchange_obesity_asthmatics,
                       foldchange_act)
row.names(foldchange.df) <- c("Asthma", "Age Group", "Obesity", "Asthmatic Obesity", "ACT
↪  Class")
# wilcoxon results for stats
```

```r
stat.wilcoxon.age <- data.frame(readRDS(file =
↪  "metacycpwy_AgeGroup_wilcox_test_fdr.rds"))
sig.pathways.age <- as.character(stat.wilcoxon.age[stat.wilcoxon.age$padj<.2,"PWY"])
row.names(stat.wilcoxon.age) <- stat.wilcoxon.age$PWY
stat.wilcoxon.age  <- stat.wilcoxon.age[as.character(stat.wilcoxon$PWY),]

stat.wilcoxon.weightclass <- data.frame(readRDS(file =
↪  "metacycpwy_weightClass_wilcox_test_fdr.rds"))
sig.pathways.weightclass <-
↪  as.character(stat.wilcoxon.weightclass[stat.wilcoxon.weightclass$padj<.2,"PWY"])
row.names(stat.wilcoxon.weightclass) <- stat.wilcoxon.weightclass$PWY
stat.wilcoxon.weightclass  <- stat.wilcoxon.weightclass[as.character(stat.wilcoxon$PWY),]

stat.wilcoxon.weightclassASTHMATICSONLY <- data.frame(readRDS(file =
↪  "metacycpwy_weightClass_asthmaticsOnly_wilcox_test_fdr.rds"))
sig.pathways.weightclassASTHMATICSONLY <-
↪  as.character(stat.wilcoxon.weightclassASTHMATICSONLY[stat.wilcoxon.weightclassASTHMATICSONLY$padj<.
row.names(stat.wilcoxon.weightclassASTHMATICSONLY) <-
↪  stat.wilcoxon.weightclassASTHMATICSONLY$PWY
stat.wilcoxon.weightclassASTHMATICSONLY  <-
↪  stat.wilcoxon.weightclassASTHMATICSONLY[as.character(stat.wilcoxon$PWY),]

stat.wilcoxon.act <- data.frame(readRDS(file =
↪  "metacycpwy_ACTClass_wilcox_test_fdr.rds"))
sig.pathways.act <- as.character(stat.wilcoxon.act[stat.wilcoxon.act$padj<.2,"PWY"])
row.names(stat.wilcoxon.act) <- stat.wilcoxon.act$PWY
stat.wilcoxon.act  <- stat.wilcoxon.act[as.character(stat.wilcoxon$PWY),]

stat.wilcoxon.age[sig.pathways, "padj"] < 0.2 # are any of the asthma pwys also sig by
↪  age groups
```

```
## [1] FALSE FALSE FALSE FALSE FALSE FALSE FALSE FALSE
```

```r
top_sig_pathways <- c(sig.pathways, sig.pathways.age[1:5], sig.pathways.weightclass,
↪  sig.pathways.act)
all_sig_pathways <- c(sig.pathways, sig.pathways.age, sig.pathways.weightclass,
↪  sig.pathways.weightclassASTHMATICSONLY, sig.pathways.act)
pval.df <- data.frame(as.character(stat.wilcoxon$PWY), stat.wilcoxon$padj,
↪  stat.wilcoxon.age$padj, stat.wilcoxon.weightclass$padj,
↪  stat.wilcoxon.weightclassASTHMATICSONLY$padj, stat.wilcoxon.act$padj)
names(pval.df) <- c("PWY", "Asthma", "AgeGroup","weightClass", "weightClassASTHMATICS",
↪  "ACT")
row.names(pval.df) <- pval.df$PWY
# View(pval.df[top_sig_pathways,])
tmp <- pval.df[top_sig_pathways,]
tmp$PWY <-  NULL
tmp <- ifelse(tmp<0.2, "<0.2", ifelse(tmp<0.5, "<0.5", "n.s.")) # ifelse(tmp<0.8, "<0.8",
↪  "n.s.")
# tmp

plotme <- foldchange.df[,top_sig_pathways[c(6,8,1,5,2,7,4,3)]] # plot the ones different
↪  for asthma
## Make vector of colors for values below threshold
# num_below_zero = sum(plotme<=0.94)
```

```r
num_below_zero = 12
# num_below_zero = min(plotme)*52
rc1 = colorRampPalette(colors = c("navy", "white"), space="Lab")(num_below_zero)
## Make vector of colors for values above threshold
rc2 = colorRampPalette(colors = c("white", "darkred"), space="Lab")(52-num_below_zero)
rampcols = c(rc1, rc2)
## In your example, this line sets the color for values between 49 and 51.
rampcols[c(num_below_zero, num_below_zero+1)] = rgb(t(col2rgb("white")),
↪  maxColorValue=256)

pdf(file = "./DApwys_Wilcoxon_foldchange_heatmap.pdf")
heatmap.2(as.matrix(as.matrix(plotme)),
                  Colv = NA, Rowv = NA,  #Colv = NA, Rowv = NA gets rid of clustering
                  ↪  order
                  dendrogram='none' ,
                  scale = "none",
                  xlab = "Model",
                  margins=c(15,5),
                  cexRow = 1, #label size
                  cexCol = 1, #label size
                  # labCol="",
                  trace = "none",
                  density.info = "none",
                  # srt=30, # I think this only works with heatmap.2()
                  # add.expr = text(x = seq_along(colnames(compareTabForHeatmapRanks)),
                  ↪  y=-0.2, srt=45,
                      # labels=colnames(compareTabForHeatmapRanks), xpd=TRUE, cex = 2),
                  # RowSideColors=colSide,  # this is the l2fc part
                  col=rampcols)   # colors for ranks
dev.off()
```

```
## pdf
##   2
```

### Figure 1E, S2B, S2C:

Stacked barplots - averaged

```r
rm(list = ls())
source("./humann3_210613_functions_ngw.R")

library(tidyverse)
library(ggpubr)
library(ggplot2)
library(ggsignif)
library(RColorBrewer)
library(ggbeeswarm)

stat.wilcoxon <- data.frame(readRDS(file = "metacycpwy_Asthma_wilcox_test_fdr.rds"))
sig.pwys <- as.character(stat.wilcoxon[stat.wilcoxon$padj < 0.2, "PWY"])
datatable<-read.table("/Users/naomiwilson/Library/CloudStorage/Box-Box/Kau
↪  Lab/Results/MARS/FecalMetagenomics/humann3_210613/MARSpathabundanceCPM_forPlotting.txt",
↪  sep="\t", comment.char="", quote="", header = T) # output from humann with taxonomic
↪  marker gene information
```

```r
row.names(datatable) <- as.character(datatable$SAMPLE)
datatable$SAMPLE <- NULL
names(datatable) <- gsub(names(datatable), pattern = "\\.", replacement = "-")
names(datatable) <- gsub(names(datatable), pattern = "_interleaved_Abundance",
↪  replacement = "")
asthmarow <- datatable[1,]

row.names(datatable)[grepl(pattern = str_split(sig.pwys[1], pattern = ":")[[1]][1],
↪  row.names(datatable))] # quotes with backlashes are causing issues, so remove them:
```

```
##  [1] "\"PWY-7388: octanoyl-[acyl-carrier protein] biosynthesis (mitochondria, yeast)\""
##  [2] "\"PWY-7388: octanoyl-[acyl-carrier protein] biosynthesis (mitochondria, yeast)|g__Bacteroides.s
##  [3] "\"PWY-7388: octanoyl-[acyl-carrier protein] biosynthesis (mitochondria, yeast)|g__Bacteroides.s
##  [4] "\"PWY-7388: octanoyl-[acyl-carrier protein] biosynthesis (mitochondria, yeast)|g__Bacteroides.s
##  [5] "\"PWY-7388: octanoyl-[acyl-carrier protein] biosynthesis (mitochondria, yeast)|g__Enterobacter
##  [6] "\"PWY-7388: octanoyl-[acyl-carrier protein] biosynthesis (mitochondria, yeast)|g__Escherichia.s
##  [7] "\"PWY-7388: octanoyl-[acyl-carrier protein] biosynthesis (mitochondria, yeast)|g__Haemophilus.s
##  [8] "\"PWY-7388: octanoyl-[acyl-carrier protein] biosynthesis (mitochondria, yeast)|g__Klebsiella.s
##  [9] "\"PWY-7388: octanoyl-[acyl-carrier protein] biosynthesis (mitochondria, yeast)|g__Streptococcu
## [10] "\"PWY-7388: octanoyl-[acyl-carrier protein] biosynthesis (mitochondria, yeast)|unclassified\""
```

```r
row.names(datatable) <- gsub(row.names(datatable), pattern = "\"", replacement = "")
row.names(datatable)[grepl(pattern = str_split(sig.pwys[1], pattern = ":")[[1]][1],
↪  row.names(datatable))]
```

```
##  [1] "PWY-7388: octanoyl-[acyl-carrier protein] biosynthesis (mitochondria, yeast)"
##  [2] "PWY-7388: octanoyl-[acyl-carrier protein] biosynthesis (mitochondria, yeast)|g__Bacteroides.s_
##  [3] "PWY-7388: octanoyl-[acyl-carrier protein] biosynthesis (mitochondria, yeast)|g__Bacteroides.s_
##  [4] "PWY-7388: octanoyl-[acyl-carrier protein] biosynthesis (mitochondria, yeast)|g__Bacteroides.s_
##  [5] "PWY-7388: octanoyl-[acyl-carrier protein] biosynthesis (mitochondria, yeast)|g__Enterobacter.s
##  [6] "PWY-7388: octanoyl-[acyl-carrier protein] biosynthesis (mitochondria, yeast)|g__Escherichia.s_
##  [7] "PWY-7388: octanoyl-[acyl-carrier protein] biosynthesis (mitochondria, yeast)|g__Haemophilus.s_
##  [8] "PWY-7388: octanoyl-[acyl-carrier protein] biosynthesis (mitochondria, yeast)|g__Klebsiella.s__
##  [9] "PWY-7388: octanoyl-[acyl-carrier protein] biosynthesis (mitochondria, yeast)|g__Streptococcus.
## [10] "PWY-7388: octanoyl-[acyl-carrier protein] biosynthesis (mitochondria, yeast)|unclassified"
```

```r
all(!grepl(pattern = "\"", row.names(datatable), fixed = TRUE)) # is true if quotes
↪  removed
```

```
## [1] TRUE
```

```r
sig.pwys.toMatch <- comprehenr::to_vec(for (i in sig.pwys) str_split(i, pattern =
↪  ":")[[1]][1])
row_pathway <- row.names(datatable)[grepl(pattern = paste(sig.pwys.toMatch,collapse="|"),
↪  x = row.names(datatable))]
# row_pathway <- row.names(datatable)[grep("PWY-7664|PWY-6282", row.names(datatable))]
filteredtable<-datatable[row_pathway,]
filteredtable<-sapply(filteredtable, function(x) as.numeric(as.character(x)))
row.names(filteredtable)<-row_pathway

# Add an "Community" column for difference between community-level abundance and sum of
↪  stratified taxa abundance
pwy_root_names <- comprehenr::to_vec(for (i in row.names(filteredtable)) str_split(i,
↪  pattern = "\\|")[[1]][1])
# grep(row.names(filteredtable), pattern = paste0(pwy_root_names[1], "|"))
```

```r
for (i in unique(pwy_root_names)) {
  tmpdat <- filteredtable[grep(row.names(filteredtable), pattern = i, fixed=T),]
  filteredtable <- rbind(filteredtable, tmpdat[grep(row.names(tmpdat), pattern = "\\|",
↪  invert = T),] - colSums(tmpdat[grep(row.names(tmpdat), pattern = "\\|"),]))
  row.names(filteredtable)[length(row.names(filteredtable))] <- paste0(i, "|Community")
  # print % unmapped:
  print(i)
  # print("Avg unmapped abundance")
  # print(mean(tmpdat[grep(row.names(tmpdat), pattern = "\\|", invert = T),] -
  ↪  colSums(tmpdat[grep(row.names(tmpdat), pattern = "\\|"),]))) # total - species
  ↪  ("community", i.e. full pathways unmapped to a species)
  # print("Avg mapped abundance")
  print(mean(colSums(tmpdat[grep(row.names(tmpdat), pattern = "\\|"),]))) # total only
  ↪  species
    # print("% Unmapped (Abundance)")
    print(100*mean(tmpdat[grep(row.names(tmpdat), pattern = "\\|", invert = T),] -
          colSums(tmpdat[grep(row.names(tmpdat), pattern =
          ↪  "\\|"),]))/mean(tmpdat[grep(row.names(tmpdat), pattern = "\\|", invert =
          ↪  T),])) # % unmapped
}
```

```
## [1] "BIOTIN-BIOSYNTHESIS-PWY: biotin biosynthesis I"
## [1] 9.425372
## [1] 84.68794
## [1] "FASYN-ELONG-PWY: fatty acid elongation -- saturated"
## [1] 15.70108
## [1] 73.86733
## [1] "PWY-2942: L-lysine biosynthesis III"
## [1] 289.9439
## [1] 24.79573
## [1] "PWY-6282: palmitoleate biosynthesis I (from (5Z)-dodec-5-enoate)"
## [1] 14.54368
## [1] 70.83382
## [1] "PWY-6519: 8-amino-7-oxononanoate biosynthesis I"
## [1] 9.612836
## [1] 82.19912
## [1] "PWY-7388: octanoyl-[acyl-carrier protein] biosynthesis (mitochondria, yeast)"
## [1] 10.81352
## [1] 82.10532
## [1] "PWY-7664: oleate biosynthesis IV (anaerobic)"
## [1] 15.05876
## [1] 72.66493
## [1] "PWY0-862: (5Z)-dodecenoate biosynthesis I"
## [1] 14.57128
## [1] 70.70463
```

```r
humann3_mf_filtered <- readRDS("./humann3_mf_filtered.rds")
row.names(humann3_mf_filtered) <- humann3_mf_filtered$SampleName
humann3_mf = humann3_mf_filtered
# remove disqualifieds
filteredtable <- filteredtable[, humann3_mf$SampleName]
# get means:
dat <- as.data.frame(t(mean_by_group_pwys(my_data= filteredtable, group_by_me = "ASTHMA",
↪  sample_mf = humann3_mf, basedir="./")))
```

```r
dat$PWY <- row.names(dat)
# remove sum rows (no stratification)
dat <- dat[grep(row.names(dat), pattern = "\\|"),]

#Assigning asthma Divide
dat.melted <- reshape2::melt(dat)
```

```
## Using PWY as id variables
```

```r
dat.melted$variable <- factor(dat.melted$variable, levels = c("Healthy", "Asthmatic"))
dat.melted$AsthmaPWY <- paste0(comprehenr::to_vec(for (i in dat.melted$PWY) str_split(i,
↪  pattern = ":")[[1]][1]), "_", as.numeric(dat.melted$variable))
dat.melted$AsthmaPWY <- factor(dat.melted$AsthmaPWY)
# dat.melted$PWYshort <- comprehenr::to_vec(for (i in dat.melted$PWY) str_split(i,
↪  pattern = ":")[[1]][1])
dat.melted$PWYshort <- comprehenr::to_vec(for (i in dat.melted$PWY) str_split(i, pattern
↪  = "[|]")[[1]][1])
dat.melted$PWYshort <- comprehenr::to_vec(for (i in dat.melted$PWYshort) str_split(i,
↪  pattern = "[:] ")[[1]][2])
dat.melted$PWYshort <- comprehenr::to_vec(for (i in dat.melted$PWYshort) str_split(i,
↪  pattern = " [(][a|m]")[[1]][1])
dat.melted$taxa <-gsub(".*s__","", perl=T, gsub(".*\\|","", perl=T, dat.melted$PWY))
sum(is.na(dat.melted))/2  #should match number of Pathways being plotted and no more.
```

```
## [1] 0
```

```r
dat.melted <- na.omit(dat.melted)

library(viridis)
```

```
## Loading required package: viridisLite
```

```r
library(RColorBrewer)
makeStackedBarPlot <- function(indat, legend_length=8) {  #CPMtable,
  require(viridis)
  require(ggplot2)

  swr = function(string, nwrap=10) {
    # create line breaks in facet grid labels
  paste(strwrap(string, width=nwrap), collapse="\n")}
  swr = Vectorize(swr)
  indat$PWYshort <- swr(indat$PWYshort)

  # reorder taxa
  # re-order by abundance and bin low abundance species into "Other" bin for better
  ↪  visual
  factorOrder <- c(rev(indat %>% group_by(taxa) %>% summarize(sum=sum(value)) %>%
↪  arrange(sum) %>% pull(taxa))[1:legend_length], "Other Bacteria")
  factorOrder <- factorOrder[factorOrder %notin% "unclassified"] # bin unclassified with
↪  Other
  indat$taxa <- ifelse(indat$taxa %in% factorOrder, indat$taxa, "Other Bacteria")
  indat$taxa <- factor(indat$taxa, levels = factorOrder)
  levels(indat$taxa) = gsub(levels(indat$taxa), pattern = "_", replacement = " ")
  prettylist = levels(indat$taxa)
```

```r
  # pathways in order of abundance of top species
  PWYorder <- indat %>% filter(taxa %in% prettylist, variable=="Asthmatic") %>%
↪  group_by(PWYshort) %>% summarize(sum=sum(value)) %>% arrange(sum) %>% pull(PWYshort)
  indat$PWYshort <- factor(indat$PWYshort, levels=rev(PWYorder))

  out = ggplot(indat, aes(fill=taxa, x = variable, y = value)) +
  geom_bar(stat="identity", position = "stack", width = 0.8) +
  theme_classic() +
  facet_grid( ~ PWYshort)+#, scales = "free_x", space = "free_x") +
  # scale_y_continuous(limits=c(0,1), expand = c(0,0))  +
  scale_y_continuous(expand = c(0, 0))+
  xlab(label = "Cohort") + ylab("Pathway Copies Per Million") +
  scale_fill_viridis(option = "turbo", discrete = T, begin = 1, end = 0) + # rainbow
  # scale_fill_manual(values = brewer.pal(7, "BrBG"))+
  theme(axis.title = element_text(size=12, color = "black"),
        axis.text.x = element_text(size=12, color = "black", angle=45, vjust=1, hjust=1),
        axis.text.y = element_text(size=12, color = "black"),
        strip.text.x = element_text(size = 10, color = "black", face = "bold"),
        legend.title = element_text(face = "bold", size = 14),
        legend.text = element_text(size = 14, face = "italic", color = "black"),
        legend.key.size = unit(x = 0.25, units = "in"))
  return(out)
}

# with Community
myPlot <- makeStackedBarPlot(dat.melted, legend_length = 16)
myPlot # all pathways with the "community" bar
```

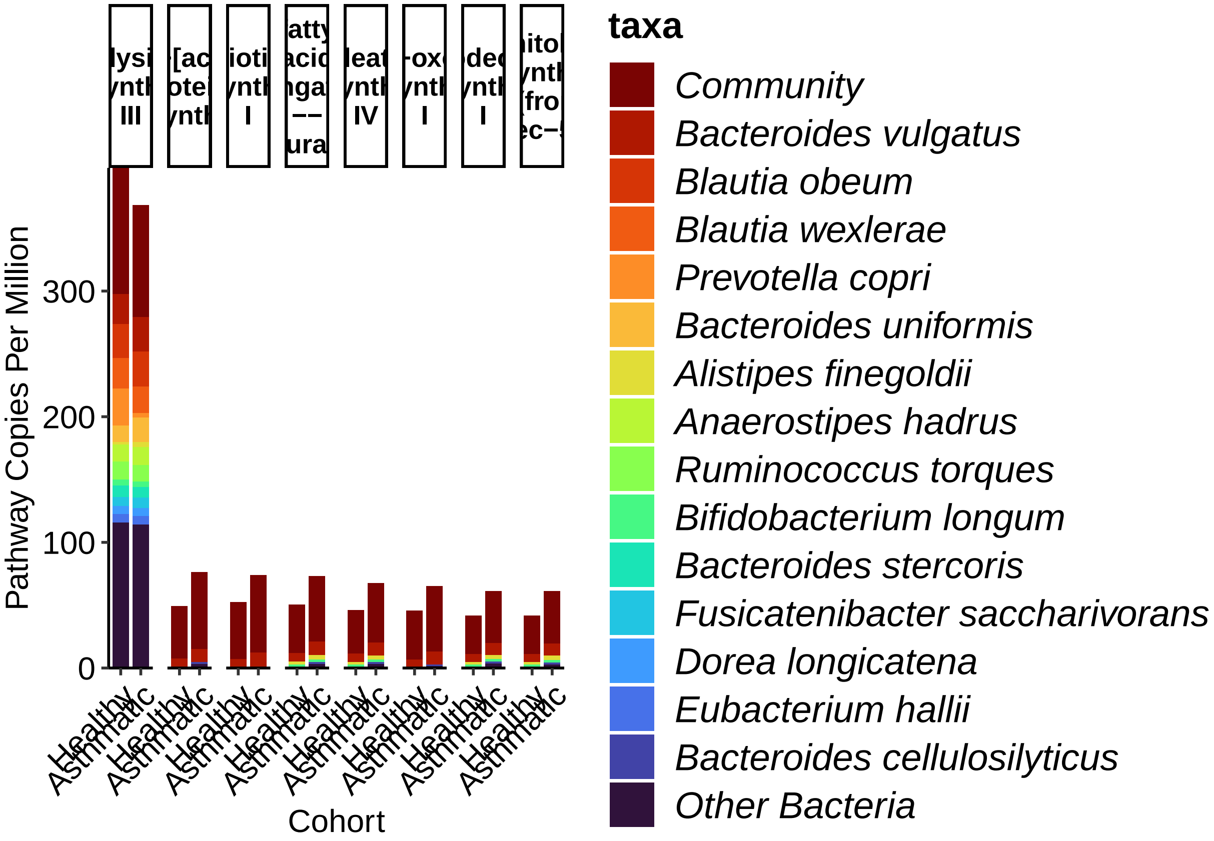


```r
# ggsave(plot = myPlot, filename = "Stacked_Barplots_Averaged_wCommunity_DApwys.jpg",
↪  width = 12, height = 6, units = "in", device = "jpg")
# ggsave(plot = myPlot, filename = "Stacked_Barplots_Averaged_wCommunity_DApwys.pdf",
↪  width = 12, height = 6, units = "in", device = "pdf", useDingbats=F)

# take out Community ####
dat.melted.noOther <- dat.melted[dat.melted$taxa %notin% "Community",]
myPlot2 <- makeStackedBarPlot(dat.melted.noOther, legend_length = 20)
# ggsave(plot = myPlot2, filename = "Stacked_Barplots_Averaged_DApwys.jpg", width = 12,
↪  height = 6, units = "in", device = "jpg")
# ggsave(plot = myPlot2, filename = "Stacked_Barplots_Averaged_DApwys.pdf", width = 12,
↪  height = 6, units = "in", device = "pdf", useDingbats=FALSE)

# take out Lysine ####
dat.melted.noLysine <- dat.melted.noOther[!grepl(pattern = "L-lysine",
↪  x=dat.melted.noOther$PWY),]
myPlot_noLysine <- makeStackedBarPlot(indat = dat.melted.noLysine, legend_length = 11)
print(myPlot_noLysine)
```

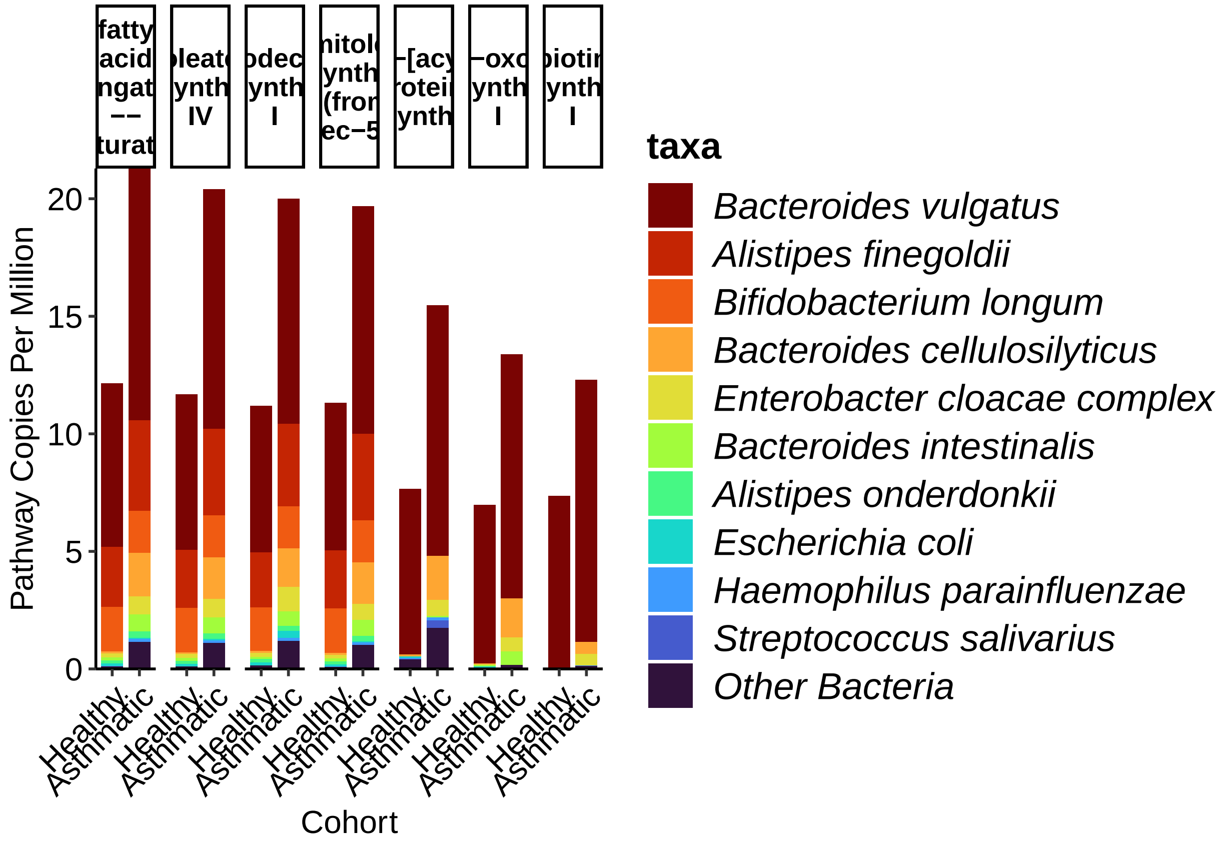


```
dat.melted.noLysine.wComm <- dat.melted[!grepl(pattern = "L-lysine", x=dat.melted$PWY),]
myPlot_noLysine_wComm <- makeStackedBarPlot(indat = dat.melted.noLysine.wComm,
↪  legend_length = 11)
print(myPlot_noLysine_wComm)
```

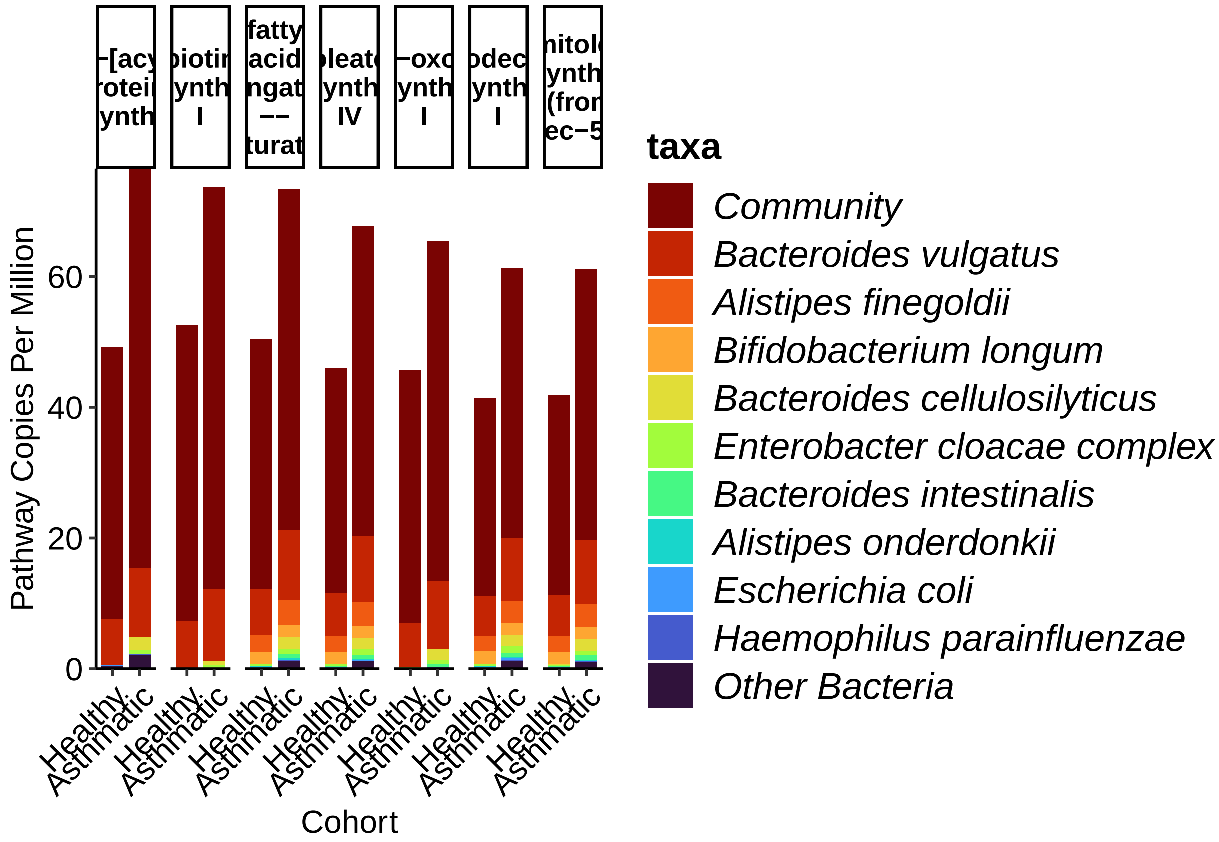


```r
# ggsave(plot = myPlot_noLysine, filename =
↪   "Stacked_Barplots_Averaged_DApwys_noLysine.jpg", width = 12, height = 6, units =
↪   "in", device = "jpg")
# ggsave(plot = myPlot_noLysine, filename =
↪   "Stacked_Barplots_Averaged_DApwys_noLysine.pdf", width = 12, height = 6, units =
↪   "in", device = "pdf", useDingbats=F)
# ggsave(plot = myPlot_noLysine_wComm, filename =
↪   "Stacked_Barplots_Averaged_DApwys_noLysine_wCommunity.jpg", width = 12, height = 6,
↪   units = "in", device = "jpg")
# ggsave(plot = myPlot_noLysine_wComm, filename =
↪   "Stacked_Barplots_Averaged_DApwys_noLysine_wCommunity.pdf", width = 12, height = 6,
↪   units = "in", device = "pdf", useDingbats=FALSE)

# only Lysine ####
dat.melted.onlyLysine <- dat.melted.noOther[grepl(pattern = "L-lysine",
↪   x=dat.melted.noOther$PWY),]
myPlot_onlyLysine <- makeStackedBarPlot(indat = dat.melted.onlyLysine, legend_length =
↪   15)
dat.melted.onlyLysine.wComm <- dat.melted[grepl(pattern = "L-lysine", x=dat.melted$PWY),]
myPlot_onlyLysine_wComm <- makeStackedBarPlot(indat = dat.melted.onlyLysine.wComm,
↪   legend_length = 15)
print(myPlot_onlyLysine)
```

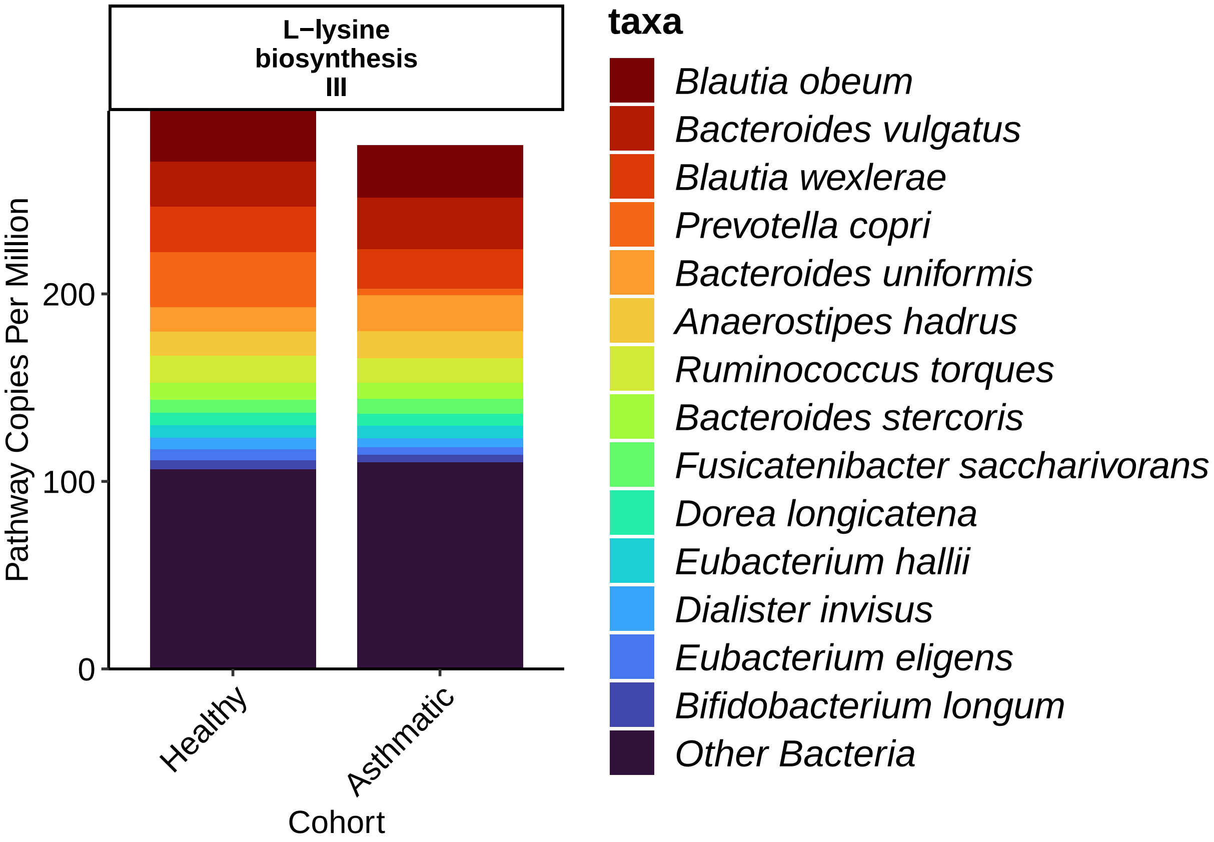


```
print(myPlot_onlyLysine_wComm)
```

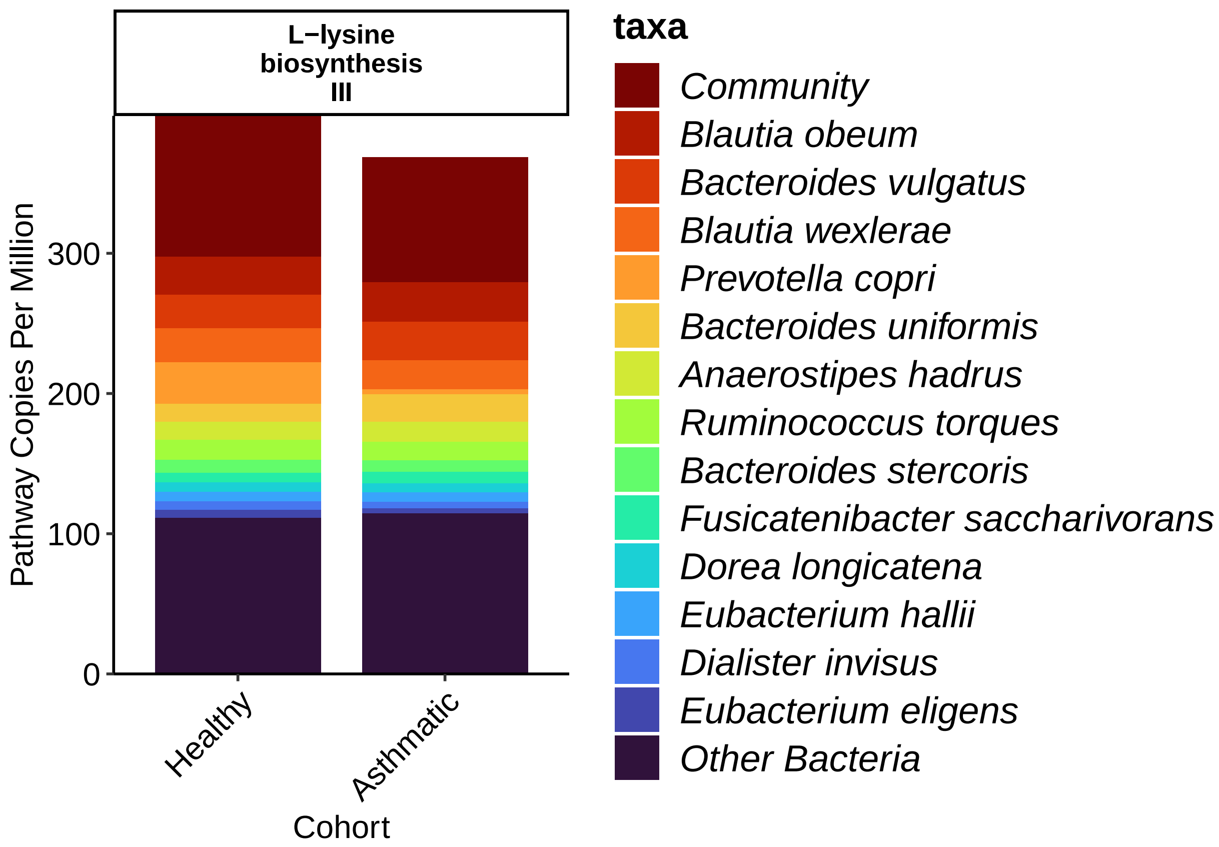


```
# ggsave(plot = myPlot_onlyLysine, filename =
↪   "Stacked_Barplots_Averaged_DApwys_onlyLysine.jpg", width = 5, height = 6, units =
↪   "in", device = "jpg")
# ggsave(plot = myPlot_onlyLysine, filename =
↪   "Stacked_Barplots_Averaged_DApwys_onlyLysine.pdf", width = 5, height = 6, units =
↪   "in", device = "pdf", useDingbats=F)
# ggsave(plot = myPlot_onlyLysine_wComm, filename =
↪   "Stacked_Barplots_Averaged_DApwys_onlyLysine_wCommunity.jpg", width = 5, height = 6,
↪   units = "in", device = "jpg")
# ggsave(plot = myPlot_onlyLysine_wComm, filename =
↪   "Stacked_Barplots_Averaged_DApwys_onlyLysine_wCommunity.pdf", width = 5, height = 6,
↪   units = "in", device = "pdf", useDingbats=FALSE)

# WILCOXONS ####
# sig.pwys
speciesOfInterest <- "Bacteroides_vulgatus"
# speciesOfInterest <- "Alistipes_finegoldii"
filteredtable.melted <-
↪   reshape2::melt(filteredtable[row.names(filteredtable)[grep(row.names(filteredtable),
↪   pattern = "\\|")], ])
filteredtable.melted$taxa <- gsub(".*s__","", perl=T, gsub(".*\\|","", perl=T,
↪   filteredtable.melted$Var1))
filteredtable.melted$Bvul <- ifelse(filteredtable.melted$taxa == speciesOfInterest, TRUE,
↪   FALSE)
filteredtable.melted$PWYshort <- comprehenr::to_vec(for (i in filteredtable.melted$Var1)
↪   str_split(i, pattern = ":")[[1]][1])
filteredtable.melted$PWYshort <- factor(filteredtable.melted$PWYshort)
```

```r
# filteredtable.melted$value <- as.numeric(filteredtable.melted$value)
filteredtable.melted$value <- sapply(filteredtable.melted$value, function(x)
↪  as.numeric(as.character(x)))

filteredtable.melted$BvulvCommunity <- ifelse(filteredtable.melted$taxa ==
↪  speciesOfInterest, speciesOfInterest, ifelse(filteredtable.melted$taxa ==
↪  "Community", "Community", NA))
# filteredtable.melted$BvulvCommunity <- factor(filteredtable.melted$BvulvCommunity)
tmpdf=na.omit(filteredtable.melted) %>%
  group_by(PWYshort) %>%
  summarise(unique(BvulvCommunity)) %>%
  summarise(duplicated(PWYshort)) # you need enough data to have 2 groups to test between
```

```
## `summarise()` has grouped output by 'PWYshort'. You can override using the
## `.groups` argument.
```

```
## `summarise()` has grouped output by 'PWYshort'. You can override using the
## `.groups` argument.
```

```r
filtered_pwys <- tmpdf$PWYshort[tmpdf$`duplicated(PWYshort)`]
filteredtable.melted <- filteredtable.melted[filteredtable.melted$PWYshort %in%
↪  as.character(filtered_pwys),]
Bvul_v_Community <- cbind(na.omit(filteredtable.melted) %>%
  group_by(PWYshort) %>%
  do(w = wilcox.test(value~BvulvCommunity, data=., paired=FALSE)) %>%
  summarise(PWYshort, Wilcox.p = w$p.value),
na.omit(filteredtable.melted) %>%
  group_by(PWYshort, BvulvCommunity) %>%
  summarise(mean=mean(value)) %>%
  summarize(fold_change = mean[1] / mean[2])) # Bvul / Unmapped
```

```
## `summarise()` has grouped output by 'PWYshort'. You can override using the
## `.groups` argument.
```

```r
Bvul_v_Community$Wilcox.q = p.adjust(Bvul_v_Community$Wilcox.p, method = "fdr")
Bvul_v_Community <- Bvul_v_Community[,c(1,4,2,5)]
Bvul_v_Community
```

```
##                 PWYshort fold_change     Wilcox.p     Wilcox.q
## 1 BIOTIN-BIOSYNTHESIS-PWY   0.1708015 1.477891e-22 5.911566e-22
## 2         FASYN-ELONG-PWY   0.1925642 8.985456e-21 1.437673e-20
## 3                PWY-2942   0.2643254 5.447038e-27 4.357630e-26
## 4                PWY-6282   0.2178560 2.161780e-19 2.161780e-19
## 5                PWY-6519   0.1855021 4.117104e-21 1.097894e-20
## 6                PWY-7388   0.1718651 6.974365e-21 1.394873e-20
## 7                PWY-7664   0.2029898 5.590800e-20 7.454399e-20
## 8                PWY0-862   0.2171864 1.690396e-19 1.931881e-19
```

```r
saveRDS(Bvul_v_Community,
↪  "metacyc_stratified_by_species_Bvul_v_Community_wilcox.rds.rds")

# split by asthma
filteredtable.melted$Asthma <-
↪  factor(humann3_mf_filtered[as.character(filteredtable.melted$Var2), "ASTHMA"])
Bvul_v_Community_split <- cbind(na.omit(filteredtable.melted) %>%
```

```
  group_by(PWYshort, Asthma) %>%
  do(w = wilcox.test(value~BvulvCommunity, data=., paired=FALSE)) %>%
  summarise(PWYshort, Wilcox.p = w$p.value, asthma=Asthma),
na.omit(filteredtable.melted) %>%
  group_by(PWYshort, BvulvCommunity) %>%
  summarise(mean=mean(value)) %>%
  summarize(fold_change = mean[1] / mean[2])) # Bvul / Unmapped
```

```
## Warning in wilcox.test.default(x = c(9.56061, 0, 34.0121, 18.8599, 12.101, :
## cannot compute exact p-value with ties

## Warning in wilcox.test.default(x = c(9.1211, 0, 31.3205, 17.7644, 11.4007, :
## cannot compute exact p-value with ties

## Warning in wilcox.test.default(x = c(17.9691, 13.626, 53.8094, 30.2285, : cannot
## compute exact p-value with ties

## Warning in wilcox.test.default(x = c(8.44994, 0, 29.2709, 15.4128, 10.0779, :
## cannot compute exact p-value with ties

## Warning in wilcox.test.default(x = c(9.21939, 0, 31.6519, 17.0672, 10.9877, :
## cannot compute exact p-value with ties

## Warning in wilcox.test.default(x = c(9.94065, 0, 32.5556, 17.1007, 11.2006, :
## cannot compute exact p-value with ties

## Warning in wilcox.test.default(x = c(8.83369, 0, 30.5575, 16.6512, 10.6911, :
## cannot compute exact p-value with ties

## Warning in wilcox.test.default(x = c(8.47752, 0, 29.5962, 15.3672, 9.87188, :
## cannot compute exact p-value with ties

## `summarise()` has grouped output by 'PWYshort'. You can override using the
## `.groups` argument.
```

```
Bvul_v_Community_split$Wilcox.q = p.adjust(Bvul_v_Community_split$Wilcox.p, method =
↪  "fdr")


Bvul_v_allelse <- cbind(filteredtable.melted %>%
  group_by(PWYshort) %>%
  do(w = wilcox.test(value~Bvul, data=., paired=FALSE)) %>%
  summarise(PWYshort, Wilcox = w$p.value),
filteredtable.melted %>%
  group_by(PWYshort, Bvul) %>%
  summarise(mean=mean(value)) %>%
  summarize(fold_change = mean[2] / mean[1])) # Bvul / All else (including unmapped)
```

```
## `summarise()` has grouped output by 'PWYshort'. You can override using the
## `.groups` argument.
```

```
# species to species between asthma/healthy ####
filteredtable.melted$asthma <- humann3_mf[as.character(filteredtable.melted$Var2),
↪  "ASTHMA"]
filteredtable.melted$PWYasthma <- paste0(filteredtable.melted$Var1, "_",
↪  filteredtable.melted$asthma)
speciesAvH <- cbind(filteredtable.melted %>%
  group_by(PWYshort, taxa) %>%
```

```r
  do(w = wilcox.test(value~asthma, data=., paired=FALSE)) %>%
  summarise(PWYshort, taxa, Wilcox.p = w$p.value),
filteredtable.melted %>%
  group_by(PWYshort, taxa, asthma) %>%
  summarise(mean=mean(value), count=sum(value>0)) %>%
  summarize(fold_change = mean[1] / mean[2]) %>% pull(fold_change)) # asthma / healthy
```

```
## `summarise()` has grouped output by 'PWYshort', 'taxa'. You can override using
## the `.groups` argument.
## `summarise()` has grouped output by 'PWYshort'. You can override using the
## `.groups` argument.
```

```r
names(speciesAvH)[dim(speciesAvH)[2]] <- "fold_change_AvH"
speciesAvH$Wilcox.q <- p.adjust(speciesAvH$Wilcox.p, method = "fdr")
# speciesAvH
saveRDS(speciesAvH, "metacyc_stratified_by_species_AvH_wilcox.rds")
write.csv(speciesAvH, "metacyc_stratified_by_species_AvH_wilcox.csv")
speciesAvH[speciesAvH$taxa == "Bacteroides_vulgatus", ]
```

```
##                     PWYshort                 taxa   Wilcox.p fold_change_AvH
## 2   BIOTIN-BIOSYNTHESIS-PWY Bacteroides_vulgatus 0.03300846        1.520993
## 13          FASYN-ELONG-PWY Bacteroides_vulgatus 0.03273270        1.539291
## 59                 PWY-2942 Bacteroides_vulgatus 0.38784203        1.150373
## 181                PWY-6282 Bacteroides_vulgatus 0.03005927        1.545388
## 193                PWY-6519 Bacteroides_vulgatus 0.02839428        1.543572
## 203                PWY-7388 Bacteroides_vulgatus 0.03273270        1.512636
## 215                PWY-7664 Bacteroides_vulgatus 0.03273270        1.539150
## 229                PWY0-862 Bacteroides_vulgatus 0.03273270        1.538672
##      Wilcox.q
## 2   0.2872959
## 13  0.2872959
## 59  0.6323697
## 181 0.2872959
## 193 0.2872959
## 203 0.2872959
## 215 0.2872959
## 229 0.2872959
```

```r
p.adjust(speciesAvH[speciesAvH$taxa == "Bacteroides_vulgatus", "Wilcox.p"], method =
↪  "fdr")
```

```
## [1] 0.03772396 0.03772396 0.38784203 0.03772396 0.03772396 0.03772396 0.03772396
## [8] 0.03772396
```

```r
# speciesAvH[order(speciesAvH$Wilcox.p), ]

# speciesAvH[speciesAvH$fold_change_AvH<Inf &
↪  speciesAvH$Wilcox.p<0.05,][order(speciesAvH[speciesAvH$fold_change_AvH<Inf &
↪  speciesAvH$Wilcox.p<0.05, "fold_change_AvH"], decreasing = T), ]
# speciesAvH[speciesAvH$Wilcox.p<0.05,][order(speciesAvH[speciesAvH$Wilcox.p<0.05,
↪  "fold_change_AvH"], decreasing = T), ]
```

#### Figure S2A:

Looking for asthma-associated pathways

```
source("./humann3_210613_functions_ngw.R")
rm(list = ls()[ls() %notin% c("%notin%", "pwy_barplot", "filter_my_data",
↪   "my_data_summary", "renameMe", "makeBarplotDifferentialPairs")])

library(tidyr)
library(dplyr)
library(reshape2)
library(ggplot2)
library(ggsignif)
library(RColorBrewer)
library(stringr)

# load universal data
humann3_mf_filtered <- readRDS("./humann3_mf_filtered.rds")
humann3_mf = humann3_mf_filtered
row.names(humann3_mf) = humann3_mf$filenamePeriod

my_height = 8
my_width = 9

data_table_name_list = c("eggnogCPM", "koCPM","goCPM", "pfamCPM", "ecCPM") #
↪   c("eggnogRPK", "pfamRPK", "ecRPK", "koRPK","goRPK") # "rxnRPK has no mapping file

# my_data_table_name = data_table_name_list[i]
sort_by = "p-value"
sample_mf = humann3_mf
group_by_me="ASTHMA"
group_by_me2 = "Asthma"
mytest = "wilcox_test"
mycorrection = "fdr"
my_norm_type= "CPM"
basedir="./"
my_color_scheme = brewer.pal(7, "BrBG")[c(7,5)]
i=2
data_table_name_list = "koCPM"

for (my_data_table_name in data_table_name_list){
  print(my_data_table_name)
  stat.wilcoxon.test.save <- readRDS(paste0("~/Library/CloudStorage/Box-Box/Kau
↪   Lab/Results/MARS/FecalMetagenomics/humann3_210613/geneAnalysis/", my_data_table_name,
↪   "_", group_by_me2, "_", mytest, "_", mycorrection, ".rds"))

  my_mappingfile <- read.csv(paste0("~/Library/CloudStorage/Box-Box/Kau
↪   Lab/Results/MARS/FecalMetagenomics/humann3_210613/regrouped_data/map_", gsub(pattern
↪   = "(RPK)|(CPM)", replacement = "", my_data_table_name), "_name.txt"), sep="\t",
↪   header = FALSE, quote="")
  row.names(my_mappingfile) <- my_mappingfile$V1
  TMP.plotdat <- readRDS(file=paste0(basedir, my_data_table_name, ".plotdat.rds"))
# my.df.tmp.means <- readRDS(paste0(basedir, my_data_table_name, ".my.df.means.",
↪   group_by_me, ".rds"))
```

```
  my.df.tmp <- as.data.frame(t(TMP.plotdat))
  my.df.tmp$Group = sample_mf[as.character(row.names(my.df.tmp)), group_by_me]

  # used koala to find Kegg ids for epoxide hydrolase using sequences from paper pmid:
  ↪  31332384
  # see ~/Library/CloudStorage/Box-Box/Kau
  ↪  Lab/Results/MARS/FecalMetagenomics/scaffold_annotation/EHs
if("fullName" %in% names(stat.wilcoxon.test.save)){
  asthmaPathways = c()
  asthmaPathways <- c(stat.wilcoxon.test.save$variables[grep(pattern = "[b|B]ile acid",
↪  stat.wilcoxon.test.save$fullName)],
                      stat.wilcoxon.test.save$variables[grep(pattern = "[h|H]istidine
↪  decarboxylase", stat.wilcoxon.test.save$fullName)],
                      stat.wilcoxon.test.save$variables[grep(pattern = "[e|E]poxide
↪  hydrolase", stat.wilcoxon.test.save$fullName)],
                      stat.wilcoxon.test.save$variables[grep(pattern =
↪  "[t|T]ryptophanase", stat.wilcoxon.test.save$fullName)],
                      stat.wilcoxon.test.save$variables[grep(pattern = "[i|I]ndoleamine",
↪  stat.wilcoxon.test.save$fullName)],
                      # indoleamine 2,3-dioxygenase (IDO) pathway
                      # cytochrome  P450  epoxygenase
                      stat.wilcoxon.test.save$variables[grep(pattern = "[s|S]hort chain
↪  fatty acid", stat.wilcoxon.test.save$fullName)])
    if (grepl(x=my_data_table_name, pattern = "ko")) {
    eh_list <- c("K12952", "K01091", "K07015", "K21064")
    hdc_list <- c("K03758", # antiporter hdcP
                  "K01590") # hdcA
    asthmaPathways <- c(asthmaPathways, eh_list, hdc_list)
  }
  } else {
    asthmaPathways = c()
  asthmaPathways <- c(stat.wilcoxon.test.save$variables[grep(pattern = "[b|B]ile acid",
↪  stat.wilcoxon.test.save$variables)],
                      stat.wilcoxon.test.save$variables[grep(pattern = "[h|H]istidine
↪  decarboxylase", stat.wilcoxon.test.save$variables)],
                      stat.wilcoxon.test.save$variables[grep(pattern = "[e|E]poxide
↪  hydrolase", stat.wilcoxon.test.save$variables)],
                      stat.wilcoxon.test.save$variables[grep(pattern =
↪  "[t|T]ryptophanase", stat.wilcoxon.test.save$variables)],
                      stat.wilcoxon.test.save$variables[grep(pattern = "[i|I]ndoleamine",
↪  stat.wilcoxon.test.save$variables)],
                      # indoleamine 2,3-dioxygenase (IDO) pathway
                      # cytochrome  P450  epoxygenase
                      stat.wilcoxon.test.save$variables[grep(pattern = "[s|S]hort chain
↪  fatty acid", stat.wilcoxon.test.save$variables)])
     if (grepl(x=my_data_table_name, pattern = "ko")) {
    eh_list <- c("K12952", "K01091", "K07015", "K21064")
    hdc_list <- c("K03758", # antiporter hdcP
                  "K01590") # hdcA
    asthmaPathways <- c(asthmaPathways,
                        stat.wilcoxon.test.save$variables[grep(pattern = eh_list[1],
↪  stat.wilcoxon.test.save$variables)],
                        stat.wilcoxon.test.save$variables[grep(pattern = eh_list[2],
↪  stat.wilcoxon.test.save$variables)],
```

```
                        stat.wilcoxon.test.save$variables[grep(pattern = eh_list[3],
↪   stat.wilcoxon.test.save$variables)],
                        stat.wilcoxon.test.save$variables[grep(pattern = eh_list[4],
↪   stat.wilcoxon.test.save$variables)],
                        stat.wilcoxon.test.save$variables[grep(pattern = hdc_list[1],
↪   stat.wilcoxon.test.save$variables)],
                        stat.wilcoxon.test.save$variables[grep(pattern = hdc_list[2],
↪   stat.wilcoxon.test.save$variables)])
     }
    stat.wilcoxon.test.save$fullName <- stat.wilcoxon.test.save$variables
}

                      # stat.wilcoxon.test.save$fullName[grep(pattern = "lipid",
                      ↪   stat.wilcoxon.test.save$fullName)] # 12,13-DiHOME?
  # stat.wilcoxon.test.save$fullName[grep(pattern = "[t|T]oxin",
  ↪   stat.wilcoxon.test.save$fullName)] # toxins?
  library(ggpubr)

  if(length(asthmaPathways)>1){
  print(stat.wilcoxon.test.save[stat.wilcoxon.test.save$variables %in% asthmaPathways,
  ↪   c("fullName", "p", "p.adj", "p.adj.signif")]) # print pvalues

  any(asthmaPathways  %in% names(my.df.tmp))
  my.df.tmp.AsthmaAss <- my.df.tmp[,asthmaPathways[asthmaPathways  %in%
↪   names(my.df.tmp)]]
  renameMe(names(my.df.tmp.AsthmaAss)[1], my_mappingfile)
  # eggnames <- lapply(X = names(my.df.tmp.AsthmaAss), FUN = renameMe, my_mappingfile)
  # eggnames <- unlist(eggnames)
  # colnames(my.df.tmp.AsthmaAss) <- eggnames
  my.df.tmp.AsthmaAss$SampleName <- row.names(my.df.tmp.AsthmaAss)
  my.df.tmp.AsthmaAss$Group = humann3_mf[as.character(row.names(my.df.tmp.AsthmaAss)),
↪   group_by_me]

  my.df.tmp.AsthmaAss

  my.df.tmp.melted.sig <- reshape2::melt(my.df.tmp.AsthmaAss)
  # make bar plot with error bars - ignore the warning here:
  library(reshape2)
  my.df.tmp.melted.sig.forplotErrorBars <- my_data_summary(my.df.tmp.melted.sig, varname
↪   = "value", groupnames = c("Group", "variable"))
  # add se for eror bars
  if(length(my.df.tmp$Group) > 0){
  my.df.tmp.melted.sig.forplotErrorBars$se <-
    as.numeric(ifelse(my.df.tmp.melted.sig.forplotErrorBars$Group ==
    ↪   levels(factor(my.df.tmp.melted.sig.forplotErrorBars$Group))[1],
           my.df.tmp.melted.sig.forplotErrorBars$sd/sqrt(sum(my.df.tmp$Group ==
↪   levels(factor(my.df.tmp.melted.sig.forplotErrorBars$Group))[1])),
           ifelse(my.df.tmp.melted.sig.forplotErrorBars$Group ==
            ↪   levels(factor(my.df.tmp.melted.sig.forplotErrorBars$Group))[2],
           my.df.tmp.melted.sig.forplotErrorBars$sd/sqrt(sum(my.df.tmp$Group ==
↪   levels(factor(my.df.tmp.melted.sig.forplotErrorBars$Group))[2])), "ERROR")))
  } else {print("my.df.tmp missing Group column.")}
```

```
my.df.tmp.melted.sig.forplotErrorBars$variable <-
↪  factor(my.df.tmp.melted.sig.forplotErrorBars$variable, levels =
↪  rev(unique(my.df.tmp.melted.sig.forplotErrorBars$variable)))
AsthmaAssgeombarError <- ggplot(my.df.tmp.melted.sig.forplotErrorBars,
           aes(x=variable, y=value, fill=Group)) +
  geom_bar(stat="identity", color="black",
           position=position_dodge()) +
  geom_errorbar(aes(ymin=value-se, ymax=value+se), width=.2,
                 position=position_dodge(.9)) +
  stat_compare_means(data=my.df.tmp.melted.sig, method="wilcox.test", aes(label =
   ↪  ..p.signif..), label.y = max(my.df.tmp.melted.sig.forplotErrorBars$value)*1.1) +
  labs(title=paste0(gsub(my_data_table_name, pattern = "(RPK)|(CPM)", replacement =
   ↪  ""), " Asthma-associated"), x=my_data_table_name, y=my_norm_type)+
   theme_classic() +
  scale_x_discrete(labels = function(x) str_wrap(x, width = 30)) +
   scale_fill_manual(values=my_color_scheme) +
    coord_flip() +
  theme(legend.text = element_text(size = 18), axis.text=element_text(size=16, color =
   ↪  "black"),
          axis.title = element_text(size = 18), text=element_text(size = 16, color =
           ↪  "black"))
print(AsthmaAssgeombarError)
  # ggsave(AsthmaAssgeombarError,
  #         filename = paste0("./", my_data_table_name,
   ↪  "_AsthmaAssGeombarSE_ASTHMA_wTryp.jpg"),
  #         height = my_height, width = my_width, units = "in",
  #         device = "jpg")
  # ggsave(AsthmaAssgeombarError,
  #         filename = paste0("./", my_data_table_name,
   ↪  "_AsthmaAssGeombarSE_ASTHMA_wTryp.pdf"),
  #         height = my_height, width = my_width, units = "in",
  #         device = "pdf", useDingbats = FALSE)
} else {
  if(length(asthmaPathways)==1){
    my.df.tmp.AsthmaAss <- my.df.tmp[,asthmaPathways[asthmaPathways  %in%
↪  names(my.df.tmp)]]
    my.df.tmp.AsthmaAss <- data.frame(as.numeric(my.df.tmp.AsthmaAss),
↪  row.names(my.df.tmp), as.character(humann3_mf[as.character(row.names(my.df.tmp)),
↪  group_by_me]), rep(asthmaPathways, length(my.df.tmp.AsthmaAss)))
    names(my.df.tmp.AsthmaAss) <- c("value", "SampleName", "Group" , "Function")
    singlefunctionname = renameMe(asthmaPathways, my_mappingfile)
    my.df.tmp.AsthmaAss$Function <- rep(singlefunctionname,
↪  length(my.df.tmp.AsthmaAss$Function))
    my.df.tmp.melted.sig <- reshape2::melt(my.df.tmp.AsthmaAss)
# make bar plot with error bars - ignore the warning here:
my.df.tmp.melted.sig.forplotErrorBars <- my_data_summary(my.df.tmp.melted.sig, varname
↪  = "value", groupnames = c("Group", "Function"))
# add se for eror bars
if(length(my.df.tmp$Group) > 0){
my.df.tmp.melted.sig.forplotErrorBars$se <-
  as.numeric(ifelse(my.df.tmp.melted.sig.forplotErrorBars$Group ==
   ↪  levels(factor(my.df.tmp.melted.sig.forplotErrorBars$Group))[1],
         my.df.tmp.melted.sig.forplotErrorBars$sd/sqrt(sum(my.df.tmp$Group ==
↪  levels(factor(my.df.tmp.melted.sig.forplotErrorBars$Group))[1])),
```

```
          ifelse(my.df.tmp.melted.sig.forplotErrorBars$Group ==
           ↪  levels(factor(my.df.tmp.melted.sig.forplotErrorBars$Group))[2],
          my.df.tmp.melted.sig.forplotErrorBars$sd/sqrt(sum(my.df.tmp$Group ==
↪  levels(factor(my.df.tmp.melted.sig.forplotErrorBars$Group))[2])), "ERROR")))
  } else {print("my.df.tmp missing Group column.")}

  my.df.tmp.melted.sig.forplotErrorBars$Function <-
↪  factor(my.df.tmp.melted.sig.forplotErrorBars$Function, levels =
↪  rev(unique(my.df.tmp.melted.sig.forplotErrorBars$Function)))

  AsthmaAssgeombarError <- ggplot(my.df.tmp.melted.sig.forplotErrorBars,
            aes(x=Function, y=value, fill=Group)) +
    geom_bar(stat="identity", color="black",
            position=position_dodge()) +
    geom_errorbar(aes(ymin=value-se, ymax=value+se), width=.2,
                  position=position_dodge(.9)) +
    stat_compare_means(data=my.df.tmp.melted.sig, method="wilcox.test", aes(label =
    ↪  ..p.signif..), label.y = max(my.df.tmp.melted.sig.forplotErrorBars$value)*1.1) +
    labs(title=paste0(gsub(my_data_table_name, pattern = "RPK", replacement = ""), "
    ↪  Asthma-associated"), x=my_data_table_name, y="Relative Abundance (mean RPK)")+
     theme_classic() +
    scale_x_discrete(labels = function(x) str_wrap(x, width = 30)) +
     scale_fill_manual(values=my_color_scheme) +
      coord_flip() +
    theme(legend.text = element_text(size = 18), axis.text=element_text(size=16, color =
    ↪  "black"),
          axis.title = element_text(size = 18), text=element_text(size = 16, color =
          ↪  "black"))
  print(AsthmaAssgeombarError)
    # ggsave(AsthmaAssgeombarError,
    #         filename = paste0("./", my_data_table_name, "_AsthmaAssGeombarSE.jpg"),
    #        height = my_height, width = my_width, units = "in",
    #        device = "jpg")
    # ggsave(AsthmaAssgeombarError,
    #         filename = paste0("./", my_data_table_name,
    ↪  "_AsthmaAssGeombarSE_ASTHMA.pdf"),
    #        height = my_height, width = my_width, units = "in",
    #        device = "pdf", useDingbats = FALSE)

    } else {
      print("No pathways matched.")
     }
  }
}
```

```
## [1] "koCPM"
##                                                fullName       p     p.adj
## 342  K01091: phosphoglycolate phosphatase [EC:3.1.3.18] 0.91100 0.9891843
## 487       K01590: histidine decarboxylase [EC:4.1.1.22] 0.86300 0.9803710
## 526                 K01667: tryptophanase [EC:4.1.99.1] 0.00258 0.4594180
## 1176       K03453: bile acid:Na+ symporter, BASS family 0.24800 0.9025384
## 1342             K03758: arginine:ornithine antiporter 0.52800 0.9473664
## 1713                                    K07015: NO_NAME 0.41400 0.9316563
```

```
## 2194                                    K12952: NO_NAME 0.79400 0.9716005
## 2299   K15868: bile acid-coenzyme A ligase [EC:6.2.1.7] 0.38900 0.9283348
## 2516                                    K21064: NO_NAME 0.76300 0.9650477
##      p.adj.signif
## 342            ns
## 487            ns
## 526            ns
## 1176           ns
## 1342           ns
## 1713           ns
## 2194           ns
## 2299           ns
## 2516           ns

## Using SampleName, Group as id variables
```

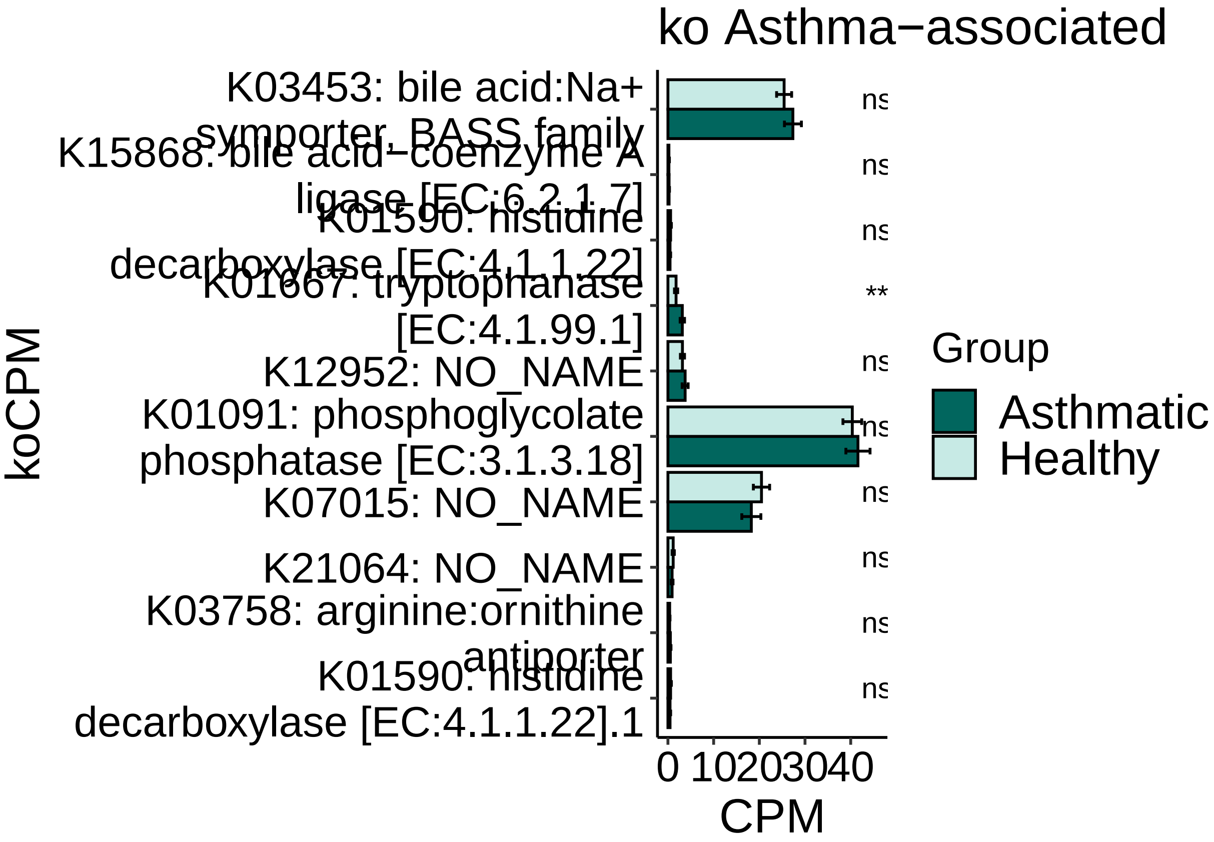


#### Figure 1G:

Asthma by Age Group Barplots

```
rm(list = ls())
source("./humann3_210613_functions_ngw.R")

library(tidyverse)
library(reshape2)
library(RColorBrewer)
library(ggpubr)
```

```r
library(stringr)

stat.wilcoxon <- readRDS("metacycpwy_Asthma_wilcox_test_fdr.rds")
stat.wilcoxon <- stat.wilcoxon[order(stat.wilcoxon$pval, decreasing = FALSE),]
stat.wilcoxon <- data.frame(stat.wilcoxon)
stat.wilcoxon.sig <- stat.wilcoxon[stat.wilcoxon$padj < 0.2,]
mypwys <- readRDS(file = "MARS_metacyc_pwys_CPM_filtered.rds")
mean_by_group_pwys_asthma <- readRDS(file = paste0("METACYC_PWYS", ".my.df.means.",
↪  "ASTHMA", ".rds"))
humann3_mf_filtered <- readRDS("./humann3_mf_filtered.rds")
row.names(humann3_mf_filtered) <- humann3_mf_filtered$SampleName
humann3_mf <- humann3_mf_filtered
humann3_mf$Cohort = paste0(as.character(humann3_mf$ASTHMA), " ", humann3_mf$AgeGroup)

topSigPWYS <- as.character(stat.wilcoxon.sig[, "PWY"]) #
↪  unique(c(as.character(stat.wilcoxon[1:10, "PWY"]), superpwys, moreFFAs))
topSigPWYS <- names(mean_by_group_pwys_asthma[,
↪  topSigPWYS][,order(colSums(mean_by_group_pwys_asthma[,topSigPWYS]), decreasing = T)])
↪  #reorder by abundance
# topSigPWYS <- names(mean_by_group_pwys_asthma[,
↪  topSigPWYS][,order(stat.wilcoxon.sig[topSigPWYS, "padj"], decreasing = T)]) #reorder
↪  by sig

group_by_me = "ASTHMA"
# my_pair_test = "wilcox_test"
# my_padjust_method="fdr"
my_color_scheme = c(brewer.pal(7, "BrBG")[c(7,1,5,3)])

forTtests = as.data.frame(t(mypwys))
# add asthma labels as last column
forTtests$Group = as.character(humann3_mf[row.names(forTtests), group_by_me])
forTtests$AgeGroup = as.character(humann3_mf[row.names(forTtests), "AgeGroup"])
forTtests$Cohort = paste0(forTtests$Group, " ", forTtests$AgeGroup)
forTtests$Sample = row.names(forTtests)

forTtests.melted <- reshape2::melt(forTtests)
```

```
## Using Group, AgeGroup, Cohort, Sample as id variables
```

```r
# rpkm.aov <- aov(value~Group*AgeGroup, forTtests.melted)
# summary(rpkm.aov)
asthma_pvals <- c()
agegroup_pvals <- c()
interaction_pvals <- c()
incidences_VFs <- c()
for (virfac in topSigPWYS) {
  forTtests.virfac <- forTtests[,c(virfac, "Group", "AgeGroup", "Sample")]
  forTtests.virfac$Cohort <- paste0(forTtests.virfac$Group, " ",
↪  forTtests.virfac$AgeGroup)
  forTtests.virfac.melted <- reshape2::melt(forTtests.virfac)
  incidence.virfac <- sum(forTtests.virfac.melted$value>0)
  incidences_VFs <- c(incidences_VFs, incidence.virfac)
  rpkm.aov.virfac <- aov(value~Group*AgeGroup, forTtests.virfac.melted)
  # rpkm.aov.virfac <- aov(value~Cohort, forTtests.virfac.melted)
```

```
  p.Asthma <- summary(rpkm.aov.virfac)[[1]][["Pr(>F)"]][1]
  asthma_pvals = c(asthma_pvals, p.Asthma)
  p.Age <- summary(rpkm.aov.virfac)[[1]][["Pr(>F)"]][2]
  agegroup_pvals <- c(agegroup_pvals, p.Age)
  p.Int <- summary(rpkm.aov.virfac)[[1]][["Pr(>F)"]][3]
  interaction_pvals <- c(interaction_pvals, p.Int)
}
```

```
## Using Group, AgeGroup, Sample, Cohort as id variables
```

```
## Using Group, AgeGroup, Sample, Cohort as id variables
## Using Group, AgeGroup, Sample, Cohort as id variables
## Using Group, AgeGroup, Sample, Cohort as id variables
## Using Group, AgeGroup, Sample, Cohort as id variables
## Using Group, AgeGroup, Sample, Cohort as id variables
## Using Group, AgeGroup, Sample, Cohort as id variables
## Using Group, AgeGroup, Sample, Cohort as id variables
```

```
rpkm_anova <- data.frame(topSigPWYS, asthma_pvals, agegroup_pvals, interaction_pvals,
↪  incidences_VFs)
rpkm_anova[as.numeric(as.character(rpkm_anova$asthma_pvals)) <0.05, ]
```

```
##                                                                    topSigPWYS
## 1                                          PWY-2942: L-lysine biosynthesis III
## 2                               BIOTIN-BIOSYNTHESIS-PWY: biotin biosynthesis I
## 3 PWY-7388: octanoyl-[acyl-carrier protein] biosynthesis (mitochondria, yeast)
## 4                           FASYN-ELONG-PWY: fatty acid elongation -- saturated
## 5                                  PWY-7664: oleate biosynthesis IV (anaerobic)
## 6                               PWY-6519: 8-amino-7-oxononanoate biosynthesis I
## 7             PWY-6282: palmitoleate biosynthesis I (from (5Z)-dodec-5-enoate)
## 8                                     PWY0-862: (5Z)-dodecenoate biosynthesis I
##   asthma_pvals agegroup_pvals interaction_pvals incidences_VFs
## 1  0.003851652      0.9130231         0.2187927             95
## 2  0.007798932      0.6785836         0.3472974             91
## 3  0.003337180      0.6350080         0.6523733             91
## 4  0.006487946      0.4722420         0.4122261             91
## 5  0.006255332      0.4554774         0.4206873             91
## 6  0.008038307      0.5321971         0.3697993             91
## 7  0.007273058      0.5018286         0.3892233             91
## 8  0.006291923      0.4114734         0.4481150             93
```

```
saveRDS(rpkm_anova, file = "metacyc_agegroup_twoWayAnovas.rds")
write.csv(rpkm_anova, file = "metacyc_agegroup_twoWayAnovas.csv")

rpkm_anova <- data.frame(rpkm_anova)
list_to_plot <- as.character(topSigPWYS)

my.df.tmp = as.data.frame(t(mypwys[,row.names(humann3_mf)]))
my.df.tmp.list_to_plot <- my.df.tmp[,list_to_plot]
my.df.tmp.list_to_plot$SampleName <- row.names(my.df.tmp.list_to_plot)
group_by_me = "Cohort"
my.df.tmp.list_to_plot$Group =
↪  humann3_mf[as.character(row.names(my.df.tmp.list_to_plot)), group_by_me]
# Take top pathways, melt for ggplot
my.df.tmp.melted.sig <- reshape2::melt(my.df.tmp.list_to_plot)
```

```
## Using SampleName, Group as id variables
```

```
my.df.tmp.melted.sig$Group <- factor(my.df.tmp.melted.sig$Group)
# make bar plot with error bars - ignore the warning here:
my.df.tmp.melted.sig.forplotErrorBars <- my_data_summary(my.df.tmp.melted.sig, varname =
↪  "value", groupnames = c("Group", "variable"))
my.df.tmp$Group = humann3_mf[row.names(my.df.tmp), group_by_me]
# add se for eror bars
my.df.tmp.melted.sig.forplotErrorBars$se <-
  as.numeric(ifelse(my.df.tmp.melted.sig.forplotErrorBars$Group ==
   ↪  levels(factor(my.df.tmp.melted.sig.forplotErrorBars$Group))[1],
                      my.df.tmp.melted.sig.forplotErrorBars$sd/sqrt(sum(my.df.tmp$Group
↪  == levels(factor(my.df.tmp.melted.sig.forplotErrorBars$Group))[1])),
                      ifelse(my.df.tmp.melted.sig.forplotErrorBars$Group ==
                      ↪  levels(factor(my.df.tmp.melted.sig.forplotErrorBars$Group))[2],
                      ↪

↪  my.df.tmp.melted.sig.forplotErrorBars$sd/sqrt(sum(my.df.tmp$Group ==
↪  levels(factor(my.df.tmp.melted.sig.forplotErrorBars$Group))[2])),
                             ifelse(my.df.tmp.melted.sig.forplotErrorBars$Group ==
                             ↪  levels(factor(my.df.tmp.melted.sig.forplotErrorBars$Group))[3],
                             ↪

↪  my.df.tmp.melted.sig.forplotErrorBars$sd/sqrt(sum(my.df.tmp$Group ==
↪  levels(factor(my.df.tmp.melted.sig.forplotErrorBars$Group))[3])),
                             ifelse(my.df.tmp.melted.sig.forplotErrorBars$Group ==
                             ↪  levels(factor(my.df.tmp.melted.sig.forplotErrorBars$Group))[4],
                             ↪

↪  my.df.tmp.melted.sig.forplotErrorBars$sd/sqrt(sum(my.df.tmp$Group ==
↪  levels(factor(my.df.tmp.melted.sig.forplotErrorBars$Group))[4])), "ERROR")))))

  my.df.tmp.melted.sig.forplotErrorBars$variable <-
↪  factor(my.df.tmp.melted.sig.forplotErrorBars$variable, levels =
↪  rev(unique(my.df.tmp.melted.sig.forplotErrorBars$variable)))
  library(stringr)
  top25geombarError<- ggplot(my.df.tmp.melted.sig.forplotErrorBars,
                             aes(x=variable, y=value, fill=Group)) +
    geom_bar(stat="identity", color="black",
             position=position_dodge()) +
    geom_errorbar(aes(ymin=value-se, ymax=value+se), width=.2,
                  position=position_dodge(.9)) +
    labs(title="Significant in Asthma",
         x="MetaCyc Pathway",
         y="Relative Abundance (CPM)")+
    theme_classic() +
    theme(legend.text = element_text(size = 18), axis.text=element_text(size=12, color =
    ↪  "black"),
          axis.title = element_text(size = 18))+
    scale_fill_manual(values=my_color_scheme) +
    coord_flip()
print(top25geombarError)
```

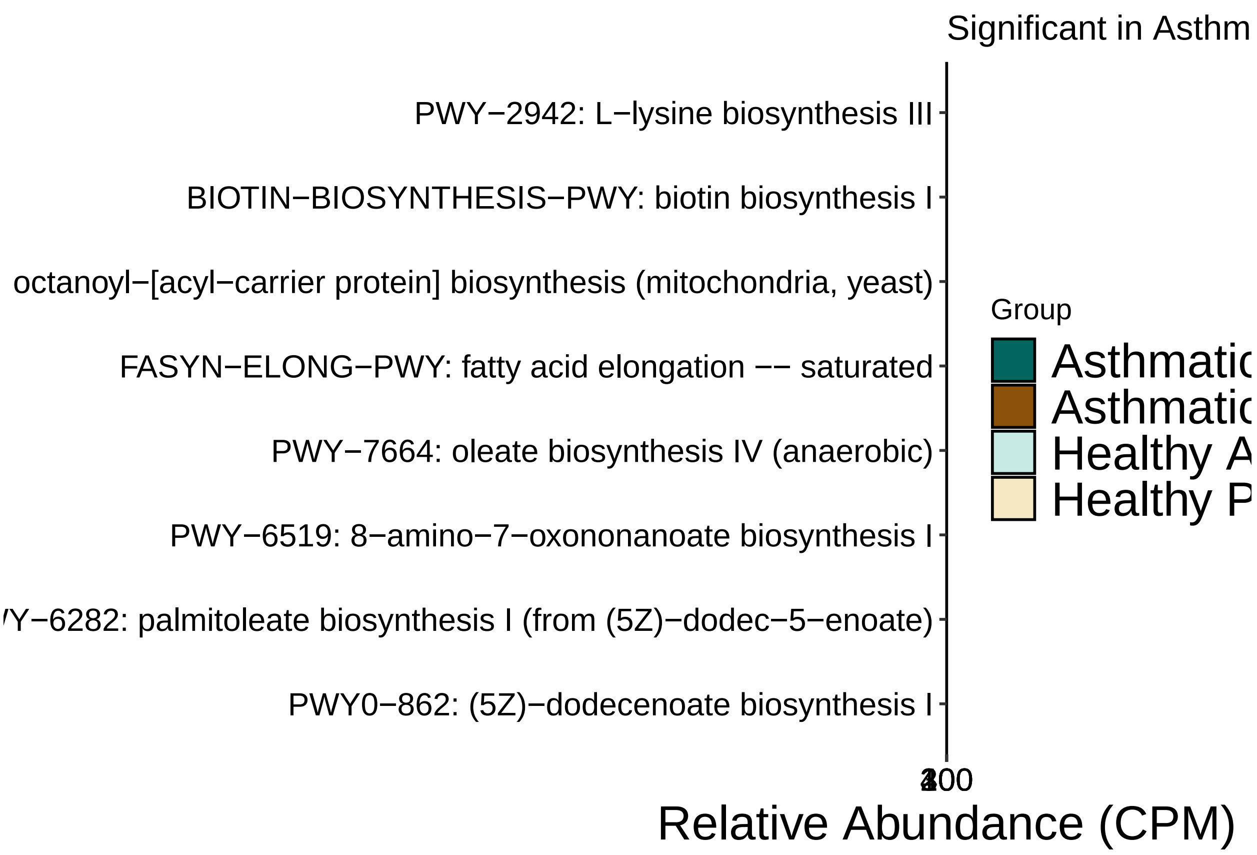


```
# ggsave(plot = top25geombarError, filename =
↪  paste0("metacyc_cpm_barplot_ANOVA_fattyAcids_4Group_",group_by_me,".jpg"), width=14,
↪  height=7, units="in", device = "jpg")
# ggsave(plot = top25geombarError, filename =
↪  paste0("metacyc_cpm_barplot_ANOVA_fattyAcids_4Group_",group_by_me,".pdf"), width=14,
↪  height=7, units="in", device = "pdf", useDingbats = FALSE)
```

#Libraries

```
##
## Attaching package: 'ape'

## The following object is masked from 'package:ggpubr':
##
##     rotate
```

#Functions for metaphlan analysis

```
#### FUNCTIONS ####
# rm(list=ls()[ls() %notin% "%notin%"])
rm(list=ls())
source("./humann3_210613_functions_ngw.R")

makeMARSPlot <- function(datframe, x_column, y_column, y_title="", expt_column,
↪  microbiome_column, shape_int=16){
  require(RColorBrewer)
  # ex usage:
  # makeMARSPlot(igeplotdat, "asthma", "final_ngml", "OVA-IgE", "experiment",
  ↪  "microbiome")
```

```
  # x_column can be any two values (I use "asthma" and "Healthy")
  # y_column are the numeric values
  # expt_column must be "HGF2" etc
  # microbiome_column is "MARS0022" and "MARS0043"
  # all column names in strings

  outplot <- ggplot(datframe, aes(x=eval(parse(text=x_column)),
↪   y=eval(parse(text=y_column)))) +
    geom_boxplot(outlier.shape=NA, lwd=0.75) +
    geom_point(position=position_beeswarm(cex=4),
    ↪  aes(shape=eval(parse(text=expt_column)),

                                                  ↪  fill=eval(parse(text=expt_column))),
                                                  ↪  size=3.5) +
    scale_fill_manual(name="asthmaPatient",
                      labels=c("Healthy", "Asthmatic"),
                       values = c(brewer.pal(7, "BrBG")[c(5,7)]))+
                      # values=c("#DE4968FF", "#51127CFF"))+# "#DE4968FF", "#51127CFF",
                      ↪  "#51127CFF")) +
    scale_shape_manual(name="asthmaPatient",
                      labels=c("Healthy", "Asthmatic"),
                      values=c(21, 21)) +
    scale_color_manual(name="asthmaPatient",
                      labels=c("Healthy", "Asthmatic"),
                      values=c("black", "black")) +
    scale_x_discrete(name = "Microbiota") +
    scale_y_continuous(name=y_title) +
    theme(panel.grid.major = element_blank(),
          panel.grid.minor = element_blank(),
          axis.line.x = element_line(size = 0, colour = "black"),
          axis.line.y = element_line(size = 0, colour = "black"),
          axis.line = element_line(size=1, colour = "black"),
          panel.border = element_rect(fill = NA, colour = "black", size = 1),
          panel.background = element_blank(),
          text=element_text(size = 16),
          axis.text.x=element_text(colour="black", size = 12),
          axis.text.y=element_text(colour="black", size = 12),
          axis.title.x=element_blank(),
          legend.title=element_blank(),
          legend.key = element_rect(color=NA, fill=NA)
    ) +
    stat_compare_means(method = "t.test", aes(label = "p.signif"), label.x = 1.5)##,
    ↪  label.x = 1.5, label.y = 3.5) # or use label = ..p.signif.. for just the asterisk

  outplot
}
```

#Loading in the Data

```
#Import all merged abundance table
indir <- "~/Library/CloudStorage/Box-Box/Kau
↪  Lab/Results/MARS/FecalMetagenomics/humann3_210613/"
allTaxa <- read.table(file=paste0(indir, "MARS_bugs_relabundance_table.txt"), header =
↪  TRUE, sep = "\t")
```

```r
names(allTaxa) <- c(names(allTaxa)[1:2], gsub(pattern =
↪   "_interleaved_metaphlan_bugs_list", replacement = "",
↪   names(allTaxa)[3:length(names(allTaxa))]))
allGenera <- read.table(file=paste0(indir, "merged_abundance_genera_fullnames.txt"),
↪   header = FALSE, sep = "\t")
colnames(allGenera) <- colnames(allTaxa)
allSpecies = allTaxa[grep(pattern = "\\|s__", x = allTaxa$clade_name),]
allGeneraImported = allGenera
#Import Human Sample Database
# humanSamples <- read.table(file = "~/Library/CloudStorage/Box-Box/Kau
↪   Lab/Results/MARS/FecalMetagenomics/191219_MARSxFecal_Nextera_mf.txt",
#                              sep = "\t", header = TRUE)

##Adjust merged abundance table
row.names(allSpecies) <- allSpecies$clade_name # or ID by default
allSpecies$clade_name <- NULL
allSpecies$NCBI_tax_id <- NULL
allSpecies <- allSpecies/colSums(allSpecies) # get relative abundance between 0 and 1
saveRDS(allSpecies, "allSpecies_104_relab_tab.rds")

row.names(allGenera) <- allGenera$clade_name # or ID by default
allGenera$clade_name <- NULL
allGenera$NCBI_tax_id <- NULL
allGenera <- allGenera/colSums(allGenera) # get relative abundance between 0 and 1
saveRDS(allGenera, "allGenera_104_relab_tab.rds")
#### drop all disqualified samples from all dataframes: ####
humann3_mf_filtered <- readRDS("./humann3_mf_filtered.rds")
row.names(humann3_mf_filtered) <- gsub(humann3_mf_filtered$SampleName, pattern = "\\-",
↪   replacement = ".")
allGenera <- allGenera[,colnames(allGenera) %in% row.names(humann3_mf_filtered)]
allSpecies <- allSpecies[,colnames(allSpecies) %in% row.names(humann3_mf_filtered)]
allTaxa <- allTaxa[,colnames(allTaxa) %in% c(row.names(humann3_mf_filtered),
↪   "clade_name", "NCBI_tax_id")]
sum(colnames(allSpecies) %in% row.names(humann3_mf_filtered)) == 95
```

```
## [1] TRUE
```

```r
sum(colnames(allGenera) %in% row.names(humann3_mf_filtered)) == 95
```

```
## [1] TRUE
```

```r
sum(names(allTaxa) %in% row.names(humann3_mf_filtered)) == 95
```

```
## [1] TRUE
```

```r
saveRDS(allSpecies, "allSpecies_95_relab_tab.rds")
saveRDS(allTaxa, "allTaxa_95_relab_tab.rds")
```

### Figure S1D:

Plot top 20 Genera

```r
rm(list=ls()[ls() %notin% c("allGenera", "makeMARSPlot")])
plotAsvs <- allGenera
```

```r
row.names(plotAsvs) <- gsub(pattern = "^.*g__", replacement = "", row.names(plotAsvs))
plotAsvs <- plotAsvs[order(rowSums(plotAsvs), decreasing=T),] #reorder by abundance
plotAsvs <- plotAsvs[, order(plotAsvs[1,], decreasing = T)] # reorder by top genre
↪  abundance
sampleFactor <- names(plotAsvs)
top25toPlot <- plotAsvs[1:19,]
top25OthersSum <- 1-colSums(top25toPlot)
# sampleFactor <- names(top25OthersSum[order(top25OthersSum, decreasing=T)])
top25toPlot <- rbind(top25toPlot, top25OthersSum)
row.names(top25toPlot)[dim(top25toPlot)[1]] <- "Others"
myFactor <- as.character(row.names(top25toPlot))
top25toPlot[,"Genus"] <- myFactor
top25toPlot$Genus <- factor(gsub(top25toPlot$Genus, pattern = "_unclassified",
↪  replacement=""), levels = rev(gsub(myFactor, pattern = "_unclassified",
↪  replacement="")))
# top25toPlot.melted$Genus <- gsub(top25toPlot.melted$Genus, pattern = "_unclassified",
↪  replacement="")
top25toPlot.melted <- melt(top25toPlot, id.vars = "Genus", na.rm = F)
humanSamples <- readRDS("./humann3_mf_filtered.rds")
row.names(humanSamples) <- humanSamples$SampleName
top25toPlot.melted$Asthma <-
↪  factor(humanSamples[as.character(gsub(top25toPlot.melted$variable,
                                                                    pattern="[.]",
                                                                     ↪  replacement="-")),"ASTHMA"],
                                                                     ↪
                                     levels = c("Healthy", "Asthmatic"),
                                     labels=c("Healthy", "Asthmatic"))
top25toPlot.melted$AgeGroup <-
  factor(humanSamples[as.character(gsub(top25toPlot.melted$variable, pattern="[.]",
   ↪  replacement="-")), "AgeGroup"])
top25toPlot.melted$variable <- factor(top25toPlot.melted$variable, levels=sampleFactor)
sum(is.na(top25toPlot.melted))
```

```
## [1] 0
```

```r
row.has.na <- apply(top25toPlot.melted, 1, function(x){any(is.na(x))})
sum(row.has.na)
```

```
## [1] 0
```

```r
top25toPlot.melted <- na.omit(top25toPlot.melted)
sum(is.na(top25toPlot.melted))
```

```
## [1] 0
```

```r
row.has.na <- apply(top25toPlot.melted, 1, function(x){any(is.na(x))})
sum(row.has.na)
```

```
## [1] 0
```

```r
library(viridis)
library(viridisLite)
# by age and asthma

top25toPlot.melted$cohort <- factor(paste0(top25toPlot.melted$Asthma," ",
↪  top25toPlot.melted$AgeGroup), levels=c("Healthy Pediatric", "Healthy Adult",
↪  "Asthmatic Pediatric", "Asthmatic Adult"))
```

```
ageasthmaTrampStamp <- ggplot(top25toPlot.melted, aes(color=Genus, fill=Genus, x =
↪  variable, y = value)) +
  geom_bar(stat="identity", position = "stack", width = 0.8) +
  theme_classic() +
  scale_fill_viridis(option="turbo", discrete = T)+
  scale_color_viridis(option="turbo", discrete = T)+
  facet_grid( ~ cohort, scales = "free_x", space = "free_x") +
  scale_y_continuous(limits=c(0,1), expand = c(0,0))  +
  xlab(label = "MARS Subjects") + ylab("Relative Abundance")+
  theme(text = element_text(size=rel(3.5)),
        strip.text.x = element_text(size=rel(3.5)),
        axis.ticks.x = element_blank(),
        axis.text.x=element_blank(),
        legend.title = element_text(face = "bold", size = 10),
        legend.text = element_text(size = 10, face = "italic"),
        legend.key.size = unit(x = 0.4, units = "cm"),
        legend.position='bottom',
        legend.direction = "horizontal"
        )
ageasthmaTrampStamp
```

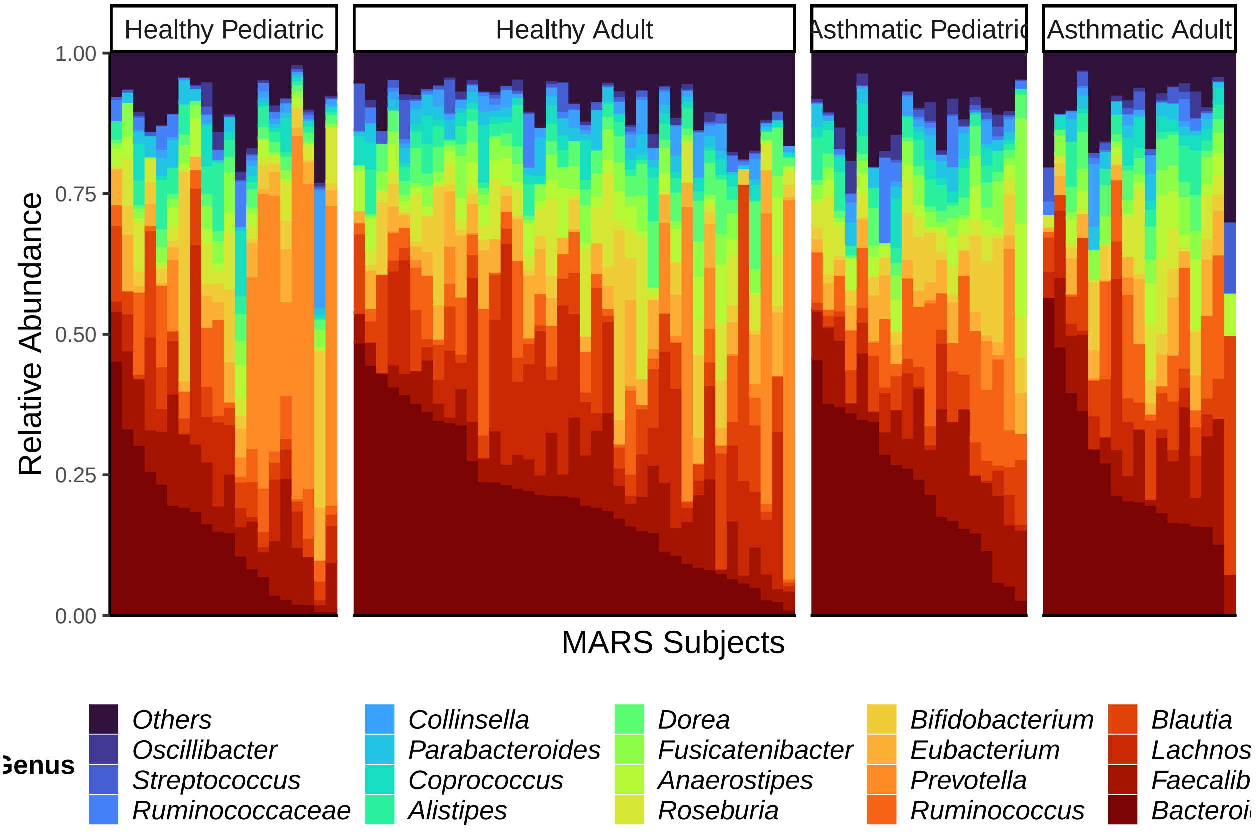


```
# ggsave(filename =
↪  "metaphlan3_topGeneraOrderedBarPlotMARSMetaphlan3_byAgeAndAsthma.jpg", plot =
↪  ageasthmaTrampStamp, device = "jpg", height = 10, width = 20, units = "cm")
# ggsave(filename =
↪  "metaphlan3_topGeneraOrderedBarPlotMARSMetaphlan3_byAgeAndAsthma.pdf", plot =
↪  ageasthmaTrampStamp, device = "pdf", height = 10, width = 20, units = "cm",
↪  useDingbats=F)
```

#### Figure S1E:

plot metaphlan alpha diversity

```
humann3_mf_filtered <- readRDS("./humann3_mf_filtered.rds")
row.names(humann3_mf_filtered) <- humann3_mf_filtered$SampleName
allSpecies <- readRDS("allSpecies_95_relab_tab.rds")
# Naomi try to plot metaphlan alpha diversity
allSpeciesBinary = allSpecies
allSpeciesBinary[allSpeciesBinary > 0] = 1
# allSpecies1000 = allSpeciesBinary*1000
obsSpecies = as.data.frame(colSums(allSpeciesBinary))
obsSpecies$Asthma = humann3_mf_filtered[gsub(row.names(obsSpecies), pattern="[.]",
↪  replacement="-"), "ASTHMA"]
names(obsSpecies) = c("observed_species", "asthma")
obsSpecies$asthma <- factor(obsSpecies$asthma, levels = c("Healthy", "Asthmatic"))
obsSpecies$simpson <- diversity(t(allSpecies), "simpson")
# obsSpecies$shannon <- diversity(t(allSpecies), "shannon")
# boxplot(observed_species~asthma, data = obsSpecies)

library(RColorBrewer)
# obsSpeciesAlphaDivPlot <- makeMARSPlot(datframe = obsSpecies, y_title="Number of
↪  Species Observed", y_column = "observed_species", x_column = "asthma", expt_column =
↪  "asthma", microbiome_column = "asthma", shape_int = 16)
# obsSpeciesAlphaDivPlot
# ggsave(plot = obsSpeciesAlphaDivPlot, filename = "obsSpeciesAlphaDivPlot.pdf",device =
↪  "pdf", width = 5, height = 5, units = "in", useDingbats=FALSE)
# ggsave(plot = obsSpeciesAlphaDivPlot, filename = "obsSpeciesAlphaDivPlot.jpeg",device =
↪  "jpg", width = 5, height = 5, units = "in")

# shannonAlphaDivPlot <- makeMARSPlot(datframe = obsSpecies, y_title="Shannon Diversity",
↪  y_column = "shannon", x_column = "asthma", expt_column = "asthma", microbiome_column
↪  = "asthma", shape_int = 16)
# ggsave(plot = shannonAlphaDivPlot, filename = "shannonAlphaDivPlot.pdf",device = "pdf",
↪  width = 5, height = 5, units = "in", useDingbats=FALSE)

# simpsonAlphaDivPlot <- makeMARSPlot(datframe = obsSpecies, y_title="Simpson Diversity",
↪  y_column = "simpson", x_column = "asthma", expt_column = "asthma", microbiome_column
↪  = "asthma", shape_int = 16)
# ggsave(plot = simpsonAlphaDivPlot, filename = "simpsonAlphaDivPlot.pdf",device = "pdf",
↪  width = 5, height = 5, units = "in", useDingbats=FALSE)

# print(obsSpeciesAlphaDivPlot)
# print(shannonAlphaDivPlot)
# print(simpsonAlphaDivPlot)

# include read depth as a covariate
# readDepth <- read.table("~/Library/CloudStorage/Box-Box/Kau
↪  Lab/Results/MARS/FecalMetagenomics/kneaddata/kneaddata_bypassTRF_on_raw_data/merged_kneaddata_table
↪  header = T, sep="\t", stringsAsFactors=F) # added Dec 11 2022 for normalizing alpha
↪  diversity by read depth
# row.names(readDepth) <- gsub(readDepth$Sample, pattern = "_L002_R1_001_kneaddata",
↪  replacement = "")
```

```r
# length(unique(gsub(humanSamples$File1, pattern="_L002_R[1|2]_001.fastq.gz", replacement
↪  = "")))
obsSpecies$readDepth <- humann3_mf_filtered[gsub(row.names(obsSpecies), pattern="[.]",
↪  replacement = "-"), "readDepth"]
obsSpecies$AgeGroup <- humann3_mf_filtered[gsub(row.names(obsSpecies), pattern="[.]",
↪  replacement = "-"), "AgeGroup"]
# summary(aov(simpson~asthma*AgeGroup+asthma*readDepth, data=obsSpecies))
# summary(aov(observed_species~asthma*AgeGroup+asthma*readDepth, data=obsSpecies))
# summary(aov(shannon~asthma*AgeGroup+asthma*readDepth, data=obsSpecies))
# car::Anova(lm(shannon~asthma*readDepth+asthma*AgeGroup, data=obsSpecies), type=2)
car::Anova(lm(simpson~readDepth+asthma*AgeGroup, data=obsSpecies), type=2)
```

```
## Anova Table (Type II tests)
##
## Response: simpson
##                   Sum Sq Df F value  Pr(>F)
## readDepth        0.00293  1  0.6441 0.42434
## asthma           0.02996  1  6.5919 0.01189 *
## AgeGroup         0.00471  1  1.0356 0.31157
## asthma:AgeGroup  0.00335  1  0.7365 0.39307
## Residuals        0.40901 90
## ---
## Signif. codes:  0 '***' 0.001 '**' 0.01 '*' 0.05 '.' 0.1 ' ' 1
```

```r
# car::Anova(lm(observed_species~readDepth+asthma*AgeGroup, data=obsSpecies), type=2)
# wilcox.test(readDepth~asthma, data=obsSpecies)

# 4 groups
simpsonAlphaDivPlot <- makeMARSPlot(datframe = obsSpecies, y_title="Simpson Diversity",
↪  y_column = "simpson", x_column = "asthma", expt_column = "asthma", microbiome_column
↪  = "asthma", shape_int = 16)
obsSpecies$cohort <- factor(paste0(obsSpecies$asthma, " ", obsSpecies$AgeGroup),
↪  levels=c("Healthy Pediatric", "Healthy Adult", "Asthmatic Pediatric", "Asthmatic
↪  Adult"), labels= c("Healthy Pediatric", "Healthy Adult", "Asthmatic Pediatric",
↪  "Asthmatic Adult"))

psimp <- ggplot(obsSpecies, aes(x=cohort, y=simpson)) +
    geom_boxplot(outlier.shape=NA) +
  ggbeeswarm::geom_quasirandom(aes(shape=cohort, fill=cohort), size=3.5, alpha=0.9)+
    # geom_point(position=position_beeswarm(cex=4), aes(shape=cohort,  fill=cohort),
    ↪  size=3.5) +
    scale_fill_manual(values = c(brewer.pal(7, "BrBG")[c(3,5,1,7)]))+
                      # values=c("#DE4968FF", "#51127CFF"))+# "#DE4968FF", "#51127CFF",
                      ↪  "#51127CFF")) +
    scale_shape_manual(values=c(22, 22, 21, 21)) +
    scale_color_manual(values=c("black", "black","black", "black")) +
    scale_x_discrete(name = "Microbiota") +
    scale_y_continuous(name="Simpson Alpha Diversity") +
    theme(panel.grid.major = element_blank(),
          panel.grid.minor = element_blank(),
          axis.line.x = element_line(size = 0, colour = "black"),
          axis.line.y = element_line(size = 0, colour = "black"),
          axis.line = element_line(size=1, colour = "black"),
          panel.border = element_rect(fill = NA, colour = "black", size = 1),
```

```
            panel.background = element_blank(),
            text=element_text(size = 8),
            axis.text.x=element_text(colour="black", size = 8),
            axis.text.y=element_text(colour="black", size = 8),
            axis.title.x=element_blank(),
            legend.title=element_blank(),
            legend.key = element_rect(color=NA, fill=NA)
    )
psimp
```

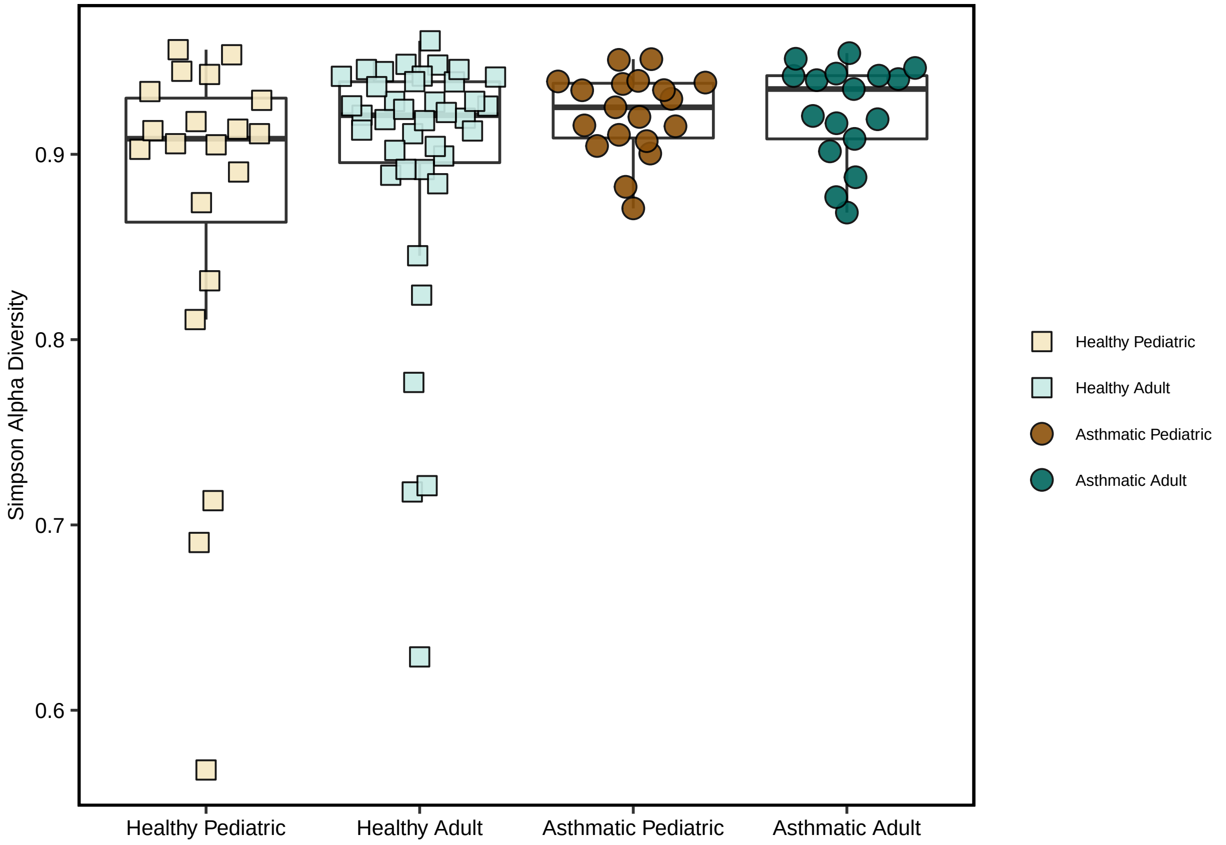


```
# ggsave(plot = psimp, filename = "simpsonAlphaDivPlot_AgeAsthmaSplit.pdf",device =
↪  "pdf", width = 5.5, height = 4, units = "in", useDingbats=FALSE)
```

### Figure S1F:

Plot Beta Diversity on species level

```
basedir="./"
humann3_mf_filtered_perm <- readRDS(file = paste0(basedir, "humann3_mf_filtered.rds"))
row.names(humann3_mf_filtered_perm) <- humann3_mf_filtered_perm$SampleName
df.meta <- humann3_mf_filtered_perm
# filter out absent species
sampleNames <- gsub(rownames(df.meta), pattern = "[-]", replacement=".")
asv.clean <- allSpecies[,sampleNames]
asv.clean <- t(asv.clean[rowSums(asv.clean)>0,])
```

```r
# NMDS
brayDiv.msds <- metaMDS(asv.clean, distance = "bray", k = 6, trymax = 100) # 0.09549306
```

```
## Run 0 stress 0.09731988
## Run 1 stress 0.09704167
## ... New best solution
## ... Procrustes: rmse 0.01137672  max resid 0.03443683
## Run 2 stress 0.09556318
## ... New best solution
## ... Procrustes: rmse 0.03863043  max resid 0.1811506
## Run 3 stress 0.09724935
## Run 4 stress 0.09775868
## Run 5 stress 0.09655825
## Run 6 stress 0.09553775
## ... New best solution
## ... Procrustes: rmse 0.003348857  max resid 0.01380841
## Run 7 stress 0.09826559
## Run 8 stress 0.09658767
## Run 9 stress 0.09549874
## ... New best solution
## ... Procrustes: rmse 0.004765412  max resid 0.01962335
## Run 10 stress 0.09566222
## ... Procrustes: rmse 0.005130193  max resid 0.02408612
## Run 11 stress 0.09551763
## ... Procrustes: rmse 0.004799967  max resid 0.01319955
## Run 12 stress 0.09549076
## ... New best solution
## ... Procrustes: rmse 0.002020785  max resid 0.008851071
## ... Similar to previous best
## Run 13 stress 0.09557864
## ... Procrustes: rmse 0.005598881  max resid 0.01912742
## Run 14 stress 0.09557937
## ... Procrustes: rmse 0.004965896  max resid 0.02109199
## Run 15 stress 0.09556539
## ... Procrustes: rmse 0.005334548  max resid 0.02146025
## Run 16 stress 0.09551874
## ... Procrustes: rmse 0.00322114  max resid 0.01031611
## Run 17 stress 0.09550711
## ... Procrustes: rmse 0.002462385  max resid 0.009720045
## ... Similar to previous best
## Run 18 stress 0.09552262
## ... Procrustes: rmse 0.002965329  max resid 0.01000008
## Run 19 stress 0.09721356
## Run 20 stress 0.09663139
## *** Solution reached
```

```r
saveRDS(object = brayDiv.msds, file = "allSpeciesRelAbBrayCurtis.msds")

# make asthma v healthy NMDS plot
row.names(df.meta) <- gsub(row.names(df.meta), pattern = "-", replacement = ".")
# makeNMDSplot(data.msds = brayDiv.msds, mymetadata = df.meta, confidenceEllipse = 0.95,
↪  my_data_table_name = "metaphlan_Species", basedir = "./")

####
```

```
# 4 grouper (Age:Asthma####
basedir="./"
my_data_table_name = "metaphlan_species"
mytitle = "Bray Curtis Dissimilarity of species relative abundance"
confidenceEllipse = 0.95
brayCurtisMDS <- data.frame(brayDiv.msds$points)
brayCurtisMDS$AgeGroup <-
↪  factor(df.meta[as.character(row.names(brayCurtisMDS)),"AgeGroup"])
brayCurtisMDS$Asthma <- factor(df.meta[as.character(row.names(brayCurtisMDS)),"ASTHMA"])
brayCurtisMDS$Cohort <- factor(paste0(brayCurtisMDS$Asthma, brayCurtisMDS$AgeGroup))
my_color_scheme = c(brewer.pal(7, "BrBG")[c(7,1,5,3)])
# library(viridis)
betaDivPlot4 <- ggplot(data = brayCurtisMDS,
                        aes(x = MDS1, y = MDS2,
                            fill = Cohort,
                            color = Cohort,
                            shape = Asthma)) +
  stat_ellipse(aes(x=MDS1, y=MDS2, color=Cohort, group=Cohort), show.legend = TRUE, type
  ↪  = "t", level = confidenceEllipse, geom = "polygon", alpha = 0.2, size = 1)+
  # geom_point(size = 6, shape = 21, color ="black") +
  geom_point(size = 4.5, color = "black") +
  scale_color_manual(values = my_color_scheme) +
  scale_fill_manual(values = my_color_scheme) +
  scale_shape_manual(values=c(21, 22)) +
  theme_classic() +
  xlab(label = "NMDS1") + ylab("NMDS2") + ggtitle(mytitle) +
  theme(axis.title.x = element_text(size = 21, vjust = -0.9),
        axis.title.y = element_text(size = 21, vjust = 1),
        axis.text.x = element_blank(),
        axis.ticks = element_blank(),
        axis.text.y = element_blank(),
        legend.text = element_text(size = 16),
        legend.title = element_text(size = 16, face = "bold"),
        legend.position = "bottom", legend.box = "horizontal")#,
betaDivPlot4
```

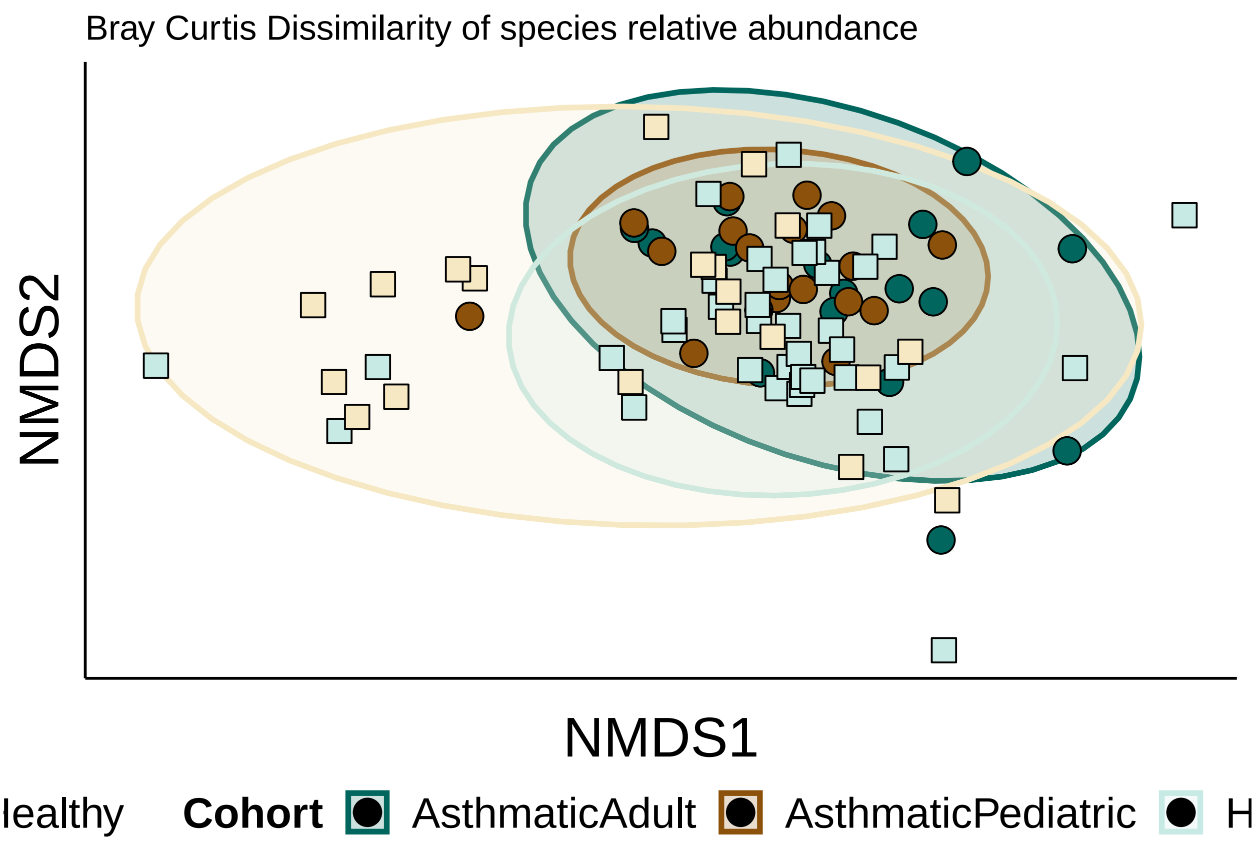


```
# ggsave(plot=betaDivPlot4, filename = paste0(basedir, my_data_table_name,
↪   "_4Group_NMDS_1and2.jpeg"), device = "jpg", width = 5, height = 5, units = "in")
# ggsave(plot=betaDivPlot4, filename = paste0(basedir, my_data_table_name,
↪   "_4Group_NMDS_1and2.pdf"), device = "pdf", width = 5, height = 5, units = "in",
↪   useDingbats = FALSE)
```

##### Figure S1G:

PERMANOVA Species Bray-Curtis distances barplot

```
rm(list=ls())
source("./humann3_210613_functions_ngw.R")

library(ggplot2)
library(ape)
library(stats)
library(vegan)
library(ggfortify)
library(RColorBrewer)

allSpecies <- readRDS("allSpecies_95_relab_tab.rds")

gene_analysis_name = "metaphlan_species"
basedir_humann3="./"
humann3_mf_filtered_perm <- readRDS(file = paste0(basedir_humann3,
↪   "humann3_mf_filtered.rds"))
row.names(humann3_mf_filtered_perm) <- humann3_mf_filtered_perm$SampleName
```

```r
df.meta <- humann3_mf_filtered_perm
df.meta$readDepth <- df.meta[row.names(df.meta), "readDepth"]

sampleNames <- gsub(rownames(df.meta), pattern = "[-]", replacement=".")
asv.clean <- allSpecies[,sampleNames]
asv.clean <- t(asv.clean[rowSums(asv.clean)>0,])

##ADONIS
#On Bray-Curtis
brayDiv <- vegdist(asv.clean, method="bray", binary=FALSE, diag=FALSE, upper=FALSE, na.rm
↪  = FALSE)

#Permanova on Distances
adonisAllIndpRaceBinaryPATHWAYS <- adonis(brayDiv ~
↪  readDepth+ASTHMA*AgeGroup+ASTHMA*AbxWithinYr+raceBinary+weightClassBinary+sex+subjectTobaccoUse,
↪  data = df.meta, permutations = 100000)

# make table
adonisAllIndpRaceBinaryPATHWAYSTable <-
↪  data.frame(adonisAllIndpRaceBinaryPATHWAYS$aov.tab[c("R2","Pr(>F)")])
names(adonisAllIndpRaceBinaryPATHWAYSTable) <- c("R2", "pval")
adonisAllIndpRaceBinaryPATHWAYSTable$features <-
↪  row.names(adonisAllIndpRaceBinaryPATHWAYSTable)
adonisAllIndpRaceBinaryPATHWAYSTable
↪  <-adonisAllIndpRaceBinaryPATHWAYSTable[!adonisAllIndpRaceBinaryPATHWAYSTable$features
↪  %in% c("Residuals","Total"),]
adonisAllIndpRaceBinaryPATHWAYSTable$pval <-
↪  round(adonisAllIndpRaceBinaryPATHWAYSTable$pval, 5)
adonisAllIndpRaceBinaryPATHWAYSTable$R2 <- round(adonisAllIndpRaceBinaryPATHWAYSTable$R2,
↪  3)
row.names(adonisAllIndpRaceBinaryPATHWAYSTable) <- NULL
adonisAllIndpRaceBinaryPATHWAYSTable <- adonisAllIndpRaceBinaryPATHWAYSTable[,c(3,1,2)]

basedir="./"
write.table(adonisAllIndpRaceBinaryPATHWAYSTable, file = paste0(basedir,
↪  gene_analysis_name, "_adonisAllIndpRaceBinaryweightClassBinary10000.tsv"), sep="\t",
↪  row.names = FALSE, col.names = TRUE)
adonisAllIndpRaceBinaryPATHWAYSTable$neglogpval <-
↪  -log(adonisAllIndpRaceBinaryPATHWAYSTable$pval, base = 10)
# adonisAllIndpRaceBinaryPATHWAYSTable$features <-
↪  reorder(adonisAllIndpRaceBinaryPATHWAYSTable$features,
↪  adonisAllIndpRaceBinaryPATHWAYSTable$R2)
adonisAllIndpRaceBinaryPATHWAYSTable$features <-
↪  factor(adonisAllIndpRaceBinaryPATHWAYSTable$features, levels=rev(c("readDepth",
↪  "ASTHMA", "AgeGroup", "ASTHMA:AgeGroup", "AbxWithinYr", "ASTHMA:AbxWithinYr",
↪  "raceBinary", "weightClassBinary",  "sex", "subjectTobaccoUse")), labels =
↪  rev(c("Read Depth", "Asthma", "Age", "Asthma:Age", "Recent Antibiotics",
↪  "Asthma:Recent\nAntibiotics", "Race","Obesity", "Sex", "Tobacco Use")))
AdonisBarplotForPaperBinaryRacePATHWAYS <- makeAdonisBarplot(adonis_dat =
↪  adonisAllIndpRaceBinaryPATHWAYSTable, y_colname = "neglogpval", features_colname =
↪  "features", fill_colname = "R2", max_fill_value =
↪  max(adonisAllIndpRaceBinaryPATHWAYSTable$R2)*1.1)
print(AdonisBarplotForPaperBinaryRacePATHWAYS)
```

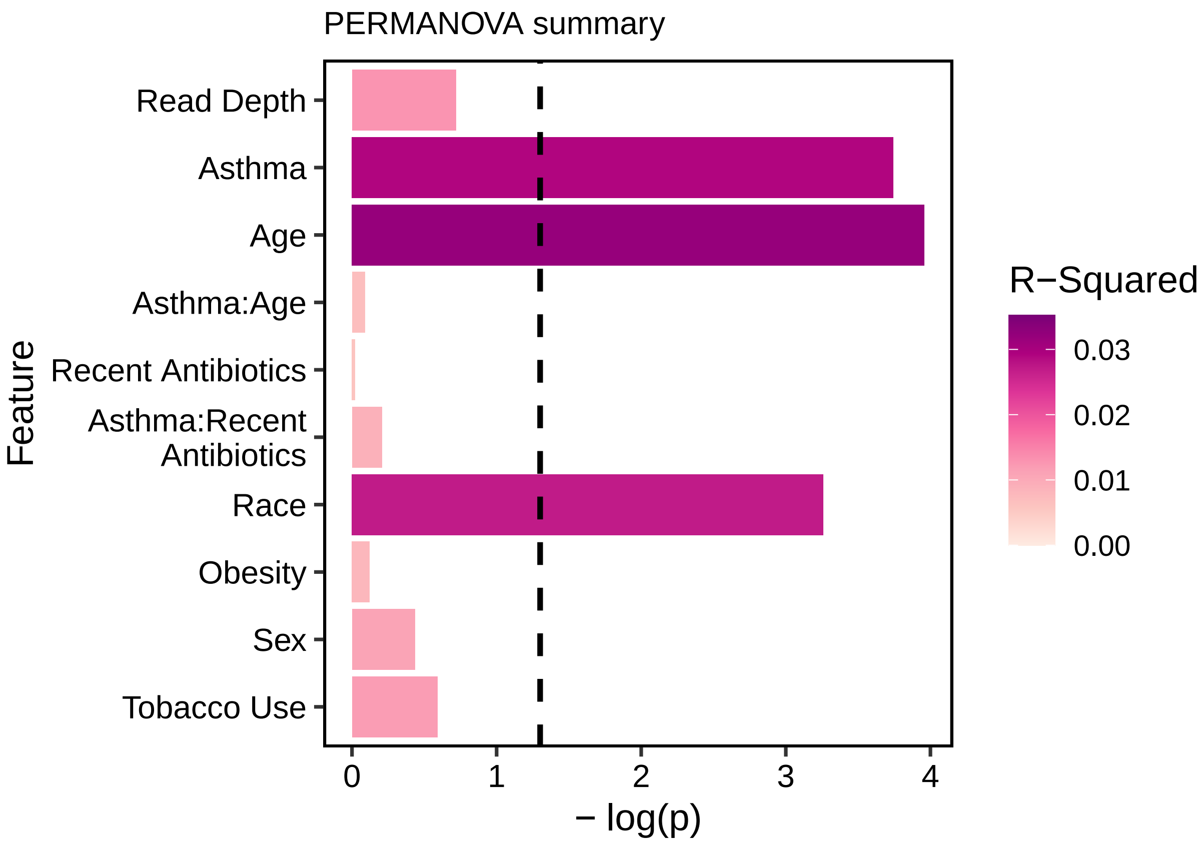


```
# ggsave(AdonisBarplotForPaperBinaryRacePATHWAYS,
#        filename = paste0(basedir, gene_analysis_name,
↪   "_adonisAllIndpRaceBinaryweightClassBinary100000.pdf"),
#        height = 4, width = 5,
#        units = "in",
#        device = "pdf", useDingbats=FALSE)
# ggsave(AdonisBarplotForPaperBinaryRacePATHWAYS,
#        filename = paste0(basedir, gene_analysis_name,
↪   "_adonisAllIndpRaceBinaryweightClassBinary100000.jpg"),
#        height = 4, width = 5,
#        units = "in",
#        device = "jpg")
```

#### Figure S1H:

Make the PCL file rows are the data, columns are the sample, file must be tab-delimited, sample names are first data, followed by rest of relevant metadata, followed by otus/asvs.

```
rm(list=ls())
options(stringsAsFactors=F)
allTaxa <- readRDS("allTaxa_95_relab_tab.rds")

humanSamples <- readRDS("./humann3_mf_filtered.rds")
maaslinMetaData <- t(humanSamples)
humanSamples$SampleNamePeriod = gsub(pattern = "-", replacement = ".",
↪  humanSamples$SampleName)
```

```
row.names(humanSamples) <- humanSamples$MARSID
metadataSamples <- humanSamples[, c("SampleNamePeriod")]
row.names(allTaxa) = allTaxa$clade_name
maaslinASVs <- allTaxa[,metadataSamples]
maaslinASVs <- apply(X = maaslinASVs, MARGIN = 2, FUN = function(x) {x/sum(x)})
maaslinPCSFile <- rbind(maaslinMetaData, maaslinASVs)
write.table(x = maaslinPCSFile, file = "maaslinPCSFile95.txt", append = FALSE, sep =
↪  "\t")
```

run maaslin for metaphlan output (random=age and race; fixed=asthma)

```
rm(list=ls())
options(stringsAsFactors=F)
maaslinPCSFile <- read.csv(file = "maaslinPCSFile95.txt", header=TRUE, sep="\t", as.is =
↪  T)

metadata_categories = rownames(maaslinPCSFile)[1:13]
input_metadata = data.frame((t(maaslinPCSFile))[,1:13])
input_metadata$Population = as.numeric(factor(input_metadata$AgeGroup))
input_metadata$raceBinary = as.numeric(as.factor(input_metadata$raceBinary))
input_metadata$weightClass = as.numeric(factor(input_metadata$weightClass))
input_metadata$AbxWithinYr = as.numeric(as.factor(input_metadata$AbxWithinYr))
input_metadata$gender = ifelse(input_metadata$sex == "Female", 1, 0)

input_data_strings = data.frame(t(maaslinPCSFile)[,14:dim(maaslinPCSFile)[1]])
input_data_numeric = mapply(as.numeric, input_data_strings)
rownames(input_data_numeric) = rownames(input_data_strings)

library(Maaslin2) # Used the dev version 1.5.1  Maaslin2_1.5.1
utils::capture.output(fit_data <- Maaslin2(
    input_data_numeric,
    input_metadata,
    output='maaslin_fixed_asthma_with_adonis_randomeffects',
    transform = "AST",
    fixed_effects = c('ASTHMA'),
    random_effects = c('Population', 'raceBinary'), #
    ↪  metadata_categories[3:length(metadata_categories)]
    normalization = 'NONE',
    standardize = FALSE))
```

```
## boundary (singular) fit: see help('isSingular')
## boundary (singular) fit: see help('isSingular')
## boundary (singular) fit: see help('isSingular')
## boundary (singular) fit: see help('isSingular')
## boundary (singular) fit: see help('isSingular')
## boundary (singular) fit: see help('isSingular')
## boundary (singular) fit: see help('isSingular')
## boundary (singular) fit: see help('isSingular')
## boundary (singular) fit: see help('isSingular')
## boundary (singular) fit: see help('isSingular')
## boundary (singular) fit: see help('isSingular')
## boundary (singular) fit: see help('isSingular')
## boundary (singular) fit: see help('isSingular')
## boundary (singular) fit: see help('isSingular')
```

```
## boundary (singular) fit: see help('isSingular')
## boundary (singular) fit: see help('isSingular')
## boundary (singular) fit: see help('isSingular')
## boundary (singular) fit: see help('isSingular')
## boundary (singular) fit: see help('isSingular')
## boundary (singular) fit: see help('isSingular')
## boundary (singular) fit: see help('isSingular')
## boundary (singular) fit: see help('isSingular')
## boundary (singular) fit: see help('isSingular')
## boundary (singular) fit: see help('isSingular')
## boundary (singular) fit: see help('isSingular')
## boundary (singular) fit: see help('isSingular')
## boundary (singular) fit: see help('isSingular')
## boundary (singular) fit: see help('isSingular')
## boundary (singular) fit: see help('isSingular')
## boundary (singular) fit: see help('isSingular')
## boundary (singular) fit: see help('isSingular')
## boundary (singular) fit: see help('isSingular')
## boundary (singular) fit: see help('isSingular')
## boundary (singular) fit: see help('isSingular')
## boundary (singular) fit: see help('isSingular')
## boundary (singular) fit: see help('isSingular')
## boundary (singular) fit: see help('isSingular')
## boundary (singular) fit: see help('isSingular')
## boundary (singular) fit: see help('isSingular')
## boundary (singular) fit: see help('isSingular')
## boundary (singular) fit: see help('isSingular')
## boundary (singular) fit: see help('isSingular')
## boundary (singular) fit: see help('isSingular')
## boundary (singular) fit: see help('isSingular')
## boundary (singular) fit: see help('isSingular')
## boundary (singular) fit: see help('isSingular')
## boundary (singular) fit: see help('isSingular')
## boundary (singular) fit: see help('isSingular')
## boundary (singular) fit: see help('isSingular')
## boundary (singular) fit: see help('isSingular')
## boundary (singular) fit: see help('isSingular')
## boundary (singular) fit: see help('isSingular')
## boundary (singular) fit: see help('isSingular')
## boundary (singular) fit: see help('isSingular')
## boundary (singular) fit: see help('isSingular')
## boundary (singular) fit: see help('isSingular')
## boundary (singular) fit: see help('isSingular')
## boundary (singular) fit: see help('isSingular')
## boundary (singular) fit: see help('isSingular')
## boundary (singular) fit: see help('isSingular')
## boundary (singular) fit: see help('isSingular')
## boundary (singular) fit: see help('isSingular')

## Warning in checkConv(attr(opt, "derivs"), opt$par, ctrl = control$checkConv, :
## Model failed to converge with max|grad| = 0.0232542 (tol = 0.002, component 1)

## boundary (singular) fit: see help('isSingular')
## boundary (singular) fit: see help('isSingular')
## boundary (singular) fit: see help('isSingular')
```

```
## boundary (singular) fit: see help('isSingular')
## boundary (singular) fit: see help('isSingular')
## boundary (singular) fit: see help('isSingular')
## boundary (singular) fit: see help('isSingular')
## boundary (singular) fit: see help('isSingular')
## boundary (singular) fit: see help('isSingular')
## boundary (singular) fit: see help('isSingular')
## boundary (singular) fit: see help('isSingular')
## boundary (singular) fit: see help('isSingular')
## boundary (singular) fit: see help('isSingular')
## boundary (singular) fit: see help('isSingular')
## boundary (singular) fit: see help('isSingular')
## boundary (singular) fit: see help('isSingular')
## boundary (singular) fit: see help('isSingular')
## boundary (singular) fit: see help('isSingular')
## boundary (singular) fit: see help('isSingular')
## boundary (singular) fit: see help('isSingular')
## boundary (singular) fit: see help('isSingular')
## boundary (singular) fit: see help('isSingular')
## boundary (singular) fit: see help('isSingular')
## boundary (singular) fit: see help('isSingular')
## boundary (singular) fit: see help('isSingular')
## boundary (singular) fit: see help('isSingular')
## boundary (singular) fit: see help('isSingular')
## boundary (singular) fit: see help('isSingular')
## boundary (singular) fit: see help('isSingular')
## boundary (singular) fit: see help('isSingular')
## boundary (singular) fit: see help('isSingular')
## boundary (singular) fit: see help('isSingular')
## boundary (singular) fit: see help('isSingular')
## boundary (singular) fit: see help('isSingular')
## boundary (singular) fit: see help('isSingular')
## boundary (singular) fit: see help('isSingular')
## boundary (singular) fit: see help('isSingular')
## boundary (singular) fit: see help('isSingular')
## boundary (singular) fit: see help('isSingular')
## boundary (singular) fit: see help('isSingular')
## boundary (singular) fit: see help('isSingular')
## boundary (singular) fit: see help('isSingular')
## boundary (singular) fit: see help('isSingular')
## boundary (singular) fit: see help('isSingular')
## boundary (singular) fit: see help('isSingular')
## boundary (singular) fit: see help('isSingular')
## boundary (singular) fit: see help('isSingular')
## boundary (singular) fit: see help('isSingular')
## boundary (singular) fit: see help('isSingular')
## boundary (singular) fit: see help('isSingular')
## boundary (singular) fit: see help('isSingular')
## boundary (singular) fit: see help('isSingular')
## boundary (singular) fit: see help('isSingular')
## boundary (singular) fit: see help('isSingular')
## boundary (singular) fit: see help('isSingular')
## boundary (singular) fit: see help('isSingular')
## boundary (singular) fit: see help('isSingular')
```

```
## boundary (singular) fit: see help('isSingular')
## boundary (singular) fit: see help('isSingular')
## boundary (singular) fit: see help('isSingular')
## boundary (singular) fit: see help('isSingular')
## boundary (singular) fit: see help('isSingular')
## boundary (singular) fit: see help('isSingular')
## boundary (singular) fit: see help('isSingular')
## boundary (singular) fit: see help('isSingular')
## boundary (singular) fit: see help('isSingular')
## boundary (singular) fit: see help('isSingular')
## boundary (singular) fit: see help('isSingular')
## boundary (singular) fit: see help('isSingular')
## boundary (singular) fit: see help('isSingular')
## boundary (singular) fit: see help('isSingular')
## boundary (singular) fit: see help('isSingular')
## boundary (singular) fit: see help('isSingular')
## boundary (singular) fit: see help('isSingular')
## boundary (singular) fit: see help('isSingular')
## boundary (singular) fit: see help('isSingular')
## boundary (singular) fit: see help('isSingular')
## boundary (singular) fit: see help('isSingular')
## boundary (singular) fit: see help('isSingular')
## boundary (singular) fit: see help('isSingular')
## boundary (singular) fit: see help('isSingular')
## boundary (singular) fit: see help('isSingular')
## boundary (singular) fit: see help('isSingular')
## boundary (singular) fit: see help('isSingular')
## boundary (singular) fit: see help('isSingular')
## boundary (singular) fit: see help('isSingular')
## boundary (singular) fit: see help('isSingular')
## boundary (singular) fit: see help('isSingular')
## boundary (singular) fit: see help('isSingular')
## boundary (singular) fit: see help('isSingular')
## boundary (singular) fit: see help('isSingular')
## boundary (singular) fit: see help('isSingular')
## boundary (singular) fit: see help('isSingular')
## boundary (singular) fit: see help('isSingular')
## boundary (singular) fit: see help('isSingular')
## boundary (singular) fit: see help('isSingular')
## boundary (singular) fit: see help('isSingular')
## boundary (singular) fit: see help('isSingular')
## boundary (singular) fit: see help('isSingular')
## boundary (singular) fit: see help('isSingular')
## boundary (singular) fit: see help('isSingular')
## boundary (singular) fit: see help('isSingular')
## boundary (singular) fit: see help('isSingular')
## boundary (singular) fit: see help('isSingular')
## boundary (singular) fit: see help('isSingular')
## boundary (singular) fit: see help('isSingular')
## boundary (singular) fit: see help('isSingular')
## boundary (singular) fit: see help('isSingular')
## boundary (singular) fit: see help('isSingular')
## boundary (singular) fit: see help('isSingular')
## boundary (singular) fit: see help('isSingular')
```

```
## boundary (singular) fit: see help('isSingular')
## boundary (singular) fit: see help('isSingular')
## boundary (singular) fit: see help('isSingular')
## boundary (singular) fit: see help('isSingular')
## boundary (singular) fit: see help('isSingular')
## boundary (singular) fit: see help('isSingular')
## boundary (singular) fit: see help('isSingular')
## boundary (singular) fit: see help('isSingular')
## boundary (singular) fit: see help('isSingular')
## boundary (singular) fit: see help('isSingular')
## boundary (singular) fit: see help('isSingular')
## boundary (singular) fit: see help('isSingular')
## boundary (singular) fit: see help('isSingular')
## boundary (singular) fit: see help('isSingular')
## boundary (singular) fit: see help('isSingular')
## boundary (singular) fit: see help('isSingular')
## boundary (singular) fit: see help('isSingular')
## boundary (singular) fit: see help('isSingular')
## boundary (singular) fit: see help('isSingular')
## boundary (singular) fit: see help('isSingular')
## boundary (singular) fit: see help('isSingular')
## boundary (singular) fit: see help('isSingular')
## boundary (singular) fit: see help('isSingular')
## boundary (singular) fit: see help('isSingular')
## boundary (singular) fit: see help('isSingular')
## boundary (singular) fit: see help('isSingular')
## boundary (singular) fit: see help('isSingular')
## boundary (singular) fit: see help('isSingular')
## boundary (singular) fit: see help('isSingular')
## boundary (singular) fit: see help('isSingular')
## boundary (singular) fit: see help('isSingular')
## boundary (singular) fit: see help('isSingular')
## boundary (singular) fit: see help('isSingular')
## boundary (singular) fit: see help('isSingular')
## boundary (singular) fit: see help('isSingular')
## boundary (singular) fit: see help('isSingular')
## boundary (singular) fit: see help('isSingular')
## boundary (singular) fit: see help('isSingular')
## boundary (singular) fit: see help('isSingular')
## boundary (singular) fit: see help('isSingular')
## boundary (singular) fit: see help('isSingular')
## boundary (singular) fit: see help('isSingular')
## boundary (singular) fit: see help('isSingular')
## boundary (singular) fit: see help('isSingular')
## boundary (singular) fit: see help('isSingular')
## boundary (singular) fit: see help('isSingular')
## boundary (singular) fit: see help('isSingular')
## boundary (singular) fit: see help('isSingular')
## boundary (singular) fit: see help('isSingular')
## boundary (singular) fit: see help('isSingular')
```

```
##   [1] "[1] \"Warning: Deleting existing log file: maaslin_fixed_asthma_with_adonis_randomeffects/maas
##   [2] "2023-01-17 09:03:23 INFO::Writing function arguments to log file"
##   [3] "2023-01-17 09:03:23 INFO::Verifying options selected are valid"
```

```
##   [4] "2023-01-17 09:03:23 INFO::Determining format of input files"
##   [5] "2023-01-17 09:03:23 INFO::Input format is data samples as rows and metadata samples as rows"
##   [6] "2023-01-17 09:03:23 INFO::Formula for random effects: expr ~ (1 | Population) + (1 | raceBina
##   [7] "2023-01-17 09:03:23 INFO::Formula for fixed effects: expr ~  ASTHMA"
##   [8] "2023-01-17 09:03:23 INFO::Filter data based on min abundance and min prevalence"
##   [9] "2023-01-17 09:03:23 INFO::Total samples in data: 95"
##  [10] "2023-01-17 09:03:23 INFO::Min samples required with min abundance for a feature not to be fil
##  [11] "2023-01-17 09:03:23 INFO::Total filtered features: 373"
##  [12] "2023-01-17 09:03:23 INFO::Filtered feature names from abundance and prevalence filtering:"
##  [13] "2023-01-17 09:03:23 INFO::Total filtered features with variance filtering: 0"
##  [14] "2023-01-17 09:03:23 INFO::Filtered feature names from variance filtering:"
##  [15] "2023-01-17 09:03:23 INFO::Running selected normalization method: NONE"
##  [16] "2023-01-17 09:03:23 INFO::Bypass z-score application to metadata"
##  [17] "2023-01-17 09:03:23 INFO::Running selected transform method: AST"
##  [18] "2023-01-17 09:03:23 INFO::Running selected analysis method: LM"
##  [19] "2023-01-17 09:03:23 INFO::Fitting model to feature number 1, k__Archaea"
##  [20] "2023-01-17 09:03:24 INFO::Fitting model to feature number 2, k__Archaea.p__Euryarchaeota"
##  [21] "2023-01-17 09:03:24 INFO::Fitting model to feature number 3, k__Archaea.p__Euryarchaeota.c__M
##  [22] "2023-01-17 09:03:24 INFO::Fitting model to feature number 4, k__Archaea.p__Euryarchaeota.c__M
##  [23] "2023-01-17 09:03:24 INFO::Fitting model to feature number 5, k__Archaea.p__Euryarchaeota.c__M
##  [24] "2023-01-17 09:03:24 INFO::Fitting model to feature number 6, k__Archaea.p__Euryarchaeota.c__M
##  [25] "2023-01-17 09:03:24 INFO::Fitting model to feature number 7, k__Archaea.p__Euryarchaeota.c__M
##  [26] "2023-01-17 09:03:24 INFO::Fitting model to feature number 8, k__Bacteria"
##  [27] "2023-01-17 09:03:24 INFO::Fitting model to feature number 9, k__Bacteria.p__Actinobacteria"
##  [28] "2023-01-17 09:03:24 INFO::Fitting model to feature number 10, k__Bacteria.p__Actinobacteria.c
##  [29] "2023-01-17 09:03:24 INFO::Fitting model to feature number 11, k__Bacteria.p__Actinobacteria.c
##  [30] "2023-01-17 09:03:24 INFO::Fitting model to feature number 12, k__Bacteria.p__Actinobacteria.c
##  [31] "2023-01-17 09:03:24 INFO::Fitting model to feature number 13, k__Bacteria.p__Actinobacteria.c
##  [32] "2023-01-17 09:03:24 INFO::Fitting model to feature number 14, k__Bacteria.p__Actinobacteria.c
##  [33] "2023-01-17 09:03:25 INFO::Fitting model to feature number 15, k__Bacteria.p__Actinobacteria.c
##  [34] "2023-01-17 09:03:25 INFO::Fitting model to feature number 16, k__Bacteria.p__Actinobacteria.c
##  [35] "2023-01-17 09:03:25 INFO::Fitting model to feature number 17, k__Bacteria.p__Actinobacteria.c
##  [36] "2023-01-17 09:03:25 INFO::Fitting model to feature number 18, k__Bacteria.p__Actinobacteria.c
##  [37] "2023-01-17 09:03:25 INFO::Fitting model to feature number 19, k__Bacteria.p__Actinobacteria.c
##  [38] "2023-01-17 09:03:25 INFO::Fitting model to feature number 20, k__Bacteria.p__Actinobacteria.c
##  [39] "2023-01-17 09:03:25 INFO::Fitting model to feature number 21, k__Bacteria.p__Actinobacteria.c
##  [40] "2023-01-17 09:03:25 INFO::Fitting model to feature number 22, k__Bacteria.p__Actinobacteria.c
##  [41] "2023-01-17 09:03:25 INFO::Fitting model to feature number 23, k__Bacteria.p__Actinobacteria.c
##  [42] "2023-01-17 09:03:25 INFO::Fitting model to feature number 24, k__Bacteria.p__Actinobacteria.c
##  [43] "2023-01-17 09:03:25 INFO::Fitting model to feature number 25, k__Bacteria.p__Actinobacteria.c
##  [44] "2023-01-17 09:03:25 INFO::Fitting model to feature number 26, k__Bacteria.p__Actinobacteria.c
##  [45] "2023-01-17 09:03:26 INFO::Fitting model to feature number 27, k__Bacteria.p__Actinobacteria.c
##  [46] "2023-01-17 09:03:26 INFO::Fitting model to feature number 28, k__Bacteria.p__Actinobacteria.c
##  [47] "2023-01-17 09:03:26 INFO::Fitting model to feature number 29, k__Bacteria.p__Actinobacteria.c
##  [48] "2023-01-17 09:03:26 INFO::Fitting model to feature number 30, k__Bacteria.p__Actinobacteria.c
##  [49] "2023-01-17 09:03:26 INFO::Fitting model to feature number 31, k__Bacteria.p__Actinobacteria.c
##  [50] "2023-01-17 09:03:26 INFO::Fitting model to feature number 32, k__Bacteria.p__Actinobacteria.c
##  [51] "2023-01-17 09:03:26 INFO::Fitting model to feature number 33, k__Bacteria.p__Actinobacteria.c
##  [52] "2023-01-17 09:03:26 INFO::Fitting model to feature number 34, k__Bacteria.p__Actinobacteria.c
##  [53] "2023-01-17 09:03:26 INFO::Fitting model to feature number 35, k__Bacteria.p__Actinobacteria.c
##  [54] "2023-01-17 09:03:26 INFO::Fitting model to feature number 36, k__Bacteria.p__Actinobacteria.c
##  [55] "2023-01-17 09:03:26 INFO::Fitting model to feature number 37, k__Bacteria.p__Actinobacteria.c
##  [56] "2023-01-17 09:03:26 INFO::Fitting model to feature number 38, k__Bacteria.p__Actinobacteria.c
##  [57] "2023-01-17 09:03:26 INFO::Fitting model to feature number 39, k__Bacteria.p__Actinobacteria.c
```

```
##  [58] "2023-01-17 09:03:27 INFO::Fitting model to feature number 40, k__Bacteria.p__Actinobacteria.c_
##  [59] "2023-01-17 09:03:27 INFO::Fitting model to feature number 41, k__Bacteria.p__Actinobacteria.c_
##  [60] "2023-01-17 09:03:27 INFO::Fitting model to feature number 42, k__Bacteria.p__Actinobacteria.c_
##  [61] "2023-01-17 09:03:27 INFO::Fitting model to feature number 43, k__Bacteria.p__Actinobacteria.c_
##  [62] "2023-01-17 09:03:27 INFO::Fitting model to feature number 44, k__Bacteria.p__Actinobacteria.c_
##  [63] "2023-01-17 09:03:27 INFO::Fitting model to feature number 45, k__Bacteria.p__Actinobacteria.c_
##  [64] "2023-01-17 09:03:27 INFO::Fitting model to feature number 46, k__Bacteria.p__Actinobacteria.c_
##  [65] "2023-01-17 09:03:27 INFO::Fitting model to feature number 47, k__Bacteria.p__Bacteroidetes"
##  [66] "2023-01-17 09:03:27 INFO::Fitting model to feature number 48, k__Bacteria.p__Bacteroidetes.c_
##  [67] "2023-01-17 09:03:27 INFO::Fitting model to feature number 49, k__Bacteria.p__Bacteroidetes.c_
##  [68] "2023-01-17 09:03:28 INFO::Fitting model to feature number 50, k__Bacteria.p__Bacteroidetes.c_
##  [69] "2023-01-17 09:03:28 INFO::Fitting model to feature number 51, k__Bacteria.p__Bacteroidetes.c_
##  [70] "2023-01-17 09:03:28 INFO::Fitting model to feature number 52, k__Bacteria.p__Bacteroidetes.c_
##  [71] "2023-01-17 09:03:28 INFO::Fitting model to feature number 53, k__Bacteria.p__Bacteroidetes.c_
##  [72] "2023-01-17 09:03:28 INFO::Fitting model to feature number 54, k__Bacteria.p__Bacteroidetes.c_
##  [73] "2023-01-17 09:03:28 INFO::Fitting model to feature number 55, k__Bacteria.p__Bacteroidetes.c_
##  [74] "2023-01-17 09:03:28 INFO::Fitting model to feature number 56, k__Bacteria.p__Bacteroidetes.c_
##  [75] "2023-01-17 09:03:28 INFO::Fitting model to feature number 57, k__Bacteria.p__Bacteroidetes.c_
##  [76] "2023-01-17 09:03:28 INFO::Fitting model to feature number 58, k__Bacteria.p__Bacteroidetes.c_
##  [77] "2023-01-17 09:03:28 INFO::Fitting model to feature number 59, k__Bacteria.p__Bacteroidetes.c_
##  [78] "2023-01-17 09:03:28 INFO::Fitting model to feature number 60, k__Bacteria.p__Bacteroidetes.c_
##  [79] "2023-01-17 09:03:28 INFO::Fitting model to feature number 61, k__Bacteria.p__Bacteroidetes.c_
##  [80] "2023-01-17 09:03:29 INFO::Fitting model to feature number 62, k__Bacteria.p__Bacteroidetes.c_
##  [81] "2023-01-17 09:03:29 INFO::Fitting model to feature number 63, k__Bacteria.p__Bacteroidetes.c_
##  [82] "2023-01-17 09:03:29 INFO::Fitting model to feature number 64, k__Bacteria.p__Bacteroidetes.c_
##  [83] "2023-01-17 09:03:29 INFO::Fitting model to feature number 65, k__Bacteria.p__Bacteroidetes.c_
##  [84] "2023-01-17 09:03:29 INFO::Fitting model to feature number 66, k__Bacteria.p__Bacteroidetes.c_
##  [85] "2023-01-17 09:03:29 INFO::Fitting model to feature number 67, k__Bacteria.p__Bacteroidetes.c_
##  [86] "2023-01-17 09:03:29 INFO::Fitting model to feature number 68, k__Bacteria.p__Bacteroidetes.c_
##  [87] "2023-01-17 09:03:29 INFO::Fitting model to feature number 69, k__Bacteria.p__Bacteroidetes.c_
##  [88] "2023-01-17 09:03:29 INFO::Fitting model to feature number 70, k__Bacteria.p__Bacteroidetes.c_
##  [89] "2023-01-17 09:03:29 INFO::Fitting model to feature number 71, k__Bacteria.p__Bacteroidetes.c_
##  [90] "2023-01-17 09:03:29 INFO::Fitting model to feature number 72, k__Bacteria.p__Bacteroidetes.c_
##  [91] "2023-01-17 09:03:29 INFO::Fitting model to feature number 73, k__Bacteria.p__Bacteroidetes.c_
##  [92] "2023-01-17 09:03:30 INFO::Fitting model to feature number 74, k__Bacteria.p__Bacteroidetes.c_
##  [93] "2023-01-17 09:03:30 INFO::Fitting model to feature number 75, k__Bacteria.p__Bacteroidetes.c_
##  [94] "2023-01-17 09:03:30 INFO::Fitting model to feature number 76, k__Bacteria.p__Bacteroidetes.c_
##  [95] "2023-01-17 09:03:30 INFO::Fitting model to feature number 77, k__Bacteria.p__Bacteroidetes.c_
##  [96] "2023-01-17 09:03:30 INFO::Fitting model to feature number 78, k__Bacteria.p__Bacteroidetes.c_
##  [97] "2023-01-17 09:03:30 INFO::Fitting model to feature number 79, k__Bacteria.p__Bacteroidetes.c_
##  [98] "2023-01-17 09:03:30 INFO::Fitting model to feature number 80, k__Bacteria.p__Bacteroidetes.c_
##  [99] "2023-01-17 09:03:30 INFO::Fitting model to feature number 81, k__Bacteria.p__Bacteroidetes.c_
## [100] "2023-01-17 09:03:30 INFO::Fitting model to feature number 82, k__Bacteria.p__Bacteroidetes.c_
## [101] "2023-01-17 09:03:30 INFO::Fitting model to feature number 83, k__Bacteria.p__Bacteroidetes.c_
## [102] "2023-01-17 09:03:30 INFO::Fitting model to feature number 84, k__Bacteria.p__Bacteroidetes.c_
## [103] "2023-01-17 09:03:30 INFO::Fitting model to feature number 85, k__Bacteria.p__Bacteroidetes.c_
## [104] "2023-01-17 09:03:30 INFO::Fitting model to feature number 86, k__Bacteria.p__Bacteroidetes.c_
## [105] "2023-01-17 09:03:31 INFO::Fitting model to feature number 87, k__Bacteria.p__Bacteroidetes.c_
## [106] "2023-01-17 09:03:31 INFO::Fitting model to feature number 88, k__Bacteria.p__Bacteroidetes.c_
## [107] "2023-01-17 09:03:31 INFO::Fitting model to feature number 89, k__Bacteria.p__Bacteroidetes.c_
## [108] "2023-01-17 09:03:31 INFO::Fitting model to feature number 90, k__Bacteria.p__Bacteroidetes.c_
## [109] "2023-01-17 09:03:31 INFO::Fitting model to feature number 91, k__Bacteria.p__Bacteroidetes.c_
## [110] "2023-01-17 09:03:31 INFO::Fitting model to feature number 92, k__Bacteria.p__Bacteroidetes.c_
## [111] "2023-01-17 09:03:31 INFO::Fitting model to feature number 93, k__Bacteria.p__Bacteroidetes.c_
```

```
## [112] "2023-01-17 09:03:31 INFO::Fitting model to feature number 94, k__Bacteria.p__Bacteroidetes.c__
## [113] "2023-01-17 09:03:31 INFO::Fitting model to feature number 95, k__Bacteria.p__Firmicutes"
## [114] "2023-01-17 09:03:31 INFO::Fitting model to feature number 96, k__Bacteria.p__Firmicutes.c__Bac
## [115] "2023-01-17 09:03:31 INFO::Fitting model to feature number 97, k__Bacteria.p__Firmicutes.c__Bac
## [116] "2023-01-17 09:03:31 INFO::Fitting model to feature number 98, k__Bacteria.p__Firmicutes.c__Bac
## [117] "2023-01-17 09:03:32 INFO::Fitting model to feature number 99, k__Bacteria.p__Firmicutes.c__Bac
## [118] "2023-01-17 09:03:32 INFO::Fitting model to feature number 100, k__Bacteria.p__Firmicutes.c__Ba
## [119] "2023-01-17 09:03:32 INFO::Fitting model to feature number 101, k__Bacteria.p__Firmicutes.c__Ba
## [120] "2023-01-17 09:03:32 INFO::Fitting model to feature number 102, k__Bacteria.p__Firmicutes.c__Ba
## [121] "2023-01-17 09:03:32 INFO::Fitting model to feature number 103, k__Bacteria.p__Firmicutes.c__Ba
## [122] "2023-01-17 09:03:32 INFO::Fitting model to feature number 104, k__Bacteria.p__Firmicutes.c__Ba
## [123] "2023-01-17 09:03:32 INFO::Fitting model to feature number 105, k__Bacteria.p__Firmicutes.c__Ba
## [124] "2023-01-17 09:03:32 INFO::Fitting model to feature number 106, k__Bacteria.p__Firmicutes.c__Ba
## [125] "2023-01-17 09:03:32 INFO::Fitting model to feature number 107, k__Bacteria.p__Firmicutes.c__Ba
## [126] "2023-01-17 09:03:32 INFO::Fitting model to feature number 108, k__Bacteria.p__Firmicutes.c__Ba
## [127] "2023-01-17 09:03:32 INFO::Fitting model to feature number 109, k__Bacteria.p__Firmicutes.c__Ba
## [128] "2023-01-17 09:03:32 INFO::Fitting model to feature number 110, k__Bacteria.p__Firmicutes.c__Ba
## [129] "2023-01-17 09:03:32 INFO::Fitting model to feature number 111, k__Bacteria.p__Firmicutes.c__Ba
## [130] "2023-01-17 09:03:33 INFO::Fitting model to feature number 112, k__Bacteria.p__Firmicutes.c__Ba
## [131] "2023-01-17 09:03:33 INFO::Fitting model to feature number 113, k__Bacteria.p__Firmicutes.c__Ba
## [132] "2023-01-17 09:03:33 INFO::Fitting model to feature number 114, k__Bacteria.p__Firmicutes.c__Ba
## [133] "2023-01-17 09:03:33 INFO::Fitting model to feature number 115, k__Bacteria.p__Firmicutes.c__Ba
## [134] "2023-01-17 09:03:33 INFO::Fitting model to feature number 116, k__Bacteria.p__Firmicutes.c__Ba
## [135] "2023-01-17 09:03:33 INFO::Fitting model to feature number 117, k__Bacteria.p__Firmicutes.c__Ba
## [136] "2023-01-17 09:03:33 INFO::Fitting model to feature number 118, k__Bacteria.p__Firmicutes.c__Cl
## [137] "2023-01-17 09:03:33 INFO::Fitting model to feature number 119, k__Bacteria.p__Firmicutes.c__Cl
## [138] "2023-01-17 09:03:33 INFO::Fitting model to feature number 120, k__Bacteria.p__Firmicutes.c__Cl
## [139] "2023-01-17 09:03:33 INFO::Fitting model to feature number 121, k__Bacteria.p__Firmicutes.c__Cl
## [140] "2023-01-17 09:03:33 INFO::Fitting model to feature number 122, k__Bacteria.p__Firmicutes.c__Cl
## [141] "2023-01-17 09:03:33 INFO::Fitting model to feature number 123, k__Bacteria.p__Firmicutes.c__Cl
## [142] "2023-01-17 09:03:34 INFO::Fitting model to feature number 124, k__Bacteria.p__Firmicutes.c__Cl
## [143] "2023-01-17 09:03:34 INFO::Fitting model to feature number 125, k__Bacteria.p__Firmicutes.c__Cl
## [144] "2023-01-17 09:03:34 INFO::Fitting model to feature number 126, k__Bacteria.p__Firmicutes.c__Cl
## [145] "2023-01-17 09:03:34 INFO::Fitting model to feature number 127, k__Bacteria.p__Firmicutes.c__Cl
## [146] "2023-01-17 09:03:34 INFO::Fitting model to feature number 128, k__Bacteria.p__Firmicutes.c__Cl
## [147] "2023-01-17 09:03:34 INFO::Fitting model to feature number 129, k__Bacteria.p__Firmicutes.c__Cl
## [148] "2023-01-17 09:03:34 INFO::Fitting model to feature number 130, k__Bacteria.p__Firmicutes.c__Cl
## [149] "2023-01-17 09:03:34 INFO::Fitting model to feature number 131, k__Bacteria.p__Firmicutes.c__Cl
## [150] "2023-01-17 09:03:34 INFO::Fitting model to feature number 132, k__Bacteria.p__Firmicutes.c__Cl
## [151] "2023-01-17 09:03:34 INFO::Fitting model to feature number 133, k__Bacteria.p__Firmicutes.c__Cl
## [152] "2023-01-17 09:03:34 INFO::Fitting model to feature number 134, k__Bacteria.p__Firmicutes.c__Cl
## [153] "2023-01-17 09:03:34 INFO::Fitting model to feature number 135, k__Bacteria.p__Firmicutes.c__Cl
## [154] "2023-01-17 09:03:34 INFO::Fitting model to feature number 136, k__Bacteria.p__Firmicutes.c__Cl
## [155] "2023-01-17 09:03:35 INFO::Fitting model to feature number 137, k__Bacteria.p__Firmicutes.c__Cl
## [156] "2023-01-17 09:03:35 INFO::Fitting model to feature number 138, k__Bacteria.p__Firmicutes.c__Cl
## [157] "2023-01-17 09:03:35 INFO::Fitting model to feature number 139, k__Bacteria.p__Firmicutes.c__Cl
## [158] "2023-01-17 09:03:35 INFO::Fitting model to feature number 140, k__Bacteria.p__Firmicutes.c__Cl
## [159] "2023-01-17 09:03:35 INFO::Fitting model to feature number 141, k__Bacteria.p__Firmicutes.c__Cl
## [160] "2023-01-17 09:03:35 INFO::Fitting model to feature number 142, k__Bacteria.p__Firmicutes.c__Cl
## [161] "2023-01-17 09:03:35 INFO::Fitting model to feature number 143, k__Bacteria.p__Firmicutes.c__Cl
## [162] "2023-01-17 09:03:35 INFO::Fitting model to feature number 144, k__Bacteria.p__Firmicutes.c__Cl
## [163] "2023-01-17 09:03:35 INFO::Fitting model to feature number 145, k__Bacteria.p__Firmicutes.c__Cl
## [164] "2023-01-17 09:03:35 INFO::Fitting model to feature number 146, k__Bacteria.p__Firmicutes.c__Cl
## [165] "2023-01-17 09:03:35 INFO::Fitting model to feature number 147, k__Bacteria.p__Firmicutes.c__Cl
```

```
## [166] "2023-01-17 09:03:35 INFO::Fitting model to feature number 148, k__Bacteria.p__Firmicutes.c__Cl
## [167] "2023-01-17 09:03:36 INFO::Fitting model to feature number 149, k__Bacteria.p__Firmicutes.c__Cl
## [168] "2023-01-17 09:03:36 INFO::Fitting model to feature number 150, k__Bacteria.p__Firmicutes.c__Cl
## [169] "2023-01-17 09:03:36 INFO::Fitting model to feature number 151, k__Bacteria.p__Firmicutes.c__Cl
## [170] "2023-01-17 09:03:36 INFO::Fitting model to feature number 152, k__Bacteria.p__Firmicutes.c__Cl
## [171] "2023-01-17 09:03:36 INFO::Fitting model to feature number 153, k__Bacteria.p__Firmicutes.c__Cl
## [172] "2023-01-17 09:03:36 INFO::Fitting model to feature number 154, k__Bacteria.p__Firmicutes.c__Cl
## [173] "2023-01-17 09:03:36 INFO::Fitting model to feature number 155, k__Bacteria.p__Firmicutes.c__Cl
## [174] "2023-01-17 09:03:36 INFO::Fitting model to feature number 156, k__Bacteria.p__Firmicutes.c__Cl
## [175] "2023-01-17 09:03:36 INFO::Fitting model to feature number 157, k__Bacteria.p__Firmicutes.c__Cl
## [176] "2023-01-17 09:03:36 INFO::Fitting model to feature number 158, k__Bacteria.p__Firmicutes.c__Cl
## [177] "2023-01-17 09:03:36 INFO::Fitting model to feature number 159, k__Bacteria.p__Firmicutes.c__Cl
## [178] "2023-01-17 09:03:36 INFO::Fitting model to feature number 160, k__Bacteria.p__Firmicutes.c__Cl
## [179] "2023-01-17 09:03:36 INFO::Fitting model to feature number 161, k__Bacteria.p__Firmicutes.c__Cl
## [180] "2023-01-17 09:03:37 INFO::Fitting model to feature number 162, k__Bacteria.p__Firmicutes.c__Cl
## [181] "2023-01-17 09:03:37 INFO::Fitting model to feature number 163, k__Bacteria.p__Firmicutes.c__Cl
## [182] "2023-01-17 09:03:37 INFO::Fitting model to feature number 164, k__Bacteria.p__Firmicutes.c__Cl
## [183] "2023-01-17 09:03:37 INFO::Fitting model to feature number 165, k__Bacteria.p__Firmicutes.c__Cl
## [184] "2023-01-17 09:03:37 INFO::Fitting model to feature number 166, k__Bacteria.p__Firmicutes.c__Cl
## [185] "2023-01-17 09:03:37 INFO::Fitting model to feature number 167, k__Bacteria.p__Firmicutes.c__Cl
## [186] "2023-01-17 09:03:37 INFO::Fitting model to feature number 168, k__Bacteria.p__Firmicutes.c__Cl
## [187] "2023-01-17 09:03:37 INFO::Fitting model to feature number 169, k__Bacteria.p__Firmicutes.c__Cl
## [188] "2023-01-17 09:03:37 INFO::Fitting model to feature number 170, k__Bacteria.p__Firmicutes.c__Cl
## [189] "2023-01-17 09:03:37 INFO::Fitting model to feature number 171, k__Bacteria.p__Firmicutes.c__Cl
## [190] "2023-01-17 09:03:37 INFO::Fitting model to feature number 172, k__Bacteria.p__Firmicutes.c__Cl
## [191] "2023-01-17 09:03:37 INFO::Fitting model to feature number 173, k__Bacteria.p__Firmicutes.c__Cl
## [192] "2023-01-17 09:03:38 INFO::Fitting model to feature number 174, k__Bacteria.p__Firmicutes.c__Cl
## [193] "2023-01-17 09:03:38 INFO::Fitting model to feature number 175, k__Bacteria.p__Firmicutes.c__Cl
## [194] "2023-01-17 09:03:38 INFO::Fitting model to feature number 176, k__Bacteria.p__Firmicutes.c__Cl
## [195] "2023-01-17 09:03:38 INFO::Fitting model to feature number 177, k__Bacteria.p__Firmicutes.c__Cl
## [196] "2023-01-17 09:03:38 INFO::Fitting model to feature number 178, k__Bacteria.p__Firmicutes.c__Cl
## [197] "2023-01-17 09:03:38 INFO::Fitting model to feature number 179, k__Bacteria.p__Firmicutes.c__Cl
## [198] "2023-01-17 09:03:38 INFO::Fitting model to feature number 180, k__Bacteria.p__Firmicutes.c__Cl
## [199] "2023-01-17 09:03:38 INFO::Fitting model to feature number 181, k__Bacteria.p__Firmicutes.c__Cl
## [200] "2023-01-17 09:03:38 INFO::Fitting model to feature number 182, k__Bacteria.p__Firmicutes.c__Cl
## [201] "2023-01-17 09:03:38 INFO::Fitting model to feature number 183, k__Bacteria.p__Firmicutes.c__Cl
## [202] "2023-01-17 09:03:38 INFO::Fitting model to feature number 184, k__Bacteria.p__Firmicutes.c__Cl
## [203] "2023-01-17 09:03:38 INFO::Fitting model to feature number 185, k__Bacteria.p__Firmicutes.c__Cl
## [204] "2023-01-17 09:03:38 INFO::Fitting model to feature number 186, k__Bacteria.p__Firmicutes.c__Cl
## [205] "2023-01-17 09:03:39 INFO::Fitting model to feature number 187, k__Bacteria.p__Firmicutes.c__Cl
## [206] "2023-01-17 09:03:39 INFO::Fitting model to feature number 188, k__Bacteria.p__Firmicutes.c__Cl
## [207] "2023-01-17 09:03:39 INFO::Fitting model to feature number 189, k__Bacteria.p__Firmicutes.c__Cl
## [208] "2023-01-17 09:03:39 INFO::Fitting model to feature number 190, k__Bacteria.p__Firmicutes.c__Cl
## [209] "2023-01-17 09:03:39 INFO::Fitting model to feature number 191, k__Bacteria.p__Firmicutes.c__Cl
## [210] "2023-01-17 09:03:39 INFO::Fitting model to feature number 192, k__Bacteria.p__Firmicutes.c__Cl
## [211] "2023-01-17 09:03:39 INFO::Fitting model to feature number 193, k__Bacteria.p__Firmicutes.c__Cl
## [212] "2023-01-17 09:03:39 INFO::Fitting model to feature number 194, k__Bacteria.p__Firmicutes.c__Cl
## [213] "2023-01-17 09:03:39 INFO::Fitting model to feature number 195, k__Bacteria.p__Firmicutes.c__Cl
## [214] "2023-01-17 09:03:39 INFO::Fitting model to feature number 196, k__Bacteria.p__Firmicutes.c__Cl
## [215] "2023-01-17 09:03:39 INFO::Fitting model to feature number 197, k__Bacteria.p__Firmicutes.c__Cl
## [216] "2023-01-17 09:03:39 INFO::Fitting model to feature number 198, k__Bacteria.p__Firmicutes.c__Cl
## [217] "2023-01-17 09:03:40 INFO::Fitting model to feature number 199, k__Bacteria.p__Firmicutes.c__Cl
## [218] "2023-01-17 09:03:40 INFO::Fitting model to feature number 200, k__Bacteria.p__Firmicutes.c__Cl
## [219] "2023-01-17 09:03:40 INFO::Fitting model to feature number 201, k__Bacteria.p__Firmicutes.c__Cl
```

```
## [220] "2023-01-17 09:03:40 INFO::Fitting model to feature number 202, k__Bacteria.p__Firmicutes.c__Cl
## [221] "2023-01-17 09:03:40 INFO::Fitting model to feature number 203, k__Bacteria.p__Firmicutes.c__Cl
## [222] "2023-01-17 09:03:40 INFO::Fitting model to feature number 204, k__Bacteria.p__Firmicutes.c__Cl
## [223] "2023-01-17 09:03:40 INFO::Fitting model to feature number 205, k__Bacteria.p__Firmicutes.c__Cl
## [224] "2023-01-17 09:03:40 INFO::Fitting model to feature number 206, k__Bacteria.p__Firmicutes.c__Cl
## [225] "2023-01-17 09:03:40 INFO::Fitting model to feature number 207, k__Bacteria.p__Firmicutes.c__Cl
## [226] "2023-01-17 09:03:40 INFO::Fitting model to feature number 208, k__Bacteria.p__Firmicutes.c__Cl
## [227] "2023-01-17 09:03:40 INFO::Fitting model to feature number 209, k__Bacteria.p__Firmicutes.c__Cl
## [228] "2023-01-17 09:03:40 INFO::Fitting model to feature number 210, k__Bacteria.p__Firmicutes.c__Cl
## [229] "2023-01-17 09:03:40 INFO::Fitting model to feature number 211, k__Bacteria.p__Firmicutes.c__Cl
## [230] "2023-01-17 09:03:41 INFO::Fitting model to feature number 212, k__Bacteria.p__Firmicutes.c__Cl
## [231] "2023-01-17 09:03:41 INFO::Fitting model to feature number 213, k__Bacteria.p__Firmicutes.c__Cl
## [232] "2023-01-17 09:03:41 INFO::Fitting model to feature number 214, k__Bacteria.p__Firmicutes.c__Cl
## [233] "2023-01-17 09:03:41 INFO::Fitting model to feature number 215, k__Bacteria.p__Firmicutes.c__Cl
## [234] "2023-01-17 09:03:41 INFO::Fitting model to feature number 216, k__Bacteria.p__Firmicutes.c__Cl
## [235] "2023-01-17 09:03:41 INFO::Fitting model to feature number 217, k__Bacteria.p__Firmicutes.c__Cl
## [236] "2023-01-17 09:03:41 INFO::Fitting model to feature number 218, k__Bacteria.p__Firmicutes.c__Cl
## [237] "2023-01-17 09:03:41 INFO::Fitting model to feature number 219, k__Bacteria.p__Firmicutes.c__Cl
## [238] "2023-01-17 09:03:41 INFO::Fitting model to feature number 220, k__Bacteria.p__Firmicutes.c__Cl
## [239] "2023-01-17 09:03:41 INFO::Fitting model to feature number 221, k__Bacteria.p__Firmicutes.c__Cl
## [240] "2023-01-17 09:03:41 INFO::Fitting model to feature number 222, k__Bacteria.p__Firmicutes.c__Cl
## [241] "2023-01-17 09:03:41 INFO::Fitting model to feature number 223, k__Bacteria.p__Firmicutes.c__Cl
## [242] "2023-01-17 09:03:42 INFO::Fitting model to feature number 224, k__Bacteria.p__Firmicutes.c__Cl
## [243] "2023-01-17 09:03:42 INFO::Fitting model to feature number 225, k__Bacteria.p__Firmicutes.c__Cl
## [244] "2023-01-17 09:03:42 INFO::Fitting model to feature number 226, k__Bacteria.p__Firmicutes.c__Cl
## [245] "2023-01-17 09:03:42 INFO::Fitting model to feature number 227, k__Bacteria.p__Firmicutes.c__Cl
## [246] "2023-01-17 09:03:42 INFO::Fitting model to feature number 228, k__Bacteria.p__Firmicutes.c__Cl
## [247] "2023-01-17 09:03:42 INFO::Fitting model to feature number 229, k__Bacteria.p__Firmicutes.c__Cl
## [248] "2023-01-17 09:03:42 INFO::Fitting model to feature number 230, k__Bacteria.p__Firmicutes.c__Cl
## [249] "2023-01-17 09:03:42 INFO::Fitting model to feature number 231, k__Bacteria.p__Firmicutes.c__Cl
## [250] "2023-01-17 09:03:42 INFO::Fitting model to feature number 232, k__Bacteria.p__Firmicutes.c__Cl
## [251] "2023-01-17 09:03:42 INFO::Fitting model to feature number 233, k__Bacteria.p__Firmicutes.c__Cl
## [252] "2023-01-17 09:03:42 INFO::Fitting model to feature number 234, k__Bacteria.p__Firmicutes.c__Cl
## [253] "2023-01-17 09:03:42 INFO::Fitting model to feature number 235, k__Bacteria.p__Firmicutes.c__Cl
## [254] "2023-01-17 09:03:42 INFO::Fitting model to feature number 236, k__Bacteria.p__Firmicutes.c__E
## [255] "2023-01-17 09:03:43 INFO::Fitting model to feature number 237, k__Bacteria.p__Firmicutes.c__E
## [256] "2023-01-17 09:03:43 INFO::Fitting model to feature number 238, k__Bacteria.p__Firmicutes.c__E
## [257] "2023-01-17 09:03:43 INFO::Fitting model to feature number 239, k__Bacteria.p__Firmicutes.c__E
## [258] "2023-01-17 09:03:43 INFO::Fitting model to feature number 240, k__Bacteria.p__Firmicutes.c__E
## [259] "2023-01-17 09:03:43 INFO::Fitting model to feature number 241, k__Bacteria.p__Firmicutes.c__E
## [260] "2023-01-17 09:03:43 INFO::Fitting model to feature number 242, k__Bacteria.p__Firmicutes.c__E
## [261] "2023-01-17 09:03:43 INFO::Fitting model to feature number 243, k__Bacteria.p__Firmicutes.c__E
## [262] "2023-01-17 09:03:43 INFO::Fitting model to feature number 244, k__Bacteria.p__Firmicutes.c__E
## [263] "2023-01-17 09:03:43 INFO::Fitting model to feature number 245, k__Bacteria.p__Firmicutes.c__E
## [264] "2023-01-17 09:03:43 INFO::Fitting model to feature number 246, k__Bacteria.p__Firmicutes.c__E
## [265] "2023-01-17 09:03:43 INFO::Fitting model to feature number 247, k__Bacteria.p__Firmicutes.c__E
## [266] "2023-01-17 09:03:43 INFO::Fitting model to feature number 248, k__Bacteria.p__Firmicutes.c__E
## [267] "2023-01-17 09:03:43 INFO::Fitting model to feature number 249, k__Bacteria.p__Firmicutes.c__E
## [268] "2023-01-17 09:03:44 INFO::Fitting model to feature number 250, k__Bacteria.p__Firmicutes.c__E
## [269] "2023-01-17 09:03:44 INFO::Fitting model to feature number 251, k__Bacteria.p__Firmicutes.c__E
## [270] "2023-01-17 09:03:44 INFO::Fitting model to feature number 252, k__Bacteria.p__Firmicutes.c__E
## [271] "2023-01-17 09:03:44 INFO::Fitting model to feature number 253, k__Bacteria.p__Firmicutes.c__E
## [272] "2023-01-17 09:03:44 INFO::Fitting model to feature number 254, k__Bacteria.p__Firmicutes.c__E
## [273] "2023-01-17 09:03:44 INFO::Fitting model to feature number 255, k__Bacteria.p__Firmicutes.c__E
```

```
## [274] "2023-01-17 09:03:44 INFO::Fitting model to feature number 256, k__Bacteria.p__Firmicutes.c__E
## [275] "2023-01-17 09:03:44 INFO::Fitting model to feature number 257, k__Bacteria.p__Firmicutes.c__E
## [276] "2023-01-17 09:03:44 INFO::Fitting model to feature number 258, k__Bacteria.p__Firmicutes.c__E
## [277] "2023-01-17 09:03:44 INFO::Fitting model to feature number 259, k__Bacteria.p__Firmicutes.c__F
## [278] "2023-01-17 09:03:44 INFO::Fitting model to feature number 260, k__Bacteria.p__Firmicutes.c__F
## [279] "2023-01-17 09:03:44 INFO::Fitting model to feature number 261, k__Bacteria.p__Firmicutes.c__F
## [280] "2023-01-17 09:03:45 INFO::Fitting model to feature number 262, k__Bacteria.p__Firmicutes.c__F
## [281] "2023-01-17 09:03:45 INFO::Fitting model to feature number 263, k__Bacteria.p__Firmicutes.c__F
## [282] "2023-01-17 09:03:45 INFO::Fitting model to feature number 264, k__Bacteria.p__Firmicutes.c__F
## [283] "2023-01-17 09:03:45 INFO::Fitting model to feature number 265, k__Bacteria.p__Firmicutes.c__F
## [284] "2023-01-17 09:03:45 INFO::Fitting model to feature number 266, k__Bacteria.p__Firmicutes.c__F
## [285] "2023-01-17 09:03:45 INFO::Fitting model to feature number 267, k__Bacteria.p__Firmicutes.c__F
## [286] "2023-01-17 09:03:45 INFO::Fitting model to feature number 268, k__Bacteria.p__Firmicutes.c__N
## [287] "2023-01-17 09:03:45 INFO::Fitting model to feature number 269, k__Bacteria.p__Firmicutes.c__N
## [288] "2023-01-17 09:03:45 INFO::Fitting model to feature number 270, k__Bacteria.p__Firmicutes.c__N
## [289] "2023-01-17 09:03:45 INFO::Fitting model to feature number 271, k__Bacteria.p__Firmicutes.c__N
## [290] "2023-01-17 09:03:45 INFO::Fitting model to feature number 272, k__Bacteria.p__Firmicutes.c__N
## [291] "2023-01-17 09:03:45 INFO::Fitting model to feature number 273, k__Bacteria.p__Firmicutes.c__N
## [292] "2023-01-17 09:03:45 INFO::Fitting model to feature number 274, k__Bacteria.p__Firmicutes.c__N
## [293] "2023-01-17 09:03:46 INFO::Fitting model to feature number 275, k__Bacteria.p__Firmicutes.c__N
## [294] "2023-01-17 09:03:46 INFO::Fitting model to feature number 276, k__Bacteria.p__Firmicutes.c__N
## [295] "2023-01-17 09:03:46 INFO::Fitting model to feature number 277, k__Bacteria.p__Firmicutes.c__N
## [296] "2023-01-17 09:03:46 INFO::Fitting model to feature number 278, k__Bacteria.p__Firmicutes.c__N
## [297] "2023-01-17 09:03:46 INFO::Fitting model to feature number 279, k__Bacteria.p__Firmicutes.c__N
## [298] "2023-01-17 09:03:46 INFO::Fitting model to feature number 280, k__Bacteria.p__Firmicutes.c__N
## [299] "2023-01-17 09:03:46 INFO::Fitting model to feature number 281, k__Bacteria.p__Firmicutes.c__N
## [300] "2023-01-17 09:03:46 INFO::Fitting model to feature number 282, k__Bacteria.p__Firmicutes.c__N
## [301] "2023-01-17 09:03:46 INFO::Fitting model to feature number 283, k__Bacteria.p__Firmicutes.c__N
## [302] "2023-01-17 09:03:46 INFO::Fitting model to feature number 284, k__Bacteria.p__Firmicutes.c__N
## [303] "2023-01-17 09:03:46 INFO::Fitting model to feature number 285, k__Bacteria.p__Firmicutes.c__N
## [304] "2023-01-17 09:03:46 INFO::Fitting model to feature number 286, k__Bacteria.p__Firmicutes.c__N
## [305] "2023-01-17 09:03:47 INFO::Fitting model to feature number 287, k__Bacteria.p__Firmicutes.c__N
## [306] "2023-01-17 09:03:47 INFO::Fitting model to feature number 288, k__Bacteria.p__Firmicutes.c__N
## [307] "2023-01-17 09:03:47 INFO::Fitting model to feature number 289, k__Bacteria.p__Firmicutes.c__T
## [308] "2023-01-17 09:03:47 INFO::Fitting model to feature number 290, k__Bacteria.p__Firmicutes.c__T
## [309] "2023-01-17 09:03:47 INFO::Fitting model to feature number 291, k__Bacteria.p__Firmicutes.c__T
## [310] "2023-01-17 09:03:47 INFO::Fitting model to feature number 292, k__Bacteria.p__Proteobacteria"
## [311] "2023-01-17 09:03:47 INFO::Fitting model to feature number 293, k__Bacteria.p__Proteobacteria.
## [312] "2023-01-17 09:03:47 INFO::Fitting model to feature number 294, k__Bacteria.p__Proteobacteria.
## [313] "2023-01-17 09:03:47 INFO::Fitting model to feature number 295, k__Bacteria.p__Proteobacteria.
## [314] "2023-01-17 09:03:47 INFO::Fitting model to feature number 296, k__Bacteria.p__Proteobacteria.
## [315] "2023-01-17 09:03:47 INFO::Fitting model to feature number 297, k__Bacteria.p__Proteobacteria.
## [316] "2023-01-17 09:03:47 INFO::Fitting model to feature number 298, k__Bacteria.p__Proteobacteria.
## [317] "2023-01-17 09:03:48 INFO::Fitting model to feature number 299, k__Bacteria.p__Proteobacteria.
## [318] "2023-01-17 09:03:48 INFO::Fitting model to feature number 300, k__Bacteria.p__Proteobacteria.
## [319] "2023-01-17 09:03:48 INFO::Fitting model to feature number 301, k__Bacteria.p__Proteobacteria.
## [320] "2023-01-17 09:03:48 INFO::Fitting model to feature number 302, k__Bacteria.p__Proteobacteria.
## [321] "2023-01-17 09:03:48 INFO::Fitting model to feature number 303, k__Bacteria.p__Proteobacteria.
## [322] "2023-01-17 09:03:48 INFO::Fitting model to feature number 304, k__Bacteria.p__Proteobacteria.
## [323] "2023-01-17 09:03:48 INFO::Fitting model to feature number 305, k__Bacteria.p__Proteobacteria.
## [324] "2023-01-17 09:03:48 INFO::Fitting model to feature number 306, k__Bacteria.p__Proteobacteria.
## [325] "2023-01-17 09:03:48 INFO::Fitting model to feature number 307, k__Bacteria.p__Proteobacteria.
## [326] "2023-01-17 09:03:48 INFO::Fitting model to feature number 308, k__Bacteria.p__Proteobacteria.
## [327] "2023-01-17 09:03:48 INFO::Fitting model to feature number 309, k__Bacteria.p__Proteobacteria.
```

```
## [328] "2023-01-17 09:03:48 INFO::Fitting model to feature number 310, k__Bacteria.p__Proteobacteria.c
## [329] "2023-01-17 09:03:48 INFO::Fitting model to feature number 311, k__Bacteria.p__Proteobacteria.c
## [330] "2023-01-17 09:03:49 INFO::Fitting model to feature number 312, k__Bacteria.p__Proteobacteria.c
## [331] "2023-01-17 09:03:49 INFO::Fitting model to feature number 313, k__Bacteria.p__Proteobacteria.c
## [332] "2023-01-17 09:03:49 INFO::Fitting model to feature number 314, k__Bacteria.p__Proteobacteria.c
## [333] "2023-01-17 09:03:49 INFO::Fitting model to feature number 315, k__Bacteria.p__Proteobacteria.c
## [334] "2023-01-17 09:03:49 INFO::Fitting model to feature number 316, k__Bacteria.p__Proteobacteria.c
## [335] "2023-01-17 09:03:49 INFO::Fitting model to feature number 317, k__Bacteria.p__Proteobacteria.c
## [336] "2023-01-17 09:03:49 INFO::Fitting model to feature number 318, k__Bacteria.p__Proteobacteria.c
## [337] "2023-01-17 09:03:49 INFO::Fitting model to feature number 319, k__Bacteria.p__Proteobacteria.c
## [338] "2023-01-17 09:03:49 INFO::Fitting model to feature number 320, k__Bacteria.p__Proteobacteria.c
## [339] "2023-01-17 09:03:49 INFO::Fitting model to feature number 321, k__Bacteria.p__Proteobacteria.c
## [340] "2023-01-17 09:03:49 INFO::Fitting model to feature number 322, k__Bacteria.p__Proteobacteria.c
## [341] "2023-01-17 09:03:49 INFO::Fitting model to feature number 323, k__Bacteria.p__Proteobacteria.c
## [342] "2023-01-17 09:03:50 INFO::Fitting model to feature number 324, k__Bacteria.p__Verrucomicrobia
## [343] "2023-01-17 09:03:50 INFO::Fitting model to feature number 325, k__Bacteria.p__Verrucomicrobia
## [344] "2023-01-17 09:03:50 INFO::Fitting model to feature number 326, k__Bacteria.p__Verrucomicrobia
## [345] "2023-01-17 09:03:50 INFO::Fitting model to feature number 327, k__Bacteria.p__Verrucomicrobia
## [346] "2023-01-17 09:03:50 INFO::Fitting model to feature number 328, k__Bacteria.p__Verrucomicrobia
## [347] "2023-01-17 09:03:50 INFO::Fitting model to feature number 329, k__Bacteria.p__Verrucomicrobia
## [348] "2023-01-17 09:03:50 INFO::Counting total values for each feature"
## [349] "2023-01-17 09:03:50 WARNING::Deleting existing residuals file: maaslin_fixed_asthma_with_adoni
## [350] "2023-01-17 09:03:50 INFO::Writing residuals to file maaslin_fixed_asthma_with_adonis_randomefl
## [351] "2023-01-17 09:03:50 WARNING::Deleting existing fitted file: maaslin_fixed_asthma_with_adonis_r
## [352] "2023-01-17 09:03:50 INFO::Writing fitted values to file maaslin_fixed_asthma_with_adonis_rando
## [353] "2023-01-17 09:03:50 WARNING::Deleting existing ranef file: maaslin_fixed_asthma_with_adonis_ra
## [354] "2023-01-17 09:03:50 INFO::Writing extracted random effects to file maaslin_fixed_asthma_with_a
## [355] "2023-01-17 09:03:50 INFO::Writing all results to file (ordered by increasing q-values): maasli
## [356] "2023-01-17 09:03:50 INFO::Writing the significant results (those which are less than or equal
## [357] "2023-01-17 09:03:50 INFO::Writing heatmap of significant results to file: maaslin_fixed_asthma
## [358] "[1] \"There is not enough metadata in the associations to create a heatmap plot. Please review
## [359] "2023-01-17 09:03:50 INFO::Writing association plots (one for each significant association) to
## [360] "2023-01-17 09:03:50 INFO::Plotting associations from most to least significant, grouped by met
## [361] "2023-01-17 09:03:50 INFO::Plotting data for metadata number 1, ASTHMA"
## [362] "2023-01-17 09:03:50 INFO::Creating boxplot for categorical data, ASTHMA vs k__Bacteria.p__Bact
## [363] "2023-01-17 09:03:50 INFO::Creating boxplot for categorical data, ASTHMA vs k__Bacteria.p__Bact
## [364] "2023-01-17 09:03:51 INFO::Creating boxplot for categorical data, ASTHMA vs k__Bacteria.p__Bact
## [365] "2023-01-17 09:03:51 INFO::Creating boxplot for categorical data, ASTHMA vs k__Bacteria.p__Firm
## [366] "2023-01-17 09:03:51 INFO::Creating boxplot for categorical data, ASTHMA vs k__Bacteria.p__Firm
## [367] "2023-01-17 09:03:51 INFO::Creating boxplot for categorical data, ASTHMA vs k__Bacteria.p__Firm
## [368] "2023-01-17 09:03:51 INFO::Creating boxplot for categorical data, ASTHMA vs k__Bacteria.p__Firm
```

```r
sig_bugs <- fit_data$results[fit_data$results[,"qval"] < 0.05,]
plotme <- sig_bugs$feature[grep(sig_bugs$feature, pattern = "s__")]

allSpecies <- readRDS("allSpecies_95_relab_tab.rds")
basedir_humann3="./"
humann3_mf_filtered_perm <- readRDS(file = paste0(basedir_humann3,
↪  "humann3_mf_filtered.rds"))
row.names(humann3_mf_filtered_perm) <- humann3_mf_filtered_perm$SampleName
sampleNames <- gsub(rownames(humann3_mf_filtered_perm), pattern = "[-]", replacement=".")
asv.clean <- allSpecies[,sampleNames]
asv.clean <- t(asv.clean[rowSums(asv.clean)>0,])
df.meta<- humann3_mf_filtered_perm
```

```
plotme.df <- asv.clean[, gsub(as.character(plotme), pattern = "[.]", replacement = "|")]
plotme.df.melted <- reshape2::melt(plotme.df)
names(plotme.df.melted) <- c("Microbiome", "variable", "value")
plotme.df.melted$asthma <- factor(df.meta[gsub(plotme.df.melted$Microbiome, pattern =
↪  "[.]", replacement = "-"), "ASTHMA"], levels = c("Healthy", "Asthmatic"),
↪  labels=c("Healthy", "Asthmatic"))
plotme.df.melted$AgeGroup <- factor(df.meta[gsub(plotme.df.melted$Microbiome, pattern =
↪  "[.]", replacement = "-"), "AgeGroup"], levels = c("Pediatric", "Adult"))
plotme.df.melted$variable <- factor(plotme.df.melted$variable, levels =
↪  c("k__Bacteria|p__Firmicutes|c__Clostridia|o__Clostridiales|f__Lachnospiraceae|g__Lachnospiraceae_u
↪  "k__Bacteria|p__Bacteroidetes|c__Bacteroidia|o__Bacteroidales|f__Prevotellaceae|g__Prevotella|s__Pr
↪  labels = c("Eubacterium rectale", "Prevotella copri"))
plotme.df.melted$cohort <- paste0(plotme.df.melted$asthma, plotme.df.melted$AgeGroup)
relabAlphaDivPlot3 <- ggplot(plotme.df.melted, aes(x=asthma, y=asin(sqrt(value)),
↪  shape=asthma)) +
  geom_boxplot(outlier.shape=NA) +
  ggbeeswarm::geom_quasirandom(aes(shape=asthma, fill=cohort), size=3.5, alpha=0.9)+
    # geom_point(position=position_beeswarm(cex=4), aes(shape=cohort,  fill=cohort),
     ↪  size=3.5) +
  scale_fill_manual(values = c(brewer.pal(7, "BrBG")[c(1,7, 3,5)]))+ #3,5,1,7
                     # values=c("#DE4968FF", "#51127CFF"))+# "#DE4968FF", "#51127CFF",
                     ↪  "#51127CFF")) +
    theme_classic()+
    facet_grid( ~ variable, scales = "free_x", space = "free_x") +
    scale_shape_manual(values=c(22, 21)) +
    scale_color_manual(values=c("black", "black","black", "black")) +
    scale_x_discrete(name = "Microbiota") +
    scale_y_continuous(name="Arcsine Transformed Relative Abundance") +
    theme(panel.grid.major = element_blank(),
          panel.grid.minor = element_blank(),
          axis.line.x = element_line(size = 0, colour = "black"),
          axis.line.y = element_line(size = 0, colour = "black"),
          axis.line = element_line(size=1, colour = "black"),
          panel.border = element_rect(fill = NA, colour = "black", size = 1),
          panel.background = element_blank(),
          text=element_text(size = 12),
          axis.text.x=element_text(colour="black", size = 12),
          axis.text.y=element_text(colour="black", size = 12),
          axis.title.x=element_blank(),
          legend.position = "none",
          legend.title=element_blank(),
          legend.key = element_rect(color=NA, fill=NA)
    )

relabAlphaDivPlot3
```

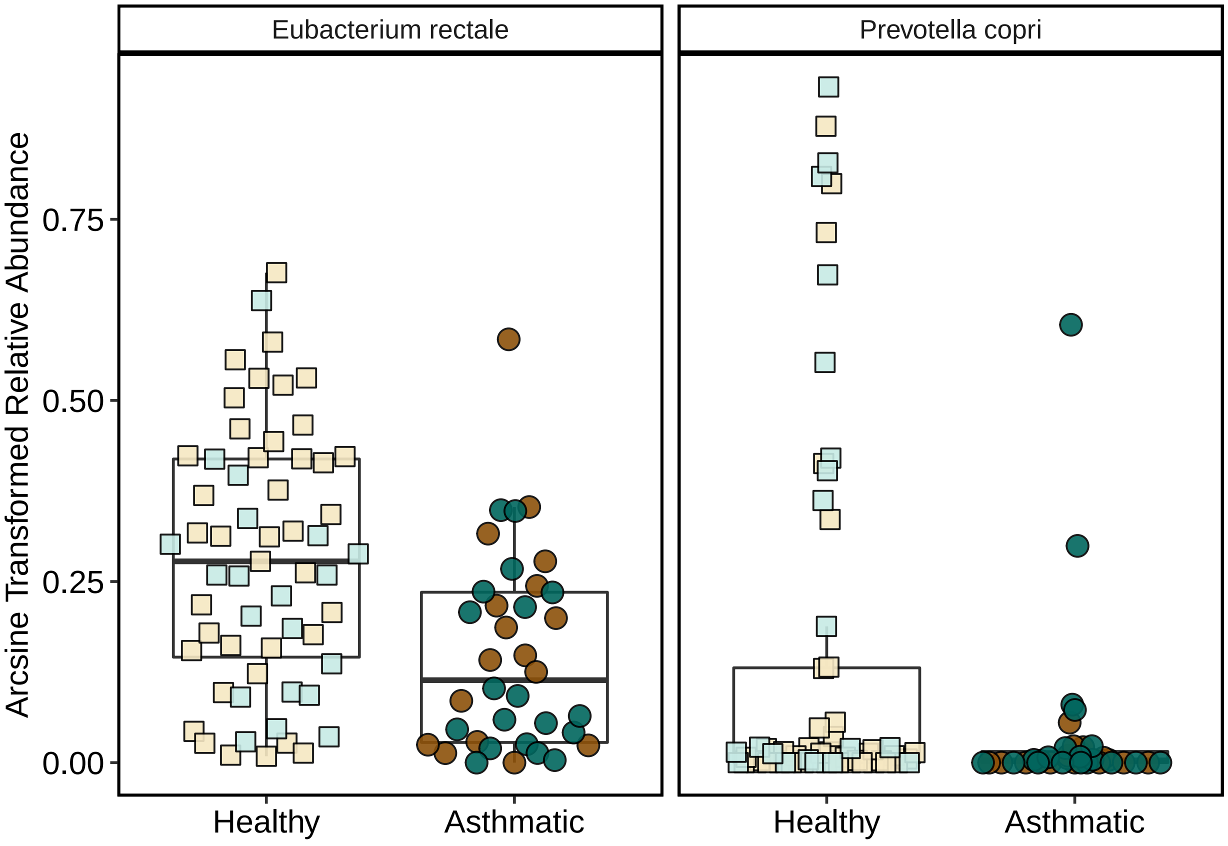


```
# ggsave(filename = "metaphlan3_maaslin_DA_bugs_4Colors.pdf", plot = relabAlphaDivPlot3,
↪  device = "pdf", height = 10, width = 12, units = "cm", useDingbats=F)
```

#### Load libraries/functions for Figures 2,3,4

```
#### load libraries ####
rm(list = ls())
library(ggplot2)
library(ggpubr)
library(ggbeeswarm)
library(magrittr)
```

```
##
## Attaching package: 'magrittr'

## The following object is masked from 'package:purrr':
##
##     set_names

## The following object is masked from 'package:tidyr':
##
##     extract
```

```
# borrowed a lot of code from here:
#
↪  https://thejacksonlaboratory.github.io/microbiome-workshop/jekyll/update/2017/11/17/Analysing-mWGS-
```

```r
#### FUNCTIONS ####

# plotting etc functions from humann3 project:
source("./humann3_210613_functions_ngw.R")

removeColumnSuffix <- function(DATAFRAME, SUFFIX) {
    # Remove the suffix from the sample labels (ex. '_shortbred_RPKM')
    names(DATAFRAME) <- gsub(SUFFIX, '', names(DATAFRAME))
    names(DATAFRAME) <- gsub('\\.', '-', names(DATAFRAME))
    return(DATAFRAME)
}

reorderColumnsToMatchMetaData <- function(DATAFRAME, METADATAFRAME) {
    # Re-order the columns so that they correspond to the order of samples in
    ↪  METADATAFRAME
    DATAFRAME <- DATAFRAME[,row.names(METADATAFRAME)]
    return(DATAFRAME)
}

filter_shortbred_data <- function(indata, mymetadata, my_suffix="_SB_CARD21_RPKM") {
  outdata <- removeColumnSuffix(indata, my_suffix) # remove suffix
  if(all(row.names(mymetadata) %in% names(outdata))){
    # drop disqualifieds and exacerbations (N=95)
    outdata <- outdata[, row.names(mymetadata)]
    # drop genes of zero abundance: goes from 977 to 95
    outdata = outdata[rowSums(outdata) > 0 ,]
    # make sure there is enough data in each column (>=7 instances)
    LowGenes = row.names(outdata)[rowSums(ifelse(outdata>0, 1, 0)) < 7]
    length(LowGenes) # 86
    outdata = outdata[row.names(outdata) %notin% LowGenes,]
    return(outdata)
  } else {
    print("Error")
  }
}

boxplotByAsthmaAndAge <- function(mydat, my_ylab="Number of CARD hits (nonzero RPKM)",
↪  myPalette=c("#C7EAE5", "#01665E")){
  outplot <- ggplot(data = mydat, aes(y=value, x=ASTHMA, fill=ASTHMA, shape=AgeGroup)) +
↪  # remove col to remove age groupings

      # what I had before:
      # geom_boxplot(outlier.shape=NA, lwd=0.5, alpha=1) +
      # geom_quasirandom(dodge.width = 0.8, width = 0.1, size=4, alpha=0.7)+

      geom_boxplot(outlier.shape=NA, lwd=0.5, alpha=0.6) +
      geom_quasirandom(dodge.width = 0.8, width = 0.1, size=4, alpha=0.8)+ # remove dodge
      ↪  width if not doing age groupings
      scale_shape_manual(values=c(21, 22)) +
      scale_fill_manual(values = myPalette) +
      scale_y_continuous(name=my_ylab) +
      theme(panel.grid.major = element_blank(),
          panel.grid.minor = element_blank(),
```

```
          axis.line.x = element_line(size = 0, colour = "black"),
          axis.line.y = element_line(size = 0, colour = "black"),
          axis.line = element_line(size=1, colour = "black"),
          panel.border = element_rect(fill = NA, colour = "black", size = 1),
          panel.background = element_blank(),
          text=element_text(size = 16),
          axis.text.x=element_text(colour="black", size = 12),
          axis.text.y=element_text(colour="black", size = 12),
          axis.title.x=element_blank(),
          legend.title=element_blank(),
          # legend.key = element_rect(color=NA, fill=NA),
          legend.key = element_rect(fill = "white"))+
    stat_compare_means(method="wilcox.test", aes(label = "p.signif"), label.y =
    ↪  (max(mydat$value)*1.1))
  # CARDRPKMDerivedHitsPlot + guides(fill="none")
  outplot <- outplot + guides(fill="none")
  return(outplot)
}

boxplotByAsthma <- function(mydat, my_ylab="Number of CARD hits (nonzero RPKM)",
↪  myPalette=c("#C7EAE5", "#01665E")){
  outplot <- ggplot(data = mydat, aes(y=value, x=ASTHMA, fill=ASTHMA, shape=AgeGroup)) +
↪   # remove col to remove age groupings
    # geom_boxplot(outlier.shape=NA, lwd=0.5, alpha=1) +
    # geom_beeswarm(dodge.width = 0.8, width = 0.3)+ # remove dodge width if not doing
    ↪  age groupings
    # geom_quasirandom(size=4, alpha=0.5, width=0.2)+ # remove dodge width if not doing
    ↪  age groupings
    geom_boxplot(mapping = aes(y=value, x=ASTHMA, fill=ASTHMA),inherit.aes = FALSE,
    ↪  outlier.shape=NA, lwd=0.5, alpha=0.6) +
    geom_quasirandom(width = 0.2, size=4, alpha=0.8)+ # remove dodge width if not doing
    ↪  age groupings
    # scale_color_manual(values = myPalette) +
    # scale_shape_manual(values=c(16, 15)) +
    # scale_shape_manual(values=c(21, 22)) +
    scale_shape_manual(values=c(21, 22)) +
    scale_fill_manual(values = myPalette) +
    scale_y_continuous(name=my_ylab) +
    theme(panel.grid.major = element_blank(),
        panel.grid.minor = element_blank(),
        axis.line.x = element_line(size = 0, colour = "black"),
        axis.line.y = element_line(size = 0, colour = "black"),
        axis.line = element_line(size=1, colour = "black"),
        panel.border = element_rect(fill = NA, colour = "black", size = 1),
        panel.background = element_blank(),
        text=element_text(size = 16),
        axis.text.x=element_text(colour="black", size = 12),
        axis.text.y=element_text(colour="black", size = 12),
        axis.title.x=element_blank(),
        legend.title=element_blank(),
        # legend.key = element_rect(color=NA, fill=NA),
        legend.key = element_rect(fill = "white"),
        legend.position = "none") +
```

```
  stat_compare_means(method="wilcox.test", aes(label = "p.signif"), label.y =
   ↪   (max(mydat$value)*1.1))
  return(outplot)
}
```

#### Filter Data

```
rm(list=ls()[ls() %notin% c("%notin%", "makePCA", "makeNMDSplot",
↪  "boxplotByAsthmaAndAge", "boxplotByAsthma", "filter_shortbred_data",
↪  "removeColumnSuffix", "reorderColumnsToMatchMetaData")])

#### import data ####
basedir="./"
humann3_mf_filtered_perm <- readRDS(file = paste0(basedir, "humann3_mf_filtered.rds"))
row.names(humann3_mf_filtered_perm) <- humann3_mf_filtered_perm$SampleName # make row
↪  names match data colnames

# Load the ShortBRED results, in which rows are antibiotic resistance markers and columns
↪  are samples
df.shortbred <- read.table('~/Library/CloudStorage/Box-Box/Kau
↪  Lab/Results/MARS/FecalMetagenomics/shortbred/CARD2021results/MARS_SB_CARD21_RPKM.tsv',
↪  row.names=1, header=T)  # 888 104
# head(df.shortbred)
length(rowSums(df.shortbred)[rowSums(df.shortbred) > 0]) # 170
```

```
## [1] 170
```

```
df.shortbredF <- filter_shortbred_data(indata = df.shortbred, mymetadata =
↪  humann3_mf_filtered_perm)
dim(df.shortbredF) # 71 ARGs
```

```
## [1] 71 95
```

```
write.csv(df.shortbredF, "SB_CARD21_RPKM_filtered.csv")
saveRDS(df.shortbredF, "SB_CARD21_RPKM_filtered.rds")

df.CARDhits <- read.table('~/Library/CloudStorage/Box-Box/Kau
↪  Lab/Results/MARS/FecalMetagenomics/shortbred/CARD2021results/MARS_SB_CARD21_HITS.tsv',
↪  row.names=1, header=T)  # 888 104
# head(df.CARDhits)
# length(rowSums(df.CARDhits)[rowSums(df.CARDhits) > 0]) # 258
df.CARDhitsF <- filter_shortbred_data(indata = df.CARDhits, mymetadata =
↪  humann3_mf_filtered_perm, my_suffix = "_SB_CARD21_HITS")
write.csv(df.CARDhitsF, "SB_CARD21_HITS_filtered.csv")
saveRDS(df.CARDhitsF, "SB_CARD21_HITS_filtered.rds")
```

#### Figure 3

#### Figure 3A:

PCA and Bray-Curtis Beta Diversity NMDS

```r
rm(list=ls()[ls() %notin% c("%notin%", "makePCA", "makeNMDSplot",
↪  "boxplotByAsthmaAndAge", "boxplotByAsthma", "filter_shortbred_data",
↪  "removeColumnSuffix")])
source("./humann3_210613_functions_ngw.R")
# setwd("~/Library/CloudStorage/Box-Box/Kau
↪  Lab/Results/MARS/FecalMetagenomics/shortbred/CARD2021results/")

library(ggplot2)
library(ape)
library(stats)
library(vegan)
library(ggfortify)
library(viridis)
library(RColorBrewer)

basedir="./"
humann3_mf_filtered_perm <- readRDS(file = paste0(basedir, "humann3_mf_filtered.rds"))
row.names(humann3_mf_filtered_perm) <- humann3_mf_filtered_perm$SampleName
df.meta <- humann3_mf_filtered_perm
df.shortbred <- readRDS("SB_CARD21_RPKM_filtered.rds")

# Total sum square the data for Bray-curtis transform
df.shortbred.tss <- apply(X = df.shortbred, MARGIN = 2, FUN = function(x){x/sum(x)})
if(!all(names(df.shortbred.tss) == row.names(df.meta))){
  print("ERROR: meta data frame is not in the same order as sample data frame")
}
# PCA: saves and plots
makePCA(mydata = df.shortbred.tss, mymetadata = df.meta,
↪  my_data_table_name="SB_CARD_TSS_RPKM", mytitle = "ShortBRED RPKM PCA", basedir="./")
```

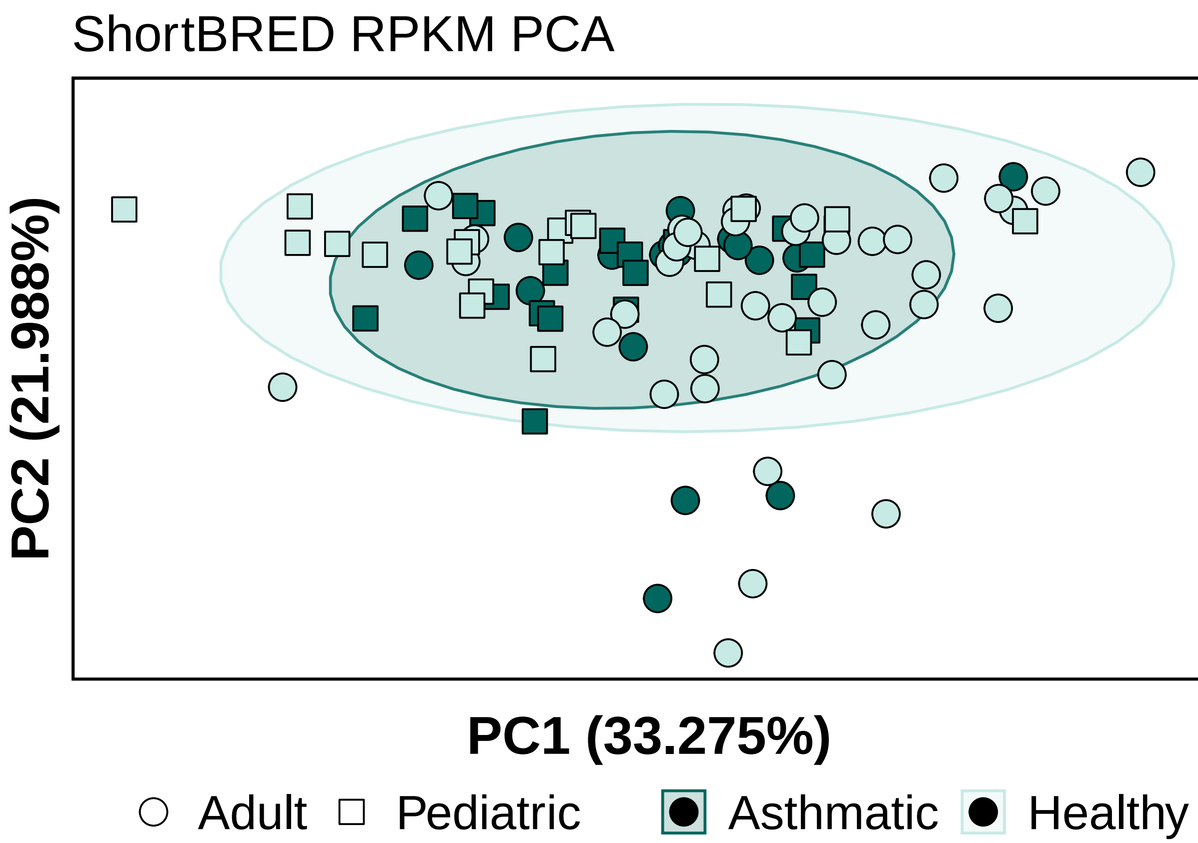


```r
# PERMANOVAs
# rpkm_euclid = vegdist(t(df.shortbred), method = "euclidean") # no tss
rpkm_euclid = vegdist(t(df.shortbred.tss), method = "euclidean") # tss
adonis2(rpkm_euclid ~ ASTHMA*AgeGroup, data = df.meta, permutations = 10000)
```

```
## Permutation test for adonis under reduced model
## Terms added sequentially (first to last)
## Permutation: free
## Number of permutations: 10000
##
## adonis2(formula = rpkm_euclid ~ ASTHMA * AgeGroup, data = df.meta, permutations = 10000)
##                 Df SumOfSqs      R2      F    Pr(>F)
## ASTHMA           1   0.2640 0.02995 3.0702  0.009599 **
## AgeGroup         1   0.5798 0.06576 6.7420 9.999e-05 ***
## ASTHMA:AgeGroup  1   0.1468 0.01665 1.7066  0.116588
## Residual        91   7.8259 0.88764
## Total           94   8.8165 1.00000
## ---
## Signif. codes:  0 '***' 0.001 '**' 0.01 '*' 0.05 '.' 0.1 ' ' 1
```

```r
# Bray-Curtis:
# dist = vegdist(t(df.shortbred), method = "bray", binary = FALSE)
dist = vegdist(t(df.shortbred.tss), method = "bray", binary = FALSE) # tss
# adonis2(dist ~ ASTHMA, data = df.meta, permutations = 5000)
adonis2(dist ~ ASTHMA*AgeGroup, data = df.meta, permutations = 100000)
```

```
## Permutation test for adonis under reduced model
## Terms added sequentially (first to last)
```

```
## Permutation: free
## Number of permutations: 1e+05
##
## adonis2(formula = dist ~ ASTHMA * AgeGroup, data = df.meta, permutations = 1e+05)
##                  Df SumOfSqs      R2      F  Pr(>F)
## ASTHMA            1   0.4073 0.02799 2.8249 0.00502 **
## AgeGroup          1   0.8107 0.05572 5.6232   2e-05 ***
## ASTHMA:AgeGroup   1   0.2133 0.01466 1.4796 0.14439
## Residual         91  13.1198 0.90164
## Total            94  14.5512 1.00000
## ---
## Signif. codes:  0 '***' 0.001 '**' 0.01 '*' 0.05 '.' 0.1 ' ' 1
```

```r
# NMDS
sbRPKM.msds <- metaMDS(t(df.shortbred.tss), distance = "bray", k = 5, trymax = 100)
```

```
## Run 0 stress 0.09572094
## Run 1 stress 0.0957223
## ... Procrustes: rmse 0.0006238534  max resid 0.002361161
## ... Similar to previous best
## Run 2 stress 0.09702725
## Run 3 stress 0.09667829
## Run 4 stress 0.0965091
## Run 5 stress 0.09608921
## ... Procrustes: rmse 0.01283582  max resid 0.06927543
## Run 6 stress 0.09834136
## Run 7 stress 0.09587901
## ... Procrustes: rmse 0.007631764  max resid 0.03574405
## Run 8 stress 0.09685854
## Run 9 stress 0.09572215
## ... Procrustes: rmse 0.0005509063  max resid 0.002216219
## ... Similar to previous best
## Run 10 stress 0.09741856
## Run 11 stress 0.09659488
## Run 12 stress 0.09572149
## ... Procrustes: rmse 0.00098679  max resid 0.003889446
## ... Similar to previous best
## Run 13 stress 0.09840121
## Run 14 stress 0.09572168
## ... Procrustes: rmse 0.001201942  max resid 0.004500401
## ... Similar to previous best
## Run 15 stress 0.09572322
## ... Procrustes: rmse 0.0004873207  max resid 0.00180228
## ... Similar to previous best
## Run 16 stress 0.0957221
## ... Procrustes: rmse 0.001385092  max resid 0.005315657
## ... Similar to previous best
## Run 17 stress 0.09638596
## Run 18 stress 0.09854131
## Run 19 stress 0.09574201
## ... Procrustes: rmse 0.003279252  max resid 0.01261642
## Run 20 stress 0.09572533
## ... Procrustes: rmse 0.001742022  max resid 0.006746942
## ... Similar to previous best
```

```
## *** Solution reached
```

```
# saveRDS(sbRPKM.msds, "sbRPKM_brayCurtis_k3.msds.rds")
saveRDS(sbRPKM.msds, "sbRPKM_brayCurtis_k5.msds.rds")
# makeNMDSplot(data.msds = sbRPKM.msds,
#              mymetadata = df.meta,
#              my_data_table_name = "SB_CARD_TSS_RPKM",
#              mytitle = "ShortBRED RPKM Bray-Curtis",
#              basedir="./")

# 4 grouper ####
basedir="./"
my_data_table_name = "SB_CARD_TSS_RPKM"
mytitle = "ShortBRED RPKM Bray-Curtis"
confidenceEllipse = 0.95
brayCurtisMDS <- data.frame(sbRPKM.msds$points)
brayCurtisMDS$AgeGroup <-
↪  factor(df.meta[as.character(row.names(brayCurtisMDS)),"AgeGroup"])
brayCurtisMDS$Asthma <- factor(df.meta[as.character(row.names(brayCurtisMDS)),"ASTHMA"])
brayCurtisMDS$Cohort <- factor(paste0(brayCurtisMDS$Asthma, brayCurtisMDS$AgeGroup))
my_color_scheme = c(brewer.pal(7, "BrBG")[c(7,1,5,3)])
# library(viridis)
betaDivPlot4 <- ggplot(data = brayCurtisMDS,
                       aes(x = MDS1, y = MDS2,
                           fill = Cohort,
                           color = Cohort,
                           shape = Asthma)) +
  stat_ellipse(aes(x=MDS1, y=MDS2, color=Cohort, group=Cohort), show.legend = TRUE, type
  ↪  = "t", level = confidenceEllipse, geom = "polygon", alpha = 0.2, size = 1)+
  # geom_point(size = 6, shape = 21, color ="black") +
  geom_point(size = 4.5, color = "black") +
  scale_color_manual(values = my_color_scheme) +
  scale_fill_manual(values = my_color_scheme) +
  scale_shape_manual(values=c(21, 22)) +
  theme_classic() +
  xlab(label = "NMDS1") + ylab("NMDS2") + ggtitle(mytitle) +
  theme(axis.title.x = element_text(size = 21, vjust = -0.9),
        axis.title.y = element_text(size = 21, vjust = 1),
        axis.text.x = element_blank(),
        axis.ticks = element_blank(),
        axis.text.y = element_blank(),
        legend.text = element_text(size = 16),
        legend.title = element_text(size = 16, face = "bold"),
        legend.position = "bottom", legend.box = "horizontal")#,
betaDivPlot4
```

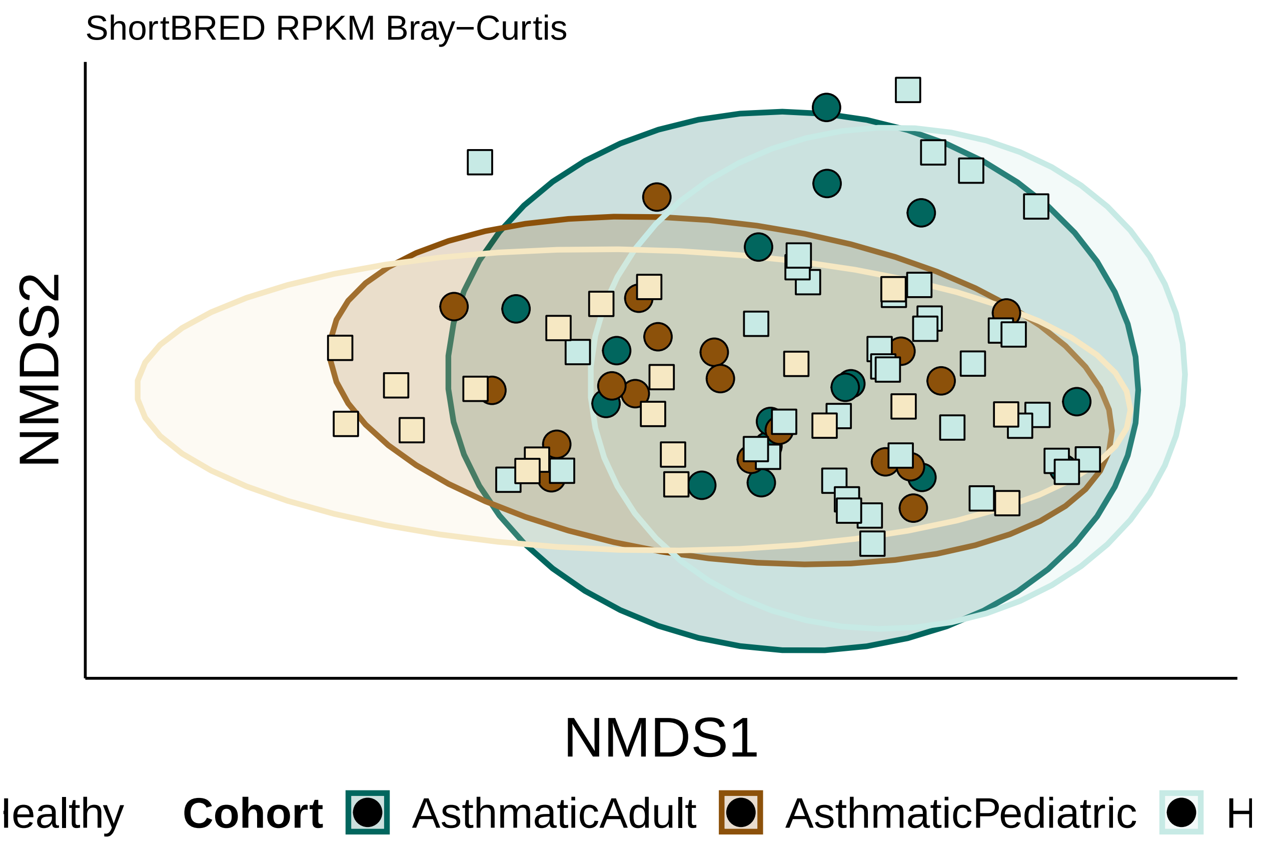


```
# ggsave(plot=betaDivPlot4, filename = paste0(basedir, my_data_table_name,
↪   "_4Group_NMDS_1and2.jpeg"), device = "jpg", width = 5, height = 5, units = "in")
# ggsave(plot=betaDivPlot4, filename = paste0(basedir, my_data_table_name,
↪   "_4Group_NMDS_1and2.pdf"), device = "pdf", width = 5, height = 5, units = "in",
↪   useDingbats = FALSE)
```

#### Figure 3B:

PERMANOVA Bray-Curtis distances barplot

```
rm(list=ls()[ls() %notin% c("%notin%", "makePCA", "makeNMDSplot",
↪   "boxplotByAsthmaAndAge", "boxplotByAsthma", "filter_shortbred_data",
↪   "removeColumnSuffix")])
source("./humann3_210613_functions_ngw.R")

library(ggplot2)
library(ape)
library(stats)
library(vegan)
library(ggfortify)
library(RColorBrewer)

gene_analysis_name = "SB_RPKM_Bray_Curtis"
basedir_humann3="./"
humann3_mf_filtered_perm <- readRDS(file = paste0(basedir_humann3,
↪   "humann3_mf_filtered.rds"))
row.names(humann3_mf_filtered_perm) <- humann3_mf_filtered_perm$SampleName
```

```
df.meta <- humann3_mf_filtered_perm
readDepth <- read.table("~/Library/CloudStorage/Box-Box/Kau
↪  Lab/Results/MARS/FecalMetagenomics/kneaddata/kneaddata_bypassTRF_on_raw_data/merged_kneaddata_table
↪  header = T, sep="\t", stringsAsFactors=F) # added Dec 11 2022 for normalizing alpha
↪  diversity by read depth
row.names(readDepth) <- gsub(readDepth$Sample, pattern = "_L002_R1_001_kneaddata",
↪  replacement = "")
df.shortbred <- readRDS("SB_CARD21_RPKM_filtered.rds")
# Total sum square the data for Bray-curtis transform
df.shortbred.tss <- apply(X = df.shortbred, MARGIN = 2, FUN = function(x){x/sum(x)})
pca.dist =vegdist(t(df.shortbred.tss), method = "bray", binary = FALSE)

df.meta$readDepth <- readDepth[row.names(df.meta), "final.pair1"]*2
adonisAllIndpRaceBinaryPATHWAYS <- adonis(pca.dist ~
↪  readDepth+ASTHMA*AgeGroup+ASTHMA*AbxWithinYr+raceBinary+weightClassBinary+sex+subjectTobaccoUse,
                        data = df.meta,
                        permutations = 100000) # edit oct 2022 to include Asthma:Abx

# testing to see if order matters for significance for abx (it doesn't)
# adonis(pca.dist ~
↪  subjectTobaccoUse+AgeGroup+AbxWithinYr+ASTHMA+raceBinary+weightClassBinary+sex+ASTHMA*AgeGroup,
↪  data = df.meta, permutations = 999) # tobacco seems to matter a bit but there are
↪  only 2 asthmatic smokers and many more healthy ones
# adonis(pca.dist ~ ASTHMA*AgeGroup+ASTHMA*AbxWithinYr, data = df.meta, permutations =
↪  999)
# adonis(pca.dist ~
↪  weightClassBinary+subjectTobaccoUse+sex+ASTHMA*AgeGroup+ASTHMA*AbxWithinYr, data =
↪  df.meta, permutations = 999) # obesity no
# adonis(pca.dist ~
↪  sex+weightClassBinary+subjectTobaccoUse+ASTHMA*AgeGroup+ASTHMA*AbxWithinYr, data =
↪  df.meta, permutations = 999) # no
# adonis(pca.dist ~
↪  subjectTobaccoUse+sex+weightClassBinary+ASTHMA*AgeGroup+ASTHMA*AbxWithinYr, data =
↪  df.meta, permutations = 999) # sex gets to p=0.05 when tobacco is first??
# Although sex and tobacco are sort of on the fence, they go back and forth whereas
↪  asthma stays significant for any of the combinations so I think it still makes sense
↪  to put it first.

# make table
adonisAllIndpRaceBinaryPATHWAYSTable <-
↪  data.frame(adonisAllIndpRaceBinaryPATHWAYS$aov.tab[c("R2","Pr(>F)")])
names(adonisAllIndpRaceBinaryPATHWAYSTable) <- c("R2", "pval")
adonisAllIndpRaceBinaryPATHWAYSTable$features <-
↪  row.names(adonisAllIndpRaceBinaryPATHWAYSTable)
adonisAllIndpRaceBinaryPATHWAYSTable
↪  <-adonisAllIndpRaceBinaryPATHWAYSTable[!adonisAllIndpRaceBinaryPATHWAYSTable$features
↪  %in% c("Residuals","Total"),]
adonisAllIndpRaceBinaryPATHWAYSTable$pval <-
↪  round(adonisAllIndpRaceBinaryPATHWAYSTable$pval, 5)
# adonisAllIndpRaceBinaryPATHWAYSTable$pval <-
↪  -log(adonisAllIndpRaceBinaryPATHWAYSTable$pval)
adonisAllIndpRaceBinaryPATHWAYSTable$R2 <- round(adonisAllIndpRaceBinaryPATHWAYSTable$R2,
↪  3)
```

```
# attr(adonisAllIndpRaceBinaryPATHWAYSTable["ASTHMA",], "label") <- "Asthma Status"
# attr(adonisAllIndpRaceBinaryPATHWAYSTable$R2, "label") <- "R-Squared"
row.names(adonisAllIndpRaceBinaryPATHWAYSTable) <- NULL
# re-order so that pvalues low are at top
adonisAllIndpRaceBinaryPATHWAYSTable <-
↪  adonisAllIndpRaceBinaryPATHWAYSTable[order(adonisAllIndpRaceBinaryPATHWAYSTable$pval,
↪  decreasing = FALSE),]
adonisAllIndpRaceBinaryPATHWAYSTable <- adonisAllIndpRaceBinaryPATHWAYSTable[,c(3,1,2)]

basedir="./"
write.table(adonisAllIndpRaceBinaryPATHWAYSTable, file = paste0(basedir,
↪  gene_analysis_name, "_adonisAllIndpRaceBinaryweightClassBinary100000.tsv"), sep="\t",
↪  row.names = FALSE, col.names = TRUE)

adonisAllIndpRaceBinaryPATHWAYSTable$neglogpval <-
↪  -log(adonisAllIndpRaceBinaryPATHWAYSTable$pval, base = 10)
adonisAllIndpRaceBinaryPATHWAYSTable$features <-
↪  reorder(adonisAllIndpRaceBinaryPATHWAYSTable$features,
↪  adonisAllIndpRaceBinaryPATHWAYSTable$R2)
adonisAllIndpRaceBinaryPATHWAYSTable$features <-
↪  factor(adonisAllIndpRaceBinaryPATHWAYSTable$features, levels=rev(c("readDepth",
↪  "ASTHMA", "AgeGroup", "ASTHMA:AgeGroup", "AbxWithinYr", "ASTHMA:AbxWithinYr",
↪  "raceBinary", "weightClassBinary",  "sex", "subjectTobaccoUse")), labels =
↪  rev(c("Read Depth", "Asthma", "Age", "Asthma:Age", "Recent\nAntibiotics",
↪  "Asthma:Recent\nAntibiotics", "Race","Obesity", "Sex", "Tobacco Use")))

AdonisBarplotForPaperBinaryRacePATHWAYS <- makeAdonisBarplot(adonis_dat =
↪  adonisAllIndpRaceBinaryPATHWAYSTable, y_colname = "neglogpval", features_colname =
↪  "features", fill_colname = "R2", max_fill_value =
↪  max(adonisAllIndpRaceBinaryPATHWAYSTable$R2)*1.1)
print(AdonisBarplotForPaperBinaryRacePATHWAYS)
```

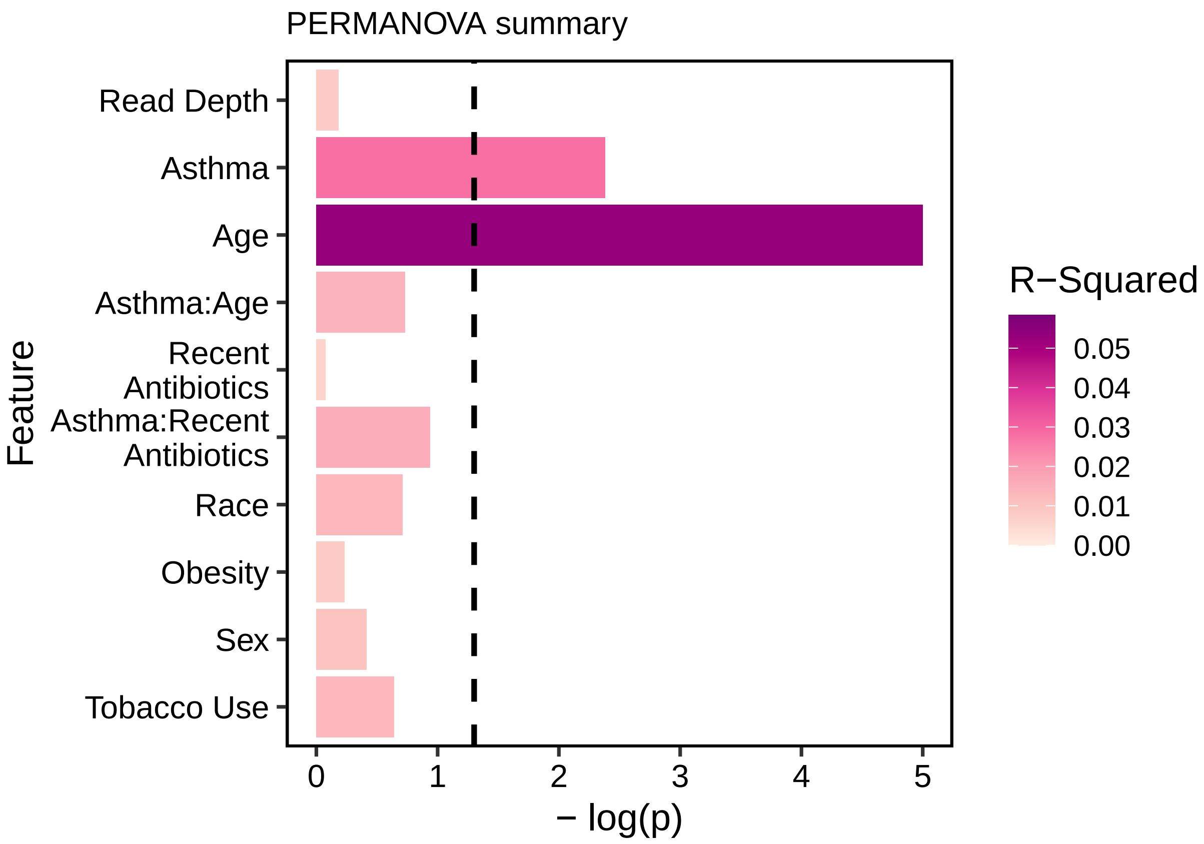


```
# ggsave(AdonisBarplotForPaperBinaryRacePATHWAYS,
#        filename = paste0(basedir, gene_analysis_name,
↪  "_adonisAllIndpRaceBinaryweightClassBinary100000.pdf"),
#        height = 4, width = 5,
#        units = "in",
#        device = "pdf", useDingbats=FALSE)
#
# ggsave(AdonisBarplotForPaperBinaryRacePATHWAYS,
#        filename = paste0(basedir, gene_analysis_name,
↪  "_adonisAllIndpRaceBinaryweightClassBinary100000.jpg"),
#        height = 4, width = 5,
#        units = "in",
#        device = "jpg")
```

#### Figure 2B: Richness (non-zero rpkm)

```
source("./split_violins_ggplot.R")
rm(list=ls()[ls() %notin% c("%notin%", "makePCA", "makeNMDSplot",
↪  "boxplotByAsthmaAndAge", "boxplotByAsthma", "GeomSplitViolin",
↪  "create_quantile_segment_frame", "geom_split_violin", "filter_shortbred_data",
↪  "removeColumnSuffix")])
library(ggplot2)
library(RColorBrewer)
basedir="./"
humann3_mf_filtered_perm <- readRDS(file = "humann3_mf_filtered.rds")
```

```r
row.names(humann3_mf_filtered_perm) <- humann3_mf_filtered_perm$SampleName
df.meta <- humann3_mf_filtered_perm
df.shortbred <- readRDS("SB_CARD21_RPKM_filtered.rds")

#Next, generate a simple table that indicates the presence/absence of antibiotic
↪  resistance genes in each sample.
# Convert our data to presence/absence
df.shortbred.b <- df.shortbred
df.shortbred.b[df.shortbred.b!=0] <- 1
# collect "hits" per sample
hits <- data.frame(colSums(df.shortbred.b))
#hits <- hits[row.names(df.meta),]
hits$ASTHMA <- df.meta[row.names(hits), "ASTHMA"]
hits$AgeGroup <- df.meta[row.names(hits), "AgeGroup"]
hits$MARSID <- df.meta[row.names(hits), "ID"]
hits$AbxWithinYr <- df.meta[row.names(hits), "AbxWithinYr"]
# table(hits$ASTHMA, hits$colSums.df.shortbred.b.)
hits$readDepth <- df.meta[row.names(hits), "readDepth"]
hits$coverage <- df.meta[row.names(hits), "coverage"]

# ANOVA types:
↪  (https://www.r-bloggers.com/2011/03/anova-%E2%80%93-type-iiiiii-ss-explained/)
library(car)
summary(aov(colSums.df.shortbred.b.~ASTHMA*readDepth+ASTHMA*AgeGroup, data=hits)) # int
↪  is 0.9, 0.4 so can do type 32
```

```
##                   Df Sum Sq Mean Sq F value Pr(>F)
## ASTHMA             1    184  183.50   4.786 0.0313 *
## readDepth          1    177  176.67   4.608 0.0345 *
## AgeGroup           1    193  192.58   5.022 0.0275 *
## ASTHMA:readDepth   1      0    0.49   0.013 0.9102
## ASTHMA:AgeGroup    1     30   29.86   0.779 0.3799
## Residuals         89   3413   38.34
## ---
## Signif. codes:  0 '***' 0.001 '**' 0.01 '*' 0.05 '.' 0.1 ' ' 1
```

```r
Anova(lm(colSums.df.shortbred.b. ~ASTHMA*readDepth+ASTHMA*AgeGroup, data = hits),type=2)
```

```
## Anova Table (Type II tests)
##
## Response: colSums.df.shortbred.b.
##                   Sum Sq Df F value   Pr(>F)
## ASTHMA             268.9  1  7.0141 0.009567 **
## readDepth          113.9  1  2.9695 0.088323 .
## AgeGroup           192.5  1  5.0207 0.027536 *
## ASTHMA:readDepth     0.1  1  0.0020 0.964644
## ASTHMA:AgeGroup     29.9  1  0.7787 0.379919
## Residuals         3412.6 89
## ---
## Signif. codes:  0 '***' 0.001 '**' 0.01 '*' 0.05 '.' 0.1 ' ' 1
```

```r
Anova(lm(colSums.df.shortbred.b. ~ASTHMA*coverage+ASTHMA*AgeGroup, data = hits),type=2)
```

```
## Anova Table (Type II tests)
```

```
##
## Response: colSums.df.shortbred.b.
##                    Sum Sq Df F value   Pr(>F)
## ASTHMA              261.1  1  7.3309 0.008126 **
## coverage            328.2  1  9.2159 0.003146 **
## AgeGroup            138.1  1  3.8770 0.052062 .
## ASTHMA:coverage      29.0  1  0.8136 0.369503
## ASTHMA:AgeGroup      32.8  1  0.9199 0.340100
## Residuals          3169.4 89
## ---
## Signif. codes:  0 '***' 0.001 '**' 0.01 '*' 0.05 '.' 0.1 ' ' 1
```

```
summary(glm(colSums.df.shortbred.b. ~ASTHMA*readDepth+ASTHMA*AgeGroup, data = hits))
```

```
##
## Call:
## glm(formula = colSums.df.shortbred.b. ~ ASTHMA * readDepth +
##     ASTHMA * AgeGroup, data = hits)
##
## Deviance Residuals:
##      Min        1Q    Median        3Q       Max
## -10.4054   -3.7740   -0.8938    4.1191   18.2713
##
## Coefficients:
##                                  Estimate Std. Error t value Pr(>|t|)
## (Intercept)                     2.585e+01  2.519e+00  10.261   <2e-16 ***
## ASTHMAHealthy                  -4.717e+00  3.100e+00  -1.522   0.1317
## readDepth                       7.128e-08  7.312e-08   0.975   0.3323
## AgeGroupPediatric              -4.422e+00  2.098e+00  -2.108   0.0379 *
## ASTHMAHealthy:readDepth         4.013e-09  9.029e-08   0.044   0.9646
## ASTHMAHealthy:AgeGroupPediatric  2.402e+00  2.722e+00   0.882   0.3799
## ---
## Signif. codes:  0 '***' 0.001 '**' 0.01 '*' 0.05 '.' 0.1 ' ' 1
##
## (Dispersion parameter for gaussian family taken to be 38.3436)
##
##     Null deviance: 3995.7  on 94  degrees of freedom
## Residual deviance: 3412.6  on 89  degrees of freedom
## AIC: 623.83
##
## Number of Fisher Scoring iterations: 2
```

```
# cor.test(hits$colSums.df.shortbred.b. , hits$readDepth)

# Transpose the binary table
df.shortbred.b <- data.frame(t(df.shortbred.b))
# Add some extra columns to our table
df.shortbred.b$ASTHMA <- df.meta[row.names(df.shortbred.b), "ASTHMA"]
df.shortbred.b$AgeGroup <- df.meta[row.names(df.shortbred.b), "AgeGroup"]
df.shortbred.b$Sample <-  row.names(df.shortbred.b)

# plot CARD richness
hits$value <- hits$colSums.df.shortbred.b.
hits$ASTHMA <- factor(hits$ASTHMA, levels = c("Healthy", "Asthmatic"))
# plot CARD richness - by age and asthma
```

```r
# CARDRPKMDerivedHitsPlot <- boxplotByAsthmaAndAge(mydat = hits)
# plot CARD richness - only asthma groups
hits$AgeGroup <- factor(hits$AgeGroup)
# CARDRPKMDerivedHitsPlotASTHMA <- boxplotByAsthma(mydat = hits)

# VIOLIN PLOTS
library(ggbeeswarm)
library(ggpubr)
CARDrichnessViolin <- ggplot(data = hits, aes(x=ASTHMA, y=value, fill =
↪  interaction(ASTHMA,AgeGroup))) +
  geom_violin(aes(x=ASTHMA, y=value, fill=ASTHMA), draw_quantiles = c(0.5), inherit.aes =
  ↪  FALSE, scale="width", trim=FALSE, alpha=0.8, show.legend = T) + # draw_quantiles =
  ↪  c(0.5), c(0.25, 0.5, 0.75)
  geom_split_violin(draw_quantiles = c(0.5), scale="width", width=0.5, trim=FALSE,
  ↪  alpha=0.8, show.legend = T) +  # add this to draw quantiles: draw_quantiles =
  ↪  c(0.25, 0.5, 0.75)
  theme_classic() +
  scale_fill_manual(name="Cohort",
                    values = c("#01665E", brewer.pal(7, "BrBG")[1], brewer.pal(7,
                    ↪  "BrBG")[3],
                               "#C7EAE5", brewer.pal(7, "BrBG")[1], brewer.pal(7,
                               ↪  "BrBG")[3]),
                    labels = c('Asthmatic','Adult', 'Pediatric',
                               'Healthy', 'Adult', 'Pediatric'))+ # color order:
                               ↪  A,AA,AP,H,HA,HP
  theme(panel.grid.major = element_blank(),
        panel.grid.minor = element_blank(),
        axis.line.x = element_line(size = 0, colour = "black"),
        axis.line.y = element_line(size = 0, colour = "black"),
        axis.line = element_line(size=1, colour = "black"),
        panel.border = element_rect(fill = NA, colour = "black", size = 1),
        panel.background = element_blank(),
        text=element_text(size = 16),
        axis.text.x=element_text(colour="black", size = 12),
        axis.text.y=element_text(colour="black", size = 12),
        axis.title.x=element_blank())+
        # legend.position = "none")+
  ylab("Number of Unique ARGs")
# ggsave(plot = CARDrichnessViolin, filename =
↪  "./sb_CARD_nonzeroRPKMHits_Asthma_doubleviolin.pdf", width = 6, height = 6, units =
↪  "in", useDingbats = FALSE)
# ggsave(filename = "./sb_CARD_nonzeroRPKMHits_Asthma_doubleviolin.jpeg", plot =
↪  CARDrichnessViolin, units = "in", width =6, height = 6, device = "jpg")

hits$AgeAsthma <- factor(paste0(hits$AgeGroup, hits$ASTHMA))
CARDrichnessViolin + ggpubr::stat_compare_means(data=hits, mapping = aes(x=ASTHMA,
↪  y=value), inherit.aes = F, label.y = 55)
```

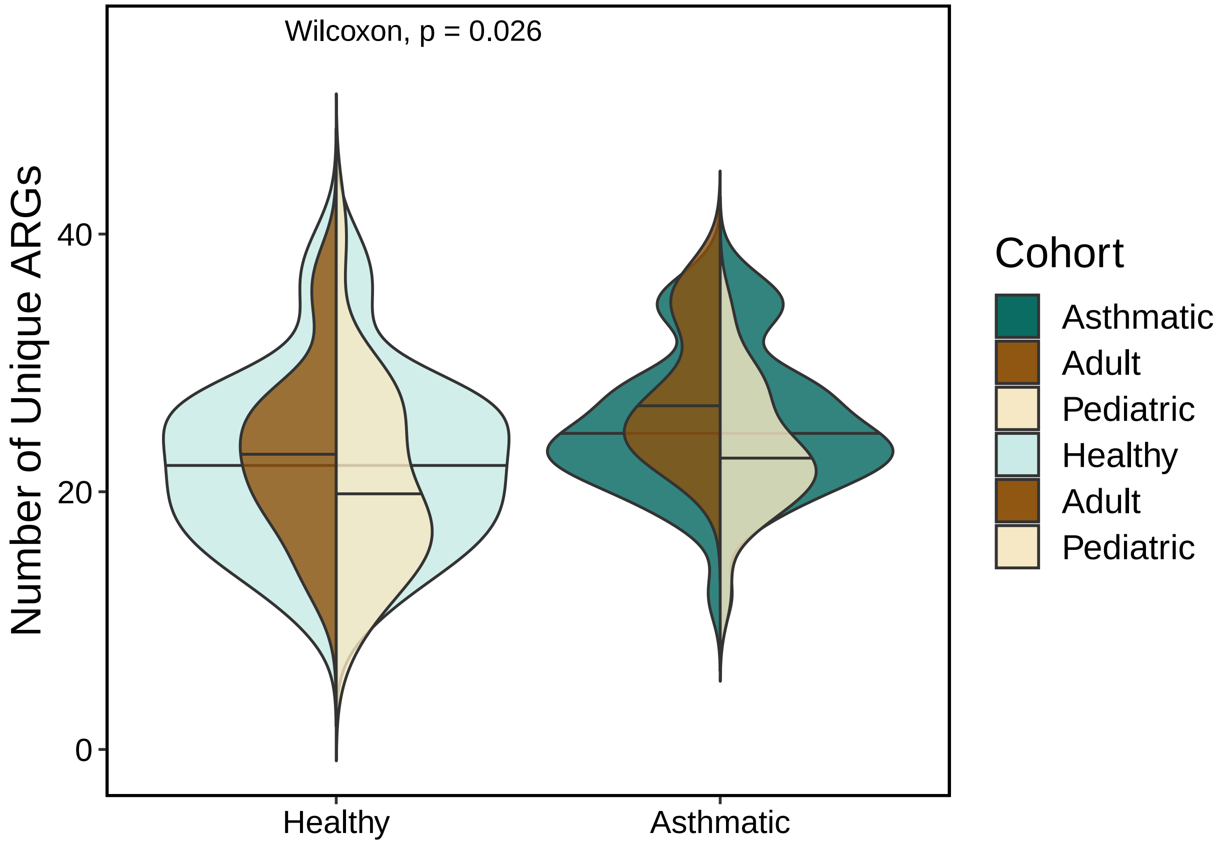


#### Figure 2B: Sum of RPKMs

```
source("/Users/naomiwilson/Library/CloudStorage/Box-Box/Kau
↪  Lab/Results/MARS/FecalMetagenomics/shortbred/CARD2021results/split_violins_ggplot.R")
rm(list=ls()[ls() %notin% c("%notin%", "makePCA", "makeNMDSplot",
↪  "boxplotByAsthmaAndAge", "boxplotByAsthma", "GeomSplitViolin",
↪  "create_quantile_segment_frame", "geom_split_violin", "filter_shortbred_data",
↪  "removeColumnSuffix")])

library(ggplot2)
library(ggbeeswarm)
library(ggpubr)

basedir="./"
humann3_mf_filtered_perm <- readRDS(file = paste0(basedir, "humann3_mf_filtered.rds"))
row.names(humann3_mf_filtered_perm) <- humann3_mf_filtered_perm$SampleName
df.meta <- humann3_mf_filtered_perm
df.shortbred <- readRDS("SB_CARD21_RPKM_filtered.rds")
# plot sum of RPKMs
df.CARD <- df.shortbred[,row.names(df.meta)]
df.rpkm.sum <- as.data.frame(colSums(df.CARD))
df.rpkm.sum$ASTHMA <- df.meta[row.names(df.rpkm.sum), "ASTHMA"]
df.rpkm.sum$ASTHMA <- factor(df.rpkm.sum$ASTHMA, levels=c("Healthy", "Asthmatic"))
df.rpkm.sum$MARSID <- df.meta[row.names(df.rpkm.sum), "MARSID"]
df.rpkm.sum$AgeGroup <- df.meta[row.names(df.rpkm.sum), "AgeGroup"]
```

```r
df.rpkm.sum$value <- df.rpkm.sum$`colSums(df.CARD)`
# hits$AbxWithinYr <- df.meta[row.names(hits), "AbxWithinYr"]
# table(hits$ASTHMA, hits$colSums.df.shortbred.b.)
df.rpkm.sum$readDepth <- df.meta[row.names(df.rpkm.sum), "readDepth"]
df.rpkm.sum$coverage <- df.meta[row.names(df.rpkm.sum), "coverage"]

summedRPKMplot <- boxplotByAsthma(mydat = df.rpkm.sum, my_ylab = "CARD ARG Load (Summed
↪  RPKM)")
summedRPKMplotByAge <- boxplotByAsthmaAndAge(mydat = df.rpkm.sum, my_ylab = "CARD ARG
↪  Load (Summed RPKM)")
# print(summedRPKMplot)
# print(summedRPKMplotByAge)

# ANOVA types:
↪  (https://www.r-bloggers.com/2011/03/anova-%E2%80%93-type-iiiiii-ss-explained/)
library(car)
# internet says if interaction is not significant in Type 1, then you should run a type 2
↪  ANOVA
summary(aov(value~ASTHMA*readDepth + ASTHMA*AgeGroup , data=df.rpkm.sum)) # type 1: int
↪  is 0.3
```

```
##                  Df  Sum Sq Mean Sq F value   Pr(>F)
## ASTHMA            1     943     943   0.023 0.880160
## readDepth         1  678551  678551  16.454 0.000107 ***
## AgeGroup          1  819914  819914  19.882  2.4e-05 ***
## ASTHMA:readDepth  1   48697   48697   1.181 0.280118
## ASTHMA:AgeGroup   1   67885   67885   1.646 0.202820
## Residuals        89 3670314   41239
## ---
## Signif. codes:  0 '***' 0.001 '**' 0.01 '*' 0.05 '.' 0.1 ' ' 1
```

```r
Anova(lm(value ~ ASTHMA*readDepth + ASTHMA*AgeGroup, data = df.rpkm.sum),type=2) # type 2
```

```
## Anova Table (Type II tests)
##
## Response: value
##                   Sum Sq Df F value    Pr(>F)
## ASTHMA             28772  1  0.6977  0.405800
## readDepth         423301  1 10.2645  0.001883 **
## AgeGroup          821255  1 19.9143 2.362e-05 ***
## ASTHMA:readDepth   69285  1  1.6801  0.198267
## ASTHMA:AgeGroup    67885  1  1.6461  0.202820
## Residuals        3670314 89
## ---
## Signif. codes:  0 '***' 0.001 '**' 0.01 '*' 0.05 '.' 0.1 ' ' 1
```

```r
# Anova(lm(value ~ASTHMA*AgeGroup, data=df.rpkm.sum, contrasts=list(topic=contr.sum,
↪  sys=contr.sum)), type=3) # type 3
Anova(lm(value ~ ASTHMA*coverage + ASTHMA*AgeGroup, data = df.rpkm.sum),type=2) # type 2
```

```
## Anova Table (Type II tests)
##
## Response: value
##                  Sum Sq Df F value    Pr(>F)
```

```
## ASTHMA             25097  1  0.5952  0.442477
## coverage          407143  1  9.6550  0.002535 **
## AgeGroup          739056  1 17.5260 6.644e-05 ***
## ASTHMA:coverage     2699  1  0.0640  0.800845
## ASTHMA:AgeGroup    32928  1  0.7809  0.379258
## Residuals       3753057 89
## ---
## Signif. codes:  0 '***' 0.001 '**' 0.01 '*' 0.05 '.' 0.1 ' ' 1
```

```
# VIOLIN PLOTS
CARDLoadViolin <- ggplot(data = df.rpkm.sum, aes(x=ASTHMA, y=value, fill =
↪  interaction(ASTHMA,AgeGroup))) +
  geom_violin(aes(x=ASTHMA, y=value, fill=ASTHMA), draw_quantiles = c(0.5), inherit.aes =
  ↪  FALSE, scale="width", trim=FALSE, alpha=0.8, show.legend = T) + # draw_quantiles =
  ↪  c(0.5), c(0.25, 0.5, 0.75)
  geom_split_violin(draw_quantiles = c(0.5), scale="width", width=0.5, trim=FALSE,
  ↪  alpha=0.8, show.legend = T) +  # add this to draw quantiles: draw_quantiles =
  ↪  c(0.25, 0.5, 0.75)
  theme_classic() +
  scale_fill_manual(name="Cohort",
                    values = c("#01665E", brewer.pal(7, "BrBG")[1], brewer.pal(7,
                    ↪  "BrBG")[3],
                               "#C7EAE5", brewer.pal(7, "BrBG")[1], brewer.pal(7,
                               ↪  "BrBG")[3]),
                    labels = c('Asthmatic','Adult', 'Pediatric',
                               'Healthy', 'Adult', 'Pediatric'))+ # color order:
                               ↪  A,AA,AP,H,HA,HP
  theme(panel.grid.major = element_blank(),
        panel.grid.minor = element_blank(),
        axis.line.x = element_line(size = 0, colour = "black"),
        axis.line.y = element_line(size = 0, colour = "black"),
        axis.line = element_line(size=1, colour = "black"),
        panel.border = element_rect(fill = NA, colour = "black", size = 1),
        panel.background = element_blank(),
        text=element_text(size = 16),
        axis.text.x=element_text(colour="black", size = 12),
        axis.text.y=element_text(colour="black", size = 12),
        axis.title.x=element_blank())+
        # legend.position = "none")+
  ylab("CARD ARG Load (summed RPKM)")
CARDLoadViolin + ggpubr::stat_compare_means(mapping = aes(x=ASTHMA, y=value), inherit.aes
↪  = F, label.y = 1800)
```

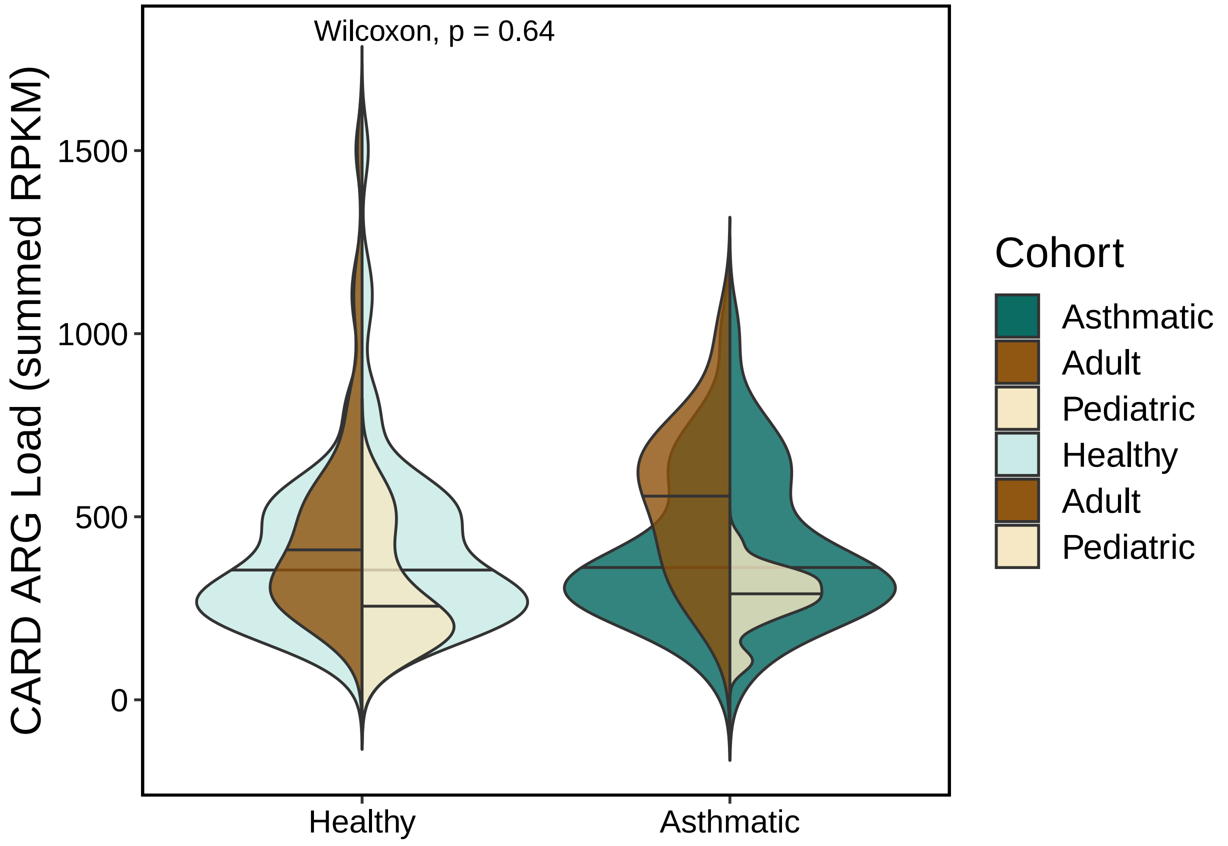


```
# ggsave(plot = CARDLoadViolin, filename =
↪  "./sb_CARD_summedRPKM_Asthma_doubleviolin.pdf", width = 6, height = 6, units = "in",
↪  useDingbats = FALSE)
# ggsave(filename = "./sb_CARD_summedRPKM_Asthma_doubleviolin.jpeg", plot =
↪  CARDLoadViolin, units = "in", width =6, height = 6, device = "jpg")
```

#### Make mapping file with functional categories (AMR family, drug class, resistance mechanism)

```
rm(list=ls()[ls() %notin% c("%notin%", "makePCA", "makeNMDSplot",
↪  "boxplotByAsthmaAndAge", "boxplotByAsthma", "filter_shortbred_data",
↪  "removeColumnSuffix")])
library(ggplot2)
library(RColorBrewer)

basedir="./"
humann3_mf_filtered_perm <- readRDS(file = paste0(basedir, "humann3_mf_filtered.rds"))
row.names(humann3_mf_filtered_perm) <- humann3_mf_filtered_perm$SampleName
df.meta <- humann3_mf_filtered_perm

# df.shortbred.U <- read.table('MARS_SB_CARD21_RPKM.tsv', row.names=1, header=T)  # 888
↪  104
df.shortbred <- readRDS('SB_CARD21_RPKM_filtered.rds')  # 71 95
aro.df = read.csv("~/Library/CloudStorage/Box-Box/Kau
↪  Lab/Results/MARS/FecalMetagenomics/shortbred/databases/CARD/card-data/aro_index.tsv",
↪  sep = "\t", header = TRUE) #[1] 3326   12
```

```r
aro.df$AccessionUnderscore = gsub(aro.df$ARO.Accession, pattern = ":", replacement = "_")

aro_hits = c()
for (i in 1:length(row.names(df.shortbred))) {
  aro_hits = c(aro_hits, strsplit(row.names(df.shortbred), split = "\\|")[[i]][3])
}
# aro_hits %in% aro.df$AccessionUnderscore
df.shortbred$fullName = row.names(df.shortbred)
row.names(df.shortbred[!(aro_hits %in% aro.df$AccessionUnderscore),]) # 9 (now 0) missing
↪  from database download
```

```
## character(0)
```

```r
# let's move on with the ones that *can be annotated:
df.shortbred <- df.shortbred[(aro_hits %in% aro.df$AccessionUnderscore),] # 162 (71)
aro_hits2 = c()
for (i in 1:length(row.names(df.shortbred))) {
  aro_hits2 = c(aro_hits2, strsplit(row.names(df.shortbred), split = "\\|")[[i]][3])
}
all(aro_hits2 %in% aro.df$AccessionUnderscore)
```

```
## [1] TRUE
```

```r
df.shortbred$AccessionNo = aro_hits2

subset = aro.df[(aro.df$AccessionUnderscore %in% df.shortbred$AccessionNo),]
row.names(subset) = subset$AccessionUnderscore
# subset[df.shortbred$AccessionNo,]
df.shortbred$DrugClass = subset[df.shortbred$AccessionNo, "Drug.Class"]
df.shortbred$ResistanceMechanism = subset[df.shortbred$AccessionNo,
↪  "Resistance.Mechanism"]
df.shortbred$AMRGeneFamily = subset[df.shortbred$AccessionNo, "AMR.Gene.Family"]

df.shortbred$DrugClassBins <- as.character(df.shortbred$DrugClass)
df.shortbred$DrugClassBins[grep(";", (df.shortbred$DrugClass))] <- "Multiple"
#
↪  barplot(table(df.shortbred[,"DrugClassBins"])[order(table(df.shortbred[,"DrugClassBins"]))],
↪  horiz = FALSE, las=2, xlab = "Antibiotic")

df.shortbred$ResistanceMechanismBins <- as.character(df.shortbred$ResistanceMechanism)
df.shortbred$ResistanceMechanismBins[grep(";", (df.shortbred$ResistanceMechanismBins))]
↪  <- "Multiple"
#
↪  barplot(table(df.shortbred[,"ResistanceMechanismBins"])[order(table(df.shortbred[,"ResistanceMechan
↪  horiz = FALSE, las=2, xlab = "Antibiotic")

df.shortbred$ResistanceMechanismBins <- as.character(df.shortbred$ResistanceMechanism)
df.shortbred$ResistanceMechanismBins[grep(";", (df.shortbred$ResistanceMechanismBins))]
↪  <- "Multiple"
#
↪  barplot(table(df.shortbred[,"ResistanceMechanismBins"])[order(table(df.shortbred[,"ResistanceMechan
↪  horiz = FALSE, las=2, xlab = "Antibiotic")

write.csv(x = df.shortbred, file =
↪  "SB_CARD21_with_AMRFam_ARO_Drug_Mechanism_filtered.csv", row.names = TRUE)
```

```
saveRDS(object = df.shortbred, file =
↪  "SB_CARD21_with_AMRFam_ARO_Drug_Mechanism_filtered.rds")
```

### Figure 2C:

Stacked barplots

```
rm(list=ls()[ls() %notin% c("%notin%", "makePCA", "makeNMDSplot",
↪  "boxplotByAsthmaAndAge", "boxplotByAsthma", "filter_shortbred_data",
↪  "removeColumnSuffix")])
source("./humann3_210613_functions_ngw.R")
# setwd("/Users/naomiwilson/Library/CloudStorage/Box-Box/Kau
↪  Lab/Results/MARS/FecalMetagenomics/shortbred/CARD2021results")

# Make stacked barplots of "richness" on average per cohort. Meaning, each stack height
↪  represents how many unique ARGs belong to each designated subgroup on average within
↪  asthma or healthy cohorts. Total height should equal the mean of the richness.

library(tidyverse)
library(ggpubr)
library(ggplot2)
library(ggsignif)
library(RColorBrewer)
library(ggbeeswarm)

basedir="./"
humann3_mf_filtered_perm <- readRDS(file = paste0(basedir, "humann3_mf_filtered.rds"))
row.names(humann3_mf_filtered_perm) <- humann3_mf_filtered_perm$SampleName
df.meta <- humann3_mf_filtered_perm

# make richness table per category.
listofcategories <- c("DrugClassBins", "AMRGeneFamily", "ResistanceMechanismBins")
listofcohorts <- c("Asthmatic", "Healthy")
orderofcohorts <- c("Healthy", "Asthmatic")
mythreshold <- 0.0
viridis_scale_name <- "plasma" #turbo for rainbow
library(viridis)
num_to_plot <- 10

df.shortbred <- readRDS(file = "SB_CARD21_with_AMRFam_ARO_Drug_Mechanism_filtered.rds")
df.shortbred <- df.shortbred[order(rowSums(df.shortbred[,1:95]), decreasing = TRUE),] #
↪  re-order by overall abundance
# make richness dataframe
df.shortbred.bin <- data.frame(ifelse(df.shortbred[,row.names(df.meta)] !=0, 1, 0)) #
↪  make a binary matrix to count detected unique ARGs ("richness")

for (category in listofcategories){
  # i=2
  # category = listofcategories[i]
    df.shortbred[,category] <- factor(df.shortbred[,category],
↪  levels=unique(df.shortbred[,category])) # make sure these are factors
  df.shortbred.bin$ARGcategory <- df.shortbred[row.names(df.shortbred.bin), category]
  df.shortbred.bin.sum <- aggregate(. ~ ARGcategory, data = df.shortbred.bin, FUN = sum)
```

```
  names(df.shortbred.bin.sum) <- gsub(x = names(df.shortbred.bin.sum), pattern =
   ↪  "\\.",replacement = "-")
  row.names(df.shortbred.bin.sum) <- df.shortbred.bin.sum$ARGcategory
  df.shortbred.bin.sum$ARGcategory <- NULL
  # myOrder <-
   ↪  row.names(df.shortbred.bin.sum)[order(as.numeric(rowSums(df.shortbred.bin.sum)),
   ↪  decreasing = FALSE)]
  dat <- as.data.frame(t(mean_by_group_pwys(my_data= df.shortbred.bin.sum, group_by_me =
↪   "ASTHMA", sample_mf = df.meta)))
  saveRDS(dat, paste0("meanRichness_", category, ".rds"))
  print(colSums(dat))
  # only plot the top 10 if there are more than 10 groupings
  print(paste0("There are ", dim(dat)[1], " subgroups."))
  if(dim(dat)[1] > num_to_plot){
    rowOrder <- row.names(dat[order(rowMeans(dat), decreasing = TRUE),])
    dat <- dat[rowOrder,]
    dat.short <- dat[rowOrder[1:num_to_plot],]
    dat.short <- rbind(dat.short, c(sum(dat[(num_to_plot+1):dim(dat)[1], 1]),
↪   sum(dat[(num_to_plot+1):dim(dat)[1], 2])))
    row.names(dat.short)[(num_to_plot+1)] <- "Other"
    dat.short$ARGcategory <- row.names(dat.short)
    rowOrderShort <- c(rev(rowOrder[1:num_to_plot]), "Other")
    dat.melted <- reshape2::melt(dat.short)
    dat.melted$ARGcategory <- factor(dat.melted$ARGcategory, levels = rowOrderShort)
    dat.melted$variable <- factor(dat.melted$variable, levels = orderofcohorts)
  } else {
    dat <- as.data.frame(t(mean_by_group_pwys(my_data= df.shortbred.bin.sum, group_by_me
↪   = "ASTHMA", sample_mf = df.meta)))
    rowOrder <- rev(row.names(dat[order(rowMeans(dat), decreasing = TRUE),]))
    dat$ARGcategory <- row.names(dat)
    dat.melted <- reshape2::melt(dat)
    dat.melted$ARGcategory <- factor(dat.melted$ARGcategory, levels = rowOrder)
    dat.melted$variable <- factor(dat.melted$variable, levels = orderofcohorts)
}

library(viridis)
library(RColorBrewer)
myPlot <- ggplot(dat.melted, aes(fill=ARGcategory, x = variable, y = value)) +
  geom_bar(stat="identity", position = "stack", width = 0.8) +
  theme_classic() +
  # facet_grid( ~ PWYshort)+#, scales = "free_x", space = "free_x") +
  # scale_y_continuous(limits=c(0,1), expand = c(0,0))  +
  scale_y_continuous(expand = c(0, 0))+
  xlab(label = "Cohort") + ylab("Average Richness (number of unique ARGs)") +
  scale_fill_viridis(option = "turbo", discrete = T, begin = 0, end = 1, name=category,
                     labels = function(x) str_wrap(x, width = 30)) + # rainbow
  ggtitle(label = "Richness") +
  # scale_fill_manual(values = brewer.pal(7, "BrBG"))+
  theme(axis.title = element_text(size=12, color = "black"),
        axis.text.x = element_text(size=12, color = "black", angle=45, vjust=1, hjust=1),
        axis.text.y = element_text(size=12, color = "black"),
        legend.title = element_text(face = "bold", size = 14),
        legend.text = element_text(size = 14, color = "black"), #face = "italic",
```

```
        legend.key.size = unit(x = 0.25, units = "in")) #+
  # guides(fill = guide_legend(nrow = 2))
print(myPlot)
# ggsave(plot = myPlot, filename = paste0(category, "_",
↪   "averageRichnessByAsthma_Stacked_Barplot", ".jpg"), width = 6, height = 6, units =
↪   "in", device = "jpg")
# ggsave(plot = myPlot, filename = paste0(category, "_",
↪   "averageRichnessByAsthma_Stacked_Barplot", ".pdf"), width = 6, height = 6, units =
↪   "in", device = "pdf", useDingbats=F)
}
```

```
## Asthmatic   Healthy
##  25.30556  22.44068
## [1] "There are 15 subgroups."
```

```
## Using ARGcategory as id variables
```

```
## Asthmatic   Healthy
##  25.30556  22.44068
## [1] "There are 32 subgroups."
```

```
## Using ARGcategory as id variables
```

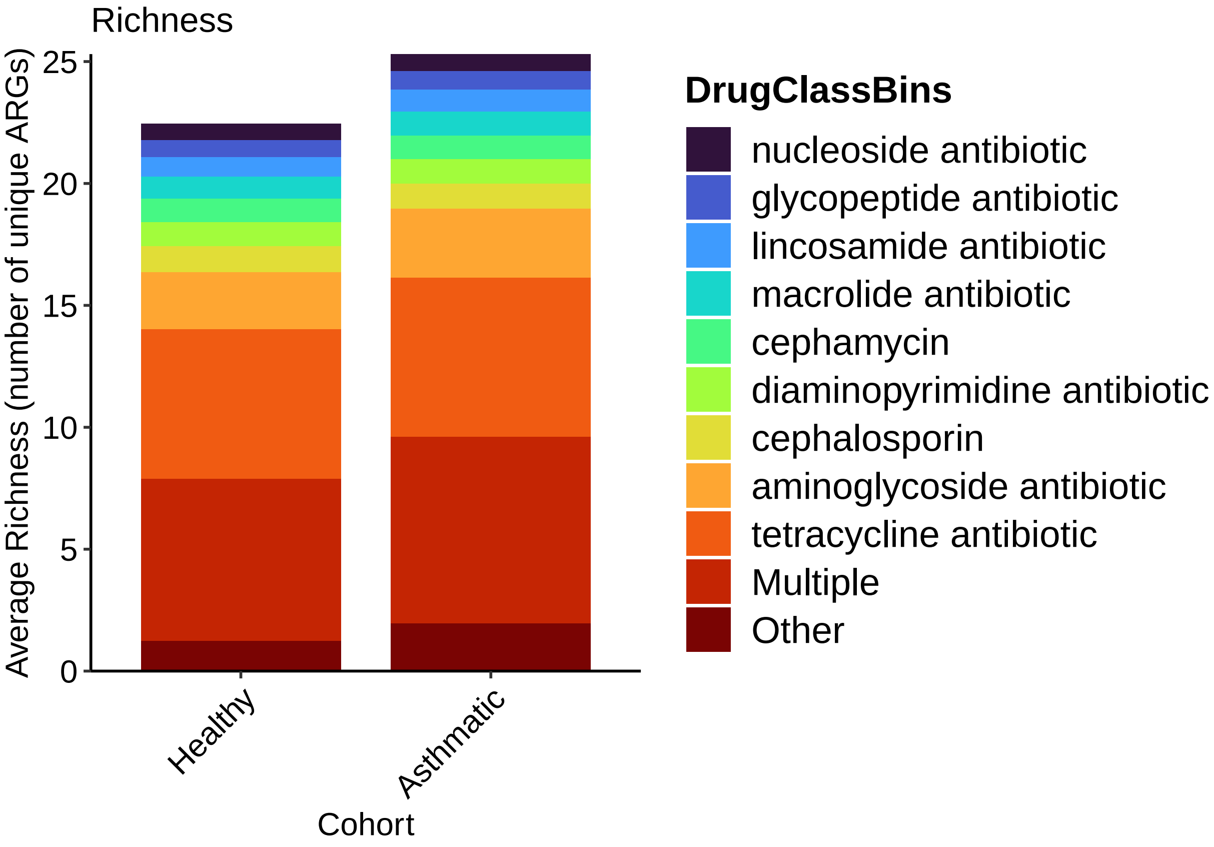


```
## Asthmatic   Healthy
##  25.30556  22.44068
## [1] "There are 6 subgroups."
```

```
## Using ARGcategory as id variables
```

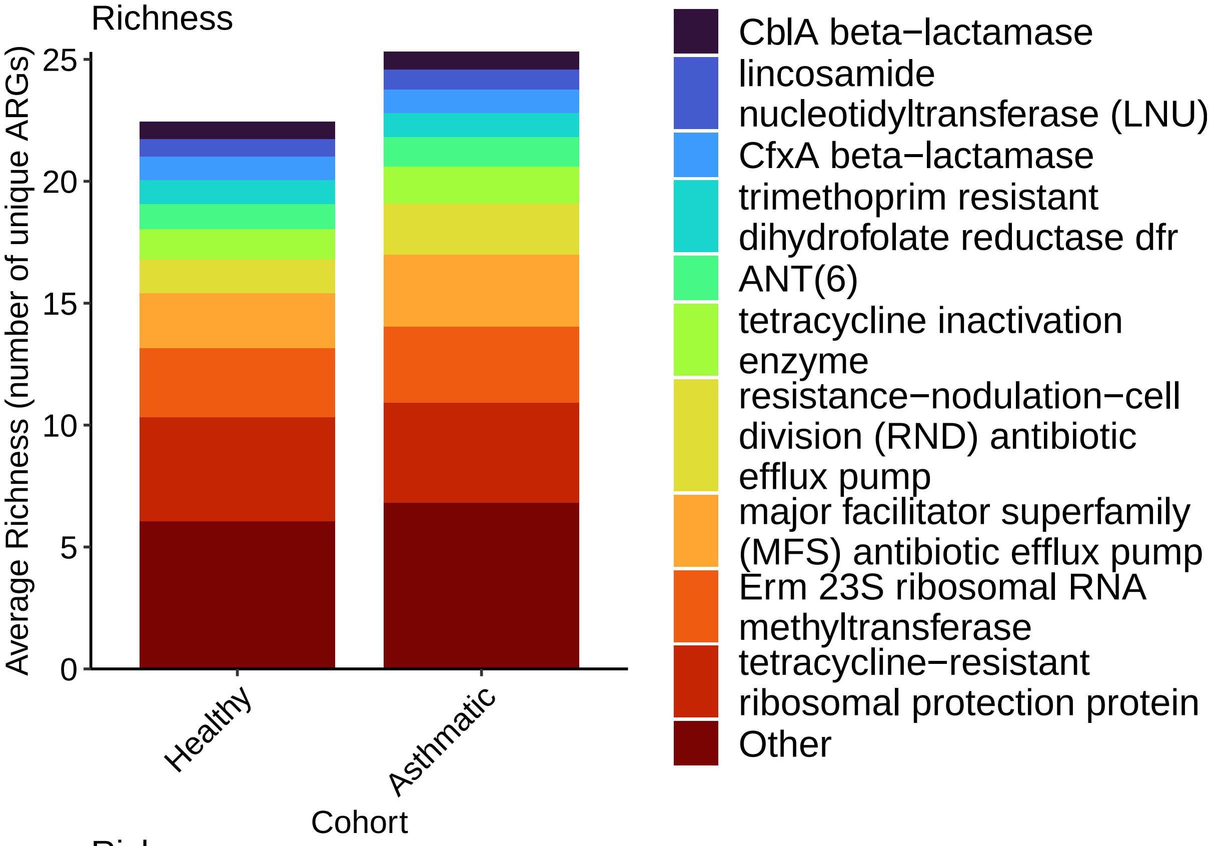
Richness
Average Richness (number of unique ARGs)
0
5
10
15
20
25
Healthy
Asthmatic
Cohort
CblA beta−lactamase
lincosamide nucleotidyltransferase (LNU)
CfxA beta−lactamase
trimethoprim resistant dihydrofolate reductase dfr
ANT(6)
tetracycline inactivation enzyme
resistance−nodulation−cell division (RND) antibiotic efflux pump
major facilitator superfamily (MFS) antibiotic efflux pump
Erm 23S ribosomal RNA methyltransferase
tetracycline−resistant ribosomal protection protein
Other

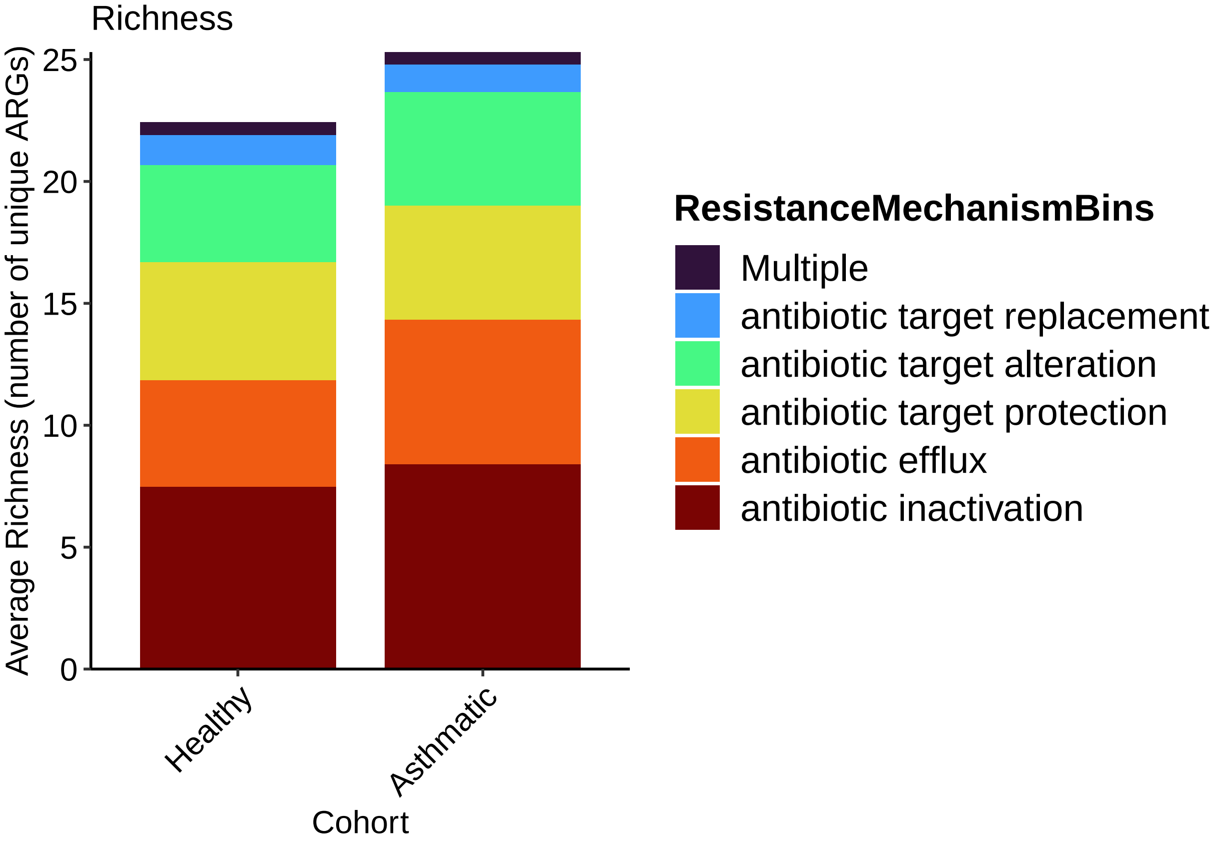
Richness
Average Richness (number of unique ARGs)
0
5
10
15
20
25
Healthy
Asthmatic
Cohort
ResistanceMechanismBins
Multiple
antibiotic target replacement
antibiotic target alteration
antibiotic target protection
antibiotic efflux
antibiotic inactivation

#### Figure 4A:

Get abundance means by asthma

```
rm(list=ls()[ls() %notin% c("%notin%", "makePCA", "makeNMDSplot",
↪  "boxplotByAsthmaAndAge", "boxplotByAsthma", "filter_shortbred_data",
↪  "removeColumnSuffix")])
library(ggplot2)
library(RColorBrewer)
library(ggpubr)
library(stringr)
library(tidyr)
library(dplyr)
library(reshape2)

source("./humann3_210613_functions_ngw.R")

basedir="./"
humann3_mf_filtered_perm <- readRDS(file = "humann3_mf_filtered.rds")
row.names(humann3_mf_filtered_perm) <- humann3_mf_filtered_perm$SampleName
df.meta <- humann3_mf_filtered_perm

df.shortbred <- readRDS(file = "SB_CARD21_with_AMRFam_ARO_Drug_Mechanism_filtered.rds")

group_by_me = "ASTHMA"
df.shortbred.plot <- as.data.frame(t(df.shortbred[,row.names(df.meta)]))
df.shortbred.plot$Group = df.meta[row.names(df.shortbred.plot), group_by_me]

# tibble is supposed to be fast:
my.tbl.tmp <- tibble::as_tibble(df.shortbred.plot)
my.tbl.tmp.means <- my.tbl.tmp %>%
  group_by(Group) %>%
  summarise_all("mean")
# make it a data frame again
my.df.tmp.means <- as.data.frame(my.tbl.tmp.means)
row.names(my.df.tmp.means) <- my.df.tmp.means$Group
my.df.tmp.means$Group = NULL
# my.df.tmp.means

saveRDS(my.df.tmp.means, paste0("./CARD21_RPKM_means_by_", group_by_me, ".rds"))
```

Negative binomial tests

```
rm(list=ls()[ls() %notin% c("%notin%", "makePCA", "makeNMDSplot",
↪  "boxplotByAsthmaAndAge", "boxplotByAsthma", "filter_shortbred_data",
↪  "removeColumnSuffix")])
library(ggplot2)
library(RColorBrewer)
library(ggpubr)
library(stringr)
library(tidyr)
library(dplyr)
library(reshape2)
library(tidyverse)
library(rstatix)
```

```
##
## Attaching package: 'rstatix'

## The following objects are masked from 'package:plyr':
##
##     desc, mutate

## The following object is masked from 'package:stats':
##
##     filter
```

```
group_by_me = "ASTHMA"
my_pair_test = "nb_test"
my_padjust_method="fdr"

basedir="./"
humann3_mf_filtered_perm <- readRDS(file = paste0(basedir, "humann3_mf_filtered.rds"))
row.names(humann3_mf_filtered_perm) <- humann3_mf_filtered_perm$SampleName
df.meta <- humann3_mf_filtered_perm
df.shortbred <- readRDS(file = "./SB_CARD21_with_AMRFam_ARO_Drug_Mechanism_filtered.rds")
sum(vegan::specnumber(t(df.shortbred[,1:dim(df.meta)[1]])) > 71*.1) # all samples have a
↪  good number of genes
```

```
## [1] 95
```

```
pseudocount =
↪  min(df.shortbred[,1:dim(df.meta)[1]][df.shortbred[,1:dim(df.meta)[1]]>0])/20
df.shortbred <- df.shortbred[,1:dim(df.meta)[1]]+pseudocount # add a pseudocount that is
↪  equal to the smallest value divided by 20
my.df.tmp.means <- readRDS(paste0("./CARD21_RPKM_means_by_", group_by_me, ".rds"))

mytest <- "negative_binomial"
mycorrection <- "fdr"

sample_mf = df.meta
features_to_drop = c()
my_pair_test = mytest
my_padjust_method = "fdr" #choose from p.adjust.methods

# choose which groups you want to use:
#df.shortbred$AMRGeneFamily
df.shortbred.t <- data.frame(t(df.shortbred[,row.names(df.meta)]))
df.shortbred.t$ASTHMA <- df.meta$ASTHMA

# Transform the data into long format
# Put all variables in the same column except `Species`, the grouping variable
mydata.long <- df.shortbred.t %>%
  pivot_longer(-ASTHMA, names_to = "variables", values_to = "value")
mydata.long %>% sample_n(6)
```

```
## # A tibble: 6 x 3
##   ASTHMA    variables                                                    value
##   <chr>     <chr>                                                        <dbl>
## 1 Healthy   gb.BAA16344_1.ARO_3000491.acrD__Escherichia_coli_str__K.12_~ 1.48e-3
## 2 Healthy   gb.AAC36915_1.ARO_3000593.ErmQ__Clostridium_perfringens_     1.48e-3
## 3 Asthmatic gb.AAY32951_1.ARO_3002837.lnuC__Streptococcus_agalactiae_    5.21e+1
```

```
## 4 Asthmatic gb.CBH51823_1.ARO_3000556.tet44__Campylobacter_fetus_subsp_~ 5.62e+0
## 5 Asthmatic gb.CAA10975_1.ARO_3000194.tetW__Butyrivibrio_fibrisolvens_   1.39e+1
## 6 Healthy   gb.BAE77933_1.ARO_3000518.CRP__Escherichia_coli_str__K.12_s~ 1.48e-3
```

```
min(mydata.long$value[mydata.long$value>0])
```

```
## [1] 0.001478695
```

```
# make integer version
mydata.long.int <- mydata.long
mydata.long.int$value <-
↪  round(mydata.long$value*(1/min(mydata.long$value[mydata.long$value>0]))) #
sum(mydata.long.int$value<1 & mydata.long.int$value>0)/length(mydata.long.int$value)
```

```
## [1] 0
```

```
mydata.long.int$ASTHMA <- factor(mydata.long.int$ASTHMA)
sum(mydata.long$value<1 & mydata.long.int$value>0)/length(mydata.long$value)
```

```
## [1] 0.7552261
```

```
library(MASS)
```

```
##
## Attaching package: 'MASS'

## The following object is masked from 'package:rstatix':
##
##     select

## The following object is masked from 'package:dplyr':
##
##     select
```

```
stat.test <- mydata.long.int %>%
  group_by(variables) %>%
  mutate(p = as.numeric(coef(summary(MASS::glm.nb(value~ASTHMA)))[2,"Pr(>|z|)"]))
# since this repeated the same value over and over and I'm a dplyr noob, i just want one
↪  instance of the test for each ARG. Thus the janky code below
rows_list_dplyr <- apply( stat.test[ , c(2, 4) ] , 1 , paste , collapse = "-" )
length(unique(rows_list_dplyr)) ==  71
```

```
## [1] TRUE
```

```
sum(!duplicated(rows_list_dplyr)) == 71
```

```
## [1] TRUE
```

```
stat.nb <- stat.test[!duplicated(rows_list_dplyr), c(1,2,4)]
stat.nb$padj <- p.adjust(stat.nb$p, method = my_padjust_method)

# check some of these
as.numeric(coef(summary(MASS::glm.nb(value~ASTHMA, data =
↪  mydata.long.int[mydata.long.int$variables ==
↪  "gb.AAA20116_1.ARO_3000180.tetA.P.__Clostridium_perfringens_",])))[2,"Pr(>|z|)"]) ==
↪  stat.nb[stat.nb$variables ==
↪  "gb.AAA20116_1.ARO_3000180.tetA.P.__Clostridium_perfringens_","p"]
```

```
##          p
## [1,] TRUE
```

```
as.numeric(coef(summary(MASS::glm.nb(value~ASTHMA, data =
↪  mydata.long.int[mydata.long.int$variables ==
↪  "gb.AAA27471_1.ARO_3000205.tetX__Bacteroides_fragilis_",])))[2,"Pr(>|z|)"]) ==
↪  stat.nb[stat.nb$variables ==
↪  "gb.AAA27471_1.ARO_3000205.tetX__Bacteroides_fragilis_","p"]
```

```
##          p
## [1,] TRUE
```

```
incidence <- data.frame(rowSums(df.shortbred[, 1:dim(df.meta)[1]] > pseudocount))
rownames(incidence) <- gsub(rownames(incidence), pattern = "[|]", replacement = ".")
rownames(incidence) <- gsub(rownames(incidence), pattern = "[(|)|-]", replacement = ".")
stat.nb$SIG <- ifelse(stat.nb$padj<0.05, "**", ifelse(stat.nb$padj<0.2, "*", "n.s."))
stat.nb$incidence <- incidence[as.character(stat.nb$variables),]
variable.order <- order(stat.nb$padj, decreasing = FALSE)
stat.nb <- stat.nb[variable.order,]

saveRDS(stat.nb, file = paste0("ARGwPseudocount", "_Asthma_", mytest, "_", mycorrection,
↪  ".rds"))
write.table(stat.nb, file = paste0("ARGwPseudocount", "_Asthma_", mytest, "_",
↪  mycorrection, ".tsv"), sep = "\t")

# add wilcoxons
stat.wilcox <- mydata.long %>%
      group_by(variables) %>%
      wilcox_test(value ~ ASTHMA) %>%  # t_test or wilcox_test
      adjust_pvalue(method = my_padjust_method) %>%
      add_significance(cutpoints = c(1e-04, 0.001, 0.01, 0.05, 0.2, 1), symbols =
       ↪  c("****", "***", "**", "*", "ns"))
# stat.wilcox[variable.order.names, "variables"]
stat.nb.wilcox <- data.frame(stat.nb)
stat.wilcox <- data.frame(stat.wilcox)
row.names(stat.wilcox) <- stat.wilcox$variables
stat.nb.wilcox$p.wilcoxon = stat.wilcox[as.character(stat.nb$variables), "p"]
stat.nb.wilcox$padj.wilcoxon = stat.wilcox[as.character(stat.nb$variables),  "p.adj"]
stat.nb.wilcox$SIG.wilcox <- ifelse(stat.nb.wilcox$padj.wilcoxon<0.05, "**",
↪  ifelse(stat.nb.wilcox$padj.wilcoxon<0.2, "*", "n.s."))

mytest = "wilcox_test"
saveRDS(stat.nb.wilcox, file = paste0("ARGwPseudocount", "_Asthma_", mytest, "_",
↪  mycorrection, ".rds"))
write.table(stat.nb.wilcox, file = paste0("ARGwPseudocount", "_Asthma_", mytest, "_",
↪  mycorrection, ".tsv"), sep = "\t")
```

Make boxplots negative binomials

```
source("./humann3_210613_functions_ngw.R")
rm(list=ls()[ls() %notin% c("%notin%", "makePCA", "makeNMDSplot",
↪  "boxplotByAsthmaAndAge", "boxplotByAsthma", "my_data_summary",
↪  "makeBarplotDifferentialPairs", "filter_shortbred_data", "removeColumnSuffix")])
# setwd("~/Library/CloudStorage/Box-Box/Kau
↪  Lab/Results/MARS/FecalMetagenomics/shortbred/CARD2021results")
```

```r
library(ggplot2)
library(RColorBrewer)
detach(package:plyr)
detach(package:dplyr)
detach(package:tidyverse)
library(plyr); library(dplyr)
```

```
##
## Attaching package: 'plyr'

## The following objects are masked from 'package:rstatix':
##
##     desc, mutate

## The following object is masked from 'package:purrr':
##
##     compact

## The following object is masked from 'package:ggpubr':
##
##     mutate

##
## Attaching package: 'dplyr'

## The following objects are masked from 'package:plyr':
##
##     arrange, count, desc, failwith, id, mutate, rename, summarise,
##     summarize

## The following object is masked from 'package:MASS':
##
##     select

## The following object is masked from 'package:car':
##
##     recode

## The following objects are masked from 'package:stats':
##
##     filter, lag

## The following objects are masked from 'package:base':
##
##     intersect, setdiff, setequal, union
```

```r
library(tidyverse)
# library(plyr); library(dplyr)

group_by_me = "ASTHMA"
mytest = "negative_binomial"
mycorrection="fdr"
my_color_scheme = brewer.pal(7, "BrBG")[c(7,5)]

basedir="./"
humann3_mf_filtered_perm <- readRDS(file = paste0(basedir, "humann3_mf_filtered.rds"))
row.names(humann3_mf_filtered_perm) <- humann3_mf_filtered_perm$SampleName
df.meta <- humann3_mf_filtered_perm
```

```r
df.meta$ASTHMA <- ifelse(df.meta$ASTHMA == "Asthmatic", "asthmatic", "healthy")

df.shortbred <- readRDS(file = "SB_CARD21_with_AMRFam_ARO_Drug_Mechanism_filtered.rds")
df.shortbred <- df.shortbred[,1:dim(df.meta)[1]]
row.names(df.shortbred) = comprehenr::to_vec(for (i in row.names(df.shortbred))
↪  stringr::str_split(i, pattern = "__[[a-z]|[A-Z]]")[[1]][1]) # remove the taxonomic
↪  inffo
row.names(df.shortbred) <- gsub(row.names(df.shortbred), pattern = "[|]", replacement =
↪  ".")
row.names(df.shortbred) <- gsub(row.names(df.shortbred), pattern =
↪  "[[-]|[\"]|[(]|[)]|[|]]", replacement = ".")
row.names(df.shortbred) <- str_wrap(row.names(df.shortbred), width = 40)
pseudocount =
↪  min(df.shortbred[,1:dim(df.meta)[1]][df.shortbred[,1:dim(df.meta)[1]]>0])/20
logdat <- log(df.shortbred+pseudocount, base = 10) # add a pseudocount that is far less
↪  than the min value for log scale plot

stat.nb.wilcox <- readRDS(file = paste0("ARGwPseudocount", "_Asthma_", mytest, "_",
↪  mycorrection, ".rds"))

stat.nb.wilcox[order(stat.nb.wilcox$incidence, decreasing = T),]
```

```
## # A tibble: 71 x 6
## # Groups:   variables [71]
##    ASTHMA    variables                                     p    padj SIG   incidence
##    <fct>     <chr>                                     <dbl>   <dbl> <chr>     <dbl>
##  1 Asthmatic gb.CAA10975_1.ARO_3000194.tetW__But~ 0.519   0.714  n.s.         95
##  2 Asthmatic gb.AAA23033_2.ARO_3000190.tetO__Cam~ 0.143   0.352  n.s.         94
##  3 Asthmatic gb.AAD01868_1.ARO_3002867.dfrF__Ent~ 0.224   0.466  n.s.         94
##  4 Asthmatic gb.ACT97371_1.ARO_3003097.CfxA6__un~ 0.534   0.714  n.s.         92
##  5 Asthmatic gb.CAA79727_1.ARO_3000191.tetQ__Bac~ 0.705   0.824  n.s.         88
##  6 Asthmatic gb.AAF86219_1.ARO_3000375.ErmB__Ent~ 0.00122 0.0144 **           87
##  7 Asthmatic gb.AAF74725_1.ARO_3004659.Mef.En2._~ 0.362   0.643  n.s.         77
##  8 Asthmatic gb.AAA88675_1.ARO_3000498.ErmF__Bac~ 0.0156  0.0964 *            75
##  9 Asthmatic gb.CAM12479_1.ARO_3000567.tet.40.__~ 0.0875  0.296  n.s.         74
## 10 Asthmatic gb.AAC37034_1.ARO_3000522.ErmG__Bac~ 0.112   0.332  n.s.         71
## # ... with 61 more rows
```

```r
stat.nb.wilcox.sig <- stat.nb.wilcox[stat.nb.wilcox$padj<0.2,]
stat.nb.wilcox.sig[order(stat.nb.wilcox.sig$incidence, decreasing = TRUE),]
```

```
## # A tibble: 17 x 6
## # Groups:   variables [17]
##    ASTHMA    variables                                    p    padj SIG   incidence
##    <fct>     <chr>                                    <dbl>   <dbl> <chr>     <dbl>
##  1 Asthmatic gb.AAF86219_1.ARO_3000375.ErmB__En~ 1.22e-3 1.44e-2 **           87
##  2 Asthmatic gb.AAA88675_1.ARO_3000498.ErmF__Ba~ 1.56e-2 9.64e-2 *            75
##  3 Asthmatic gb.AHA41505_1.ARO_3002930.vanRO__R~ 2.25e-2 1.06e-1 *            37
##  4 Asthmatic gb.AAL59753_1.ARO_3000412.sul2__Vi~ 4.51e-2 1.88e-1 *            21
##  5 Asthmatic gb.AGV10818_1.ARO_3002669.APH.2__.~ 9.24e-3 7.29e-2 *            19
##  6 Asthmatic gb.AAD51345_1.ARO_3003052.smeB__St~ 2.19e-2 1.06e-1 *            19
##  7 Asthmatic gb.BAE78083_1.ARO_3003549.mdtO__Es~ 5.05e-4 7.17e-3 **           12
##  8 Asthmatic gb.AAC23556_1.ARO_3002660.APH.6..I~ 5.45e-5 1.93e-3 **           11
##  9 Asthmatic gb.AAP43109_1.ARO_3003922.oqxA__Es~ 1.23e-7 8.77e-6 **           10
```

```
## 10 Asthmatic gb.AET10445_1.ARO_3004033.tetB.46.~ 8.37e-3 7.29e-2 *          10
## 11 Asthmatic gb.AAB60941_1.ARO_3005099.23S_rRNA~ 1.77e-2 9.64e-2 *          10
## 12 Asthmatic gb.BAA15221_2.ARO_3000263.marA__Es~ 1.08e-2 7.64e-2 *           9
## 13 Asthmatic gb.ACS83559_1.ARO_3004601.LnuP__Cl~ 1.40e-4 3.31e-3 **          8
## 14 Asthmatic gb.BAE77933_1.ARO_3000518.CRP__Esc~ 4.40e-4 7.17e-3 **          7
## 15 Asthmatic gb.AAC77110_1.ARO_3004290.Escheric~ 2.73e-3 2.77e-2 **          7
## 16 Asthmatic gb.BAA16344_1.ARO_3000491.acrD__Es~ 1.75e-2 9.64e-2 *           7
## 17 Asthmatic gb.AAK76136_1.ARO_3000025.patB__St~ 3.10e-2 1.37e-1 *           7
```

```
numSamples <- dim(df.meta)[1]
incidence_treshold = numSamples*.1 # for plot, show those who occur in 10% or more
↪  samples

df.shortbred.meta <- data.frame(t(df.shortbred))
df.shortbred.meta$asthma <- as.character(df.meta[row.names(df.shortbred.meta), "ASTHMA"])
df.shortbred.meta <- df.shortbred.meta %>%
  pivot_longer(-asthma)
directioninfo <- df.shortbred.meta %>%
  group_by(asthma, name) %>%     # if this throws an error, you may need to restart R to
  ↪  purge loaded libraries
  summarize(incidence=sum(value>0), mean=mean(value)) %>%
  arrange(name)
```

```
## `summarise()` has grouped output by 'asthma'. You can override using the
## `.groups` argument.
```

```
directioninfo
```

```
## # A tibble: 142 x 4
## # Groups:   asthma [2]
##    asthma    name                                          incidence    mean
##    <chr>     <chr>                                             <int>   <dbl>
##  1 asthmatic gb.AAA20116_1.ARO_3000180.tetA.P.                    11  0.133
##  2 healthy   gb.AAA20116_1.ARO_3000180.tetA.P.                    10  0.109
##  3 asthmatic gb.AAA20117_1.ARO_3000195.tetB.P.                     2  0.0390
##  4 healthy   gb.AAA20117_1.ARO_3000195.tetB.P.                    12  0.0818
##  5 asthmatic gb.AAA21532_1.ARO_3003559.cepA_beta.lactamase         5  0.270
##  6 healthy   gb.AAA21532_1.ARO_3003559.cepA_beta.lactamase        12  0.536
##  7 asthmatic gb.AAA23033_2.ARO_3000190.tetO                       35 103.
##  8 healthy   gb.AAA23033_2.ARO_3000190.tetO                       59 135.
##  9 asthmatic gb.AAA27459_1.ARO_3004683.aadS                       26  10.5
## 10 healthy   gb.AAA27459_1.ARO_3004683.aadS                       33   7.51
## # ... with 132 more rows
```

```
write.csv(directioninfo, "./means_incidence_asthma.csv")
fc.tb <- directioninfo %>%
  group_by(name) %>%
  summarize(fc=mean[1]/mean[2])
write.csv(fc.tb, "./means_foldchangeAvH_asthma.csv")

stat.nb.wilcox.sig[stat.nb.wilcox.sig$incidence >= incidence_treshold, ]
```

```
## # A tibble: 11 x 6
## # Groups:   variables [11]
##    ASTHMA    variables                                    p    padj SIG   incidence
```

```
##    <fct>     <chr>                                      <dbl>   <dbl> <chr>     <dbl>
##  1 Asthmatic gb.AAP43109_1.ARO_3003922.oqxA__Es~ 1.23e-7 8.77e-6 **           10
##  2 Asthmatic gb.AAC23556_1.ARO_3002660.APH.6..I~ 5.45e-5 1.93e-3 **           11
##  3 Asthmatic gb.BAE78083_1.ARO_3003549.mdtO__Es~ 5.05e-4 7.17e-3 **           12
##  4 Asthmatic gb.AAF86219_1.ARO_3000375.ErmB__En~ 1.22e-3 1.44e-2 **           87
##  5 Asthmatic gb.AET10445_1.ARO_3004033.tetB.46.~ 8.37e-3 7.29e-2 *            10
##  6 Asthmatic gb.AGV10818_1.ARO_3002669.APH.2__.~ 9.24e-3 7.29e-2 *            19
##  7 Asthmatic gb.AAA88675_1.ARO_3000498.ErmF__Ba~ 1.56e-2 9.64e-2 *            75
##  8 Asthmatic gb.AAB60941_1.ARO_3005099.23S_rRNA~ 1.77e-2 9.64e-2 *            10
##  9 Asthmatic gb.AAD51345_1.ARO_3003052.smeB__St~ 2.19e-2 1.06e-1 *            19
## 10 Asthmatic gb.AHA41505_1.ARO_3002930.vanRO__R~ 2.25e-2 1.06e-1 *            37
## 11 Asthmatic gb.AAL59753_1.ARO_3000412.sul2__Vi~ 4.51e-2 1.88e-1 *            21
```

```
negbin_sig <- stat.nb.wilcox.sig[stat.nb.wilcox.sig$incidence >= incidence_treshold,
↪  "variables"]

negbin_sig = comprehenr::to_vec(for (i in negbin_sig$variables) stringr::str_split(i,
↪  pattern = "__[[a-z]|[A-Z]]")[[1]][1])
negbin_sig <- str_wrap(negbin_sig, width = 40)

directioninfo[directioninfo$name %in% negbin_sig,] %>%
  group_by(name) %>%
  summarize(fc=mean[1]/mean[2])
```

```
## # A tibble: 11 x 2
##    name                                                                         fc
##    <chr>                                                                     <dbl>
##  1 gb.AAA88675_1.ARO_3000498.ErmF                                             2.71
##  2 gb.AAB60941_1.ARO_3005099.23S_rRNA_.adenine.2058..N.6...methyltransfe~     3.43
##  3 gb.AAC23556_1.ARO_3002660.APH.6..Id                                        0.136
##  4 gb.AAD51345_1.ARO_3003052.smeB                                             2.98
##  5 gb.AAF86219_1.ARO_3000375.ErmB                                             2.81
##  6 gb.AAL59753_1.ARO_3000412.sul2                                             0.371
##  7 gb.AAP43109_1.ARO_3003922.oqxA                                            11.4
##  8 gb.AET10445_1.ARO_3004033.tetB.46.                                         3.73
##  9 gb.AGV10818_1.ARO_3002669.APH.2__..Ig                                      0.332
## 10 gb.AHA41505_1.ARO_3002930.vanRO                                            3.06
## 11 gb.BAE78083_1.ARO_3003549.mdtO                                             5.58
```

```
if(all(negbin_sig %in% row.names(logdat))){
  p1box <- makeBarplotDifferentialPairs(my_data_table_name =
↪  data.frame(t(logdat)),list_to_plot = negbin_sig, basedir = "./", sample_mf = df.meta,
↪  group_by_me = "ASTHMA", ordered_by_abundance = TRUE, check_normalization=FALSE,
↪  my_title = "CARD Differential ARGs (q<0.2 Negative Binomial,\n10% incidence)",
↪  my_xlab="ARG",norm_type = "RPKM", plot_method = "boxplot")
  p1box <- p1box+ylab("log10(RPKM)") + geom_abline(slope = 0, intercept =
↪  log(pseudocount, base = 10), lty=2) #+ scale_x_discrete(labels=c("-3" = "LOD",
↪  "-2"="-2", "-1"="-1", "0"="0", "1"="1", "2"="2"))
}
```

```
## Using SampleName, Group as id variables
```

```
p1box
```

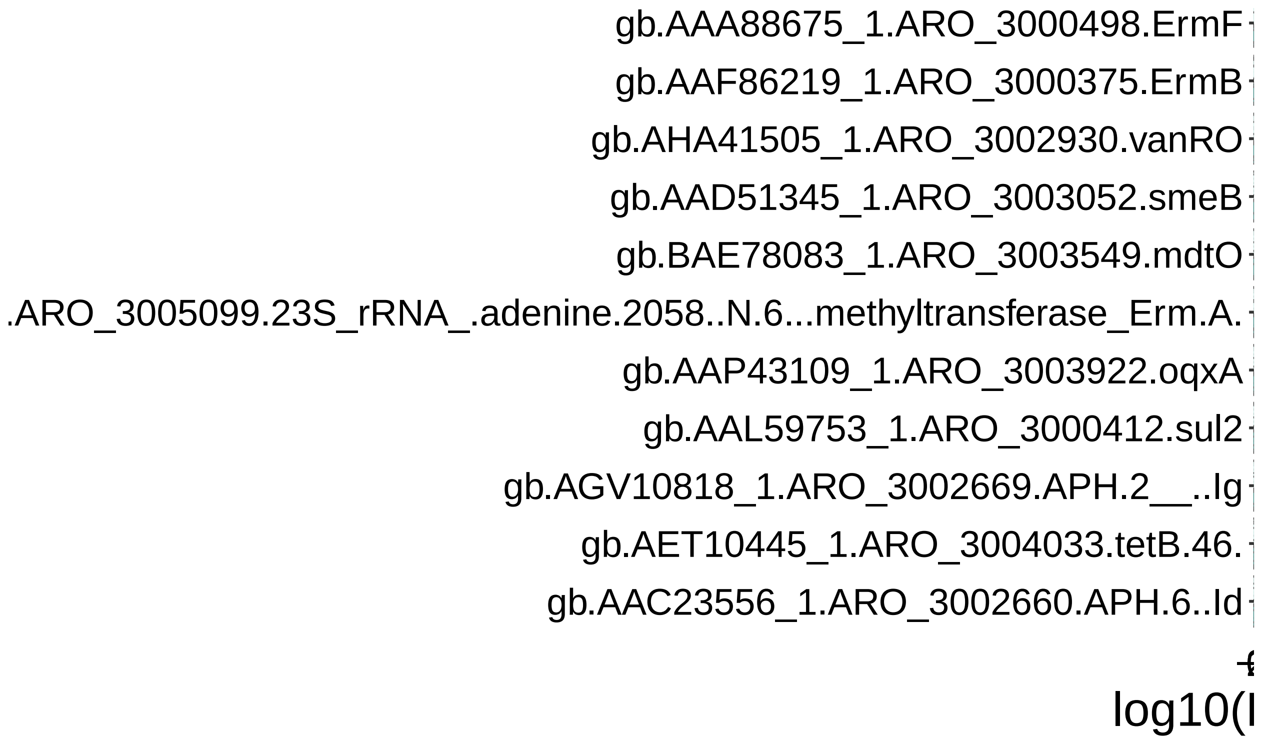


```
# ggsave(p1box, filename = "./negative_binomial_boxplots.jpg", device = "jpeg", width =
↪  12, height = 8, units = "in")
# ggsave(p1box, filename = "./negative_binomial_boxplots.pdf", device = "pdf", width =
↪  12, height = 8, units = "in", useDingbats=FALSE)
```

#### Figure 4B: table

```
# see ~/Library/CloudStorage/Box-Box/Kau
↪  Lab/Results/MARS/FecalMetagenomics/scaffold_annotation/ermF/count_genes_near_ermF.R

# Make table of ermF context in MARS metagenomes:
# plasmid-associated genes
# transposases
# antibiotic resistance genes
# within +/- 10 kilobases of ermF

# this reads in the modified gff table of contigs that had a BLAST hit to the CARD ermF
↪  sequence (prokka called it ermD)
# this table does not have any of the ermFs that were the only annotated gene on its
↪  contig since we cannot get context from those.
dat <- read.table("~/Library/CloudStorage/Box-Box/Kau
↪  Lab/Results/MARS/FecalMetagenomics/scaffold_annotation/ermF/all_CARDermF_BLAST_contigs_nodups/all_C
↪
           sep = "\t", header = FALSE)
dat$V10 = NULL
names(dat) <- c("NODE", "prodigal", "type", "start", "stop", "notsure", "strand", "zero",
↪  "GENE") # give em readable names

library(tidyverse)
```

```
ermD_lines <- dplyr::filter(dat, grepl("ermD",GENE))
# ermD_lines$NODE # all nodes that contain ermF
ermF_pos_node.tb <- table(ermD_lines$NODE) # get num of ermFs per node

# filter out low confidence nodes (no ermF annotation)
dat2 <- dat[dat$NODE %in% as.character(ermD_lines$NODE),]
all(as.character(unique(dat2$NODE)) %in% as.character(ermD_lines$NODE)) # check
↪  successful filtering - should be TRUE
```

```
## [1] TRUE
```

```
# cbind(sort(as.character(unique(dat2$NODE)))
↪  ,sort(as.character(unique(ermD_lines$NODE)))) # check visually

my_radius <- 10000 # how many amino acids to extend thru for counting ORFs

# make filtered df ####
# filter out genes outside +/- 10 kilobases of an ermF on a given contig
# this step isn't necessary except if you want a clean table with only the genes you are
↪  going to count from:
filtered.df <- data.frame(matrix(ncol = 9, nrow = 0))
↪  #data.frame(t(c(1,2,3,4,5,6,7,8,9)))
names(filtered.df) <- names(dat2)
for (myNode in as.character(ermD_lines$NODE)) {
  tmp.df <- dat2[dat2$NODE == myNode,]
  # set boundaries for respective node:
  newradius.df <- tmp.df[grep(x = tmp.df$GENE, pattern = "ermD"), c("start", "stop")]
  start_count <- ifelse((newradius.df[1,1] - my_radius) < 0, 0, (newradius.df[1,1] -
↪  my_radius))
  stop_count <- newradius.df[dim(newradius.df)[1], dim(newradius.df)[2]] + my_radius
  # filter out anything outside radius
  filtered.df <- rbind(filtered.df, tmp.df[tmp.df$start >= start_count & tmp.df$stop <=
↪  stop_count,])
}
# filtered.df
length(unique(as.character(filtered.df$NODE))) ==
↪  length(as.character(unique(ermD_lines$NODE))) # check nodes
```

```
## [1] TRUE
```

```
# saveRDS(filtered.df, "MARS_ermF_10kb_context.gff.rds")

# kmo
kmo <- dplyr::filter(filtered.df, grepl("kmo",GENE))
length(unique(kmo$NODE))  # 8 contigs
```

```
## [1] 8
```

```
length(unique(kmo$NODE))/length(unique(filtered.df$NODE))
```

```
## [1] 0.2285714
```

```
# make context tables ####
# count nearby genes by group
myrows <- c("transposase",
```

```
            "antitox",
            "replication",
            "mobilization",
            "conjugative",
            "clindamycin",
            "tet",
            "lactamase",
            "aadK")
context_tbl <- data.frame(matrix(ncol = 0, nrow = length(myrows)))
↪  #data.frame(t(c(1,2,3,4,5,6,7,8,9)))
row.names(context_tbl) <- myrows
for (i in grep(pattern = "ermD", x=filtered.df$GENE)) {
  # grab everybody at radius around erm:
  tmpstart <- ifelse((filtered.df[i, c("start")] - my_radius)<0, 0, (filtered.df[i,
↪  c("start")] - my_radius))
  tmpstop <- filtered.df[i, c("stop")] + my_radius
  tmpfiltered.df <- filtered.df[(filtered.df$NODE==as.character(filtered.df[i, "NODE"]) &
↪  filtered.df$start >= tmpstart & filtered.df$stop <= tmpstop),]
  mycounts = c(length(grep(x=tmpfiltered.df$GENE, pattern = "(transposase)|(tnp)",
↪  ignore.case = TRUE)),

                length(grep(x=tmpfiltered.df$GENE, pattern = "antitox", ignore.case =
                ↪  TRUE)),
                length(grep(x=tmpfiltered.df$GENE, pattern = "replication", ignore.case
                ↪  = TRUE)),
                length(grep(x=tmpfiltered.df$GENE, pattern = "mobilization",
                ↪  ignore.case = TRUE)),
                length(grep(x=tmpfiltered.df$GENE, pattern = "conjugative", ignore.case
                ↪  = TRUE)),
             # length(grep(x=tmpfiltered.df$GENE, pattern = "pcf", ignore.case =
             ↪  TRUE)), # pcfK and pcfJ??

                length(grep(x=tmpfiltered.df$GENE, pattern = "clindamycin", ignore.case
                ↪  = TRUE)),
                length(grep(x=tmpfiltered.df$GENE, pattern = "tet", ignore.case =
                ↪  TRUE)),
                length(grep(x=tmpfiltered.df$GENE, pattern = "lactamase", ignore.case =
                ↪  TRUE)),
                length(grep(x=tmpfiltered.df$GENE, pattern = "aadK", ignore.case =
                ↪  TRUE))) # streptomycin resistance
  context_tbl <- cbind(context_tbl, as.numeric(mycounts))
  if (as.character(filtered.df[i, "NODE"]) %in% names(context_tbl)) {
    if (max(ermF_pos_node.tb)==2){
      names(context_tbl)[length(names(context_tbl))] = paste0(as.character(filtered.df[i,
      ↪  "NODE"]), "_2ndErmF")
    }
  } else {
    names(context_tbl)[length(names(context_tbl))] = as.character(filtered.df[i, "NODE"])
  }
}

# summarize into percentages/prevalences
context_tbl.bin <- context_tbl
```

```r
context_tbl.bin[context_tbl.bin>0]=1
context_tbl_prevalence <- data.frame(rowMeans(context_tbl.bin)*100)
context_tbl_prevalence <- cbind(context_tbl_prevalence, rowSums(context_tbl.bin))
context_tbl_prevalence <- cbind(context_tbl_prevalence, dim(context_tbl.bin)[2])
context_tbl_prevalence <- cbind(context_tbl_prevalence, dim(context_tbl.bin)[2]+28)
names(context_tbl_prevalence) <- c("prevalence", "sum", "total_nonorphaned_ermF",
↪  "total_ermF_by_BLASTorProkka")

# supergroups
context_tbl_supergroups <- data.frame(matrix(nrow = 3, ncol = dim(context_tbl)[2]))
names(context_tbl_supergroups) <- names(context_tbl)
context_tbl_supergroups[1,] <- colSums(context_tbl[c("antitox", "replication",
↪  "mobilization", "conjugative"),])
context_tbl_supergroups[2,] <- colSums(context_tbl["transposase",])
context_tbl_supergroups[3,] <- colSums(context_tbl[c("clindamycin", "tet", "lactamase",
↪  "aadK"),])
row.names(context_tbl_supergroups) <- c("plasmid", "tnp", "ARG")
# summarize into percentages/prevalences
context_tbl_supergroups.bin <- context_tbl_supergroups
context_tbl_supergroups.bin[context_tbl_supergroups.bin>0]=1
context_tbl_supergroup_prevalence <-
↪  data.frame(rowMeans(context_tbl_supergroups.bin)*100)
context_tbl_supergroup_prevalence <- cbind(context_tbl_supergroup_prevalence,
↪  rowSums(context_tbl_supergroups.bin))
context_tbl_supergroup_prevalence <- cbind(context_tbl_supergroup_prevalence,
↪  dim(context_tbl_supergroups.bin)[2])
context_tbl_supergroup_prevalence <- cbind(context_tbl_supergroup_prevalence,
↪  dim(context_tbl_supergroups.bin)[2]+28) # 28 were orphaned (no ORF near ermF)
names(context_tbl_supergroup_prevalence) <- c("prevalence", "sum",
↪  "total_nonorphaned_ermF", "total_ermF_by_BLASTorProkka")
context_tbl_supergroup_prevalence
```

```
##          prevalence sum total_nonorphaned_ermF total_ermF_by_BLASTorProkka
## plasmid    57.89474  22                     38                          66
## tnp        50.00000  19                     38                          66
## ARG        50.00000  19                     38                          66
```

```r
# setwd("~/Library/CloudStorage/Box-Box/Kau
↪  Lab/Results/MARS/FecalMetagenomics/scaffold_annotation/ermF")
write.table(x = context_tbl_supergroup_prevalence, file="ermF_context_summary_table.txt",
↪  sep = "\t", row.names = TRUE)
write.table(x = context_tbl_prevalence, file="ermF_context_summary_table_genenames.txt",
↪  sep = "\t")

# Count specific transposases ####
tnp_names <- c()
for (i in filtered.df[grep(pattern = "(transposase)|(tnp)", x=filtered.df$GENE,
↪  ignore.case = TRUE), "GENE"]) {
  # grab everybody at radius around erm:
  # tmpstart <- ifelse((filtered.df[i, c("start")] - my_radius)<0, 0, (filtered.df[i,
  ↪  c("start")] - my_radius))
  # tmpstop <- filtered.df[i, c("stop")] + my_radius
  # tmpfiltered.df <- filtered.df[(filtered.df$NODE==as.character(filtered.df[i, "NODE"])
  ↪  & filtered.df$start >= tmpstart & filtered.df$stop <= tmpstop),]
```

```
  # print(i)
  # if (any(grepl(x=tmpfiltered.df$GENE, pattern = "(transposase)|(tnp)", ignore.case =
  ↪  TRUE))) {
    # print(comprehenr::to_vec(for (j in stringr::str_split(i, pattern = ";"))
    ↪  j))#[grep(j, pattern = "product")]))
    tnp_names <- c(tnp_names, comprehenr::to_vec(for (j in unlist(stringr::str_split(i,
↪  pattern = ";"))) j[grep(j, pattern = "product")]))
  # } else {
    # print("No tnp")
  # }
}
tnp_names
```

```
##  [1] "product=IS5 family transposase ISBf6"
##  [2] "product=Tnp [Prevotella sp. MGM2]"
##  [3] "product=IS5 family transposase ISBf6"
##  [4] "product=IS5 family transposase ISBf6"
##  [5] "product=IS91 family transposase ISMno24"
##  [6] "product=IS982 family transposase IS1187"
##  [7] "product=IS91 family transposase ISMno24"
##  [8] "product=IS5 family transposase ISBf6"
##  [9] "product=IS91 family transposase ISMno24"
## [10] "product=IS5 family transposase ISBf6"
## [11] "product=IS5 family transposase ISBf6"
## [12] "product=IS30 family transposase IS4351"
## [13] "product=IS5 family transposase ISBf6"
## [14] "product=IS5 family transposase ISBf6"
## [15] "product=IS5 family transposase ISBf6"
## [16] "product=Tnp [Bacteroides ovatus]"
## [17] "product=IS30 family transposase IS4351"
## [18] "product=IS91 family transposase ISMno24"
```

```
length(tnp_names) == length(filtered.df[grep(pattern = "(transposase)|(tnp)",
↪  x=filtered.df$GENE, ignore.case = TRUE), "GENE"])
```

```
## [1] TRUE
```

```
unique(tnp_names)
```

```
## [1] "product=IS5 family transposase ISBf6"
## [2] "product=Tnp [Prevotella sp. MGM2]"
## [3] "product=IS91 family transposase ISMno24"
## [4] "product=IS982 family transposase IS1187"
## [5] "product=IS30 family transposase IS4351"
## [6] "product=Tnp [Bacteroides ovatus]"
```

```
# count transposases genes by group
mytnps <- unique(tnp_names)
tnp_context_tbl <- data.frame(matrix(ncol = 0, nrow = length(mytnps)))
↪  #data.frame(t(c(1,2,3,4,5,6,7,8,9)))
row.names(tnp_context_tbl) <- mytnps
for (i in grep(pattern = "ermD", x=filtered.df$GENE)) {
  # grab everybody at radius around erm:
  tmpstart <- ifelse((filtered.df[i, c("start")] - my_radius)<0, 0, (filtered.df[i,
↪  c("start")] - my_radius))
```

```
  tmpstop <- filtered.df[i, c("stop")] + my_radius
  tmpfiltered.df <- filtered.df[(filtered.df$NODE==as.character(filtered.df[i, "NODE"]) &
↪  filtered.df$start >= tmpstart & filtered.df$stop <= tmpstop),]
  mytnpcounts = c(length(grep(x=tmpfiltered.df$GENE, pattern = "ISBf6", ignore.case =
↪  TRUE)),
                              length(grep(x=tmpfiltered.df$GENE, pattern = "Tnp
                               ↪  [[]Bacteroides ovatus[]]", ignore.case = TRUE)),
                              length(grep(x=tmpfiltered.df$GENE, pattern = "ISMno24",
                               ↪  ignore.case = TRUE)),
                              length(grep(x=tmpfiltered.df$GENE, pattern = "IS4351",
                               ↪  ignore.case = TRUE)),
                              length(grep(x=tmpfiltered.df$GENE, pattern = "Tnp
                               ↪  [[]Prevotella sp. MGM2[]]", ignore.case = TRUE)),
                              length(grep(x=tmpfiltered.df$GENE, pattern = "IS1187",
                               ↪  ignore.case = TRUE)))

    tnp_context_tbl <- cbind(tnp_context_tbl, as.numeric(mytnpcounts))
    if (as.character(filtered.df[i, "NODE"]) %in% names(tnp_context_tbl)) {
      if (max(ermF_pos_node.tb)==2){
        names(tnp_context_tbl)[length(names(tnp_context_tbl))] =
         ↪  paste0(as.character(filtered.df[i, "NODE"]), "_2ndErmF")
      }
    } else {
      names(tnp_context_tbl)[length(names(tnp_context_tbl))] =
       ↪  as.character(filtered.df[i, "NODE"])
    }
  }

# # summarize into percentages/prevalences
tnp_context_tbl.bin <- tnp_context_tbl
tnp_context_tbl.bin[tnp_context_tbl.bin>0]=1
context_tnptbl_prevalence <- data.frame(rowMeans(tnp_context_tbl.bin)*100)
context_tnptbl_prevalence <- cbind(context_tnptbl_prevalence,
↪  rowSums(tnp_context_tbl.bin))
context_tnptbl_prevalence <- cbind(context_tnptbl_prevalence,
↪  dim(tnp_context_tbl.bin)[2])
context_tnptbl_prevalence <- cbind(context_tnptbl_prevalence,
↪  dim(tnp_context_tbl.bin)[2]+28)
names(context_tnptbl_prevalence) <- c("prevalence", "sum", "total_nonorphaned_ermF",
↪  "total_ermF_by_BLASTorProkka")
context_tnptbl_prevalence <- cbind(context_tnptbl_prevalence,
↪  100*context_tnptbl_prevalence$sum/context_tbl_prevalence["transposase", "sum"]) # get
↪  percentage within transposase-positive contigs
names(context_tnptbl_prevalence) <- c("prevalence", "sum", "total_nonorphaned_ermF",
↪  "total_ermF_by_BLASTorProkka", "proportion_of_tnps")
# context_tnptbl_prevalence[order(context_tnptbl_prevalence$proportion_of_tnps,
↪  decreasing = TRUE),]

# Breakdown ARGs ####
arg_names <- c()
for (i in names(context_tbl_supergroups)[context_tbl_supergroups["ARG",] >0]) {
  # grab everybody at radius around erm:
  # tmpstart <- ifelse((filtered.df[i, c("start")] - my_radius)<0, 0, (filtered.df[i,
   ↪  c("start")] - my_radius))
```

```r
  # tmpstop <- filtered.df[i, c("stop")] + my_radius
  # tmpfiltered.df <- filtered.df[(filtered.df$NODE==as.character(filtered.df[i, "NODE"])
  ↪  & filtered.df$start >= tmpstart & filtered.df$stop <= tmpstop),]
  # print(i)
  current_node <- filtered.df[filtered.df$NODE == i,]
  if (any(grepl(x=current_node$GENE, pattern = "(clindamycin)|(tet)|(lactamase)|(aadK)",
  ↪  ignore.case = TRUE))) {
    # print(current_node$NODE[1])
    # print(comprehenr::to_vec(for (j in stringr::str_split(current_node, pattern = ";"))
    ↪  j)) #[grep(j, pattern = "product")]))
    current_ARGs <- grep(x=current_node$GENE, pattern =
↪  "(clindamycin)|(tet)|(lactamase)|(aadK)", ignore.case = TRUE)
    for (idx in current_ARGs) {
      arg_names <- c(arg_names, comprehenr::to_vec(for (j in
↪  unlist(stringr::str_split(current_node[idx,"GENE"], pattern = ";"))) j[grep(j,
↪  pattern = "product")]))
    }
  }# } else {
  # print("No tnp")
  # }
}
arg_names
```

```
##  [1] "product=clindamycin resistance transfer factor btgA [Bacteroidales]"
##  [2] "product=clindamycin resistance transfer factor btgA [Bacteroidales]"
##  [3] "product=clindamycin resistance transfer factor btgA [Bacteroidales]"
##  [4] "product=tetracycline resistance ribosomal protection protein Tet(Q)"
##  [5] "product=Aminoglycoside 6-adenylyltransferase"
##  [6] "product=tetracycline resistance ribosomal protection protein Tet(Q)"
##  [7] "product=clindamycin resistance transfer factor btgA [Bacteroidales]"
##  [8] "product=Aminoglycoside 6-adenylyltransferase"
##  [9] "product=Beta-lactamase OXA-10"
## [10] "product=clindamycin resistance transfer factor btgA [Bacteroidales]"
## [11] "product=clindamycin resistance transfer factor btgA [Bacteroidales]"
## [12] "product=clindamycin resistance transfer factor btgA [Bacteroidales]"
## [13] "product=clindamycin resistance transfer factor btgA [Bacteroidales]"
## [14] "product=clindamycin resistance transfer factor btgA [Bacteroidales]"
## [15] "product=clindamycin resistance transfer factor btgA [Bacteroidales]"
## [16] "product=clindamycin resistance transfer factor btgA [Bacteroidales]"
## [17] "product=Tetracycline resistance protein TetO"
```

```r
length(arg_names) == length(filtered.df[grep(pattern =
↪  "(clindamycin)|(tet)|(lactamase)|(aadK)", x=filtered.df$GENE, ignore.case = TRUE),
↪  "GENE"])
```

```
## [1] TRUE
```

```r
unique(arg_names)
```

```
## [1] "product=clindamycin resistance transfer factor btgA [Bacteroidales]"
## [2] "product=tetracycline resistance ribosomal protection protein Tet(Q)"
## [3] "product=Aminoglycoside 6-adenylyltransferase"
## [4] "product=Beta-lactamase OXA-10"
## [5] "product=Tetracycline resistance protein TetO"
```

```
# count transposases genes by group
myargs <- unique(arg_names)
arg_context_tbl <- data.frame(matrix(ncol = 0, nrow = length(myargs)))
↪  #data.frame(t(c(1,2,3,4,5,6,7,8,9)))
row.names(arg_context_tbl) <- myargs
for (i in grep(pattern = "ermD", x=filtered.df$GENE)) {
  # grab everybody at radius around erm:
  tmpstart <- ifelse((filtered.df[i, c("start")] - my_radius)<0, 0, (filtered.df[i,
↪  c("start")] - my_radius))
  tmpstop <- filtered.df[i, c("stop")] + my_radius
  tmpfiltered.df <- filtered.df[(filtered.df$NODE==as.character(filtered.df[i, "NODE"]) &
↪  filtered.df$start >= tmpstart & filtered.df$stop <= tmpstop),]
  myargcounts = c(length(grep(x=tmpfiltered.df$GENE, pattern = "clindamycin resistance
↪  transfer factor btgA", ignore.case = FALSE)),
                  length(grep(x=tmpfiltered.df$GENE, pattern = "Tetracycline resistance
                   ↪  protein TetO", ignore.case = TRUE)),
                  length(grep(x=tmpfiltered.df$GENE, pattern = "Aminoglycoside
                   ↪  6-adenylyltransferase", ignore.case = TRUE)),
                  length(grep(x=tmpfiltered.df$GENE, pattern = "tetracycline resistance
                   ↪  ribosomal protection protein Tet[(]Q[)]", ignore.case = TRUE)),
                  length(grep(x=tmpfiltered.df$GENE, pattern = "Beta-lactamase OXA",
                   ↪  ignore.case = TRUE)))

  arg_context_tbl <- cbind(arg_context_tbl, as.numeric(myargcounts))
  if (as.character(filtered.df[i, "NODE"]) %in% names(arg_context_tbl)) {
    if (max(ermF_pos_node.tb)==2){
      names(arg_context_tbl)[length(names(arg_context_tbl))] =
       ↪  paste0(as.character(filtered.df[i, "NODE"]), "_2ndErmF")
    }
  } else {
    names(arg_context_tbl)[length(names(arg_context_tbl))] = as.character(filtered.df[i,
     ↪  "NODE"])
  }
}

# # summarize into percentages/prevalences
arg_context_tbl.bin <- arg_context_tbl
arg_context_tbl.bin[arg_context_tbl.bin>0]=1
context_argtbl_prevalence <- data.frame(rowMeans(arg_context_tbl.bin)*100)
context_argtbl_prevalence <- cbind(context_argtbl_prevalence,
↪  rowSums(arg_context_tbl.bin))
context_argtbl_prevalence <- cbind(context_argtbl_prevalence,
↪  dim(arg_context_tbl.bin)[2])
context_argtbl_prevalence <- cbind(context_argtbl_prevalence,
↪  dim(arg_context_tbl.bin)[2]+28)
names(context_argtbl_prevalence) <- c("prevalence", "sum", "total_nonorphaned_ermF",
↪  "total_ermF_by_BLASTorProkka")
context_argtbl_prevalence <- cbind(context_argtbl_prevalence,
↪  100*context_argtbl_prevalence$sum/context_tbl_supergroup_prevalence["ARG", "sum"]) #
↪  get percentage within ARG-positive contigs
names(context_argtbl_prevalence) <- c("prevalence", "sum", "total_nonorphaned_ermF",
↪  "total_ermF_by_BLASTorProkka", "proportion_of_tnps")
# context_argtbl_prevalence[order(context_argtbl_prevalence$proportion_of_tnps,
↪  decreasing = TRUE),]
```

```
# Breakdown Plasmid-related genes ####
plasmid_names <- c()
for (i in names(context_tbl_supergroups)[context_tbl_supergroups["plasmid",] >0]) {
  # grab everybody at radius around erm:
  # tmpstart <- ifelse((filtered.df[i, c("start")] - my_radius)<0, 0, (filtered.df[i,
  ↪  c("start")] - my_radius))
  # tmpstop <- filtered.df[i, c("stop")] + my_radius
  # tmpfiltered.df <- filtered.df[(filtered.df$NODE==as.character(filtered.df[i, "NODE"])
  ↪  & filtered.df$start >= tmpstart & filtered.df$stop <= tmpstop),]
  # print(i)
  current_node <- filtered.df[filtered.df$NODE == i,]
  if (any(grepl(x=current_node$GENE, pattern =
  ↪  "(antitox)|(replication)|(mobilization)|(conjugative)", ignore.case = TRUE))) {
    print(current_node$NODE[1])
    # print(comprehenr::to_vec(for (j in stringr::str_split(current_node, pattern = ";"))
    ↪  j)) #[grep(j, pattern = "product")]))
    current_ARGs <- grep(x=current_node$GENE, pattern =
↪   "(antitox)|(replication)|(mobilization)|(conjugative)", ignore.case = TRUE)
    for (idx in current_ARGs) {
      plasmid_names <- c(plasmid_names, comprehenr::to_vec(for (j in
↪   unlist(stringr::str_split(current_node[idx,"GENE"], pattern = ";"))) j[grep(j,
↪   pattern = "product")]))
    }
  }# } else {
  # print("No tnp")
  # }
}
```

```
## [1] "NODE_774_length_8428_cov_8.073764"
## [1] "NODE_44_length_102108_cov_10.369897"
## [1] "NODE_1680_length_10244_cov_21.728927"
## [1] "NODE_997_length_18930_cov_223.500345"
## [1] "NODE_313_length_34664_cov_138.792494"
## [1] "NODE_1791_length_7324_cov_12.197185"
## [1] "NODE_532_length_18853_cov_55.769440"
## [1] "NODE_1065_length_12050_cov_5.852836"
## [1] "NODE_1830_length_8428_cov_113.077955"
## [1] "NODE_718_length_11885_cov_128.725356"
## [1] "NODE_5251_length_5056_cov_367.594095"
## [1] "NODE_5371_length_6032_cov_1985.945928"
## [1] "NODE_2184_length_8428_cov_190.211232"
## [1] "NODE_210_length_59732_cov_13.678351"
## [1] "NODE_2775_length_8428_cov_153.802778"
## [1] "NODE_1328_length_8486_cov_25.845047"
## [1] "NODE_1069_length_10540_cov_17.095193"
## [1] "NODE_1559_length_7645_cov_12.707320"
## [1] "NODE_449_length_34665_cov_211.027379"
```

```
plasmid_names
```

```
##  [1] "product=replication initiation protein [Bacteroidales]"
##  [2] "product=type II toxin-antitoxin system Phd/YefM family antitoxin [Bacteroidales]"
##  [3] "product=Txe/YoeB family addiction module toxin [Bacteria]"
```

```
##  [4] "product=nucleotidyl transferase AbiEii/AbiGii toxin family protein [Bacteroidales]"
##  [5] "product=type IV toxin-antitoxin system AbiEi family antitoxin domain-containing protein [Bacte
##  [6] "product=nucleotidyl transferase AbiEii/AbiGii toxin family protein [Bacteroidales]"
##  [7] "product=replication initiation protein [Bacteroidales]"
##  [8] "product=type II toxin-antitoxin system Phd/YefM family antitoxin [Bacteroidales]"
##  [9] "product=Txe/YoeB family addiction module toxin [Bacteria]"
## [10] "product=replication initiation protein [Bacteroidales]"
## [11] "product=type II toxin-antitoxin system Phd/YefM family antitoxin [Bacteroidales]"
## [12] "product=Txe/YoeB family addiction module toxin [Bacteria]"
## [13] "product=relaxase/mobilization nuclease domain-containing protein [Bacteroidetes]"
## [14] "product=plasmid mobilization relaxosome protein MobC [Bacteroidetes]"
## [15] "product=plasmid mobilization relaxosome protein MobC [Bacteroidales]"
## [16] "product=relaxase/mobilization nuclease domain-containing protein [Bacteroidales]"
## [17] "product=nucleotidyl transferase AbiEii/AbiGii toxin family protein [Bacteroides fragilis]"
## [18] "product=nucleotidyl transferase AbiEii/AbiGii toxin family protein [Bacteroidales]"
## [19] "product=replication initiation protein [Bacteroidales]"
## [20] "product=type II toxin-antitoxin system Phd/YefM family antitoxin [Bacteroidales]"
## [21] "product=plasmid mobilization relaxosome protein MobC [Bacteroidetes]"
## [22] "product=relaxase/mobilization nuclease domain-containing protein [Bacteroidetes]"
## [23] "product=nucleotidyl transferase AbiEii/AbiGii toxin family protein [Bacteroidales]"
## [24] "product=nucleotidyl transferase AbiEii/AbiGii toxin family protein [Bacteroides fragilis]"
## [25] "product=relaxase/mobilization nuclease domain-containing protein [Bacteroidales]"
## [26] "product=plasmid mobilization relaxosome protein MobC [Bacteroidales]"
## [27] "product=replication initiation protein [Bacteroidales]"
## [28] "product=type II toxin-antitoxin system Phd/YefM family antitoxin [Bacteroidales]"
## [29] "product=Txe/YoeB family addiction module toxin [Bacteria]"
## [30] "product=replication initiation protein [Bacteroidales]"
## [31] "product=type II toxin-antitoxin system Phd/YefM family antitoxin [Bacteroidales]"
## [32] "product=Txe/YoeB family addiction module toxin [Bacteria]"
## [33] "product=Txe/YoeB family addiction module toxin [Bacteria]"
## [34] "product=type II toxin-antitoxin system Phd/YefM family antitoxin [Bacteroidales]"
## [35] "product=replication initiation protein [Bacteroidales]"
## [36] "product=Txe/YoeB family addiction module toxin [Bacteria]"
## [37] "product=type II toxin-antitoxin system Phd/YefM family antitoxin [Bacteroidales]"
## [38] "product=replication initiation protein [Bacteroidales]"
## [39] "product=replication initiation protein [Bacteroidales]"
## [40] "product=type II toxin-antitoxin system Phd/YefM family antitoxin [Bacteroidales]"
## [41] "product=Txe/YoeB family addiction module toxin [Bacteria]"
## [42] "product=nucleotidyl transferase AbiEii/AbiGii toxin family protein [Bacteroides fragilis]"
## [43] "product=DNA replication and repair protein RecF"
## [44] "product=Txe/YoeB family addiction module toxin [Bacteria]"
## [45] "product=type II toxin-antitoxin system Phd/YefM family antitoxin [Bacteroidales]"
## [46] "product=replication initiation protein [Bacteroidales]"
## [47] "product=Txe/YoeB family addiction module toxin [Bacteria]"
## [48] "product=type II toxin-antitoxin system Phd/YefM family antitoxin [Bacteroidales]"
## [49] "product=replication initiation protein [Bacteroidales]"
## [50] "product=replication initiation protein [Bacteroidales]"
## [51] "product=type II toxin-antitoxin system Phd/YefM family antitoxin [Bacteroidales]"
## [52] "product=Txe/YoeB family addiction module toxin [Bacteria]"
## [53] "product=nucleotidyl transferase AbiEii/AbiGii toxin family protein [Bacteroides fragilis]"
## [54] "product=nucleotidyl transferase AbiEii/AbiGii toxin family protein [Bacteroidales]"
## [55] "product=nucleotidyl transferase AbiEii/AbiGii toxin family protein [Bacteroidales]"
## [56] "product=nucleotidyl transferase AbiEii/AbiGii toxin family protein [Bacteroides fragilis]"
## [57] "product=relaxase/mobilization nuclease domain-containing protein [Bacteroidales]"
```

```
## [58] "product=plasmid mobilization relaxosome protein MobC [Bacteroidales]"
```

```
length(plasmid_names) == length(filtered.df[grep(pattern =
↪  "(antitox)|(replication)|(mobilization)|(conjugative)", x=filtered.df$GENE,
↪  ignore.case = TRUE), "GENE"])
```

```
## [1] TRUE
```

```
unique(plasmid_names)
```

```
##  [1] "product=replication initiation protein [Bacteroidales]"
##  [2] "product=type II toxin-antitoxin system Phd/YefM family antitoxin [Bacteroidales]"
##  [3] "product=Txe/YoeB family addiction module toxin [Bacteria]"
##  [4] "product=nucleotidyl transferase AbiEii/AbiGii toxin family protein [Bacteroidales]"
##  [5] "product=type IV toxin-antitoxin system AbiEi family antitoxin domain-containing protein [Bacte
##  [6] "product=relaxase/mobilization nuclease domain-containing protein [Bacteroidetes]"
##  [7] "product=plasmid mobilization relaxosome protein MobC [Bacteroidetes]"
##  [8] "product=plasmid mobilization relaxosome protein MobC [Bacteroidales]"
##  [9] "product=relaxase/mobilization nuclease domain-containing protein [Bacteroidales]"
## [10] "product=nucleotidyl transferase AbiEii/AbiGii toxin family protein [Bacteroides fragilis]"
## [11] "product=DNA replication and repair protein RecF"
```

```
# count plasmid-related genes by group
myplas <- sort(unique(plasmid_names))
myplas <- myplas[c(1, 3, 4, 6, 8, 9, 10, 11)] # hard-coded dropping of similar names
plasmid_context_tbl <- data.frame(matrix(ncol = 0, nrow = length(myplas)))
↪  #data.frame(t(c(1,2,3,4,5,6,7,8,9)))
row.names(plasmid_context_tbl) <- myplas
for (i in grep(pattern = "ermD", x=filtered.df$GENE)) {
  # grab everybody at radius around erm:
  tmpstart <- ifelse((filtered.df[i, c("start")] - my_radius)<0, 0, (filtered.df[i,
↪  c("start")] - my_radius))
  tmpstop <- filtered.df[i, c("stop")] + my_radius
  tmpfiltered.df <- filtered.df[(filtered.df$NODE==as.character(filtered.df[i, "NODE"]) &
↪  filtered.df$start >= tmpstart & filtered.df$stop <= tmpstop),]
  myplascounts = c(length(grep(x=tmpfiltered.df$GENE, pattern = "DNA replication and
↪  repair protein RecF", ignore.case = FALSE)),
                   length(grep(x=tmpfiltered.df$GENE, pattern = "AbiEii[/]AbiGii toxin
                    ↪  family protein [[]Bacteroid", ignore.case = FALSE)),
                   length(grep(x=tmpfiltered.df$GENE, pattern = "plasmid mobilization
                    ↪  relaxosome protein MobC", ignore.case = FALSE)),
                   length(grep(x=tmpfiltered.df$GENE, pattern = "relaxase[/]mobilization
                    ↪  nuclease domain[-]containing protein [[]Bacteroid", ignore.case =
                    ↪  FALSE)),
                  length(grep(x=tmpfiltered.df$GENE, pattern = "replication initiation
                   ↪  protein [[]Bacteroid", ignore.case = FALSE)),
                  length(grep(x=tmpfiltered.df$GENE, pattern = "Txe[/]YoeB family
                   ↪  addiction module toxin [[]Bacteria[]]", ignore.case = FALSE)),
                  length(grep(x=tmpfiltered.df$GENE, pattern = "type II toxin[-]antitoxin
                   ↪  system Phd[/]YefM family antitoxin", ignore.case = FALSE)),
                  length(grep(x=tmpfiltered.df$GENE, pattern = "AbiEi family antitoxin
                   ↪  domain[-]containing protein", ignore.case = FALSE)))

  plasmid_context_tbl <- cbind(plasmid_context_tbl, as.numeric(myplascounts))
  if (as.character(filtered.df[i, "NODE"]) %in% names(plasmid_context_tbl)) {
```

```r
    if (max(ermF_pos_node.tb)==2){
      names(plasmid_context_tbl)[length(names(plasmid_context_tbl))] =
       ↪  paste0(as.character(filtered.df[i, "NODE"]), "_2ndErmF")
    }
  } else {
    names(plasmid_context_tbl)[length(names(plasmid_context_tbl))] =
     ↪  as.character(filtered.df[i, "NODE"])
  }
}

# # summarize into percentages/prevalences
plasmid_context_tbl.bin <- plasmid_context_tbl
plasmid_context_tbl.bin[plasmid_context_tbl.bin>0]=1
context_plastbl_prevalence <- data.frame(rowMeans(plasmid_context_tbl.bin)*100)
context_plastbl_prevalence <- cbind(context_plastbl_prevalence,
 ↪  rowSums(plasmid_context_tbl.bin))
context_plastbl_prevalence <- cbind(context_plastbl_prevalence,
 ↪  dim(plasmid_context_tbl.bin)[2])
context_plastbl_prevalence <- cbind(context_plastbl_prevalence,
 ↪  dim(plasmid_context_tbl.bin)[2]+28)
names(context_plastbl_prevalence) <- c("prevalence", "sum", "total_nonorphaned_ermF",
 ↪  "total_ermF_by_BLASTorProkka")
context_plastbl_prevalence <- cbind(context_plastbl_prevalence,
 ↪  100*context_plastbl_prevalence$sum/context_tbl_supergroup_prevalence["plasmid",
 ↪  "sum"]) # get percentage within ARG-positive contigs
names(context_plastbl_prevalence) <- c("prevalence", "sum", "total_nonorphaned_ermF",
 ↪  "total_ermF_by_BLASTorProkka", "proportion_of_tnps")
# context_plastbl_prevalence[order(context_plastbl_prevalence$proportion_of_tnps,
 ↪  decreasing = TRUE),]
# context_plastbl_prevalence[order(context_plastbl_prevalence$proportion_of_tnps,
 ↪  decreasing = TRUE),]
# View(context_plastbl_prevalence[order(context_plastbl_prevalence$proportion_of_tnps,
 ↪  decreasing = TRUE),])
```

### Figure 4D and 4E tables

```r
# ermf+ are contigs with a shortbred hit and bft+ are positive by PCR screen in MARS 1
 ↪  paper (2023 iscience)
dat <- read.csv("/Users/naomiwilson/Library/CloudStorage/Box-Box/Kau
 ↪  Lab/Results/MARS/FecalMetagenomics/scaffold_annotation/bft/bft_ermf_cooccurrence.csv")
dat <- data.frame(dat)
# dat
contab <- table(dat$bft_binary, dat$ermf_binary)
fisher.test(contab)
```

```
##
##  Fisher's Exact Test for Count Data
##
## data:  contab
## p-value = 0.2172
## alternative hypothesis: true odds ratio is not equal to 1
## 95 percent confidence interval:
```

```
##   0.6070085 16.2380785
## sample estimates:
## odds ratio
##   2.649433
```

```
fisher.test(contab, alternative = "greater")
```

```
##
##  Fisher's Exact Test for Count Data
##
## data:  contab
## p-value = 0.1272
## alternative hypothesis: true odds ratio is greater than 1
## 95 percent confidence interval:
##  0.7347945       Inf
## sample estimates:
## odds ratio
##   2.649433
```

```
# drop disqualifieds
humanSamples <- readRDS(file="./humann3_mf_filtered.rds")
row.names(humanSamples) <- humanSamples$MARSID
samplenames <- paste0(row.names(humanSamples), "_",
↪  ifelse(humanSamples$ASTHMA=="Asthmatic", "asthmatic", "healthy"))
row.names(dat) <- as.character(dat$Individual)
dat_95 <- dat[samplenames,]

library(readxl)
bft_screen <- read_excel("/Users/naomiwilson/Library/CloudStorage/Box-Box/Kau
↪  Lab/Results/MARS/Fragilysin_Toxin_Test/Frag_Test_for_qPCR.xlsx",
↪  sheet="FragPosSamples")
```

```
## New names:
## * `` -> `...1`
## * `` -> `...18`
## * `` -> `...19`
## * `` -> `...20`
## * `` -> `...21`
## * `` -> `...22`
## * `` -> `...23`
## * `` -> `...24`
```

```
bft_screen <- data.frame(bft_screen[,1:16])
row.names(bft_screen) <- paste0(bft_screen$...1, "_", bft_screen$asthma_status)
anyPositive <- bft_screen$everPositive + bft_screen$blast_metagenomes > 0
bft_screen$anyPositive <- anyPositive
# fix mars 1 data column
dat_95$anyPositiveBFT <- bft_screen[row.names(dat_95), "anyPositive"]
dat_95$anyPositiveBFT[is.na(dat_95$anyPositiveBFT)] <- FALSE
dat_95$bft_MARS1 <- bft_screen[row.names(dat_95), "everPositive"]
dat_95$bft_MARS1[is.na(dat_95$bft_MARS1)] <- FALSE

# UPDATE OCT 2022: consider bft PCR results (everPositive), and shortbred ermf detection
↪  ####
args_shortbred <- readRDS("/Users/naomiwilson/Library/CloudStorage/Box-Box/Kau
↪  Lab/Results/MARS/FecalMetagenomics/shortbred/CARD2021results/SB_CARD21_RPKM_filtered.rds")
```

```r
vegan::specnumber(args_shortbred[grepl(row.names(args_shortbred), pattern =
↪  "ErmF__Bacteroides_fragilis_"), as.character(dat_95$Sample.Name)]) # ARO_3000498
```

```
## gb|AAA88675_1|ARO_3000498|ErmF__Bacteroides_fragilis_
##                                                      75
```

```r
args_shortbred[args_shortbred>0]=1
args_shortbred <- t(args_shortbred)
ermf_rpkm <- args_shortbred[as.character(dat_95$Sample.Name),
↪  grepl(colnames(args_shortbred), pattern = "ErmF__Bacteroides_fragilis_")]
dat_95$ermf_shortbred <- ermf_rpkm
# dat_95[, c("ermf_shortbred", "bft_MARS1")]
contab_95_MARS2_final <- table(dat_95$bft_MARS1, dat_95$ermf_shortbred)
fisher.test(contab_95_MARS2_final, alternative = "greater") # p=0.6
```

```
##
##  Fisher's Exact Test for Count Data
##
## data:  contab_95_MARS2_final
## p-value = 0.6478
## alternative hypothesis: true odds ratio is greater than 1
## 95 percent confidence interval:
##  0.2350879       Inf
## sample estimates:
## odds ratio
##   1.073826
```

```r
# of only ermF+ samples, does bft occur more in asthma?
dat_95_ermFpos_only <- dat_95[dat_95$ermf_shortbred == 1,]
contab_95_MARS2_final_ermFPos <- table(dat_95_ermFpos_only$ASTHMA,
↪  dat_95_ermFpos_only$bft_MARS1)
contab_95_MARS2_final_ermFPos
```

```
##
##              FALSE TRUE
##   asthmatic    25    7
##   healthy      42    1
```

```r
fisher.test(contab_95_MARS2_final_ermFPos)
```

```
##
##  Fisher's Exact Test for Count Data
##
## data:  contab_95_MARS2_final_ermFPos
## p-value = 0.009202
## alternative hypothesis: true odds ratio is not equal to 1
## 95 percent confidence interval:
##  0.001849415 0.745409950
## sample estimates:
## odds ratio
## 0.08761096
```

```r
# of only ermF- samples, does bft occur more in asthma?
dat_95_ermFneg_only <- dat_95[dat_95$ermf_shortbred == 0,]
contab_95_MARS2_final_ermFNeg <- table(dat_95_ermFneg_only$ASTHMA,
↪  dat_95_ermFneg_only$bft_MARS1)
```

```
contab_95_MARS2_final_ermFNeg
```

```
##
##             FALSE TRUE
##   asthmatic     3    1
##   healthy      15    1
```

```
fisher.test(contab_95_MARS2_final_ermFNeg)
```

```
##
##  Fisher's Exact Test for Count Data
##
## data:  contab_95_MARS2_final_ermFNeg
## p-value = 0.3684
## alternative hypothesis: true odds ratio is not equal to 1
## 95 percent confidence interval:
##   0.002403181 20.893364831
## sample estimates:
## odds ratio
##   0.223606
```

### Filter Data

```
rm(list=ls()[ls() %notin% c("%notin%", "filter_shortbred_data", "boxplotByAsthmaAndAge",
↪  "removeColumnSuffix")])

#### import data ####
basedir="./"
humann3_mf_filtered_perm <- readRDS(file = paste0(basedir, "humann3_mf_filtered.rds"))
row.names(humann3_mf_filtered_perm) <- humann3_mf_filtered_perm$SampleName # make row
↪  names match data colnames

# Load the ShortBRED results, in which rows are antibiotic resistance markers and columns
↪  are samples
df.shortbred <- as.data.frame(read.table('~/Library/CloudStorage/Box-Box/Kau
↪  Lab/Results/MARS/FecalMetagenomics/shortbred/VFDB21results/MARS_SB_VF_RPKM.tsv',
↪  row.names=1, header=T))  # 12919 104
# head(df.shortbred)
length(rowSums(df.shortbred)[rowSums(df.shortbred) > 0]) # 616
```

```
## [1] 616
```

```
# names(df.shortbred) <- gsub(names(df.shortbred), pattern = "_SB_VF_RPKM",
↪  replacement="")
df.shortbredF <- filter_shortbred_data(indata = df.shortbred, my_suffix = "_SB_VF_RPKM",
↪  mymetadata = humann3_mf_filtered_perm)

dim(df.shortbredF) # 139 VFs
```

```
## [1] 139  95
```

```
write.csv(df.shortbredF, "SB_VF_RPKM_filtered.csv")
saveRDS(df.shortbredF, "SB_VF_RPKM_filtered.rds")
```

```
df.shortbredUF <- filter_shortbred_data(indata = df.shortbred, my_suffix = "_SB_VF_RPKM",
↪  mymetadata = humann3_mf_filtered_perm)
dim(df.shortbredUF) # 596 VFs
```

```
## [1] 139  95
```

```
write.csv(df.shortbredUF, "SB_VF_RPKM_unfiltered.csv") # dropped disqualifieds and all
↪  zeroes
saveRDS(df.shortbredUF, "SB_VF_RPKM_unfiltered.rds")

df.VFhits <- read.table('~/Library/CloudStorage/Box-Box/Kau
↪  Lab/Results/MARS/FecalMetagenomics/shortbred/VFDB21results/MARS_SB_VF_HITS.tsv',
↪  row.names=1, header=T)  # 12919 104
# head(df.VFhits)
# length(rowSums(df.VFhits)[rowSums(df.VFhits) > 0]) # 258
df.VFhitsF <- filter_shortbred_data(indata = df.VFhits, mymetadata =
↪  humann3_mf_filtered_perm, my_suffix = "_SB_VF_HITS")
write.csv(df.VFhitsF, "SB_VF_HITS_filtered.csv")
saveRDS(df.VFhitsF, "SB_VF_HITS_filtered.rds")
```

### Figure S4A:

PCA and Bray-Curtis Beta Diversity NMDS

```
source("./humann3_210613_functions_ngw.R")
rm(list=ls()[ls() %notin% c("%notin%", "makePCA", "makeNMDSplot", "makeAdonisBarplot",
↪  "boxplotByAsthmaAndAge")])

library(ggplot2)
library(ape)
library(stats)
library(vegan)
library(ggfortify)
library(viridis)
library(RColorBrewer)

basedir="./"
humann3_mf_filtered_perm <- readRDS(file = paste0(basedir, "humann3_mf_filtered.rds"))
row.names(humann3_mf_filtered_perm) <- humann3_mf_filtered_perm$SampleName
df.meta <- humann3_mf_filtered_perm

# use UNFILTERED DATA FOR ORDINATIONs # ******
df.shortbred <- readRDS("./SB_VF_RPKM_unfiltered.rds")

# Total sum square the data for Bray-curtis transform
df.shortbred.tss <- apply(X = df.shortbred, MARGIN = 2, FUN = function(x){x/sum(x)})

if(!all(names(df.shortbred.tss) == row.names(df.meta))){
  print("ERROR: meta data frame is not in the same order as sample data frame")
}

# NMDS
# sbRPKM.msds <- metaMDS(t(df.shortbred.tss), distance = "bray", k = 5, trymax = 1500,
↪  previous.best = TRUE) # Run 454 stress 0.08714864
```

```r
# saveRDS(sbRPKM.msds, "sbRPKM_brayCurtis.msds.rds")
# makeNMDSplot(data.msds = sbRPKM.msds,
#              mymetadata = df.meta,
#              my_data_table_name = "SB_RPKM",
#              mytitle = "ShortBRED TSS RPKM Bray-Curtis Unfiltered Data (N=95)",
#              basedir="./")

sbRPKM.msds <- readRDS("sbRPKM_brayCurtis.msds.rds")
sbRPKM_MDS <- as.data.frame(sbRPKM.msds$points)
sbRPKM_MDS$ASTHMA <- factor(df.meta[row.names(sbRPKM_MDS),"ASTHMA"], levels =
↪  c("Asthmatic", "Healthy"), c("Asthmatic", "Healthy"))
sbRPKM_MDS$AgeGroup <- factor(df.meta[row.names(sbRPKM_MDS), "AgeGroup"])
sbRPKM_MDS$AbxWithinYr <- factor(df.meta[row.names(sbRPKM_MDS), "AbxWithinYr"])
sbRPKM_MDS$Cohort <- paste0(sbRPKM_MDS$ASTHMA, " ", sbRPKM_MDS$AgeGroup)

# 4 grouper ####
my_color_scheme = c(brewer.pal(7, "BrBG")[c(7,1,5,3)])
confidenceEllipse = 0.95
mytitle = "Virulence Factor Gene Bray-Curtis Dissimilarity NMDS"
# library(viridis)
betaDivPlot4 <- ggplot(data = sbRPKM_MDS,
                        aes(x = MDS1, y = MDS2,
                            fill = Cohort,
                            color = Cohort,
                            shape = ASTHMA)) +
  stat_ellipse(aes(x=MDS1, y=MDS2, color=Cohort, group=Cohort), show.legend = TRUE, type
  ↪  = "t", level = confidenceEllipse, geom = "polygon", alpha = 0.4, size = 1)+
  # geom_point(size = 6, shape = 21, color ="black") +
  geom_point(size = 4.5, color = "black", alpha=0.8) +
    # geom_point(show.legend = FALSE, aes(brayCurtisMDS["MARS0043.F.16s", "MDS1"],
    ↪  brayCurtisMDS["MARS0043.F.16s", "MDS2"]),
    #            colour = asthma_color_hex, pch=5, size =8) +
    # geom_point(show.legend = FALSE, aes(brayCurtisMDS["MARS0022.F.16s", "MDS1"],
    ↪  brayCurtisMDS["MARS0022.F.16s", "MDS2"]),
    #            colour = healthy_color_hex, pch=5, size =8) +
    # scale_fill_viridis(breaks = levels(brayCurtisMDS$Cohort), discrete = TRUE, alpha =
    ↪  1,
    #                    labels = gsub(pattern = " ", replacement = "\n",
    #                                  x = levels(brayCurtisMDS$Cohort))) +
    # scale_fill_viridis(breaks = levels(brayCurtisMDS$Cohort), discrete = TRUE, alpha =
    ↪  1,
    #                    labels = gsub(pattern = " ", replacement = "\n",
    #                                  x = levels(brayCurtisMDS$Cohort))) +
    # scale_color_viridis(discrete = TRUE, alpha = 0.4) +
    scale_color_manual(values = my_color_scheme) +
    scale_fill_manual(values = my_color_scheme) +
    scale_shape_manual(values=c(21, 22)) +
    theme_classic() +
    xlab(label = "NMDS1") + ylab("NMDS2") + ggtitle(mytitle) +

    # geom_point(aes(shape=AgeGroup, color="black", size=6)) +
    # scale_shape_manual(values=c(21, 22)) +
```

```
    theme(axis.title.x = element_text(size = 21, vjust = -0.9),
          axis.title.y = element_text(size = 21, vjust = 1),
          axis.text.x = element_blank(),
          axis.ticks = element_blank(),
          axis.text.y = element_blank(),
          legend.text = element_text(size = 16),
          legend.title = element_text(size = 16, face = "bold"),
          legend.position = "bottom", legend.box = "horizontal")#,
  # plot.margin = margin(b = .9, l = 0.9, t = 1, r = 1, unit = "cm"))
  betaDivPlot4
```

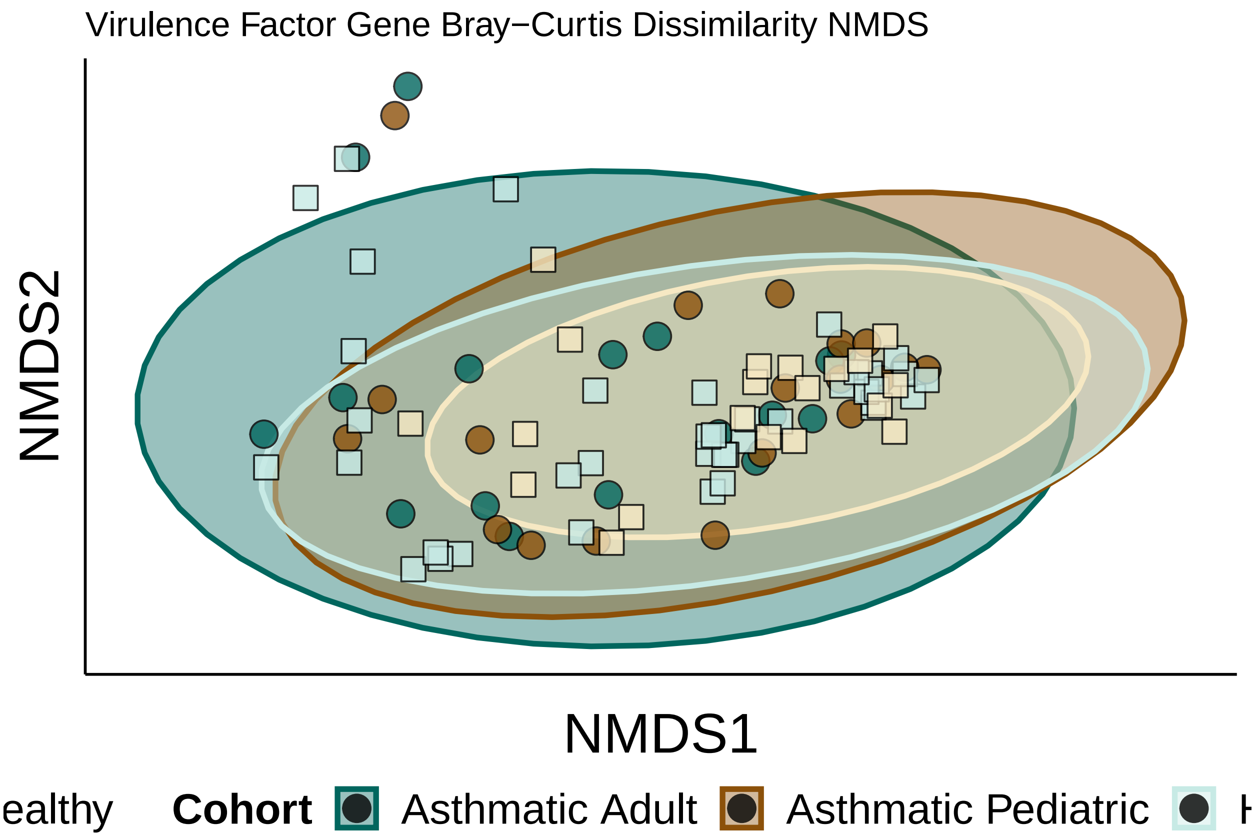


```
# ggsave(plot=betaDivPlot4, filename = paste0("./VF", "_4Group_NMDS_1and2.jpeg"), device
↪   = "jpg", width = 5, height = 5, units = "in")
# ggsave(plot=betaDivPlot4, filename = paste0("./VF", "_4Group_NMDS_1and2.pdf"), device =
↪   "pdf", width = 5, height = 5, units = "in", useDingbats = FALSE)
```

#### Figure S4B:

PERMANOVA Bray-Curtis distances barplot

```
rm(list=ls()[ls() %notin% c("%notin%", "makePCA", "makeNMDSplot", "makeAdonisBarplot",
↪   "boxplotByAsthmaAndAge")])

library(ggplot2)
library(ape)
library(stats)
```

```r
library(vegan)
library(ggfortify)

gene_analysis_name = "SB_RPKM_Bray_Curtis"
basedir_humann3="./"
humann3_mf_filtered_perm <- readRDS(file = paste0(basedir_humann3,
↪  "humann3_mf_filtered.rds"))
row.names(humann3_mf_filtered_perm) <- humann3_mf_filtered_perm$SampleName
df.meta <- humann3_mf_filtered_perm
df.shortbred <- readRDS("./SB_VF_RPKM_unfiltered.rds")

# Total sum square the data for Bray-curtis transform
df.shortbred.tss <- apply(X = df.shortbred, MARGIN = 2, FUN = function(x){x/sum(x)})

pca.dist =vegdist(t(df.shortbred.tss), method = "bray", binary = FALSE)
adonisAllIndpRaceBinaryPATHWAYS <- adonis(pca.dist ~
↪  readDepth+ASTHMA+AgeGroup+raceBinary+weightClassBinary+sex+subjectTobaccoUse+ASTHMA*AgeGroup+AbxWit[illegible]
                        data = df.meta,
                        permutations = 100000)

# make table
adonisAllIndpRaceBinaryPATHWAYSTable <-
↪  data.frame(adonisAllIndpRaceBinaryPATHWAYS$aov.tab[c("R2","Pr(>F)")])
names(adonisAllIndpRaceBinaryPATHWAYSTable) <- c("R2", "pval")
adonisAllIndpRaceBinaryPATHWAYSTable$features <-
↪  row.names(adonisAllIndpRaceBinaryPATHWAYSTable)
adonisAllIndpRaceBinaryPATHWAYSTable
↪  <-adonisAllIndpRaceBinaryPATHWAYSTable[!adonisAllIndpRaceBinaryPATHWAYSTable$features
↪  %in% c("Residuals","Total"),]
adonisAllIndpRaceBinaryPATHWAYSTable$pval <-
↪  round(adonisAllIndpRaceBinaryPATHWAYSTable$pval, 5)
adonisAllIndpRaceBinaryPATHWAYSTable$R2 <- round(adonisAllIndpRaceBinaryPATHWAYSTable$R2,
↪  3)
row.names(adonisAllIndpRaceBinaryPATHWAYSTable) <- NULL
adonisAllIndpRaceBinaryPATHWAYSTable <- adonisAllIndpRaceBinaryPATHWAYSTable[,c(3,1,2)]

basedir="./"
write.table(adonisAllIndpRaceBinaryPATHWAYSTable, file = paste0(basedir,
↪  gene_analysis_name, "_VFDBTSSRPKMadonisAllIndpRaceBinary100000"), sep="\t", row.names
↪  = FALSE, col.names = TRUE) #_filteredData.tsv

adonisAllIndpRaceBinaryPATHWAYSTable$neglogpval <-
↪  -log(adonisAllIndpRaceBinaryPATHWAYSTable$pval, base = 10)
adonisAllIndpRaceBinaryPATHWAYSTable$features <-
↪  factor(adonisAllIndpRaceBinaryPATHWAYSTable$features, levels=rev(c("readDepth",
↪  "ASTHMA", "AgeGroup", "raceBinary", "weightClassBinary", "sex", "subjectTobaccoUse",
↪  "AbxWithinYr",  "ASTHMA:AgeGroup")), labels = rev(c("Read Depth", "Asthma", "Age",
↪  "Race","Obesity", "Sex", "Tobacco Use", "Recent Antibiotics", "Asthma:Age")))
AdonisBarplotForPaperBinaryRacePATHWAYS <- makeAdonisBarplot(adonis_dat =
↪  adonisAllIndpRaceBinaryPATHWAYSTable, y_colname = "neglogpval", features_colname =
↪  "features", fill_colname = "R2", max_fill_value = 0.03)
print(AdonisBarplotForPaperBinaryRacePATHWAYS)
```

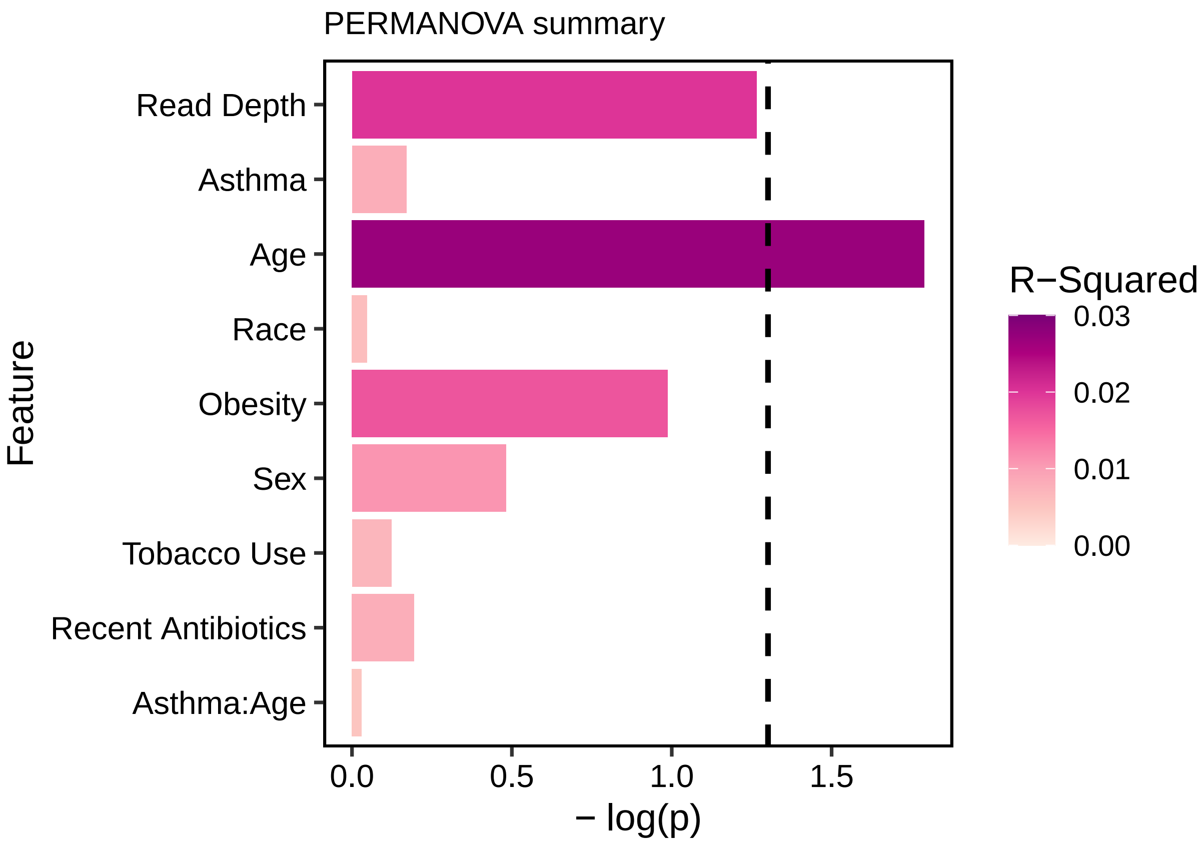


```
# ggsave(AdonisBarplotForPaperBinaryRacePATHWAYS,
       # filename = paste0("./", gene_analysis_name,
       ↪  "_VFDBTSSRPKMadonisAllIndpRaceBinary10000.pdf"),
       # #_adonisAllIndpRaceBinaryPATHWAYS100000_filteredData.jpg
       # height = 4, width = 5,
       # units = "in",
       # device = "pdf", useDingbats=FALSE)

# ggsave(AdonisBarplotForPaperBinaryRacePATHWAYS,
#        filename = paste0("./", gene_analysis_name,
↪  "_VFDBTSSRPKMadonisAllIndpRaceBinary100000.jpg"),
↪  #_adonisAllIndpRaceBinaryPATHWAYS100000_filteredData.jpg
#        height = 4, width = 5,
#        units = "in",
#        device = "jpg")
```

#### Figure S4C: Richness (non-zero rpkm)

```
rm(list=ls()[ls() %notin% c("%notin%", "unit_test1", "makePCA", "makeNMDSplot",
↪  "boxplotByAsthmaAndAge", "boxplotByAsthma")])
library(ggplot2)
library(RColorBrewer)
library(readxl)

basedir="./"
```

```r
humann3_mf_filtered_perm <- readRDS(file = paste0(basedir, "humann3_mf_filtered.rds"))
row.names(humann3_mf_filtered_perm) <- humann3_mf_filtered_perm$SampleName
df.meta <- humann3_mf_filtered_perm
df.shortbred <- readRDS("SB_VF_RPKM_filtered.rds")

#Next, generate a simple table that indicates the presence/absence of VF genes in each
↪  sample.
# Convert our data to presence/absence
df.shortbred <- readRDS("SB_VF_RPKM_filtered.rds")
df.shortbred.b <- df.shortbred
df.shortbred.b[df.shortbred.b!=0] <- 1
# collect "hits" per sample
hits <- data.frame(colSums(df.shortbred.b))
#hits <- hits[row.names(df.meta),]
hits$ASTHMA <- as.character(df.meta[row.names(hits), "ASTHMA"])
hits$AgeGroup <- as.character(df.meta[row.names(hits), "AgeGroup"])
hits$MARSID <- as.character(df.meta[row.names(hits), "MARSID"])
# table(hits$ASTHMA, hits$colSums.df.shortbred.b.)

# collect hits per ARG
hitsHealthy <- data.frame(rowSums(df.shortbred.b[, df.meta$ASTHMA=="Healthy"]))
hitsAsthma <- data.frame(rowSums(df.shortbred.b[, df.meta$ASTHMA=="Asthmatic"]))
names(hitsAsthma)<-"asthmaticHits"
names(hitsHealthy)<-"healthyHits"
IncidenceARGsByAsthma <- data.frame(hitsHealthy, hitsAsthma)
all(rowSums(IncidenceARGsByAsthma) == rowSums(df.shortbred.b)) # sanity check
```

```
## [1] TRUE
```

```r
IncidenceARGsByAsthma$ARGs <- row.names(IncidenceARGsByAsthma)

# Transpose the binary table
df.shortbred.b <- data.frame(t(df.shortbred.b))
# add metadata columns
df.shortbred.b$ASTHMA <- as.character(df.meta[row.names(df.shortbred.b), "ASTHMA"])
df.shortbred.b$AgeGroup <- as.character(df.meta[row.names(df.shortbred.b), "AgeGroup"])
df.shortbred.b$Sample <-  row.names(df.shortbred.b)
# head(df.shortbred.b)

# Melt the dataframe
library('reshape2')
df.shortbred.b.melted <- melt(df.shortbred.b, id.vars=c('ASTHMA', 'Sample'),
↪  value.name='Presence')
# Plot the presence/absence heatmap
library('ggplot2')
# pl <- ggplot(df.shortbred.b.melted, aes(x=Sample, y=variable, fill=Presence)) +
↪  geom_tile()
# pl <- pl + scale_x_discrete(labels=df.shortbred.b.melted$ASTHMA)
# pl <- pl + theme(axis.text.x = element_text(angle = 90, hjust = 1))
# plot(pl)

# add read depth
hits$readDepth <- as.numeric(df.meta[row.names(hits), "readDepth"]) # number of filtered
↪  reads (paired end sisters inclusive) that went into all post processing analyses
```

```
hits$coverage <- as.numeric(df.meta[row.names(hits), "coverage"]) # nonpareil metagenome
↪  coverage

wilcox.test(colSums.df.shortbred.b.~AgeGroup, hits)
```

```
##
##  Wilcoxon rank sum test with continuity correction
##
## data:  colSums.df.shortbred.b. by AgeGroup
## W = 1334, p-value = 0.06745
## alternative hypothesis: true location shift is not equal to 0
```

```
wilcox.test(colSums.df.shortbred.b.~ASTHMA, hits)
```

```
##
##  Wilcoxon rank sum test with continuity correction
##
## data:  colSums.df.shortbred.b. by ASTHMA
## W = 1266.5, p-value = 0.1173
## alternative hypothesis: true location shift is not equal to 0
```

```
# ANOVA: (https://www.r-bloggers.com/2011/03/anova-%E2%80%93-type-iiiiii-ss-explained/)
library(car)
summary(aov(colSums.df.shortbred.b.~ASTHMA*AgeGroup, data=hits)) #interaction p=0.6
```

```
##                  Df Sum Sq Mean Sq F value Pr(>F)
## ASTHMA            1    390   389.9   1.167  0.283
## AgeGroup          1   1465  1465.0   4.386  0.039 *
## ASTHMA:AgeGroup   1     70    70.3   0.210  0.648
## Residuals        91  30397   334.0
## ---
## Signif. codes:  0 '***' 0.001 '**' 0.01 '*' 0.05 '.' 0.1 ' ' 1
```

```
# Anova(lm(colSums.df.shortbred.b.~ASTHMA*AgeGroup, data=hits,
↪  contrasts=list(topic=contr.sum, sys=contr.sum)), type=2)
# with read depth or covreage
summary(aov(colSums.df.shortbred.b.~ASTHMA*readDepth+ASTHMA*AgeGroup, data=hits))
↪  #p=.9,.6
```

```
##                   Df Sum Sq Mean Sq F value Pr(>F)
## ASTHMA             1    390   389.9   1.144 0.2877
## readDepth          1    204   204.4   0.600 0.4407
## AgeGroup           1   1317  1316.8   3.863 0.0525 .
## ASTHMA:readDepth   1      0     0.1   0.000 0.9839
## ASTHMA:AgeGroup    1     76    76.3   0.224 0.6372
## Residuals         89  30335   340.8
## ---
## Signif. codes:  0 '***' 0.001 '**' 0.01 '*' 0.05 '.' 0.1 ' ' 1
```

```
Anova(lm(colSums.df.shortbred.b.~ASTHMA*readDepth+ASTHMA*AgeGroup, data=hits,
↪  contrasts=list(topic=contr.sum, sys=contr.sum)), type=2)
```

```
## Warning in model.matrix.default(mt, mf, contrasts): variable 'topic' is absent,
## its contrast will be ignored

## Warning in model.matrix.default(mt, mf, contrasts): variable 'sys' is absent,
```

```
## its contrast will be ignored
```

```
## Anova Table (Type II tests)
##
## Response: colSums.df.shortbred.b.
##                    Sum Sq Df F value  Pr(>F)
## ASTHMA              707.5  1  2.0757 0.15317
## readDepth            58.8  1  0.1725 0.67888
## AgeGroup           1316.9  1  3.8637 0.05246 .
## ASTHMA:readDepth      3.6  1  0.0107 0.91785
## ASTHMA:AgeGroup      76.3  1  0.2239 0.63723
## Residuals         30334.9 89
## ---
## Signif. codes:  0 '***' 0.001 '**' 0.01 '*' 0.05 '.' 0.1 ' ' 1
```

```r
Anova(lm(colSums.df.shortbred.b.~ASTHMA*coverage+ASTHMA*AgeGroup, data=hits,
↪  contrasts=list(topic=contr.sum, sys=contr.sum)), type=2)
```

```
## Warning in model.matrix.default(mt, mf, contrasts): variable 'topic' is absent,
## its contrast will be ignored
```

```
## Warning in model.matrix.default(mt, mf, contrasts): variable 'sys' is absent,
## its contrast will be ignored
```

```
## Anova Table (Type II tests)
##
## Response: colSums.df.shortbred.b.
##                   Sum Sq Df F value  Pr(>F)
## ASTHMA             689.3  1  2.1123 0.14963
## coverage          1239.4  1  3.7981 0.05446 .
## AgeGroup           887.5  1  2.7196 0.10265
## ASTHMA:coverage    115.1  1  0.3526 0.55416
## ASTHMA:AgeGroup     87.8  1  0.2690 0.60528
## Residuals        29042.9 89
## ---
## Signif. codes:  0 '***' 0.001 '**' 0.01 '*' 0.05 '.' 0.1 ' ' 1
```

```r
# Anova(lm(colSums.df.shortbred.b.~ASTHMA+readDepth+AgeGroup, data=hits,
↪  contrasts=list(topic=contr.sum, sys=contr.sum)), type=2)
# Anova(lm(colSums.df.shortbred.b.~ASTHMA+coverage+AgeGroup, data=hits,
↪  contrasts=list(topic=contr.sum, sys=contr.sum)), type=2)

library(ggbeeswarm)
library(ggpubr)
myPalette = brewer.pal(7, "BrBG")[c(7,5)] #c("red", "blue", "green", "purple")
myPaletteAge = brewer.pal(7, "BrBG")[c(1,3)]

# plot VFDB21hits
hits$value <- hits$colSums.df.shortbred.b.
# VFRPKMDerivedHitsPlot <- boxplotByAsthmaAndAge(mydat = hits)
# print(VFRPKMDerivedHitsPlot)
# # check with base R:
# myPalette=c("#01665E", "#01665E","#C7EAE5","#C7EAE5")
# boxplot(value~AgeGroup+ASTHMA, data = hits, at = c(1:2, 4:5), col = myPalette,
↪  main="sanity check")
#
```

```
# ggsave(plot = VFRPKMDerivedHitsPlot, filename =
↪  "./sb_VF_nonzeroRPKMHits_AgeGroups_boxplot.pdf", width = 5, height = 6, units = "in",
↪  useDingbats = FALSE)
# ggsave("./sb_VF_nonzeroRPKMHits_AgeGroups_boxplot.jpeg", plot = VFRPKMDerivedHitsPlot,
↪  units = "in", width = 5, height = 6, device = "jpg")

# # plot VFDB21hits - only asthma groups
# VFRPKMDerivedHitsPlotASTHMA <- boxplotByAsthma(mydat = hits)
# print(VFRPKMDerivedHitsPlotASTHMA)
# # check with base R:
# myPalette=c("#01665E", "#C7EAE5")
# boxplot(value~ASTHMA, data = hits, col=myPalette, main="sanity check")
# ggsave(plot = VFRPKMDerivedHitsPlotASTHMA, filename =
↪  "./sb_VF_nonzeroRPKMHits_Asthma_boxplot.pdf", width = 5, height = 6, units = "in",
↪  useDingbats = FALSE)
# ggsave(filename = "./sb_VF_nonzeroRPKMHits_Asthma_boxplot.jpeg", plot =
↪  VFRPKMDerivedHitsPlotASTHMA, units = "in", width = 5, height = 6, device = "jpg")

# violins?
my_color_scheme = c(brewer.pal(7, "BrBG")[c(7,1,5,3)])
# VIOLIN PLOTS
source("/Users/naomiwilson/Library/CloudStorage/Box-Box/Kau
↪  Lab/Results/MARS/FecalMetagenomics/shortbred/CARD2021results/split_violins_ggplot.R")
VFrichnessViolin <- ggplot(data = hits, aes(x=ASTHMA, y=value, fill=
↪  interaction(ASTHMA,AgeGroup))) +
  geom_violin(aes(x=ASTHMA, y=value, fill=ASTHMA), draw_quantiles = c(0.5), inherit.aes =
  ↪  FALSE, scale="width", trim=FALSE, alpha=0.8, show.legend = T) + # draw_quantiles =
  ↪  c(0.5), c(0.25, 0.5, 0.75)
  geom_split_violin(draw_quantiles = c(0.5), scale="width", width=0.5, trim=FALSE,
  ↪  alpha=0.8, show.legend = T) +  # add this to draw quantiles: draw_quantiles =
  ↪  c(0.25, 0.5, 0.75)
  theme_classic() +
  scale_fill_manual(name="Cohort",
                    values = c("#01665E", brewer.pal(7, "BrBG")[1], brewer.pal(7,
                    ↪  "BrBG")[3],
                               "#C7EAE5", brewer.pal(7, "BrBG")[1], brewer.pal(7,
                               ↪  "BrBG")[3]),
                    labels = c('Asthmatic','Adult', 'Pediatric',
                               'Healthy', 'Adult', 'Pediatric'))+ # color order:
                               ↪  A,AA,AP,H,HA,HP
  theme(panel.grid.major = element_blank(),
        panel.grid.minor = element_blank(),
        axis.line.x = element_line(size = 0, colour = "black"),
        axis.line.y = element_line(size = 0, colour = "black"),
        axis.line = element_line(size=1, colour = "black"),
        panel.border = element_rect(fill = NA, colour = "black", size = 1),
        panel.background = element_blank(),
        text=element_text(size = 16),
        axis.text.x=element_text(colour="black", size = 12),
        axis.text.y=element_text(colour="black", size = 12),
        axis.title.x=element_blank())+
        # legend.position = "none")+
  ylab("Number of Unique virulence factor genes")
```

```
VFrichnessViolin
```

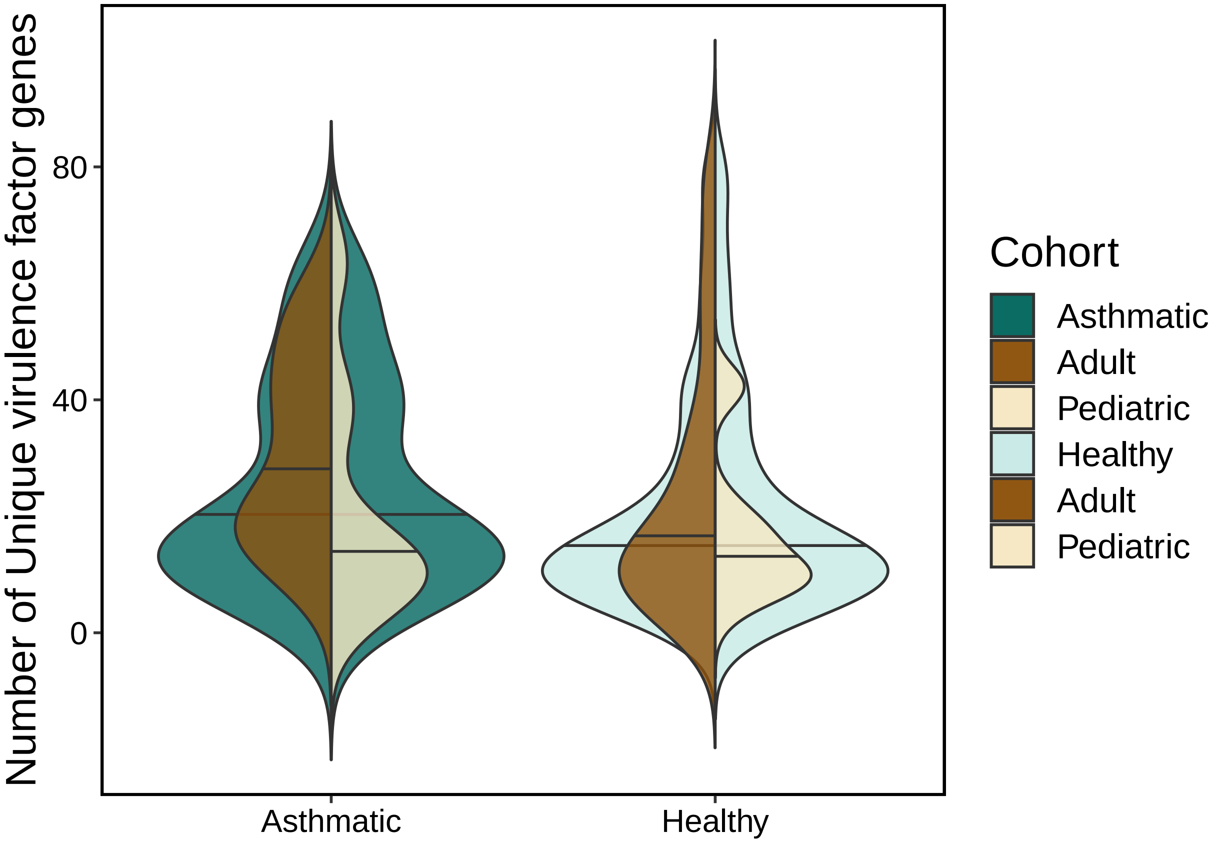


```
# ggsave(plot = VFrichnessViolin, filename = "./VF_Richness_doubleviolin.pdf", width = 6,
↪  height = 5, units = "in", useDingbats = FALSE)
# ggsave(filename = "./VF_Richness_doubleviolin.jpeg", plot = VFrichnessViolin, units =
↪  "in", width =6, height = 6, device = "jpg")
```

#### Figure 3C, 5B, S4D (procrustes)

format data

```
rm(list=ls())
CARD.file <- readRDS(file = "./SB_CARD21_with_AMRFam_ARO_Drug_Mechanism_filtered.rds")
CARD.genes <- CARD.file[!grepl(pattern = "ARO_3003577", row.names(CARD.file)),-c(96, 97,
↪  98, 99, 100, 101, 102)] # DROP UGD because it's also a VF
saveRDS(CARD.genes, file = "CARD.genes.rds")

VF.file <- readRDS(file = "~/Library/CloudStorage/Box-Box/Kau
↪  Lab/Results/MARS/FecalMetagenomics/shortbred/VFDB21results/VF_RPKM_with_drugclass_filtered.rds")
VF.genes <- VF.file[,-c(96, 97, 98, 99)]
if(!all(names(VF.genes) == names(CARD.genes))){
  VF.genes <- VF.genes[, as.character(names(CARD.genes))]
}
saveRDS(VF.genes, file = "VF.genes.rds")
```

```
metaphlan.species <- readRDS("./allSpecies_95_relab_tab.rds")
# sum(colSums(t(metaphlan.species)) == 0)
metaphlan.species <- metaphlan.species[rowSums(metaphlan.species) > 0.0001,] # filter out
↪  < 0.0001 relab
names(metaphlan.species) <- gsub(names(metaphlan.species), pattern = "[.]", replacement =
↪  "-")
if(!all(names(metaphlan.species) == names(CARD.genes))){
  metaphlan.species <- metaphlan.species[, as.character(names(CARD.genes))]
}
saveRDS(metaphlan.species, file = "metaphlan.species.rds")
```

plotting function

```
ariel_plot_edit <- function(pro, protest.res, mycolnames=c("first.1", "first.2",
↪  "second.1", "second.2"),
                            my_pallette_fill=c(RColorBrewer::brewer.pal(7,
                             ↪  "BrBG")[c(7,5)]),
                            my_pallete_col=c(RColorBrewer::brewer.pal(7,
                             ↪  "BrBG")[c(7,5)]),
                            mytitle=""){
  humann3_mf_filtered <- readRDS("./humann3_mf_filtered.rds")
  row.names(humann3_mf_filtered) <- humann3_mf_filtered$SampleName

  require(ggplot2)
  #Make This Into A GGPlot2 Figure.
  ggpro = data.frame(cbind(pro$X[,c(1,2)], pro$Yrot[,c(1,2)])) # grab first 2 dimensions
  colnames(ggpro) = mycolnames
  #bind in Sample Data
  ggpro[as.character(humann3_mf_filtered$SampleName),"ASTHMA"] =
↪  factor(as.character(humann3_mf_filtered$ASTHMA))
  gg_procrustes = ggplot(data = ggpro) +
    geom_segment(aes(x = second.1 , y = second.2, xend = first.1, yend =first.2 , color =
    ↪  ASTHMA), arrow = arrow(length = unit(0.5, "cm")), size = 1) +
    geom_point(mapping = aes(x = second.1, y = second.2, fill = ASTHMA, shape=ASTHMA),#,
               color = "black", size = 4) +
    theme_classic() +
    scale_shape_manual(values = c(21, 22)) +
    scale_fill_manual(values = my_pallete_col) +
    scale_color_manual(values = my_pallete_col, guide = 'none') +
    # guides(fill = guide_legend(override.aes = list(shape = 21))) +
    xlab(label = "PCoA1") +
    ylab(label = "PCoA2") +
    ggtitle(mytitle, subtitle = paste0("PROTEST Corr: ", round(protest.res$t0, digits =
    ↪  3), "\np-value = ", protest.res$signif)) +
   theme(text = element_text(family="Helvetica"),
         axis.title.x = element_text(size = 18, vjust = -1),
         axis.title.y = element_text(size = 18, vjust = 1),
         axis.text.x = element_blank(),
         axis.line = element_line(size = 1.0),
         axis.ticks = element_blank(),
         axis.text.y = element_blank(),
         legend.text = element_text(size = 14),
         legend.title = element_text(size = 12, face = "bold"),
         # legend.position = "none", legend.box = "verticle",
```

```
          plot.margin = margin(b = .9, l = 0.9, t = 1, r = 1, unit = "cm"))
  return(gg_procrustes)
}
```

PROCRUSTES on Hellinger transform + Bray-Curtis

```
source("./humann3_210613_functions_ngw.R")
rm(list=ls()[ls() %notin% c("ariel_plot_edit", "%notin%")])
# trying to make this analysis match the infant e coli asthma gut metagenomics paper:
# https://www.sciencedirect.com/science/article/pii/S1931312821001451?via%3Dihub#bib28
# They ran Hellinger transform on their relative abundance tables (it's just sqrt of the
↪  TSS) and then did bray-curtis in vegan

library(vegan)

CARD.genes <- readRDS(file = "CARD.genes.rds")
CARD.genes.hell <- decostand(t(CARD.genes), method = "hellinger")
CARD.genes.hell.bray <- vegdist(CARD.genes.hell, method = "bray")
# CARD.genes.bray.msds <- metaMDS(t(CARD.genes), distance = "bray", k = 4, trymax = 500)
CARD.genes.pcoa <- ape::pcoa(CARD.genes.hell.bray)
# plot(CARD.genes.pcoa$vectors[,1], CARD.genes.pcoa$vectors[,2], main="CARD")

VF.genes <- readRDS(file = "VF.genes.rds")
VF.genes.hell <- decostand(t(VF.genes), method = "hellinger")
VF.genes.hell.bray <- vegdist(VF.genes.hell, method = "bray")
VF.genes.pcoa = ape::pcoa(VF.genes.hell.bray)
# VF.genes.bray.msds <- metaMDS(VF.genes, distance = "bray", k = 10, trymax = 500)
# plot(VF.genes.pcoa$vectors[,1], VF.genes.pcoa$vectors[,2], main="VF")

metaphlan.species <- readRDS(file = "metaphlan.species.rds")
metaphlan.species.hell <- decostand(t(metaphlan.species), method = "hellinger")
metaphlan.species.hell.bray <- vegdist(metaphlan.species.hell, method = "bray")
metaphlan.species.pcoa <- ape::pcoa(metaphlan.species.hell.bray)
# plot(metaphlan.species.pcoa$vectors[,1], metaphlan.species.pcoa$vectors[,2],
↪  main="Metaphlan")

if (dim(metaphlan.species)[2]==dim(VF.genes)[2] & dim(VF.genes)[2] ==dim(CARD.genes)[2])
↪  {
  print("CARD v VF")
  pro1a.pca <- procrustes(X = CARD.genes.pcoa$vectors, Y = VF.genes.pcoa$vectors, scores
↪  = "sites", symmetric = TRUE)
  pro1a.protest <- protest(X = CARD.genes.pcoa$vectors, Y = VF.genes.pcoa$vectors, scores
↪  = "sites", permutations = how(nperm = 9999))
  # plot(pro1a.pca, main="CARD v VF")
  # text(x = 0.1, y=0.04, paste("corr= ", round(pro1a.protest$t0, digits = 3)))
  # text(x = 0.1, y=-0.04, paste("p= ", pro1a.protest$signif))

  print("Metaphlan v CARD")
  pro2.pca <- procrustes(X = metaphlan.species.pcoa$vectors, Y = CARD.genes.pcoa$vectors,
↪  scores = "sites", symmetric = TRUE)
  pro2.protest <- protest(X = metaphlan.species.pcoa$vectors, Y =
↪  CARD.genes.pcoa$vectors, scores = "sites", permutations = how(nperm = 9999))
  # plot(pro2.pca, main="metaphlan v CARD")
  # text(x = 0.1, y=0.04, paste("corr= ", round(pro2.protest$t0, digits = 3)))
```

```
  # text(x = 0.1, y=-0.04, paste("p= ", pro2.protest$signif))
  pro2.protest$signif

  print("Metaphlan v VF")
  pro3.pca <- procrustes(X = metaphlan.species.pcoa$vectors, Y = VF.genes.pcoa$vectors,
↪   scores = "sites", symmetric = TRUE)
  pro3.protest <- protest(X = metaphlan.species.pcoa$vectors, Y = VF.genes.pcoa$vectors,
↪   scores = "sites", permutations = how(nperm = 9999))
  # plot(pro3.pca, main="metaphlan v VF", kind=1)
  # text(x = 0.1, y=0.04, paste("corr= ", round(pro3.protest$t0, digits = 3)))
  # text(x = 0.1, y=-0.04, paste("p= ", pro3.protest$signif))
  pro3.protest$signif

  print(ariel_plot_edit(pro = pro1a.pca, protest.res = pro1a.protest, mytitle = "ARG v.
   ↪   VF"))
  print(ariel_plot_edit(pro = pro2.pca, protest.res = pro2.protest, mytitle = "Species
   ↪   Composition v. ARG"))
  print(ariel_plot_edit(pro = pro3.pca, protest.res = pro3.protest, mytitle = "Species
   ↪   Composition v. VF"))

  # ggsave(plot = ariel_plot_edit(pro = pro1a.pca, protest.res = pro1a.protest, mytitle =
   ↪   "ARG v. VF"), filename = "Procrustes_ARG_VF.pdf", device = "pdf",
   ↪   useDingbats=FALSE)
  # ggsave(plot = ariel_plot_edit(pro = pro2.pca, protest.res = pro2.protest, mytitle =
   ↪   "Species Composition v. ARG"), filename = "Procrustes_SpeciesComposition_ARG.pdf",
   ↪   device = "pdf", useDingbats=FALSE)
  # ggsave(plot = ariel_plot_edit(pro = pro3.pca, protest.res = pro3.protest, mytitle =
   ↪   "Species Composition v. VF"), filename = "Procrustes_SpeciesComposition_VF.pdf",
   ↪   device = "pdf", useDingbats=FALSE)


  pro1list <- c(pro1a.protest$t0, pro1a.protest$signif, pro1a.protest$ss, "ARG v. VF")
  pro2list <- c(pro2.protest$t0, pro2.protest$signif, pro2.protest$ss, "Species
↪   Composition v. ARG")
  pro3list <- c(pro3.protest$t0, pro3.protest$signif, pro3.protest$ss, "Species
↪   Composition v. VF")

  saveme <- data.frame(rbind(pro1list, pro2list, pro3list))
  names(saveme) <- c("protest_corr", "protest_p", "protest_sumofsquares", "comparison")
  saveme$type <- "Both Cohorts Hellinger transformed Bray-Curtis"
  # pro1a.protest$signif
  # pro2.protest$signif
  # pro3.protest$signif
}
```

```
## [1] "CARD v VF"

## Warning in procrustes(X = CARD.genes.pcoa$vectors, Y = VF.genes.pcoa$vectors, : X has fewer axes than

## Warning in procrustes(X, Y, symmetric = FALSE): X has fewer axes than Y: X adjusted to comform Y

## [1] "Metaphlan v CARD"
## [1] "Metaphlan v VF"
```

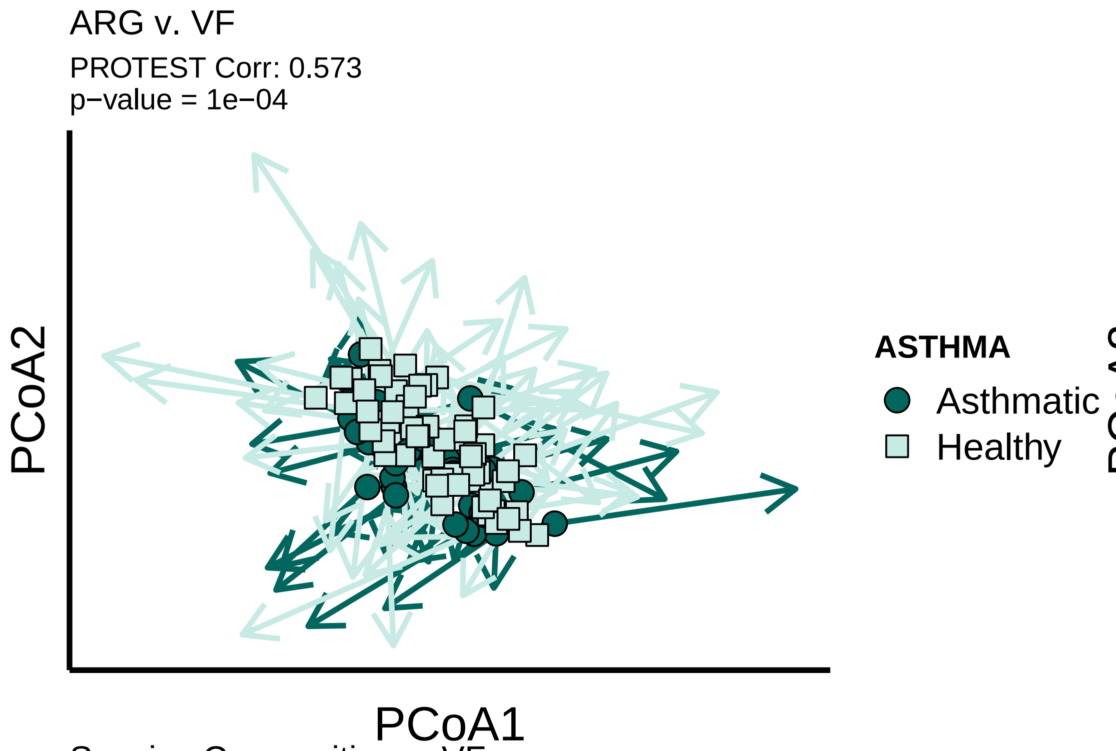


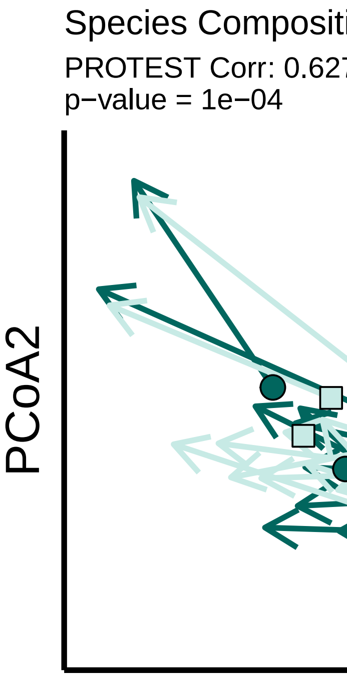


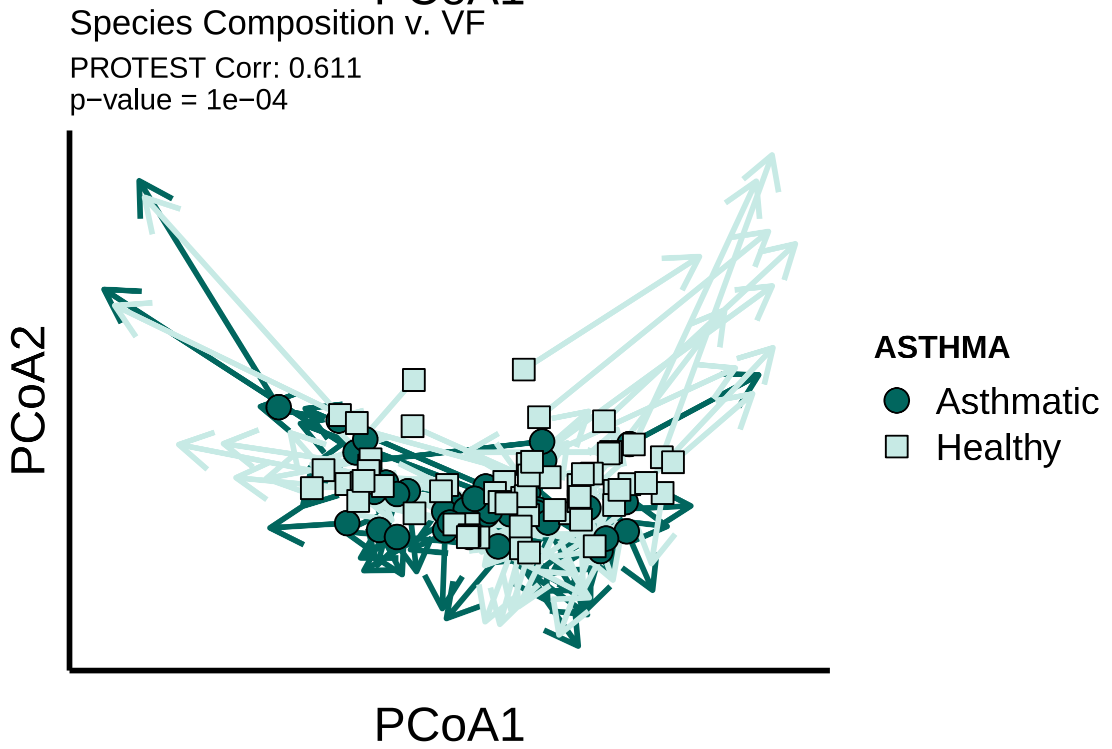


```
saveRDS(saveme, "hellingerBC_protest_results.rds")
```

##### VFs vs. ARGs co-occurrence family-level

ARGs by AMR Family based on CARD, VFs by Family but then binned if they had the same or similar name (ex. "T6SS-II (CVF861)" and "T6SS (VF0579)")

```
rm(list=ls()[ls() %notin% c("unit_test1", "%notin%", "makePCA", "makeNMDSplot",
↪  "boxplotByAsthmaAndAge", "boxplotByAsthma")])
source("./humann3_210613_functions_ngw.R")

library(tidyverse)
library(gplots)
library(RColorBrewer)
library(binovisualfields)
```

```
##
## Attaching package: 'binovisualfields'

## The following object is masked from 'package:ape':
##
##     rotate

## The following object is masked from 'package:ggpubr':
##
##     rotate
```

```
group_by_me = "Cohort"
my_pair_test = "kruskal_test"
my_padjust_method="fdr"
summarized_by = "FAMILYSUM"
my_color_scheme = brewer.pal(7, "BrBG")[c(7,5)]

# Family Rel abund:
df.VFDB <- readRDS("~/Library/CloudStorage/Box-Box/Kau
↪  Lab/Results/MARS/FecalMetagenomics/shortbred/VFDB21results/VFDB21_RPKMSummedByFamilyBins.rds")

df.CARD <- readRDS("./SB_CARD21_with_AMRFam_ARO_Drug_Mechanism_filtered.rds")

# Drop ugd in CARD
row.names(df.CARD)[grep(row.names(df.CARD), pattern = "ARO_3003577")]
```

```
## [1] "gb|AAC75089_1|ARO_3003577|ugd__Escherichia_coli_str__K-12_substr__MG1655_"
```

```
df.CARD <- df.CARD[!(row.names(df.CARD) %in% row.names(df.CARD)[grep(row.names(df.CARD),
↪  pattern = "ARO_3003577")]),]

basedir="./"
humann3_mf_filtered_perm <- readRDS(file = paste0(basedir, "humann3_mf_filtered.rds"))
row.names(humann3_mf_filtered_perm) <- humann3_mf_filtered_perm$SampleName
df.meta <- humann3_mf_filtered_perm
df.meta$Cohort <- factor(paste0(df.meta$ASTHMA, " ", df.meta$AgeGroup))
df.meta$weightClassAgeGroup <- factor(paste0(df.meta$AgeGroup, " ",
↪  df.meta$weightClassBinary))

# df.CARD
# get family summed
aggfun <- function(x, mydata=df.CARD){
  df=aggregate(. ~ AMRGeneFamily, FUN = sum, mydata[,c("AMRGeneFamily",
↪  as.character(x))])
  return(data.frame(df))
}
```

```r
# sanitycheck these are equal:
# aggfun("TAAGGCGA-CTCTCTAT_S112")
# library("tidyverse")
# data.frame(df.CARD[, c("TAAGGCGA-CTCTCTAT_S112", "AMRGeneFamily")] %>%
↪  group_by(AMRGeneFamily) %>% summarise_all("sum"))
# df.CARD %>% group_by(AMRGeneFamily) %>% summarise_all("sum")
df.CARD.Summed <- lapply(X = names(df.CARD)[1:95], FUN = aggfun)
df.CARD.Summed <- bind_cols(df.CARD.Summed, .id = "column_label")
```

```
## New names:
## * `AMRGeneFamily` -> `AMRGeneFamily...1`
## * `AMRGeneFamily` -> `AMRGeneFamily...3`
## * `AMRGeneFamily` -> `AMRGeneFamily...5`
## * `AMRGeneFamily` -> `AMRGeneFamily...7`
## * `AMRGeneFamily` -> `AMRGeneFamily...9`
## * `AMRGeneFamily` -> `AMRGeneFamily...11`
## * `AMRGeneFamily` -> `AMRGeneFamily...13`
## * `AMRGeneFamily` -> `AMRGeneFamily...15`
## * `AMRGeneFamily` -> `AMRGeneFamily...17`
## * `AMRGeneFamily` -> `AMRGeneFamily...19`
## * `AMRGeneFamily` -> `AMRGeneFamily...21`
## * `AMRGeneFamily` -> `AMRGeneFamily...23`
## * `AMRGeneFamily` -> `AMRGeneFamily...25`
## * `AMRGeneFamily` -> `AMRGeneFamily...27`
## * `AMRGeneFamily` -> `AMRGeneFamily...29`
## * `AMRGeneFamily` -> `AMRGeneFamily...31`
## * `AMRGeneFamily` -> `AMRGeneFamily...33`
## * `AMRGeneFamily` -> `AMRGeneFamily...35`
## * `AMRGeneFamily` -> `AMRGeneFamily...37`
## * `AMRGeneFamily` -> `AMRGeneFamily...39`
## * `AMRGeneFamily` -> `AMRGeneFamily...41`
## * `AMRGeneFamily` -> `AMRGeneFamily...43`
## * `AMRGeneFamily` -> `AMRGeneFamily...45`
## * `AMRGeneFamily` -> `AMRGeneFamily...47`
## * `AMRGeneFamily` -> `AMRGeneFamily...49`
## * `AMRGeneFamily` -> `AMRGeneFamily...51`
## * `AMRGeneFamily` -> `AMRGeneFamily...53`
## * `AMRGeneFamily` -> `AMRGeneFamily...55`
## * `AMRGeneFamily` -> `AMRGeneFamily...57`
## * `AMRGeneFamily` -> `AMRGeneFamily...59`
## * `AMRGeneFamily` -> `AMRGeneFamily...61`
## * `AMRGeneFamily` -> `AMRGeneFamily...63`
## * `AMRGeneFamily` -> `AMRGeneFamily...65`
## * `AMRGeneFamily` -> `AMRGeneFamily...67`
## * `AMRGeneFamily` -> `AMRGeneFamily...69`
## * `AMRGeneFamily` -> `AMRGeneFamily...71`
## * `AMRGeneFamily` -> `AMRGeneFamily...73`
## * `AMRGeneFamily` -> `AMRGeneFamily...75`
## * `AMRGeneFamily` -> `AMRGeneFamily...77`
## * `AMRGeneFamily` -> `AMRGeneFamily...79`
## * `AMRGeneFamily` -> `AMRGeneFamily...81`
## * `AMRGeneFamily` -> `AMRGeneFamily...83`
## * `AMRGeneFamily` -> `AMRGeneFamily...85`
```

```
## * `AMRGeneFamily` -> `AMRGeneFamily...87`
## * `AMRGeneFamily` -> `AMRGeneFamily...89`
## * `AMRGeneFamily` -> `AMRGeneFamily...91`
## * `AMRGeneFamily` -> `AMRGeneFamily...93`
## * `AMRGeneFamily` -> `AMRGeneFamily...95`
## * `AMRGeneFamily` -> `AMRGeneFamily...97`
## * `AMRGeneFamily` -> `AMRGeneFamily...99`
## * `AMRGeneFamily` -> `AMRGeneFamily...101`
## * `AMRGeneFamily` -> `AMRGeneFamily...103`
## * `AMRGeneFamily` -> `AMRGeneFamily...105`
## * `AMRGeneFamily` -> `AMRGeneFamily...107`
## * `AMRGeneFamily` -> `AMRGeneFamily...109`
## * `AMRGeneFamily` -> `AMRGeneFamily...111`
## * `AMRGeneFamily` -> `AMRGeneFamily...113`
## * `AMRGeneFamily` -> `AMRGeneFamily...115`
## * `AMRGeneFamily` -> `AMRGeneFamily...117`
## * `AMRGeneFamily` -> `AMRGeneFamily...119`
## * `AMRGeneFamily` -> `AMRGeneFamily...121`
## * `AMRGeneFamily` -> `AMRGeneFamily...123`
## * `AMRGeneFamily` -> `AMRGeneFamily...125`
## * `AMRGeneFamily` -> `AMRGeneFamily...127`
## * `AMRGeneFamily` -> `AMRGeneFamily...129`
## * `AMRGeneFamily` -> `AMRGeneFamily...131`
## * `AMRGeneFamily` -> `AMRGeneFamily...133`
## * `AMRGeneFamily` -> `AMRGeneFamily...135`
## * `AMRGeneFamily` -> `AMRGeneFamily...137`
## * `AMRGeneFamily` -> `AMRGeneFamily...139`
## * `AMRGeneFamily` -> `AMRGeneFamily...141`
## * `AMRGeneFamily` -> `AMRGeneFamily...143`
## * `AMRGeneFamily` -> `AMRGeneFamily...145`
## * `AMRGeneFamily` -> `AMRGeneFamily...147`
## * `AMRGeneFamily` -> `AMRGeneFamily...149`
## * `AMRGeneFamily` -> `AMRGeneFamily...151`
## * `AMRGeneFamily` -> `AMRGeneFamily...153`
## * `AMRGeneFamily` -> `AMRGeneFamily...155`
## * `AMRGeneFamily` -> `AMRGeneFamily...157`
## * `AMRGeneFamily` -> `AMRGeneFamily...159`
## * `AMRGeneFamily` -> `AMRGeneFamily...161`
## * `AMRGeneFamily` -> `AMRGeneFamily...163`
## * `AMRGeneFamily` -> `AMRGeneFamily...165`
## * `AMRGeneFamily` -> `AMRGeneFamily...167`
## * `AMRGeneFamily` -> `AMRGeneFamily...169`
## * `AMRGeneFamily` -> `AMRGeneFamily...171`
## * `AMRGeneFamily` -> `AMRGeneFamily...173`
## * `AMRGeneFamily` -> `AMRGeneFamily...175`
## * `AMRGeneFamily` -> `AMRGeneFamily...177`
## * `AMRGeneFamily` -> `AMRGeneFamily...179`
## * `AMRGeneFamily` -> `AMRGeneFamily...181`
## * `AMRGeneFamily` -> `AMRGeneFamily...183`
## * `AMRGeneFamily` -> `AMRGeneFamily...185`
## * `AMRGeneFamily` -> `AMRGeneFamily...187`
## * `AMRGeneFamily` -> `AMRGeneFamily...189`
```

```r
# df.CARD.Summed <- data.frame(df.CARD.Summed)
df.CARD.Summed <- data.frame(df.CARD.Summed[,1], df.CARD.Summed[,
↪  c(gsub(names(df.CARD)[1:95], pattern = "-", replacement = "."))])
names(df.CARD.Summed)[1] <- "AMRGeneFamily"
names(df.CARD.Summed)[2:96] <- gsub(names(df.CARD.Summed)[2:96],
                                              pattern = "[.]",
                                              replacement = "-")
row.names(df.CARD.Summed) <- df.CARD.Summed$AMRGeneFamily
df.CARD.Summed$AMRGeneFamily <- NULL

# use family summed:
df.CARD <- df.CARD.Summed

library("cooccur")
df.VFDB <- df.VFDB[, names(df.CARD)]
if (all(names(df.VFDB)[1:95] == names(df.CARD)[1:95])) {
  mymat <- apply(rbind(df.VFDB[,1:95], df.CARD[,1:95]), MARGIN = 2, FUN = as.numeric)
  row.names(mymat) <- c(row.names(df.VFDB[,1:95]), row.names(df.CARD[,1:95]))
  mymat.bin <- mymat
  mymat.bin[mymat.bin>0]=1 # make binary
  myresults.cooccur <- cooccur(mymat.bin, spp_names = TRUE)
}
```

```
##   |                                                                              |
```

```r
summary(myresults.cooccur)
```

```
## Call:
## cooccur(mat = mymat.bin, spp_names = TRUE)
##
## Of 2145 species pair combinations, 107 pairs (4.99 %) were removed from the analysis because expecte
##
## Cooccurrence Summary:
```

```
##         Species          Sites       Positive       Negative         Random
##              66             95            333             13           1692
## Unclassifiable Non-random (%)
##               0             17
## attr(,"class")
## [1] "summary.cooccur"
```

```r
# plot(myresults.cooccur)
# print(myresults.cooccur)
# dim(print(myresults.cooccur)) # 353  11

# ggsave(plot(myresults.cooccur)+ggtitle("VF v ARG co-occurrence"), filename =
↪  "VFvARG_summedByFamily_allCooccurPlot.jpg", device = "jpeg" , width =10, height = 10)
# ggsave(plot(myresults.cooccur)+ggtitle("VF v ARG co-occurrence"), filename =
↪  "VFvARG_summedByFamily_allCooccurPlot.pdf", device = "pdf" , width =10, height = 10)

family.df <-  myresults.cooccur$results
# sp1 Numeric label giving the identity of species 1, assigned based on the order inthe
↪  input matrix
# sp2 Numeric label for species
# sp1_inc Number of sites (or samples) that have species
```

```r
# sp2_inc Number of sites that have species
# obs cooccur Observed number of sites having both species
# prob cooccur Probability that both species occur at a site
# exp cooccur Expected number of sites having both species
# p_lt Probability  that  the  two  species  would  co-occur  at  a  frequency  less
↪  than  the observed number of co-occurrence sites if the two species were distributed
↪  ran-domly (independently) of one another
# p_gt Probability of co-occurrence at a frequency greater than the observed frequency
# sp1_name If  species  names  were  specified  in  the  community  data  matrix  this
↪  field  willcontain the supplied name of sp1
# sp2_name The supplied name of sp2

family.df$sp1type <- ifelse(family.df$sp1_name %in% row.names(df.VFDB[,1:95]), "VF",
↪  ifelse(family.df$sp1_name %in% row.names(df.CARD[,1:95]), "ARG", 0))
family.df$sp2type <- ifelse(family.df$sp2_name %in% row.names(df.VFDB[,1:95]), "VF",
↪  ifelse(family.df$sp2_name %in% row.names(df.CARD[,1:95]), "ARG", 0))
# family.df[(family.df$p_gt < 0.05 | family.df$p_lt < 0.05),]
family.df.sig <- family.df[(family.df$p_gt < 0.05 | family.df$p_lt < 0.05),]

family.df.VFvARG <- family.df[(family.df$sp1type == "VF" & family.df$sp2type == "ARG"),]
family.df.ARGvVF <- family.df[(family.df$sp1type == "ARG" & family.df$sp2type == "VF"),]
↪  # none of these, so is not reversible
family.df.filt <- rbind(family.df.VFvARG, family.df.ARGvVF)
family.df.filt.sig <- family.df.filt[(family.df.filt$p_gt < 0.05 | family.df.filt$p_lt <
↪  0.05),]
# family.df.filt.sig[order(family.df.filt.sig$prob_cooccur, decreasing = TRUE),]

# top positively co-occurring:
# View(family.df.filt.sig[order(family.df.filt.sig$p_gt, decreasing = FALSE),][1:25,
↪  c("p_gt", "p_lt","prob_cooccur", "obs_cooccur", "sp1_name", "sp2_name")])
# top negatively co-occurring:
# View(family.df.filt.sig[family.df.filt.sig$p_lt<0.05,][, c("p_gt",
↪  "p_lt","prob_cooccur", "obs_cooccur", "sp1_name", "sp2_name")])

saveRDS(object = myresults.cooccur, file =
↪  "VFvARG_summedByFamily_results.cooccurobject.rds")
saveRDS(object = family.df.filt[order(family.df.filt$prob_cooccur, decreasing = TRUE),],
↪  file = "VFvARG_summedByFamily_cooccur.rds")
write.table(x = family.df.filt.sig[order(family.df.filt.sig$prob_cooccur, decreasing =
↪  TRUE),], file = "VFvARG_summedByFamily_cooccur_significant.txt")

# ASTHMA v HEALTHY FAMILY ####
df.VFDB.asthma <- df.VFDB[,names(df.VFDB)[names(df.VFDB) %in%
↪  row.names(df.meta[df.meta$ASTHMA=="Asthmatic",])]]
df.CARD.asthma <- df.CARD.Summed[,names(df.CARD.Summed)[names(df.CARD.Summed) %in%
↪  row.names(df.meta[df.meta$ASTHMA=="Asthmatic",])]]

df.VFDB.healthy <- df.VFDB[,names(df.VFDB)[names(df.VFDB) %in%
↪  row.names(df.meta[df.meta$ASTHMA=="Healthy",])]]
df.CARD.healthy <- df.CARD.Summed[,names(df.CARD.Summed)[names(df.CARD.Summed) %in%
↪  row.names(df.meta[df.meta$ASTHMA=="Healthy",])]]

if (all(names(df.VFDB.asthma) == names(df.CARD.asthma))) {
```

```
  mymat <- apply(rbind(df.VFDB.asthma, df.CARD.asthma), MARGIN = 2, FUN = as.numeric)
  row.names(mymat) <- c(row.names(df.VFDB.asthma), row.names(df.CARD.asthma))
  mymat.bin <- mymat
  mymat.bin[mymat.bin>0]=1 # make binary
  asthma.myresults.cooccur <- cooccur::cooccur(mymat.bin, spp_names = TRUE)
}
```

```
##   |                                                                              |
```

```
# print(asthma.myresults.cooccur)
# plt.asthma <- plot(asthma.myresults.cooccur) + ggtitle("Asthmatic VF v ARG")
# ggsave(plt.asthma, filename = "~/Library/CloudStorage/Box-Box/Kau
↪  Lab/Results/MARS/FecalMetagenomics/shortbred/VFvARG_summedByFamily_asthmaticCooccurPlot.jpg",
↪  device = "jpeg" , width =10, height = 10)
# ggsave(plt.asthma, filename = "~/Library/CloudStorage/Box-Box/Kau
↪  Lab/Results/MARS/FecalMetagenomics/shortbred/VFvARG_summedByFamily_asthmaticCooccurPlot.pdf",
↪  device = "pdf" , width =10, height = 10)

if (all(names(df.VFDB.healthy) == names(df.CARD.healthy))) {
  mymat <- apply(rbind(df.VFDB.healthy, df.CARD.healthy), MARGIN = 2, FUN = as.numeric)
  row.names(mymat) <- c(row.names(df.VFDB.healthy), row.names(df.CARD.healthy))
  mymat.bin <- mymat
  mymat.bin[mymat.bin>0]=1 # make binary
  healthy.myresults.cooccur <- cooccur::cooccur(mymat.bin, spp_names = TRUE)
}
```

```
##   |                                                                              |
```

```
# print(healthy.myresults.cooccur)
# plt.healthy <- plot(healthy.myresults.cooccur) + ggtitle("Healthy VF v ARG")
# ggsave(plt.healthy, filename = "~/Library/CloudStorage/Box-Box/Kau
↪  Lab/Results/MARS/FecalMetagenomics/shortbred/VFvARG_summedByFamily_healthyCooccurPlot.jpg",
↪  device = "jpeg" , width =10, height = 10)
# ggsave(plt.healthy, filename = "~/Library/CloudStorage/Box-Box/Kau
↪  Lab/Results/MARS/FecalMetagenomics/shortbred/VFvARG_summedByFamily_healthyCooccurPlot.pdf",
↪  device = "pdf" , width =10, height = 10)

saveRDS(object = healthy.myresults.cooccur, file =
↪  "./VFvARG_summedByFamily_HEALTHY.cooccurobject.rds")
saveRDS(object = asthma.myresults.cooccur, file =
↪  "./VFvARG_summedByFamily_ASTHMATICS.cooccurobject.rds")
write.table(x = healthy.myresults.cooccur$results, file =
↪  "./VFvARG_summedByFamily_HEALTHY_cooccur.txt")
write.table(x = asthma.myresults.cooccur$results, file =
↪  "./VFvARG_summedByFamily_ASTHMATICS_cooccur.txt")

# homemade plots ####
results <- asthma.myresults.cooccur$results
results$pBins <- ifelse(results$p_gt < 0.05, "Pos", ifelse(results$p_lt < 0.05, "Neg",
↪  "Neither"))
```

length with ugd –> dropped ugd: 353 family.df.sig –> 346 1042 family.df.VFvARG –> 1042

#### Compare healthy v asthma ARGvVF co-occurrence

```
rm(list=ls())
source("./humann3_210613_functions_ngw.R")
library(tidyverse)
healthy.myresults.cooccur <- readRDS(file =
↪  "./VFvARG_summedByFamily_HEALTHY.cooccurobject.rds")
asthma.myresults.cooccur <- readRDS(file = "~/Library/CloudStorage/Box-Box/Kau
↪  Lab/Results/MARS/FecalMetagenomics/shortbred/VFvARG_summedByFamily_ASTHMATICS.cooccurobject.rds")
all(healthy.myresults.cooccur$spp_key == asthma.myresults.cooccur$spp_key) # names match
↪  numbered labels
```

```
## [1] TRUE
```

```
makeCategoricalSigColumn <- function(tmp.df) {
  tmp.df$sig <- ifelse(tmp.df$p_gt < 0.05, "Positive",
                       ifelse(tmp.df$p_lt < 0.05, "Negative", "Random"))
  return(tmp.df)
}

healthy.myresults.cooccur.df <-
↪  makeCategoricalSigColumn(data.frame(healthy.myresults.cooccur$results))
# healthy.myresults.cooccur.df$effect_sizes <- effect.sizes(healthy.myresults.cooccur)
healthy.myresults.cooccur.df=merge(x=healthy.myresults.cooccur.df, y =
↪  effect.sizes(healthy.myresults.cooccur), by.x = c("sp1_name", "sp2_name"), by.y =
↪  c("sp1", "sp2"))
asthma.myresults.cooccur.df <-
↪  makeCategoricalSigColumn(data.frame(asthma.myresults.cooccur$results))
asthma.myresults.cooccur.df=merge(x=asthma.myresults.cooccur.df, y =
↪  effect.sizes(asthma.myresults.cooccur), by.x = c("sp1_name", "sp2_name"), by.y =
↪  c("sp1", "sp2"))

# names of comparisons do not match, so let's make our own key: ####
# all are 65 features long
length(levels(factor(healthy.myresults.cooccur.df$sp2_name)))
```

```
## [1] 65
```

```
length(levels(factor(healthy.myresults.cooccur.df$sp1_name)))
```

```
## [1] 65
```

```
length(levels(factor(asthma.myresults.cooccur.df$sp2_name)))
```

```
## [1] 65
```

```
length(levels(factor(asthma.myresults.cooccur.df$sp1_name)))
```

```
## [1] 65
```

```
sum(levels(factor(healthy.myresults.cooccur.df$sp1_name)) %in%
↪  levels(factor(asthma.myresults.cooccur.df$sp1_name)))
```

```
## [1] 65
```

```
sum(unique(healthy.myresults.cooccur.df$sp1_name) %in%
↪  unique(asthma.myresults.cooccur.df$sp1_name))
```

```
## [1] 65
```

```
# but there's a discrepancy in matrix length
dim(healthy.myresults.cooccur.df) # 1830
```

```
## [1] 1822   13
```

```
dim(asthma.myresults.cooccur.df) # 1798
```

```
## [1] 1794   13
```

```
# there's also a discrepancy in pairs, i.e. there aren't an  exact 32 missing, there are
↪  several unique to each matrix:
# create pair columns:
asthma.myresults.cooccur.df$comp_name <- paste(asthma.myresults.cooccur.df$sp1_name,
↪  asthma.myresults.cooccur.df$sp2_name, sep = "|v|")
asthma.myresults.cooccur.df$comp_name_rev <- paste(asthma.myresults.cooccur.df$sp2_name,
↪  asthma.myresults.cooccur.df$sp1_name, sep = "|v|") # this is to check if the 2
↪  matrices have pairs that were named differently
healthy.myresults.cooccur.df$comp_name <- paste(healthy.myresults.cooccur.df$sp1_name,
↪  healthy.myresults.cooccur.df$sp2_name, sep = "|v|")
healthy.myresults.cooccur.df$comp_name_rev <-
↪  paste(healthy.myresults.cooccur.df$sp2_name, healthy.myresults.cooccur.df$sp1_name,
↪  sep = "|v|") # this is to check if the 2 matrices have pairs that were named
↪  differently

# in asthma but not in healthy:
length(unique(asthma.myresults.cooccur.df$comp_name)[unique(asthma.myresults.cooccur.df$comp_name)
↪  %notin% unique(healthy.myresults.cooccur.df$comp_name)]) # 98
```

```
## [1] 103
```

```
length(unique(asthma.myresults.cooccur.df$comp_name)[unique(asthma.myresults.cooccur.df$comp_name)
↪  %in% unique(healthy.myresults.cooccur.df$comp_name_rev)]) # 0 (means order of vf v
↪  arg was ok)
```

```
## [1] 0
```

```
# in healthy but not in asthma:
length(unique(healthy.myresults.cooccur.df$comp_name)[unique(healthy.myresults.cooccur.df$comp_name)
↪  %notin% unique(asthma.myresults.cooccur.df$comp_name)]) # 130
```

```
## [1] 131
```

```
length(unique(healthy.myresults.cooccur.df$comp_name)[unique(healthy.myresults.cooccur.df$comp_name)
↪  %in% unique(asthma.myresults.cooccur.df$comp_name_rev)]) # 0 (means order of vf v arg
↪  was ok)
```

```
## [1] 0
```

```
# so some pairs were filtered out for each matrix, presumably because of the "thresh"
↪  option in cooccur which drops pairs whose expected co-occurrence is less than 1.
uniqueAsthmaPairs <-
↪  unique(asthma.myresults.cooccur.df$comp_name)[unique(asthma.myresults.cooccur.df$comp_name)
↪  %notin% unique(healthy.myresults.cooccur.df$comp_name)] # 98
```

```r
uniqueHealthyPairs <-
↪  unique(healthy.myresults.cooccur.df$comp_name)[unique(healthy.myresults.cooccur.df$comp_name)
↪  %notin% unique(asthma.myresults.cooccur.df$comp_name)] # 130

# drop these from both matrices:
asthma.myresults.cooccur.df.filt <-
↪  asthma.myresults.cooccur.df[asthma.myresults.cooccur.df$comp_name %notin%
↪  uniqueAsthmaPairs,]
healthy.myresults.cooccur.df.filt <-
↪  healthy.myresults.cooccur.df[healthy.myresults.cooccur.df$comp_name %notin%
↪  uniqueHealthyPairs,]

dim(healthy.myresults.cooccur.df.filt) == dim(asthma.myresults.cooccur.df.filt)
```

```
## [1] TRUE TRUE
```

```r
sum(asthma.myresults.cooccur.df.filt$comp_name %in%
↪  healthy.myresults.cooccur.df.filt$comp_name) ==
↪  dim(healthy.myresults.cooccur.df.filt)[1]
```

```
## [1] TRUE
```

```r
# NOW COMPARE HvA: ####
row.names(healthy.myresults.cooccur.df.filt) <-
↪  healthy.myresults.cooccur.df.filt$comp_name
row.names(asthma.myresults.cooccur.df.filt) <- asthma.myresults.cooccur.df.filt$comp_name

if(sum(asthma.myresults.cooccur.df.filt$comp_name ==
↪  healthy.myresults.cooccur.df.filt$comp_name) ==
↪  dim(healthy.myresults.cooccur.df.filt)[1]){
  results.df <- data.frame(cbind(healthy.myresults.cooccur.df.filt$sig,
↪  asthma.myresults.cooccur.df.filt$sig))
  names(results.df) <- c("Healthy", "Asthma")
  row.names(results.df) <- healthy.myresults.cooccur.df.filt$comp_name
  results.df$match <- ifelse(results.df$Healthy == results.df$Asthma, 1, 0)
}

allMismatches <- row.names(results.df)[results.df$match ==0] # 176 mismatches
AsthmaPos <- row.names(results.df)[results.df$match ==0 & results.df$Asthma ==
↪  "Positive"] #29 (1 v Negative)
HealthyPos <- row.names(results.df)[results.df$match ==0 & results.df$Healthy ==
↪  "Positive"] #120 (all v Random)
AsthmaNeg <- row.names(results.df)[results.df$match ==0 & results.df$Asthma ==
↪  "Negative"] #14 (all v Random)
HealthyNeg <- row.names(results.df)[results.df$match ==0 & results.df$Healthy ==
↪  "Negative"] #14 (1 v Positive)

row.names(results.df)[results.df$match ==0 & results.df$Asthma == "Negative" &
↪  results.df$Healthy == "Positive"] #0
```

```
## character(0)
```

```r
row.names(results.df)[results.df$match ==0 & results.df$Asthma == "Positive" &
↪  results.df$Healthy == "Negative"] #1
```

```
## [1] "RcsAB (VF0571)|v|glycopeptide resistance gene cluster;vanR"
```

```
# row.names(results.df)[results.df$match ==0]
# asthma.myresults.cooccur.df[asthma.myresults.cooccur.df$comp_name %in% AsthmaPos,]

#filter out VFvVF and ARGvARG
# healthy.myresults.cooccur$spp_key[1:34,] # ALL VFs
# healthy.myresults.cooccur$spp_key[35:66,] # ALL ARGs

VFvARG.df <- expand.grid(healthy.myresults.cooccur$spp_key[1:34,"spp"],
↪  healthy.myresults.cooccur$spp_key[35:66,"spp"])
VFvARG_all_possible <- paste(VFvARG.df$Var1, VFvARG.df$Var2, sep = "|v|")
asthma.myresults.cooccur.df.filt.no.incest <-
↪  asthma.myresults.cooccur.df.filt[asthma.myresults.cooccur.df.filt$comp_name %in%
↪  VFvARG_all_possible,]
healthy.myresults.cooccur.df.filt.no.incest <-
↪  healthy.myresults.cooccur.df.filt[healthy.myresults.cooccur.df.filt$comp_name %in%
↪  VFvARG_all_possible,]

# Plot these and highlight which way the mismatch goes (sig positive/neg in asthma or
↪  healthy):
#
↪  asthma.myresults.cooccur.df.filt.no.incest[asthma.myresults.cooccur.df.filt.no.incest$comp_name
↪  %in% allMismatches,]
#
↪  healthy.myresults.cooccur.df.filt.no.incest[healthy.myresults.cooccur.df.filt.no.incest$comp_name
↪  %in% allMismatches,]

saveRDS(asthma.myresults.cooccur.df.filt.no.incest,
↪  "./asthma.myresults.cooccur.df.filt.no.incest.rds")
saveRDS(healthy.myresults.cooccur.df.filt.no.incest,
↪  "./healthy.myresults.cooccur.df.filt.no.incest.rds")
saveRDS(asthma.myresults.cooccur.df, "./asthma.myresults.cooccur.df.unfiltered.rds")
saveRDS(healthy.myresults.cooccur.df, "./healthy.myresults.cooccur.df.unfiltered.rds")
```

#### Make incidence matrix for CARD to use for cooccurrence that drops genes matching from VFDB

```
rm(list=ls())
# GET CARD STUFF ####
df.CARD.F <- readRDS(file = "./SB_CARD21_with_AMRFam_ARO_Drug_Mechanism_filtered.rds")
#DROP UGD since it shows up as a VF ****
df.CARD.F <- df.CARD.F[!(grepl(pattern = "ARO_3003577", row.names(df.CARD.F))),]
# most taxa are in the rownames of this DF
taxalist <- comprehenr::to_vec(for (i in row.names(df.CARD.F)) strsplit(i, "__")[[1]][2])
taxalist <- gsub(pattern = "_$", "", taxalist)
taxalist <- gsub(pattern = "_", " ", taxalist)
df.CARD.F$Source <- as.character(taxalist)
#make combined name column
df.CARD.F$familySource <- paste(df.CARD.F$AMRGeneFamily, df.CARD.F$Source, sep="__")
df.CARD.F$OG_rows <- row.names(df.CARD.F)
# get short gene names to match hash table
row.names(df.CARD.F) <- gsub(pattern = "__)", "'')", row.names(df.CARD.F))
```

```
row.names(df.CARD.F) <- gsub(pattern = "_)", "')", row.names(df.CARD.F))
shortnamelist <- comprehenr::to_vec(for (i in comprehenr::to_vec(for (i in
↪  row.names(df.CARD.F)) strsplit(i, "__[A-Z]")[[1]][1])) strsplit(i, "[|]")[[1]][4])
shortnamelist <- gsub(pattern = "_$", "", shortnamelist)
shortnamelist <- gsub(pattern = "_", " ", shortnamelist)
shortnamelist <- comprehenr::to_vec(for (i in shortnamelist) strsplit(i, "  ")[[1]][1]) #
↪  gets rid of extra text like "uncultured bacterium"

df.CARD.F$geneShortName <- shortnamelist
row.names(df.CARD.F) <- df.CARD.F$geneShortName

# count incidence
# make binary
df.CARD.F.bin <- df.CARD.F
df.CARD.F.bin[,1:95][df.CARD.F.bin[,1:95]>0] = 1

saveRDS(df.CARD.F.bin, "./df.CARD.F.bin.rds")
```

### Figure 5A:

Partial Correlations on richness

```
rm(list=ls())
source("./humann3_210613_functions_ngw.R")
# library(tidyverse)
library(ggplot2)
library(RColorBrewer)

basedir="./"
humann3_mf_filtered_perm <- readRDS(file = paste0(basedir, "humann3_mf_filtered.rds"))
row.names(humann3_mf_filtered_perm) <- humann3_mf_filtered_perm$SampleName
df.meta <- humann3_mf_filtered_perm

df.CARD.F.bin <- readRDS("./df.CARD.F.bin.rds")
df.shortbred.VFgene.Incidence <-
↪  readRDS("/Users/naomiwilson/Library/CloudStorage/Box-Box/Kau
↪  Lab/Results/MARS/FecalMetagenomics/shortbred/VFandARG21/VFvARG_taxa_constrained/datatables/MARS_VFD

uniref90_dat <- readRDS(file=paste0(basedir, "uniref90CPM", ".plotdat.rds"))
row.names(df.meta) <- df.meta$filenamePeriod
names(uniref90_dat) <- as.character(df.meta[as.character(names(uniref90_dat)),
↪  "SampleName"])
all(as.character(names(uniref90_dat)) %in% names(df.CARD.F.bin[,1:95]))
```

```
## [1] TRUE
```

```
row.names(df.meta) <- df.meta$SampleName
uniref90_dat <- uniref90_dat[, names(df.CARD.F.bin[,1:95])]
all(as.character(names(uniref90_dat)) == names(df.CARD.F.bin[,1:95]))
```

```
## [1] TRUE
```

```
all(as.character(names(uniref90_dat)) == names(df.shortbred.VFgene.Incidence[,1:95]))
```

```
## [1] FALSE
```

```r
df.shortbred.VFgene.Incidence <- df.shortbred.VFgene.Incidence[,
↪  as.character(names(df.CARD.F.bin[,1:95]))]
uniref90_dat.bin <- uniref90_dat
uniref90_dat.bin[uniref90_dat.bin>0]=1

# combine data:
dat <- cbind(colSums(df.shortbred.VFgene.Incidence[,1:95]), colSums(uniref90_dat.bin))
dat <- data.frame(cbind(dat, colSums(df.CARD.F.bin[, 1:95])))
names(dat) <- c("VF", "uniref90", "ARG")
# install.packages("ppcor")
library(ppcor)
dat_asthma <- dat
dat_asthma$ASTHMA <- df.meta[row.names(dat_asthma), "ASTHMA"]

dat_asthmatic <-dat_asthma[dat_asthma$ASTHMA=="Asthmatic", c("VF", "uniref90", "ARG")]
dat_healthy <- dat_asthma[dat_asthma$ASTHMA=="Healthy", c("VF", "uniref90", "ARG")]

pcor(dat) # defaults to pearson
```

```
## $estimate
##                  VF   uniref90       ARG
## VF        1.0000000 -0.2526939 0.7156145
## uniref90 -0.2526939  1.0000000 0.2336995
## ARG       0.7156145  0.2336995 1.0000000
##
## $p.value
##                    VF   uniref90          ARG
## VF       0.000000e+00 0.01400264 5.283680e-16
## uniref90 1.400264e-02 0.00000000 2.339185e-02
## ARG      5.283680e-16 0.02339185 0.000000e+00
##
## $statistic
##                 VF  uniref90      ARG
## VF        0.000000 -2.505053 9.826741
## uniref90 -2.505053  0.000000 2.305406
## ARG       9.826741  2.305406 0.000000
##
## $n
## [1] 95
##
## $gp
## [1] 1
##
## $method
## [1] "pearson"
```

```r
pcor(dat_asthmatic)
```

```
## $estimate
##                  VF    uniref90        ARG
## VF        1.0000000 -0.20112039 0.70057051
## uniref90 -0.2011204  1.00000000 0.08748953
## ARG       0.7005705  0.08748953 1.00000000
```

```
##
## $p.value
##                    VF  uniref90          ARG
## VF       0.000000e+00 0.2466530 2.789546e-06
## uniref90 2.466530e-01 0.0000000 6.172458e-01
## ARG      2.789546e-06 0.6172458 0.000000e+00
##
## $statistic
##                 VF   uniref90       ARG
## VF        0.000000 -1.1794489 5.6398066
## uniref90 -1.179449  0.0000000 0.5045237
## ARG       5.639807  0.5045237 0.0000000
##
## $n
## [1] 36
##
## $gp
## [1] 1
##
## $method
## [1] "pearson"
```

```
pcor(dat_healthy)
```

```
## $estimate
##                  VF   uniref90       ARG
## VF        1.0000000 -0.2715487 0.7186397
## uniref90 -0.2715487  1.0000000 0.2850089
## ARG       0.7186397  0.2850089 1.0000000
##
## $p.value
##                    VF   uniref90          ARG
## VF       0.000000e+00 0.03921212 2.124092e-10
## uniref90 3.921212e-02 0.00000000 3.012013e-02
## ARG      2.124092e-10 0.03012013 0.000000e+00
##
## $statistic
##                 VF  uniref90      ARG
## VF        0.000000 -2.111421 7.733594
## uniref90 -2.111421  0.000000 2.225097
## ARG       7.733594  2.225097 0.000000
##
## $n
## [1] 59
##
## $gp
## [1] 1
##
## $method
## [1] "pearson"
```

```
# summarise
results.all <- data.frame(rbind(pcor(dat)$estimate[upper.tri(pcor(dat)$estimate)],
↪  pcor(dat)$p.value[upper.tri(pcor(dat)$p.value)]), row.names = c("estimate",
↪  "p.value"))
```

```r
names(results.all) <- c("UniRefvVF", "VFvARG", "UniRefvARG")
results.all["estimate_asthma",] <-
↪  pcor(dat_asthmatic)$estimate[upper.tri(pcor(dat_asthmatic)$estimate)]
results.all["p.value_asthma",] <-
↪  pcor(dat_asthmatic)$p.value[upper.tri(pcor(dat_asthmatic)$p.value)]
results.all["estimate_healthy",] <-
↪  pcor(dat_healthy)$estimate[upper.tri(pcor(dat_healthy)$estimate)]
results.all["p.value_healthy",] <-
↪  pcor(dat_healthy)$p.value[upper.tri(pcor(dat_healthy)$p.value)]
results.all
```

```
##                    UniRefvVF       VFvARG UniRefvARG
## estimate         -0.25269387 7.156145e-01 0.23369949
## p.value           0.01400264 5.283680e-16 0.02339185
## estimate_asthma  -0.20112039 7.005705e-01 0.08748953
## p.value_asthma    0.24665297 2.789546e-06 0.61724584
## estimate_healthy -0.27154865 7.186397e-01 0.28500886
## p.value_healthy   0.03921212 2.124092e-10 0.03012013
```

```r
# cor.test(dat)

m1 <- lm(ARG~uniref90, data = dat)  #Create a linear model
m1_resid <- resid(m1) #List of residuals
m2 <- lm(VF~uniref90, data = dat)  #Create a linear model
m2_resid <- resid(m2) #List of residuals
# plot(m1_resid, m2_resid)
# cor.test(m1_resid, m2_resid)

m1a <- lm(ARG~uniref90, data = dat_asthmatic)  #Create a linear model
m1_resida <- resid(m1a) #List of residuals
m2a <- lm(VF~uniref90, data = dat_asthmatic)  #Create a linear model
m2_resida <- resid(m2a) #List of residuals
# plot(m1_resida, m2_resida)
# cor.test(m1_resida, m2_resida)

m1h <- lm(ARG~uniref90, data = dat_healthy)  #Create a linear model
m1_residh <- resid(m1h) #List of residuals
m2h <- lm(VF~uniref90, data = dat_healthy)  #Create a linear model
m2_residh <- resid(m2h) #List of residuals
# plot(m1_residh, m2_residh)
# cor.test(m1_residh, m2_residh)

residuals.all <- data.frame(cbind(m1_resida, m2_resida))
names(residuals.all) <- c("ARG", "VF")
residuals.all$asthma <- rep("Asthmatic", dim(residuals.all)[1])
healthy <- data.frame(cbind(m1_residh, m2_residh, rep("Healthy", length(m2_residh))))
names(healthy) <- c("ARG", "VF", "asthma")
residuals.all <- rbind(residuals.all, healthy)
residuals.all$ARG <- as.numeric(residuals.all$ARG)
residuals.all$VF <- as.numeric(residuals.all$VF)
library(RColorBrewer)
residuals.all.p <- ggplot(residuals.all, aes(x=ARG, y=VF, color=asthma, fill=asthma)) +
    geom_smooth(method="lm", se=TRUE, fullrange=FALSE, level=0.95, lty=2)+#auto makes
     ↪  squiggly loess line
```

```
    geom_point(size=4) + theme_classic() +
    scale_color_manual(values = c(brewer.pal(11, "BrBG")[c(10,7)])) +
    scale_fill_manual(values = c(brewer.pal(11, "BrBG")[c(10,7)])) +
    annotate("text", x=-10, y=20, label= paste("Asthmatic Pearson \np=",
    ↪  format(cor.test(x = residuals.all[residuals.all$asthma == "Asthmatic","VF"],
    ↪  residuals.all[residuals.all$asthma == "Asthmatic","ARG"], method =
    ↪  "pearson")$p.value, scientific=TRUE), "\n", round(cor.test(x =
    ↪  residuals.all[residuals.all$asthma == "Asthmatic","VF"],
    ↪  residuals.all[residuals.all$asthma == "Asthmatic","ARG"], method =
    ↪  "pearson")$estimate, digits = 2)), col=c(brewer.pal(11, "BrBG")[c(10)])) +
    annotate("text", x=10, y=-25, label= paste("Healthy Pearson \np=", format(cor.test(x
    ↪  = residuals.all[residuals.all$asthma == "Healthy","VF"],
    ↪  residuals.all[residuals.all$asthma == "Healthy","ARG"], method =
    ↪  "pearson")$p.value, scientific=TRUE), "\n", round(cor.test(x =
    ↪  residuals.all[residuals.all$asthma == "Healthy","VF"],
    ↪  residuals.all[residuals.all$asthma == "Healthy","ARG"], method =
    ↪  "pearson")$estimate, digits = 2)), col=c(brewer.pal(11, "BrBG")[c(7)]))

residuals.all.p <- residuals.all.p + ggtitle("Residuals accounting for gene richness") +
↪  labs(x="ARG Residuals", y="VF residuals")
# ggsave(plot = residuals.all.p, filename =
↪  "./VFvARG_richness_partial_correlations_residuals_scatterplot.pdf", device = "pdf",
↪  units = "in", useDingbats=FALSE) #Saving 4.79 x 5.47 in image
# ggsave(plot = residuals.all.p, filename =
↪  "./VFvARG_richness_partial_correlations_residuals_scatterplot.jpg", device = "jpeg",
↪  units = "in")

residuals.all.p
```

```
## `geom_smooth()` using formula 'y ~ x'
```

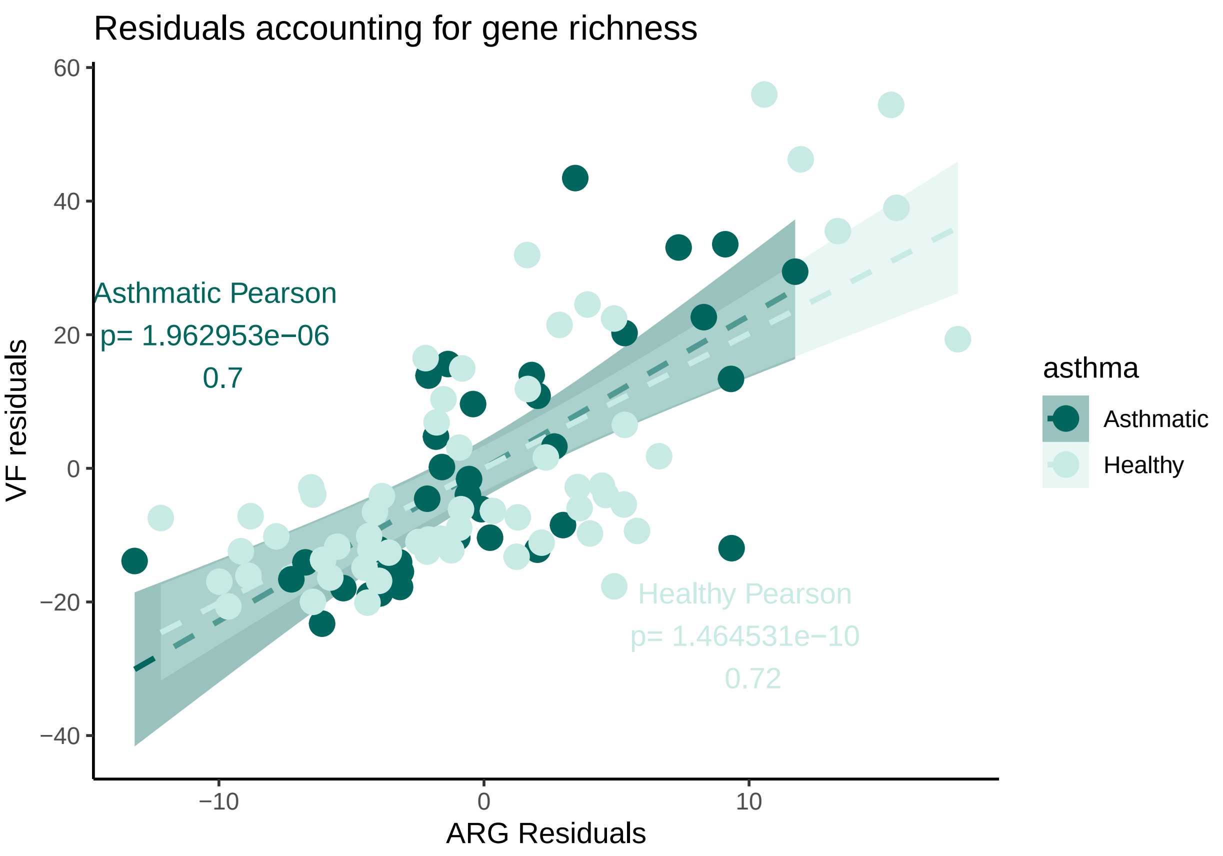


#### Figure 5C:

Heatmap all co-occurrences dropping non-sig rows/cols

```
rm(list=ls())
source("./humann3_210613_functions_ngw.R")
library(tidyverse)
library(ggplot2)
library(RColorBrewer)
asthma.myresults.cooccur.df.filt.no.incest <-
↪   readRDS("./asthma.myresults.cooccur.df.filt.no.incest.rds")
healthy.myresults.cooccur.df.filt.no.incest <-
↪   readRDS("./healthy.myresults.cooccur.df.filt.no.incest.rds")
VF_RPKM_with_names <- readRDS("~/Library/CloudStorage/Box-Box/Kau
↪   Lab/Results/MARS/FecalMetagenomics/shortbred/VFDB21results/VF_RPKM_with_VFnames_FamilyBins_filtered
ARG_PRKM_with_names <- readRDS("./SB_CARD21_with_AMRFam_ARO_Drug_Mechanism_filtered.rds")
# add supergroups for grouping on heatmap in an intuitive way:
ARG_PRKM_with_names$SUPERGROUP <- ifelse(grepl(ARG_PRKM_with_names$AMRGeneFamily, pattern
↪   = "lactamase"), "5bla",
                                         ifelse(grepl(ARG_PRKM_with_names$AMRGeneFamily,
                                         ↪   pattern = "efflux pump"), "1pump",
                                                
                                                ↪   ifelse(grepl(ARG_PRKM_with_names$AMRGeneFamily,
                                                ↪   pattern = "glycopeptide"),
                                                ↪   "4glycopeptide",
                                                       
                                                       ↪   ifelse(grepl(ARG_PRKM_with_names$AMRGeneFami
                                                       ↪   pattern = "tetracycline"),
                                                       ↪   "6tet",
```

```
                                                                    ↪  ifelse(grepl(ARG_PRKM_with_names$AMRG
                                                                    ↪  pattern =
                                                                    ↪  "transferase"),
                                                                    ↪  "2transferase",

                                                                          ↪  ifelse(grepl(ARG_PRKM_with_name
                                                                          ↪  pattern =
                                                                          ↪  "APH|ANT|AAC"),
                                                                          ↪  "3aminoglycoside",
                                                                          ↪  "7other"))))))

arg_mf <- cbind(as.character(ARG_PRKM_with_names$AMRGeneFamily),
↪  ARG_PRKM_with_names$SUPERGROUP)
arg_mf <- arg_mf[!duplicated(arg_mf[,1]), ]
arg_mf <- data.frame(arg_mf)
row.names(arg_mf) <- arg_mf$X1

names(asthma.myresults.cooccur.df.filt.no.incest) <- paste0("asthma_",
↪  names(asthma.myresults.cooccur.df.filt.no.incest))
names(healthy.myresults.cooccur.df.filt.no.incest) <- paste0("healthy_",
↪  names(healthy.myresults.cooccur.df.filt.no.incest))

catmat <- cbind(asthma.myresults.cooccur.df.filt.no.incest,
↪  healthy.myresults.cooccur.df.filt.no.incest)

# Virulence factor supergroups:
VF_RPKM_with_names$FamilyBins <- as.character(VF_RPKM_with_names$FamilyBins)
VF_RPKM_with_names$VFSUPERGROUP <- ifelse(grepl(VF_RPKM_with_names$FamilyBins, pattern =
↪  "[p|P]ili|[f|F]imbriae|[c|C]urli|Agf|[P|p]ilus|ECP|[e|E]xopolysaccharide|CVF587|CVF744|Fibronectin-
↪  protein|Choline-binding proteins|[c|C]apsule|Cell surface|RcsAB|Streptococcal plasmin
↪  receptor|[f|F]lagella"), "Adherence, Motility, & Capsule",

                                                      ↪  ifelse(grepl(VF_RPKM_with_names$FamilyBins,
                                                      ↪  pattern = "Chu|[H|h]eme|Ent
                                                      ↪  |VF0562|[E|e]nterobactin|Iron|[p|P]yoverdine|Shu|Yb
                                                      ↪  "Iron Uptake",

                                                            ↪  ifelse(grepl(VF_RPKM_with_names$FamilyBins,
                                                            ↪  pattern =
                                                            ↪  "Colicin|Enterotoxin|Tsh
                                                            ↪  |VF0233|LOS"), "Toxins",

                                                                    ↪  ifelse(grepl(VF_RPKM_with_names$Family
                                                                    ↪  pattern =
                                                                    ↪  "T7SS|T6SS|gsp "),
                                                                    ↪  "SS",

                                                                          ↪  ifelse(grepl(VF_RPKM_with_names
                                                                          ↪  pattern
                                                                          ↪  ="GroEL|Nucleoside
                                                                          ↪  diphosphate
                                                                          ↪  kinase|Potassium/proton
                                                                          ↪  antiporter|Trigger
                                                                          ↪  factor"),
                                                                          ↪  "Stress, Etc.",
                                                                          ↪  "7other")))))
```

```
vf_mf <- cbind(as.character(VF_RPKM_with_names$FamilyBins),
↪  VF_RPKM_with_names$VFSUPERGROUP)
vf_mf <- vf_mf[!duplicated(vf_mf[,1]), ]
vf_mf <- data.frame(na.omit(vf_mf))
row.names(vf_mf) <- vf_mf$X1

# make key for heatmap colors:
# asthma   | healthy:
# ------------------
# random   | random
# ran/neg  | positive
# ran/pos  | negative
# positive  | ran/neg
# negative  | ran/pos

catmat$heatmap_key <- ifelse(catmat$healthy_sig == "Random" & catmat$asthma_sig ==
↪  "Random", "Random",
                             ifelse(catmat$healthy_sig == "Positive" & catmat$asthma_sig
                             ↪  != "Positive", "HealthyPos",
                                    ifelse(catmat$healthy_sig == "Negative" &
                                    ↪  catmat$asthma_sig != "Negative", "HealthyNeg",
                                           ifelse(catmat$asthma_sig == "Negative" &
                                           ↪  catmat$healthy_sig != "Negative",
                                           ↪  "AsthmaNeg",
                                                  ifelse(catmat$asthma_sig == "Positive"
                                                  ↪  & catmat$healthy_sig != "Positive",
                                                  ↪  "AsthmaPos",
                                                         ifelse(catmat$asthma_sig ==
                                                         ↪  "Positive" &
                                                         ↪  catmat$healthy_sig
                                                         ↪  =="Positive", "BothPos",
                                                                ifelse(catmat$asthma_sig
                                                                ↪  == "Negative" &
                                                                ↪  catmat$healthy_sig
                                                                ↪  =="Negative",
                                                                ↪  "BothNeg",
                                                                ↪  "ERROR")))))))

# View(catmat[,c("heatmap_key", "asthma_sig", "healthy_sig")])

# heatmap needs numbers:
catmat$heatmap_key_numeric <- as.numeric(factor(catmat$heatmap_key)) #Levels: AsthmaNeg
↪  AsthmaPos BothPos HealthyNeg HealthyPos Random

# catmat
sum(catmat$asthma_comp_name == catmat$healthy_comp_name) == dim(catmat)[1]
```

```
## [1] TRUE
```

```
# -----------------------------------------------------------------------
# DROP rows and cols that are all p<0.05 *******
if(all(catmat[, "asthma_sp1_name"] == catmat[, "healthy_sp1_name"])){
  sp1_num_sig <- catmat %>% group_by(asthma_sp1_name) %>%
```

```
    summarize(rank=sum(asthma_p_gt<0.05 | asthma_p_lt<0.05 | healthy_p_gt<0.05 |
    ↪  healthy_p_lt<0.05))
  sp2_num_sig <- catmat %>% group_by(asthma_sp2_name) %>%
    summarize(rank=sum(asthma_p_gt<0.05 | asthma_p_lt<0.05 | healthy_p_gt<0.05 |
    ↪  healthy_p_lt<0.05))
} else {print("ERROR")}
sp1_ranks <- sp1_num_sig[sp1_num_sig$rank>0,]
sp2_ranks <- sp2_num_sig[sp2_num_sig$rank>0,]
# ---------------------------------------------------------------------------
# make better supergroups now that we've dropped some
# lilmf <- ARG_PRKM_with_names[, 96:103]
# lilmf[lilmf$AMRGeneFamily %in% sp2_ranks$asthma_sp2_name, ]
# try DrugClassBins instead of supergroups
# arg_mf <- cbind(as.character(ARG_PRKM_with_names$AMRGeneFamily),
↪  ARG_PRKM_with_names$DrugClassBins)
# arg_mf <- arg_mf[!duplicated(arg_mf[,1]), ]
# arg_mf <- data.frame(arg_mf)
# row.names(arg_mf) <- arg_mf$X1

# # try supergrouping DrugClassBins
ARG_PRKM_with_names$DrugSUPERGROUP <- ifelse(grepl(ARG_PRKM_with_names$DrugClassBins,
↪  pattern = "phenicol|glycopeptide|lincosamide|nucleoside|mupirocin"), "Other",
↪  ARG_PRKM_with_names$DrugClassBins)
ARG_PRKM_with_names$DrugSUPERGROUP <- gsub(x = ARG_PRKM_with_names$DrugSUPERGROUP,
↪  pattern = " antibiotic", replacement = "")
arg_mf <- cbind(as.character(ARG_PRKM_with_names$AMRGeneFamily),
↪  ARG_PRKM_with_names$DrugSUPERGROUP)
arg_mf <- arg_mf[!duplicated(arg_mf[,1]), ]
arg_mf <- data.frame(arg_mf)
row.names(arg_mf) <- arg_mf$X1

# ---------------------------------------------------------------------------

# re-order for heatmap by number of significant hits
tmp=expand.grid(as.character(sp1_ranks$asthma_sp1_name[order(sp1_ranks$rank, decreasing =
↪  TRUE)]), as.character(sp2_ranks$asthma_sp2_name[order(sp2_ranks$rank, decreasing =
↪  TRUE)]))
comp_name_order <- paste(tmp$Var1, tmp$Var2, sep = "|v|")
comp_name_order <- comp_name_order[(as.character(comp_name_order) %in%
↪  row.names(catmat))]
catmat.ordered <- catmat[as.character(comp_name_order), ]

# re-order by supergroups
tmp2=expand.grid(as.character(vf_mf[order(vf_mf$X2), "X1"]),
↪  as.character(arg_mf[order(arg_mf$X2), "X1"]))
comp_name_order2 <- paste(tmp$Var1, tmp$Var2, sep = "|v|")
comp_name_order2 <- comp_name_order2[(as.character(comp_name_order2) %in%
↪  row.names(catmat))]
catmat.ordered <- catmat.ordered[as.character(comp_name_order), ]

# catmat.ordered
↪  <-catmat.ordered[order(arg_mf[as.character(catmat.ordered$asthma_sp2_name),"X2"]),] #
↪  groups ARGs by arbitrary super-family
```

```
catmat.ordered$SUPERGROUP <- arg_mf[as.character(catmat.ordered$asthma_sp2_name),"X2"]
catmat.ordered$VFSUPERGROUP <- vf_mf[as.character(catmat.ordered$asthma_sp1_name),"X2"]
catmat.ordered$asthma_sp1_name <- factor(catmat.ordered$asthma_sp1_name, levels =
↪  unique(catmat.ordered$asthma_sp1_name))
catmat.ordered$asthma_sp2_name <- factor(catmat.ordered$asthma_sp2_name, levels =
↪  rev(unique(catmat.ordered$asthma_sp2_name)))
catmat.ordered$SUPERGROUP <- factor(catmat.ordered$SUPERGROUP,
↪  levels=c(as.character(unique(arg_mf[as.character(catmat.ordered$asthma_sp2_name),"X2"])[unique(arg_
↪  != "Other"]), "Other")) # labels = c("Efflux pumps", "transferases", "aminoglycoside
↪  resistant", "glycopeptide resistance", "beta-lactamases", "other"))

# assign text colour
textcol <- "black" # "grey40"
p <- ggplot(catmat.ordered, aes(x=asthma_sp1_name, y=asthma_sp2_name, fill=heatmap_key,
                        #color=as.factor(asthma_sig)
                        ))+
    #add border white colour of line thickness 0.25
  facet_grid(col=vars(VFSUPERGROUP), rows=vars(SUPERGROUP), scales='free', space="free")
  ↪  +
  geom_tile(size=0.25, width=0.9, height=0.9)+ #colour="white",
  #remove x and y axis labels
  labs(x="", y="")+
  scale_fill_manual(values=c(brewer.pal(11, "Spectral")[c(10, 1, 5, 9, 3)], "grey"),
  ↪  na.value = "grey40")+
  # scale_fill_manual(values=c("#d53e4f", "#f46d43", "#fdae61", "#fee08b", "#e6f598",
  ↪  "#abdda4", "#ddf1da"), na.value = "grey40")+
  # scale_fill_manual(values=brewer.pal(7, "BrBG")[c(7,6,2,5,4,1)], na.value="grey90")+
  ↪
  scale_color_manual(values=c("blue", "red", "white"))+
  #remove extra space
  # scale_y_discrete(expand=c(0, 0))+
  guides(fill=guide_legend(title="Co-occurrence"))+
  #set a base size for all fonts
  theme_minimal(base_size=12)+
  # coord_flip()+
  xlab("Virulence Factors")+
  ylab("Antibiotic Resistance Genes") +
  theme(legend.position="right", legend.direction="vertical",
        strip.text.y = element_text(angle = 0, vjust=0.5, hjust=0, face="bold"),
        strip.text.x = element_text(face="bold"),#angle = 45, size = 10, hjust=1),
        legend.title=element_text(colour=textcol),
        legend.margin=margin(grid::unit(0, "cm")),
        legend.text=element_text(colour=textcol, size=12, face="bold"),
        legend.key.height=grid::unit(0.8, "cm"),
        legend.key.width=grid::unit(0.4, "cm"),
        axis.text.x=element_text(size=10, colour=textcol, angle=45,hjust=1),
        axis.text.y=element_text(vjust=0.2, colour=textcol),
        axis.ticks=element_line(size=0.4),
        plot.background=element_blank(),
        panel.grid.major = element_blank(),
        panel.grid.minor = element_blank(),
        panel.border = element_rect(colour = "black", fill=NA, size=1),
```

```
        plot.margin=margin(0.7, 0.4, 0.1, 0.2, "cm"),
        plot.title=element_text(colour=textcol, hjust=0, size=14, face="bold")
      )
# ggsave(p, filename="./HvA_VFvARG_cooccurrence_filtered.jpg", device = "jpeg",
       # width=40, height=21, units = "cm")
# ggsave(p, filename="./HvA_VFvARG_cooccurrence_filtered.pdf", device = "pdf",
       # width=40, height=21, units = "cm", useDingbats=FALSE)
p
```

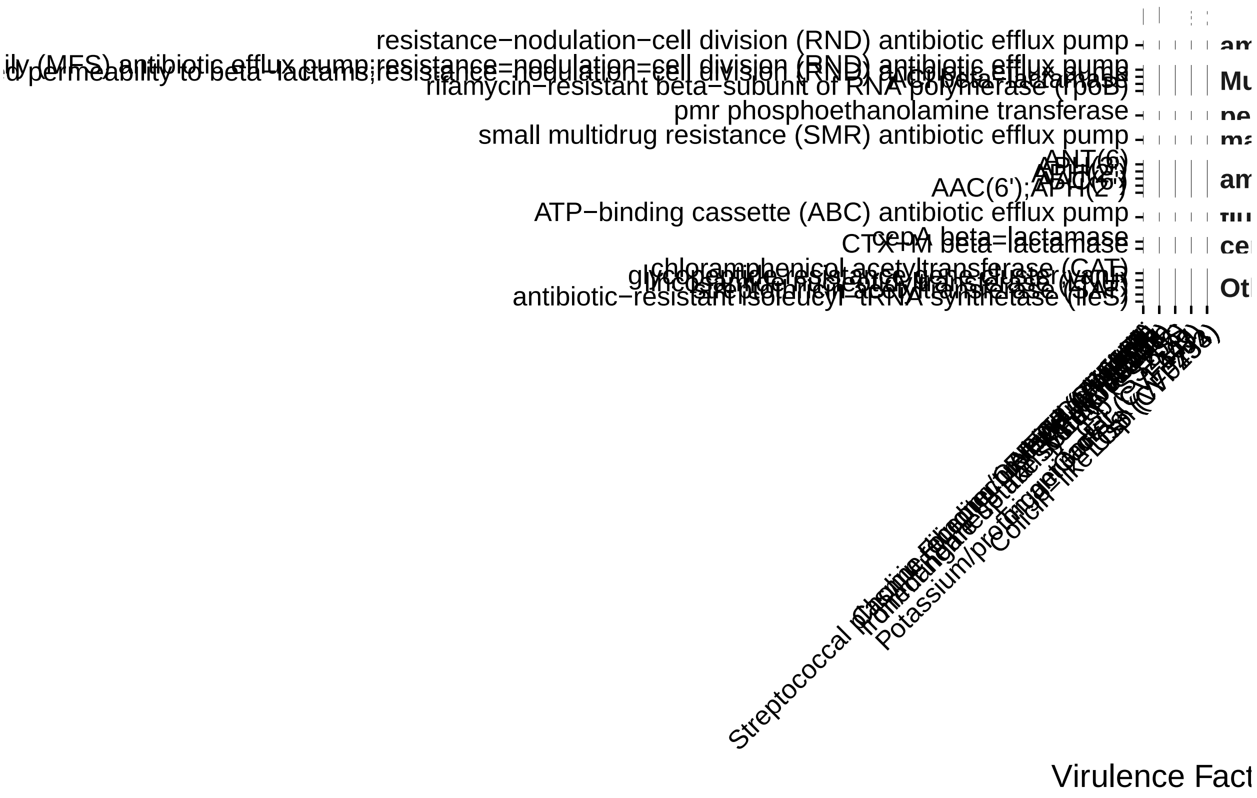


#### Figure S5:

CO-OCCURRENCE- filtering out non-taxonomic self comparisons "no incest" (constrain to looking at within species relationships) Run coooccur on data constrained to species (run previous 3 chunks first)

```
rm(list=ls())
library("cooccur")
# pseudocode
# run through the list of taxa in VF MARS data, for each taxon:
# grab all ARGs that have been found in that taxa
# cross reference that list with which ARGs we detected in MARS
# Run cooccur on those args/vf incidences in MARS
# Repeat for every taxon sourcing the VFs we detected
df.CARD.F.bin <- readRDS("./df.CARD.F.bin.rds")
df.shortbred.VFgene.Incidence <- readRDS("~/Library/CloudStorage/Box-Box/Kau
↪  Lab/Results/MARS/FecalMetagenomics/shortbred/VFandARG21/VFvARG_taxa_constrained/datatables/MARS_VFDI
MARS_CARD_taxtab <- readRDS(file = "~/Library/CloudStorage/Box-Box/Kau
↪  Lab/Results/MARS/FecalMetagenomics/shortbred/databases/CARD/ngw_make_taxa_arg_table/MARS_CARD_taxtal
```

```r
MARS_CARD_taxtab <- MARS_CARD_taxtab[MARS_CARD_taxtab$GeneName != "ugd",] # triple
↪  checking that this is gone

VFindex <- readRDS(file = "~/Library/CloudStorage/Box-Box/Kau
↪  Lab/Results/MARS/FecalMetagenomics/shortbred/databases/VFDB/ngw_VFindex.rds")

basedir="./"
humann3_mf_filtered_perm <- readRDS(file = paste0(basedir, "humann3_mf_filtered.rds"))
row.names(humann3_mf_filtered_perm) <- humann3_mf_filtered_perm$SampleName
df.meta <- humann3_mf_filtered_perm
for (VFspecies in unique(VFindex$species)) {
  if (any(MARS_CARD_taxtab$Source %in% VFspecies)) {
    print(VFspecies)
    df.CARD.F.tmp <-
↪   df.CARD.F.bin[as.character(unique(MARS_CARD_taxtab[MARS_CARD_taxtab$Source %in%
↪   VFspecies, "GeneName"])), 1:95]
    df.VFDB.F.tmp <-
↪   df.shortbred.VFgene.Incidence[as.character(unique(VFindex[VFindex$species %in%
↪   VFspecies, "VFID_strain"])), 1:95]
    df.VFDB.F.tmp <- df.VFDB.F.tmp[, as.character(names(df.CARD.F.tmp))]
    print(paste("ARGs:", dim(df.CARD.F.tmp)[1]))
    print(paste("VFs:", dim(df.VFDB.F.tmp)[1]))
    print("Running cooccur.")
    if (dim(df.CARD.F.tmp)[1] > 1 & dim(df.VFDB.F.tmp)[1] > 1) {
      if (all(names(df.VFDB.F.tmp)[1:95] == names(df.CARD.F.tmp)[1:95])) {
        # ALL SAMPLES
        mymat <- apply(rbind(df.VFDB.F.tmp[,1:95], df.CARD.F.tmp[,1:95]), MARGIN = 2, FUN
↪   = as.numeric)
        row.names(mymat) <- c(row.names(df.VFDB.F.tmp[,1:95]),
         ↪  row.names(df.CARD.F.tmp[,1:95]))
        mymat.bin <- mymat
        mymat.bin[mymat.bin>0]=1 # make binary
        myresults.cooccur <- cooccur(mymat.bin, spp_names = TRUE)
        saveRDS(myresults.cooccur, paste0("./", VFspecies,
         ↪  "_VFvARGgenelevel_AllSamples.cooccur"))
        saveRDS(mymat.bin, paste0("./", VFspecies,
         ↪  "_VFvARGgenelevel_AllSamples_cooccur_input.rds"))
        # ASTHMA ONLY
        mymat2 <- apply(rbind(
          df.VFDB.F.tmp[,names(df.VFDB.F.tmp)[names(df.VFDB.F.tmp) %in%
↪   row.names(df.meta[df.meta$ASTHMA=="Asthmatic",])]],
          df.CARD.F.tmp[,names(df.CARD.F.tmp)[names(df.CARD.F.tmp) %in%
↪   row.names(df.meta[df.meta$ASTHMA=="Asthmatic",])]]),
                      MARGIN = 2, FUN = as.numeric)
        row.names(mymat2) <- c(row.names(df.VFDB.F.tmp[,1:95]),
         ↪  row.names(df.CARD.F.tmp[,1:95]))
        mymat2.bin <- mymat2
        mymat2.bin[mymat2.bin>0]=1 # make binary
        myresults.cooccur <- cooccur(mymat2.bin, spp_names = TRUE)
        saveRDS(myresults.cooccur, paste0("./", VFspecies,
         ↪  "_VFvARGgenelevel_Asthma.cooccur"))
        saveRDS(mymat2.bin, paste0("./", VFspecies,
         ↪  "_VFvARGgenelevel_Asthma_cooccur_input.rds"))
```

```
        # HEALTHY ONLY
        mymat3 <- apply(rbind(
          df.VFDB.F.tmp[,names(df.VFDB.F.tmp)[names(df.VFDB.F.tmp) %in%
↪   row.names(df.meta[df.meta$ASTHMA=="Healthy",])]],
          df.CARD.F.tmp[,names(df.CARD.F.tmp)[names(df.CARD.F.tmp) %in%
↪   row.names(df.meta[df.meta$ASTHMA=="Healthy",])]]),
                      MARGIN = 2, FUN = as.numeric)
        row.names(mymat3) <- c(row.names(df.VFDB.F.tmp[,1:95]),
        ↪   row.names(df.CARD.F.tmp[,1:95]))
        mymat3.bin <- mymat3
        mymat3.bin[mymat3.bin>0]=1 # make binary
        myresults.cooccur <- cooccur(mymat3.bin, spp_names = TRUE)
        saveRDS(myresults.cooccur, paste0("./", VFspecies,
        ↪   "_VFvARGgenelevel_Healthy.cooccur"))
              saveRDS(mymat3.bin, paste0("./", VFspecies,
              ↪   "_VFvARGgenelevel_Healthy_cooccur_input.rds"))
      } else { print("ERROR: Samples names do not match.")}
    } else { print("There is only a 1:1 comparison. Skipping these comparisons. Do
↪   Fishers test.")}
  } else {
    print(paste("No", VFspecies, "in CARD dataset"))
  }
}
```

```
## [1] "Salmonella enterica"
## [1] "ARGs: 29"
## [1] "VFs: 6"
## [1] "Running cooccur."
##   |                                                                              |
```

```
## [1] "No Streptococcus mutans in CARD dataset"
## [1] "No Clostridium beijerinckii in CARD dataset"
## [1] "Shigella dysenteriae"
## [1] "ARGs: 16"
## [1] "VFs: 1"
## [1] "Running cooccur."
## [1] "There is only a 1:1 comparison. Skipping these comparisons. Do Fishers test."
## [1] "Haemophilus influenzae"
## [1] "ARGs: 21"
## [1] "VFs: 6"
## [1] "Running cooccur."
##   |                                                                              |
```

```
ecoli <- readRDS("./Escherichia coli_VFvARGgenelevel_Asthma.cooccur")
summary(ecoli)
```

```
## Call:
## cooccur(mat = mymat2.bin, spp_names = TRUE)
##
## Of 5886 species pair combinations, 3776 pairs (64.15 %) were removed from the analysis because expec
##
## Cooccurrence Summary:

##        Species          Sites       Positive       Negative         Random
##          109.0           36.0          534.0           16.0         1560.0
## Unclassifiable Non-random (%)
##            0.0           26.1
## attr(,"class")
## [1] "summary.cooccur"
```

```
# plot(ecoli)
# print(ecoli)

klebsiella <- readRDS("./Klebsiella pneumoniae_VFvARGgenelevel_Asthma.cooccur")
summary(klebsiella)
```

```
## Call:
## cooccur(mat = mymat2.bin, spp_names = TRUE)
##
## Of 820 species pair combinations, 426 pairs (51.95 %) were removed from the analysis because expecte
##
## Cooccurrence Summary:

##        Species          Sites       Positive       Negative         Random
##           41.0           36.0           44.0            1.0          349.0
## Unclassifiable Non-random (%)
##            0.0           11.4
## attr(,"class")
## [1] "summary.cooccur"
```

```
# plot(klebsiella)
# print(klebsiella)

# yersinia <- readRDS("./Yersinia enterocolitica_VFvARGgenelevel_Asthma.cooccur")
# summary(yersinia)
# plot(yersinia)
```

Asthma v Healthy co-occurrence on species-constrained VF/ARG comparisons - prepare data for heatmap

```
rm(list=ls())
source("./humann3_210613_functions_ngw.R")
library(tidyverse)
datadir <- "./"
cooccurdir <- "./"
VFindex <- readRDS(file = "~/Library/CloudStorage/Box-Box/Kau
↪  Lab/Results/MARS/FecalMetagenomics/shortbred/databases/VFDB/ngw_VFindex.rds")
MARS_CARD_taxtab <- readRDS(file = "~/Library/CloudStorage/Box-Box/Kau
↪  Lab/Results/MARS/FecalMetagenomics/shortbred/databases/CARD/ngw_make_taxa_arg_table/MARS_CARD_taxta
```

```r
makeCategoricalSigColumn <- function(tmp.df) {
  tmp.df$sig <- ifelse(tmp.df$p_gt < 0.05, "Positive",
                       ifelse(tmp.df$p_lt < 0.05, "Negative", "Random"))
  tmp.df$sig <- factor(tmp.df$sig, levels = c("Negative", "Positive", "Random"))
  return(tmp.df)
}

# READ IN COOCCUR RESULTS
possibleSpecies <- readRDS("~/Library/CloudStorage/Box-Box/Kau
↪  Lab/Results/MARS/FecalMetagenomics/shortbred/VFandARG21/VFvARG_taxa_constrained/datatables/all_teste
# VFspecies <- "Escherichia coli"
VFspecies = "Yersinia enterocolitica"
species_tested_list <- c()

for (VFspecies in unique(possibleSpecies)) {
  print(paste("Running:", VFspecies))
  healthy.myresults.cooccur <- readRDS(file = paste0(cooccurdir, VFspecies,
↪  "_VFvARGgenelevel_Healthy.cooccur"))
  asthma.myresults.cooccur <- readRDS(file = paste0(cooccurdir, VFspecies,
↪  "_VFvARGgenelevel_Asthma.cooccur"))
  all(healthy.myresults.cooccur$spp_key == asthma.myresults.cooccur$spp_key) # names
   ↪  match numbered labels

  # get significance bins and effect sizes
  healthy.myresults.cooccur.df <-
↪  makeCategoricalSigColumn(data.frame(healthy.myresults.cooccur$results))
  # healthy.myresults.cooccur.df$effect_sizes <- effect.sizes(healthy.myresults.cooccur)
  healthy.myresults.cooccur.df=merge(x=healthy.myresults.cooccur.df, y =
↪  effect.sizes(healthy.myresults.cooccur), by.x = c("sp1_name", "sp2_name"), by.y =
↪  c("sp1", "sp2"))
  asthma.myresults.cooccur.df <-
↪  makeCategoricalSigColumn(data.frame(asthma.myresults.cooccur$results))
  asthma.myresults.cooccur.df=merge(x=asthma.myresults.cooccur.df, y =
↪  effect.sizes(asthma.myresults.cooccur), by.x = c("sp1_name", "sp2_name"), by.y =
↪  c("sp1", "sp2"))

  # create VF/ARG pair columns:
  asthma.myresults.cooccur.df$comp_name <- paste(asthma.myresults.cooccur.df$sp1_name,
↪  asthma.myresults.cooccur.df$sp2_name, sep = "|v|")
  asthma.myresults.cooccur.df$comp_name_rev <-
↪  paste(asthma.myresults.cooccur.df$sp2_name, asthma.myresults.cooccur.df$sp1_name, sep
↪  = "|v|") # this is to check if the 2 matrices have pairs that were named differently
  healthy.myresults.cooccur.df$comp_name <- paste(healthy.myresults.cooccur.df$sp1_name,
↪  healthy.myresults.cooccur.df$sp2_name, sep = "|v|")
  healthy.myresults.cooccur.df$comp_name_rev <-
↪  paste(healthy.myresults.cooccur.df$sp2_name, healthy.myresults.cooccur.df$sp1_name,
↪  sep = "|v|") # this is to check if the 2 matrices have pairs that were named
↪  differently

  # asthma but not in healthy:
  
   ↪  length(unique(asthma.myresults.cooccur.df$comp_name)[unique(asthma.myresults.cooccur.df$comp_name)
   ↪  %notin% unique(healthy.myresults.cooccur.df$comp_name)]) # 1286
```

```
↪ length(unique(asthma.myresults.cooccur.df$comp_name)[unique(asthma.myresults.cooccur.df$comp_name)
↪ %in% unique(healthy.myresults.cooccur.df$comp_name_rev)]) # 0 (means order of vf v
↪ arg was ok)
# in healthy but not in asthma:

↪ length(unique(healthy.myresults.cooccur.df$comp_name)[unique(healthy.myresults.cooccur.df$comp_nar
↪ %notin% unique(asthma.myresults.cooccur.df$comp_name)]) # 770

↪ length(unique(healthy.myresults.cooccur.df$comp_name)[unique(healthy.myresults.cooccur.df$comp_nar
↪ %in% unique(asthma.myresults.cooccur.df$comp_name_rev)]) # 0 (means order of vf v
↪ arg was ok)

# so some pairs were filtered out for each matrix, presumably because of the "thresh"
↪ option in cooccur which drops pairs whose expected co-occurrence is less than 1.
uniqueAsthmaPairs <-
↪ unique(asthma.myresults.cooccur.df$comp_name)[unique(asthma.myresults.cooccur.df$comp_name)
↪ %notin% unique(healthy.myresults.cooccur.df$comp_name)] # 98
uniqueHealthyPairs <-
↪ unique(healthy.myresults.cooccur.df$comp_name)[unique(healthy.myresults.cooccur.df$comp_name)
↪ %notin% unique(asthma.myresults.cooccur.df$comp_name)] # 130

# drop these from both matrices:
asthma.myresults.cooccur.df.filt <-
↪ asthma.myresults.cooccur.df[asthma.myresults.cooccur.df$comp_name %notin%
↪ uniqueAsthmaPairs,]
healthy.myresults.cooccur.df.filt <-
↪ healthy.myresults.cooccur.df[healthy.myresults.cooccur.df$comp_name %notin%
↪ uniqueHealthyPairs,]

dim(healthy.myresults.cooccur.df.filt) == dim(asthma.myresults.cooccur.df.filt)
sum(asthma.myresults.cooccur.df.filt$comp_name %in%
↪ healthy.myresults.cooccur.df.filt$comp_name) ==
↪ dim(healthy.myresults.cooccur.df.filt)[1]

# NOW COMPARE HvA: ####
row.names(healthy.myresults.cooccur.df.filt) <-
↪ healthy.myresults.cooccur.df.filt$comp_name
row.names(asthma.myresults.cooccur.df.filt) <-
↪ asthma.myresults.cooccur.df.filt$comp_name

if(sum(asthma.myresults.cooccur.df.filt$comp_name ==
↪ healthy.myresults.cooccur.df.filt$comp_name) ==
↪ dim(healthy.myresults.cooccur.df.filt)[1]){
  results.df <- data.frame(cbind(healthy.myresults.cooccur.df.filt$sig,
↪ asthma.myresults.cooccur.df.filt$sig))
  names(results.df) <- c("Healthy", "Asthma")
  row.names(results.df) <- healthy.myresults.cooccur.df.filt$comp_name
  results.df$match <- ifelse(results.df$Healthy == results.df$Asthma, 1, 0)
}

allMismatches <- row.names(results.df)[results.df$match ==0] # 421 mismatches
AsthmaPos <- row.names(results.df)[results.df$match ==0 & results.df$Asthma ==
↪ "Positive"] #54 (1 v Negative)
```

```
HealthyPos <- row.names(results.df)[results.df$match ==0 & results.df$Healthy ==
↪  "Positive"] #320 (all v Random)
AsthmaNeg <- row.names(results.df)[results.df$match ==0 & results.df$Asthma ==
↪  "Negative"] #29 (all v Random)
HealthyNeg <- row.names(results.df)[results.df$match ==0 & results.df$Healthy ==
↪  "Negative"] #18 (1 v Positive)

row.names(results.df)[results.df$match ==0 & results.df$Asthma == "Negative" &
 ↪  results.df$Healthy == "Positive"] #0
row.names(results.df)[results.df$match ==0 & results.df$Asthma == "Positive" &
 ↪  results.df$Healthy == "Negative"] #0
row.names(results.df)[results.df$match ==0]
asthma.myresults.cooccur.df[asthma.myresults.cooccur.df$comp_name %in% AsthmaPos,]

#filter out VFvVF and ARGvARG # THIS IS BASED ON A GREP PATTERN THAT COULD CHANGE SO
 ↪  CHECK
healthy.myresults.cooccur$spp_key[grepl(pattern = "__",
↪  healthy.myresults.cooccur$spp_key[,"spp"]), "spp"] # ALL VFs
healthy.myresults.cooccur$spp_key[!(grepl(pattern = "__",
↪  healthy.myresults.cooccur$spp_key[,"spp"])),"spp"] # ALL ARGs

VFvARG.df <- expand.grid(healthy.myresults.cooccur$spp_key[grepl(pattern = "__",
↪  healthy.myresults.cooccur$spp_key[,"spp"]), "spp"],
↪  healthy.myresults.cooccur$spp_key[!(grepl(pattern = "__",
↪  healthy.myresults.cooccur$spp_key[,"spp"])),"spp"])
VFvARG_all_possible <- paste(VFvARG.df$Var1, VFvARG.df$Var2, sep = "|v|")
if (sum(asthma.myresults.cooccur.df.filt$comp_name %in% VFvARG_all_possible) > 0) {
asthma.myresults.cooccur.df.filt.no.incest <-
↪  asthma.myresults.cooccur.df.filt[asthma.myresults.cooccur.df.filt$comp_name %in%
↪  VFvARG_all_possible,]
saveRDS(asthma.myresults.cooccur.df.filt.no.incest, paste0(datadir, VFspecies,
 ↪  ".asthma.myresults.cooccur.df.filt.no.incest.rds"))
species_tested_list <- c(species_tested_list, VFspecies) # record list of tested
↪  species
} else { print("NO VF/ARG combinations in the asthma group survived filtering.")}
if (sum(healthy.myresults.cooccur.df.filt$comp_name %in% VFvARG_all_possible) > 0) {
healthy.myresults.cooccur.df.filt.no.incest <-
↪  healthy.myresults.cooccur.df.filt[healthy.myresults.cooccur.df.filt$comp_name %in%
↪  VFvARG_all_possible,]
  saveRDS(healthy.myresults.cooccur.df.filt.no.incest, paste0(datadir, VFspecies,
   ↪  ".healthy.myresults.cooccur.df.filt.no.incest.rds"))
} else { print("NO VF/ARG combinations in the healthy group survived filtering.")}

# Plot these and highlight which way the mismatch goes (sig positive/neg in asthma or
 ↪  healthy):
#
 ↪  asthma.myresults.cooccur.df.filt.no.incest[asthma.myresults.cooccur.df.filt.no.incest$comp_name
 ↪  %in% allMismatches,]
#
 ↪  healthy.myresults.cooccur.df.filt.no.incest[healthy.myresults.cooccur.df.filt.no.incest$comp_name
 ↪  %in% allMismatches,]

saveRDS(asthma.myresults.cooccur.df, paste0(datadir, VFspecies,
 ↪  ".asthma.myresults.cooccur.df.unfiltered.rds"))
```

```
  saveRDS(healthy.myresults.cooccur.df, paste0(datadir, VFspecies,
  ↪  ".healthy.myresults.cooccur.df.unfiltered.rds"))
}
```

```
## [1] "Running: Salmonella enterica"
## [1] "Running: Escherichia coli"
## [1] "Running: Haemophilus influenzae"
## [1] "Running: Klebsiella pneumoniae"
```

```
saveRDS(species_tested_list, paste0("./", "all_tested_species_list.rds"))
```

#### Figure S5A: Richness constrained to taxa

```
rm(list=ls())
source("./humann3_210613_functions_ngw.R")
library(tidyverse)
library(ggplot2)
library(RColorBrewer)
# library(ggprism)
# library(rstatix)

basedir="./"
humann3_mf_filtered_perm <- readRDS(file = paste0(basedir, "humann3_mf_filtered.rds"))
row.names(humann3_mf_filtered_perm) <- humann3_mf_filtered_perm$SampleName
df.meta <- humann3_mf_filtered_perm

df.CARD.F.bin <- readRDS("./df.CARD.F.bin.rds")
df.shortbred.VFgene.Incidence <- readRDS("~/Library/CloudStorage/Box-Box/Kau
↪  Lab/Results/MARS/FecalMetagenomics/shortbred/VFandARG21/VFvARG_taxa_constrained/datatables/MARS_VFDB
MARS_CARD_taxtab <- readRDS(file = "~/Library/CloudStorage/Box-Box/Kau
↪  Lab/Results/MARS/FecalMetagenomics/shortbred/databases/CARD/ngw_make_taxa_arg_table/MARS_CARD_taxtab
VFindex <- readRDS(file = "~/Library/CloudStorage/Box-Box/Kau
↪  Lab/Results/MARS/FecalMetagenomics/shortbred/databases/VFDB/ngw_VFindex.rds")
possibleSpecies <- readRDS("~/Library/CloudStorage/Box-Box/Kau
↪  Lab/Results/MARS/FecalMetagenomics/shortbred/VFandARG21/VFvARG_taxa_constrained/datatables/all_teste

# from list of species you want to look at, loop thru em
# search species thru ARG and VF taxa table
# make list of  ARGs and VFs detected in MARS per species
# count average richness of these genes for asthma and healthy cohorts
# barplot: healthy VFs, healthy ARGs, asthma VFs, asthma ARGs

VFspecies <- possibleSpecies[2]
df.richness.results <- data.frame(matrix(nrow = 0, ncol = 3))
colnames(df.richness.results) <- c("aov.p", "tukey.pvals", "VFspecies")

df.richness.wilcoxons <- data.frame(matrix(nrow = 0, ncol = 3))
colnames(df.richness.wilcoxons) <- c("species", "ARG_pval", "VF_pval")
# VFspecies = ""
for( VFspecies in possibleSpecies){

  print(paste("Running:", VFspecies))
```

```
VFlist <- VFindex[VFindex$species %in% VFspecies, "VFID_strain"]
ARGlist <- MARS_CARD_taxtab[MARS_CARD_taxtab$Source %in% VFspecies, "GeneName"]

VF_all <- df.shortbred.VFgene.Incidence[VFlist, ]
ARG_all <- df.CARD.F.bin[ARGlist, ]

VFs.asthma <- VF_all[,names(VF_all)[names(VF_all) %in%
↪  row.names(df.meta[df.meta$ASTHMA=="Asthmatic",])]]
VFs.healthy <- VF_all[,names(VF_all)[names(VF_all) %in%
↪  row.names(df.meta[df.meta$ASTHMA=="Healthy",])]]
ARGs.asthma <- ARG_all[,names(ARG_all)[names(ARG_all) %in%
↪  row.names(df.meta[df.meta$ASTHMA=="Asthmatic",])]]
ARGs.healthy <- ARG_all[,names(ARG_all)[names(ARG_all) %in%
↪  row.names(df.meta[df.meta$ASTHMA=="Healthy",])]]

mean(colSums(VFs.asthma))
mean(colSums(VFs.healthy))
mean(colSums(ARGs.asthma))
mean(colSums(ARGs.healthy))

dat <- data.frame(c(colSums(VFs.asthma),
            colSums(VFs.healthy),
            colSums(ARGs.asthma),
            colSums(ARGs.healthy)),
           c(rep("VFAsthma", dim(VFs.asthma)[2]),
             rep("VFHealthy", dim(VFs.healthy)[2]),
             rep("ARGAsthma", dim(ARGs.asthma)[2]),
             rep("ARGHealthy", dim(ARGs.healthy)[2])),
           c(rep("VF", (dim(VFs.healthy)[2]+ dim(VFs.asthma)[2])),
             rep("ARG", (dim(ARGs.healthy)[2]+ dim(ARGs.asthma)[2]))),
           c(rep("Asthma", dim(VFs.asthma)[2]),
             rep("Healthy", dim(VFs.healthy)[2]),
             rep("Asthma", dim(ARGs.asthma)[2]),
             rep("Healthy", dim(ARGs.healthy)[2])))
names(dat) <- c("richness", "Group", "Gene", "Cohort")
dat.plot <- my_data_summary(dat, varname="richness",
                   groupnames=c("Cohort", "Gene"))
# Convert dose to a factor variable
dat.plot$se <- as.numeric(ifelse(dat.plot$Cohort == "Asthma",
↪  dat.plot$sd/sqrt(dim(VFs.asthma)[2]),
                      ifelse(dat.plot$Cohort == "Healthy",
                       ↪  dat.plot$sd/sqrt(dim(VFs.healthy)[2]), "ERROR")))
# head(dat.plot)

VFp <- wilcox.test(richness~Cohort, data = dat[dat$Gene=="VF",],
↪  alternative="greater")$p.value
ARGp <- wilcox.test(richness~Cohort, data = dat[dat$Gene=="ARG",],
↪  alternative="greater")$p.value

# kruskal.test(richness~Group,data= dat)
# library(FSA)
# dunnTest(richness~Group,data= dat, method = "bh")
# print(summary(aov(richness~Cohort+Gene,data= dat)))
```

```
summary(aov(richness~Group,data= dat))
aov.p <- summary(aov(richness~Group,data= dat))[[1]][["Pr(>F)"]][1]
tukey.pvals <- TukeyHSD(aov(richness~Group,data= dat))$Group[, "p adj"]

df.add <- data.frame(aov.p, tukey.pvals, VFspecies)
df.richness.results <- rbind(df.richness.results, df.add)

df.add.wilcox <- data.frame(VFspecies, ARGp, VFp)
df.richness.wilcoxons <- rbind(df.richness.wilcoxons, df.add.wilcox)

dat.plot$Group <- paste0(dat.plot$Gene, dat.plot$Cohort)

# Default bar plot
textcol = "black"
p<- ggplot(dat.plot, aes(x=Cohort, y=richness, fill=Gene)) +
  geom_bar(stat="identity", color="black",
           position=position_dodge()) +
  geom_errorbar(aes(ymin=richness, ymax=richness+se), width=.2,
                 position=position_dodge(.9))
# Finished bar plot
p <- p  +
   labs(title=VFspecies, x="Cohort", y = "Richness")+
   theme_classic() +
   theme(axis.text.x=element_text(size=10, colour=textcol),
            axis.text.y=element_text(vjust=0.2, colour=textcol),
            axis.ticks=element_line(size=0.4),
            plot.title=element_text(colour=textcol, hjust=0, size=12, face="bold")
          ) +
   scale_fill_manual(values=c("#5AB4AC","#F5F5F5")) #+
   # ggpubr::stat_compare_means(comparisons = list(c("VF", "Asthma"), c("VF",
   ↪  "Healthy")), data=dat.plot, method="wilcox.test", aes(label = ..p.signif..),
   ↪  label.y = max(dat.plot$richness)*1.1)

# ggsave(p, filename=paste0("./", VFspecies, "_VF_ARG_richness.jpg"), device = "jpeg",
↪  width = 3, height = 3, units = "in")

dat.plot$Group <- factor(dat.plot$Group, levels = c("ARGHealthy",  "ARGAsthma",
↪  "VFHealthy", "VFAsthma"), ordered = TRUE)
dat.plot$Cohort <- factor(dat.plot$Cohort, levels = c("Healthy", "Asthma"))
p2<- ggplot(dat.plot, aes(x=Gene, y=richness, fill=Cohort)) +
  geom_bar(stat="identity", color="black",
           position=position_dodge()) +
  geom_errorbar(aes(ymin=richness, ymax=richness+se), width=.2,
                 position=position_dodge(.9)) +
  scale_fill_manual(values = brewer.pal(11, "BrBG")[c(7,10)])+
  labs(title=VFspecies, x="Gene Type", y = "Richness")+
  theme_classic() +
  theme(axis.text.x=element_text(size=10, colour=textcol),
          axis.text.y=element_text(vjust=0.2, colour=textcol),
          axis.ticks=element_line(size=0.4),
          plot.title=element_text(colour=textcol, hjust=0, size=12, face="bold")
        ) +
#  scale_fill_manual(values=c("#5AB4AC","#F5F5F5")) +
```

```
  ggpubr::stat_compare_means(data=dat[dat$Gene=="ARG",], method="wilcox.test",
↪  method.args = list(alternative="less"), aes(label = ..p.signif..), label.y =
↪  max(dat.plot$richness)*1.1, label.x = 1, hide.ns = FALSE, vjust = 0.5) +
  ggpubr::stat_compare_means(data=dat[dat$Gene=="VF",], method="wilcox.test", method.args
↪  = list(alternative="less"), aes(label = ..p.signif..), label.y =
↪  max(dat.plot$richness)*1.1, label.x = 2, hide.ns = FALSE, vjust = 0.5)

    # ggsave(p2, filename=paste0("./", VFspecies, "_VF_ARG_richness_wilcoxons_pval.jpg"),
    ↪  device = "jpeg", width = 3, height = 3, units = "in")
    # ggsave(p2, filename=paste0("./", VFspecies, "_VF_ARG_richness_wilcoxons_pval.pdf"),
    ↪  device = "pdf", width = 3, height = 3, units = "in", useDingbats=FALSE)

  }
```

```
## [1] "Running: Salmonella enterica"
## [1] "Running: Escherichia coli"
## [1] "Running: Haemophilus influenzae"
## [1] "Running: Klebsiella pneumoniae"
```

```
df.richness.results$signif <- df.richness.results$tukey.pvals < 0.05
write.table(df.richness.results, file = "taxa_constrained_richness_ANOVAs.tsv", sep="\t")
write.table(df.richness.wilcoxons, file = "taxa_constrained_richness_wilcoxons.tsv", sep
↪  = "\t")
df.richness.wilcoxons
```

```
##                VFspecies        ARGp        VFp
## 1    Salmonella enterica 0.021706595 0.02673848
## 2       Escherichia coli 0.009956279 0.09094725
## 3 Haemophilus influenzae 0.009218985 0.85942206
## 4  Klebsiella pneumoniae 0.006629688 0.00460098
```

### Figure S5B: co-occurrence heatmap of taxa-constrained VF/ARGs

```
rm(list=ls())
source("./humann3_210613_functions_ngw.R")
library(tidyverse)
library(ggplot2)
library(RColorBrewer)
datadir <- "./"
cooccurdir <- "./"
# df.CARD.F.bin <- readRDS("~/Library/CloudStorage/Box-Box/Kau
↪  Lab/Results/MARS/FecalMetagenomics/shortbred/VFandARG21/df.CARD.F.bin.rds")
# df.shortbred.VFgene.Incidence <-
↪  readRDS("VFvARG_taxa_constrained/datatables/MARS_VFDB_taxa_constrained_incidence.rds")
# MARS_CARD_taxtab <- readRDS(file = "~/Library/CloudStorage/Box-Box/Kau
↪  Lab/Results/MARS/FecalMetagenomics/shortbred/databases/CARD/ngw_make_taxa_arg_table/MARS_CARD_taxta
df.CARD.F.bin <- readRDS("./df.CARD.F.bin.rds")
# VFindex <- readRDS(file = "./ngw_VFindex.rds")

# READ IN cleaned data
# VFspecies <- "Escherichia coli"
possibleSpecies <- readRDS("/Users/naomiwilson/Library/CloudStorage/Box-Box/Kau
↪  Lab/Results/MARS/FecalMetagenomics/shortbred/VFandARG21/VFvARG_taxa_constrained/datatables/all_teste
```

```
# VFspecies <- "Escherichia coli"

possibleSpecies <- possibleSpecies[possibleSpecies!="Salmonella enterica"] # these don't
↪  show up in metaphlan
possibleSpecies <- possibleSpecies[possibleSpecies!="Haemophilus influenzae"] # these
↪  don't show up in metaphlan
for (VFspecies in unique(possibleSpecies)) {
  print(paste("Running:", VFspecies))
  asthma.myresults.cooccur.df.filt.no.incest <- readRDS(paste0(datadir, VFspecies,
↪  ".asthma.myresults.cooccur.df.filt.no.incest.rds"))
  healthy.myresults.cooccur.df.filt.no.incest <- readRDS(paste0(datadir, VFspecies,
↪  ".healthy.myresults.cooccur.df.filt.no.incest.rds"))
  if ((dim(asthma.myresults.cooccur.df.filt.no.incest)[1] +
   ↪  dim(healthy.myresults.cooccur.df.filt.no.incest)[1]) >0 ) {
    VF_RPKM_with_names <- readRDS("~/Library/CloudStorage/Box-Box/Kau
↪  Lab/Results/MARS/FecalMetagenomics/shortbred/VFDB21results/VF_RPKM_with_VFnames_FamilyBins_filtered
    VF_RPKM_with_names$VFID <- row.names(VF_RPKM_with_names)
    row.names(VF_RPKM_with_names) <- paste0(VF_RPKM_with_names$VFID, "__",
     ↪  VF_RPKM_with_names$Source)
# # try supergrouping DrugClassBins
df.CARD.F.bin$DrugSUPERGROUP <- ifelse(grepl(df.CARD.F.bin$DrugClassBins, pattern =
↪  "phenicol|glycopeptide|lincosamide|nucleoside|mupirocin"), "Other",
↪  df.CARD.F.bin$DrugClassBins)
df.CARD.F.bin$DrugSUPERGROUP <- gsub(x = df.CARD.F.bin$DrugSUPERGROUP, pattern = "
↪  antibiotic", replacement = "")
arg_mf <- cbind(as.character(df.CARD.F.bin$geneShortName), df.CARD.F.bin$DrugSUPERGROUP)
arg_mf <- arg_mf[!duplicated(arg_mf[,1]), ]
arg_mf <- data.frame(arg_mf)
row.names(arg_mf) <- arg_mf$X1

names(asthma.myresults.cooccur.df.filt.no.incest) <- paste0("asthma_",
↪  names(asthma.myresults.cooccur.df.filt.no.incest))
    names(healthy.myresults.cooccur.df.filt.no.incest) <- paste0("healthy_",
     ↪  names(healthy.myresults.cooccur.df.filt.no.incest))

    catmat <- cbind(asthma.myresults.cooccur.df.filt.no.incest,
↪  healthy.myresults.cooccur.df.filt.no.incest)

    # Virulence factor supergroups:
    VF_RPKM_with_names$FamilyBins <- as.character(VF_RPKM_with_names$FamilyBins)
    VF_RPKM_with_names$VFSUPERGROUP <- ifelse(grepl(VF_RPKM_with_names$FamilyBins,
↪  pattern =
↪  "[p|P]ili|[f|F]imbriae|[c|C]urli|Agf|[P|p]ilus|ECP|[e|E]xopolysaccharide|CVF587|CVF744|Fibronectin-
↪  protein|Choline-binding proteins|[c|C]apsule|Cell surface|RcsAB|Streptococcal plasmin
↪  receptor|[f|F]lagella"), "Adherence, Motility, & Capsule",

                                                ↪  ifelse(grepl(VF_RPKM_with_names$FamilyBins,
                                                ↪  pattern = "Chu|[H|h]eme|Ent
                                                ↪  |VF0562|[E|e]nterobactin|Iron|[p|P]yoverdine|Sh
                                                ↪  "Iron Uptake",

                                                       ↪  ifelse(grepl(VF_RPKM_with_names$FamilyBi
                                                       ↪  pattern =
                                                       ↪  "Colicin|Enterotoxin|Tsh
                                                       ↪  |VF0233|LOS"), "Toxins",
```

```
                                                            ↪  ifelse(grepl(VF_RPKM_with_names$F
                                                            ↪  pattern =
                                                            ↪  "T7SS|T6SS|gsp "),
                                                            ↪  "SS",

                                                                   ↪  ifelse(grepl(VF_RPKM_with_
                                                                   ↪  pattern
                                                                   ↪  ="GroEL|Nucleoside
                                                                   ↪  diphosphate
                                                                   ↪  kinase|Potassium/proton
                                                                   ↪  antiporter|Trigger
                                                                   ↪  factor"),
                                                                   ↪  "Stress,
                                                                   ↪  Etc.",
                                                                   ↪  "7other")))))

vf_mf <- cbind(row.names(VF_RPKM_with_names), VF_RPKM_with_names$VFSUPERGROUP)
vf_mf <- vf_mf[!duplicated(vf_mf[,1]), ]
vf_mf <- data.frame(na.omit(vf_mf))
row.names(vf_mf) <- vf_mf$X1

# make key for heatmap colors:
# asthma   | healthy:
# random   | random
# ran/neg  | positive
# ran/pos  | negative
# positive  | ran/neg
# negative  | ran/pos

catmat$heatmap_key <- ifelse(catmat$healthy_sig == "Random" & catmat$asthma_sig ==
↪  "Random", "Random",
                             ifelse(catmat$healthy_sig == "Positive" &
                             ↪  catmat$asthma_sig != "Positive", "HealthyPos",
                                    ifelse(catmat$healthy_sig == "Negative" &
                                    ↪  catmat$asthma_sig != "Negative",
                                    ↪  "HealthyNeg",
                                           ifelse(catmat$asthma_sig == "Negative" &
                                           ↪  catmat$healthy_sig != "Negative",
                                           ↪  "AsthmaNeg",
                                                  ifelse(catmat$asthma_sig ==
                                                  ↪  "Positive" & catmat$healthy_sig
                                                  ↪  != "Positive", "AsthmaPos",
                                                         ifelse(catmat$asthma_sig ==
                                                         ↪  "Positive" &
                                                         ↪  catmat$healthy_sig
                                                         ↪  =="Positive", "BothPos",
                                                         ↪

                                                              ↪  ifelse(catmat$asthma_sig
                                                              ↪  == "Negative" &
                                                              ↪  catmat$healthy_sig
                                                              ↪  =="Negative",
                                                              ↪  "BothNeg",
                                                              ↪  "ERROR")))))))
```

```
    # View(catmat[,c("heatmap_key", "asthma_sig", "healthy_sig")])

    # heatmap needs numbers:
    catmat$heatmap_key_numeric <- as.numeric(factor(catmat$heatmap_key, levels =
↪   c("AsthmaNeg", "AsthmaPos", "BothPos", "HealthyNeg", "HealthyPos", "Random")))
↪   #Levels: AsthmaNeg AsthmaPos BothPos HealthyNeg HealthyPos Random

    # catmat
    sum(catmat$asthma_comp_name == catmat$healthy_comp_name) == dim(catmat)[1]

# DROP rows and cols that are all p<0.05 *******
    if(all(catmat[, "asthma_sp1_name"] == catmat[, "healthy_sp1_name"])){
      sp1_num_sig <- catmat %>% group_by(asthma_sp1_name) %>%
        summarize(rank=sum(asthma_p_gt<0.05 | asthma_p_lt<0.05 | healthy_p_gt<0.05 |
        ↪   healthy_p_lt<0.05))
      sp2_num_sig <- catmat %>% group_by(asthma_sp2_name) %>%
        summarize(rank=sum(asthma_p_gt<0.05 | asthma_p_lt<0.05 | healthy_p_gt<0.05 |
        ↪   healthy_p_lt<0.05))
    } else {print("ERROR")}
    sp1_ranks <- sp1_num_sig[sp1_num_sig$rank>0,]
    sp2_ranks <- sp2_num_sig[sp2_num_sig$rank>0,]

    # re-order for heatmap by number of significant hits
    tmp=expand.grid(as.character(sp1_ranks$asthma_sp1_name[order(sp1_ranks$rank,
↪   decreasing = TRUE)]), as.character(sp2_ranks$asthma_sp2_name[order(sp2_ranks$rank,
↪   decreasing = TRUE)]))
    comp_name_order <- paste(tmp$Var1, tmp$Var2, sep = "|v|")
    comp_name_order <- comp_name_order[(as.character(comp_name_order) %in%
↪   row.names(catmat))]
    catmat.ordered <- catmat[as.character(comp_name_order), ]

    # re-order by supergroups
    tmp2=expand.grid(as.character(vf_mf[order(vf_mf$X2), "X1"]),
↪   as.character(arg_mf[order(arg_mf$X2), "X1"]))
    comp_name_order2 <- paste(tmp$Var1, tmp$Var2, sep = "|v|")
    comp_name_order2 <- comp_name_order2[(as.character(comp_name_order2) %in%
↪   row.names(catmat))]
    catmat.ordered <- catmat.ordered[as.character(comp_name_order), ]

    # catmat.ordered
    ↪   <-catmat.ordered[order(arg_mf[as.character(catmat.ordered$asthma_sp2_name),"X2"]),]
    ↪   # groups ARGs by arbitrary super-family

    catmat.ordered$SUPERGROUP <-
↪   as.character(arg_mf[as.character(catmat.ordered$asthma_sp2_name),"X2"])
    catmat.ordered$VFSUPERGROUP <-
↪   as.character(vf_mf[as.character(catmat.ordered$asthma_sp1_name),"X2"])
    catmat.ordered$asthma_sp1_name <- factor(catmat.ordered$asthma_sp1_name, levels =
↪   unique(catmat.ordered$asthma_sp1_name))
    catmat.ordered$asthma_plotsp1_name <-
↪   factor(VF_RPKM_with_names[as.character(catmat.ordered$asthma_sp1_name),
↪   "ShortName"],levels =
↪   unique(VF_RPKM_with_names[as.character(catmat.ordered$asthma_sp1_name),
↪   "ShortName"])) # use short names for heatmap
```

```r
catmat.ordered$asthma_sp2_name <- factor(catmat.ordered$asthma_sp2_name, levels =
↪  rev(unique(catmat.ordered$asthma_sp2_name)))
# catmat.ordered$SUPERGROUP <- factor(catmat.ordered$SUPERGROUP, levels = c("1pump",
↪  "2transferase", "3aminoglycoside", "4glycopeptide", "5bla", "6tet", "7other"),
↪  labels = c("Efflux pumps", "transferases", "aminoglycoside resistant",
↪  "glycopeptide resistance", "beta-lactamases", "tetracycline resistance",
↪  "other"))
catmat.ordered$SUPERGROUP <- factor(catmat.ordered$SUPERGROUP,
↪  levels=c(as.character(unique(arg_mf[as.character(catmat.ordered$asthma_sp2_name),"X2"])[unique(arg_[illegible]
↪  != "Other"]), "Other")) # labels = c("Efflux pumps", "transferases", "aminoglycoside
↪  resistant", "glycopeptide resistance", "beta-lactamases", "other"))

myColors <- c(brewer.pal(11, "Spectral")[c(10, 1, 5, 9, 3)], "grey")
names(myColors) <- c("AsthmaNeg", "AsthmaPos", "BothPos", "HealthyNeg", "HealthyPos",
 ↪  "Random")
# assign text colour
textcol <- "black" # "grey40"
p <- ggplot(catmat.ordered, aes(x=asthma_plotsp1_name, y=asthma_sp2_name,
↪  fill=heatmap_key,
                        #color=as.factor(asthma_sig)
                        ))+
    #add border white colour of line thickness 0.25
  facet_grid(col=vars(VFSUPERGROUP), rows=vars(SUPERGROUP), scales='free',
   ↪  space="free") +
  geom_tile(size=0.25, width=0.9, height=0.9)+ #colour="white",
  #remove x and y axis labels
  labs(x="", y="")+
  scale_fill_manual(values=myColors[levels(factor(catmat.ordered$heatmap_key))],
   ↪  na.value = "grey40")+
  scale_color_manual(values=c("blue", "red", "white"))+
  #remove extra space
  # scale_y_discrete(expand=c(0, 0))+
  guides(fill=guide_legend(title="Co-occurrence"))+
  #set a base size for all fonts
  theme_minimal(base_size=12)+
  # coord_flip()+
  xlab("Virulence Factors")+
  ylab("Antibiotic Resistance Genes") +
  theme(legend.position="right", legend.direction="vertical",
        strip.text.y = element_text(angle = 0, vjust=0.5, hjust=0, face="bold"),
        strip.text.x = element_text(face="bold"),#angle = 45, size = 10, hjust=1),
        legend.title=element_text(colour=textcol, face="bold"),
        legend.margin=margin(grid::unit(0, "cm")),
        legend.text=element_text(colour=textcol, size=12, face="bold"),
        legend.key.height=grid::unit(0.8, "cm"),
        legend.key.width=grid::unit(0.4, "cm"),
        axis.text.x=element_text(size=10,colour=textcol,angle=45,
         ↪  hjust=1,face="italic"),
        axis.text.y=element_text(vjust=0.2, colour=textcol, face="italic"),
        axis.ticks=element_line(size=0.4),
        plot.background=element_blank(),
        panel.grid.major = element_blank(),
        panel.grid.minor = element_blank(),
```

```
            panel.border = element_rect(colour = "black", fill=NA, size=1),
            plot.margin=margin(0.7, 0.4, 0.1, 0.2, "cm"),
            plot.title=element_text(colour=textcol, hjust=0, size=14, face="bold")
          ) +
      ggtitle(VFspecies)
    # ggsave(p, filename=paste0(cooccurdir, VFspecies,
    ↪  "_HvA_VFvARG_cooccurrence_sig.jpg"), device = "jpeg",
           # width=40, height=21, units = "cm")
    # ggsave(p, filename=paste0(cooccurdir, VFspecies,
    ↪  "_HvA_VFvARG_cooccurrence_sig.pdf"), device = "pdf",
           # width=40, height=21, units = "cm", useDingbats=FALSE)
    p
  } else { print("ERROR: INPUT EMPTY.")}
}
```

```
## [1] "Running: Escherichia coli"
## [1] "Running: Klebsiella pneumoniae"
```

end
